# Migrations facilitated the cultural diffusion of megaliths in Neolithic Europe

**DOI:** 10.64898/2026.09.24.753699

**Authors:** Hugh McColl, Bettina Schulz Paulsson, Stephen Shennan, Rasmus Nielsen, Thomaz Pinotti, Jessie Woodbridge, Ralph Fyfe, Maria Grazia Melis, Christian Isendahl, Rita Peyroteo Stjerna, Vitale Sparacello, Rich Potter, Catarina Costeira, Eduardo Porfirio, Bente Philippsen, Ana Maria Silva, António Faustino Carvalho, Pétrequin Pierre, Cassen Serge, Guus Kroonen, Audrey Blanchard, Jean-Noël Guyodo, Anna Bini, Jonas Paulsson, Victoria Dahl, José Moita, Marcus Hjorth, Andreas Bak Pørksen, Jesper Stenderup, Gaunitz Charleen, Lasse Vinner, Martin Sikora, Morten Allentoft, Kristian Kristiansen, Eske Willerslev

**Affiliations:** The Lundbeck Foundation GeoGenetics Centre, Globe Institute, University of Copenhagen, Copenhagen, Denmark; Centre for Ancient Environmental Genomics, Globe Institute, University of Copenhagen, Copenhagen, Denmark; Department of Historical Studies, University of Gothenburg, Gothenburg, Sweden; Institute of Archaeology, University College London; Departments of Integrative Biology and Statistics, UC Berkeley, USA; Laboratório de Biodiversidade e Evolução Molecular (LBEM), Universidade Federal de Minas Gerais, Belo Horizonte, Brazil; School of Geography, Earth and Environmental Sciences, University of Plymouth, UK; Department of History, Human Sciences and Education, Laboratory of Prehistory and Experimental Archaeology (LaPArS), University of Sassari, Italy; Human Evolution, Department of Organismal Biology, Uppsala University, Sweden; UNIARQ - Centre for Archaeology, School of Arts and Humanities, University of Lisbon, Portugal; Department of Life and Environmental Sciences, University of Cagliari, Cagliari, Italy; CEAACP - Center for Studies in Archeology, Arts and Heritage Sciences, Portugal; National Laboratory for Age Determination, NTNU University Museum, Norwegian University of Science and Technology. Sem Sælands vei 5, 7034 Trondheim, Norway; CIAS, Department of Life Sciences, University of Coimbra, Portugal; CEF, Department of Life Sciences, University of Coimbra, Portugal; Universidade do Algarve, FCHS, Campus de Gambelas, 8000-117 Faro, Portugal; ICArEHB - Interdisciplinary Center for Archaeology and Evolution of Human Behaviour; MSHE C.-N. Ledoux, Université Marie et Louis Pasteur, CNRS, Besançon, France; Nantes Université, Laboratoire archéologie et architectures, France; Leiden University Centre for Linguistics, Leiden University, Leiden, The Netherlands; Department of Nordic Studies and Linguistics, University of Copenhagen, Copenhagen, Denmark; Archeodunum SAS, France; Museo Geologico, Lisbon, Portugal; Trace and Environmental DNA (TrEnD) Laboratory, School of Molecular and Life Sciences, Curtin University, Perth, WA, Australia; Section for GeoGenetics, Globe Institute, University of Copenhagen, Copenhagen, Denmark; Department of Genetics, University of Cambridge, Cambridge, UK

## Abstract

Around 6700 BP (years before present), Neolithic Europe saw the emergence of megalithic monuments in northern France, followed by a rapid spread to the rest of the continent^1^. However, whether this was driven by the transmission of ideas (cultural diffusion) or the movement of people remains debated. To test whether this spread coincided with population movements, we reconstructed the population history of the European Neolithic (∼6500–4800 BP) by analysing genomic data from 2963 ancient individuals, including 46 unpublished genomes from megalithic and related contexts. To assess the alternative scenario whereby megaliths spread through contact networks, as expected under cultural diffusion, we integrated archaeological evidence with radiocarbon chronologies to reconstruct past trading networks. We identify northern France, where megaliths first emerged, as a zone of genetic admixture and cultural influence from the two major early Neolithic divisions: the Cardial of the Western Mediterranean and the Linear Pottery traditions of central Europe. From this admixed population, we uncover several migrations associated with the formation of the Funnel Beaker Culture in north-central Europe and the neolithisation of Scandinavia and the British Isles. We find that megaliths were spread exclusively within regions impacted by Cardial-related genetic ancestry, but only in the British Isles and Ireland did their emergence coincide with the time of the migration. The temporal delay between the migrations and the spread of megaliths elsewhere reveals that neither population movements nor cultural diffusion alone fully explains the geospatial distribution of the megalithic tradition. Rather, megaliths diffused across different Neolithic groups whose long-distance trade networks formed in the wake of earlier population movements. Our findings highlight the need to move beyond simplistic interpretations that present migrations or cultural diffusion as isolated events bringing technological and cultural innovations.

## Main

The causal mechanisms explaining the distribution of material culture, technology, art, religion, and language in prehistoric Europe have been intensely debated for over a century. During the first half of the 20th century, archaeologists tended to attribute the spread of culture to migration. However, in the post-war period, the assumptions of migrationism were increasingly challenged, in part due to their association with nationalistic and racialised models of prehistory^2^. By the end of the century, migrationism had been largely replaced by cultural diffusionism - the transmission of ideas without large-scale human movement. Recently, ancient population genomics, in combination with isotope^3,4^ and radiocarbon dates^5^, have reignited these debates by tracing fine-scale human mobility across time and space. While attempts have been made over the last decade to couple demographic movements interpreted from ancient DNA evidence with the spread of material culture and language (e.g. ^6–11^), these correlations have been criticised for being overly simplistic^12^.

The megaliths of Neolithic Europe are central to the debate on human migration versus cultural diffusion. More than 35,000 of these stone monumental structures are distributed across the coastal regions of western Europe, in the form of megalithic tombs or standing stones. They are often arranged in specific alignments or circles, as is the case for Carnac and Stonehenge, and their use as tombs represents one of the earliest examples of a widespread ritualistic burial practice. Explanations for their expanding distribution over time have historically mirrored shifting archaeological paradigms. Early diffusionist models suggested that megalith architecture and burial rites spread from the Near East or a Mediterranean point of origin to northwest Europe via migrations or small groups of missionaries^13^. Contrasting processual perspectives later argued for independent centres of origin among a series of sedentary farming communities^14^ (Supplementary Note S1.1). Most recently, radiocarbon modelling identified a single origin of megaliths around 6700 BP in Northwest France (Brittany), followed by two phases of expansion of megalithic tombs over coastal regions^1^, reviving models that incorporate population movements. After their emergence in the form of dolmens, the early megalithic tombs initially spread southward via maritime routes along the Atlantic and Western Mediterranean coasts (Fig. 1c, e, Extended Data Fig. 1c). A second phase began ∼6300 BP with the appearance of megalithic passage graves in western France, a larger grave form associated with new burial practices (Fig. 1d, e, Extended Data Fig. 1c). This phase saw a rapid expansion of megalith construction along the coasts, reaching the British Isles ∼5900 BP and later northern Europe by ∼5600 BP. Despite being associated with different regionally defined Neolithic cultures, megalith architecture is highly homogeneous (Supplementary Note S1.1, S2.2-S2.10).

**Fig. 1.**
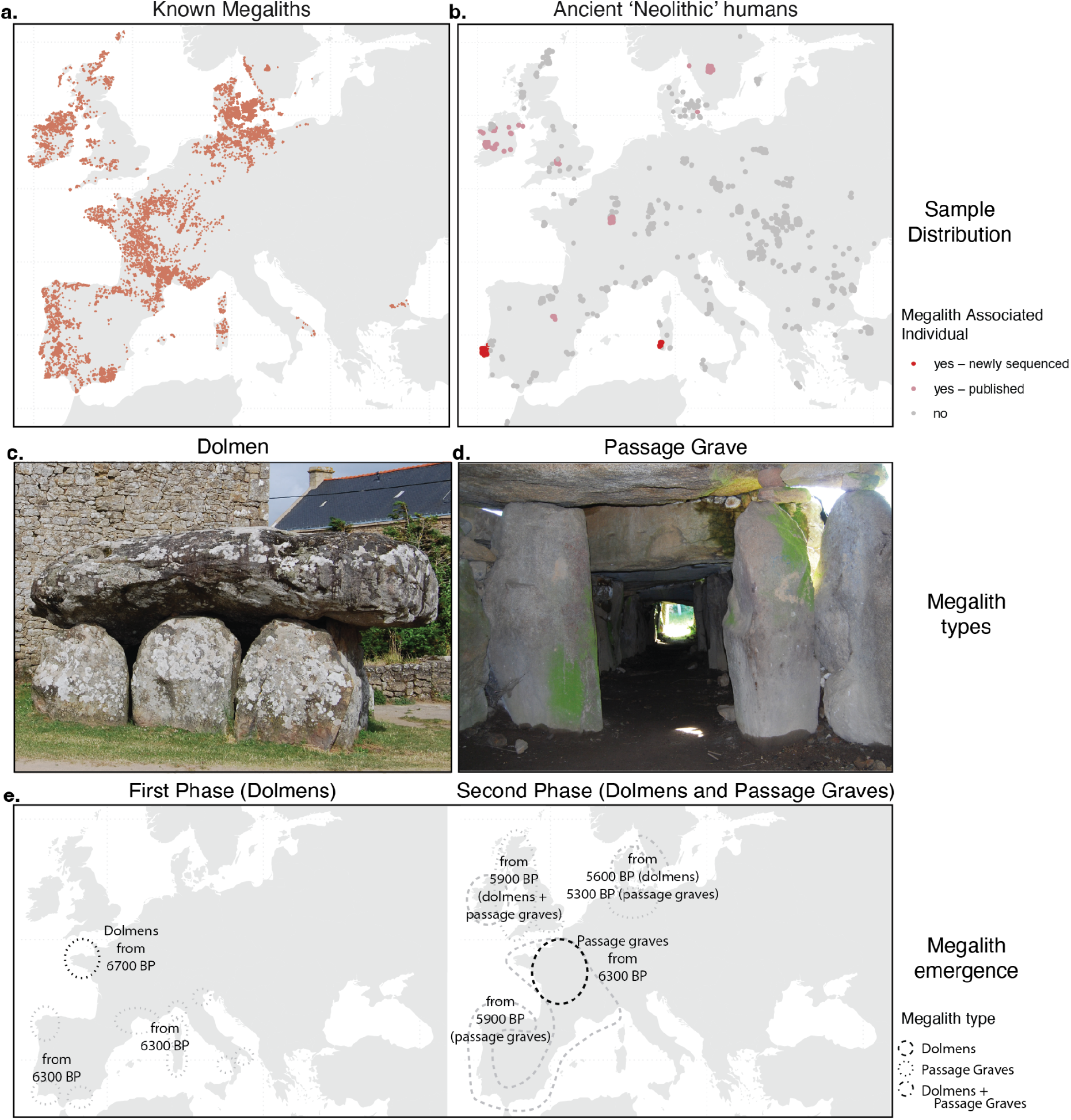
Geographic distributions of megaliths and Neolithic populations, alongside examples of the two major megalith types. **(a)** The geographic distribution of known megaliths^15^. **(b)** The geographic distribution of the 952 ancient individuals (among the 2963 individuals analysed in this study) showing >=50% Neolithic farming ancestry (mixture modelling set 1) older than 4800 BP. Newly sequenced samples associated with megaliths and related grave forms are coloured red. Published genomes associated with megaliths^16–22^ are coloured dark pink. **(c)** A dolmen (Dolmen de Plouhinec II, Brittany), displaying a simple chamber with capstone. **(d)** A passage grave (Mané Ruetual, Brittany) showing an extended passage leading to a large chamber. **(e)** The emergence and geographic extent of the first and second phases of megalithic expansion^1^.

An explicit test of the two competing hypotheses is possible, as they would leave distinct demographic patterns. The spread of megalithic traditions with a migrating population would be evidenced by observing the emergence of megaliths coinciding with a population movement from a region where the specific type of megaliths was already present. Conversely, the spread of megalithic structures through cultural diffusion alone would produce no genetic signal of population migration or admixture but require contact and communication between populations to facilitate the passing of the highly technical construction ability. To evaluate which hypothesis explains the emergence of megaliths, a continent-wide overview of changes in demographic structure and contact networks is required.

The demographic structure across Neolithic Europe prior to the emergence of megaliths is well understood. Following its origin and expansion from the Middle East, farming practices expanded within the European continent during the Early Neolithic, along two genetically and culturally distinct trajectories reflecting separate migratory routes^19,23–28^. The northern branch reached northern Germany around 7400 BP, associated with the inland Linear Pottery Culture (Linearbandkeramik, LBK) (Extended Data Fig. 1a), whose origins have archaeologically been traced to the Starčevo-Körös culture of southeast Europe. The southern branch followed a maritime route along the northern Mediterranean coast, reaching the Iberian Atlantic coast around 7500 BP^29^ via the Impressed Ware (*Impressa*) and subsequent Cardial traditions. By contrast, the demographic structure and dynamics during the millennium following the first megaliths (6700 BP) remain unclear. Some ancient DNA studies suggest that people of the Post-LBK Funnelbeaker Culture (*Trichterbecherkultur*, TRB) of Scandinavia (from ∼6100 BP) were derived from the earlier, northern LBK-related populations^28^, and the contemporaneous Neolithic populations of the British Isles and Ireland from the southern Cardial-related lineages^19,24,25^. Conflictingly, subtle genetic links between Neolithic individuals from Scandinavia and the British Isles and Ireland have been detected^19,24^, potentially linked to their shared megalithic traditions. Understanding the extent to which these populations were related and whether the suggested links are temporally consistent with the emergence of megaliths is required to uncover the underlying drivers of the spread.

Just as population movements can be traced genetically, the prehistoric contact spheres that potentially facilitated the cultural diffusion of megalith building expertise can be traced archaeologically. Indeed, extensive archaeological research has revealed a series of changes in the long-distance exchange networks connecting Neolithic Europe. For example, an expanding influence from southwest Europe is apparent from the introduction of agriculture into southern Scandinavia, including varieties of naked wheat and barley (Extended Data Fig. 1c) traced back to the Western Mediterranean around 7000 BP^30,31^, and the arrival of Alpine jadeite axes in northern Europe and the British Isles by 6000 BP^32^, that were previously only circulated in the southwest (Extended Data Fig. 1b, Supplementary Note S2.11). However, these studies tend to focus on specific regions or artefact types, necessitating a wider systematic review of Neolithic contact spheres.

To address the extent to which the emergence of megaliths fits with the spread of people, we first reconstruct the population history of Neolithic Europe by analysing 2917 published ancient individuals dating from 45000 to 2800 BP. As megalith-associated burials are primarily from western and northern Europe, we also include 46 newly sequenced genomes, all of which represent individuals from megaliths and associated burials in Portugal and Sardinia (Fig. 1b). Further, to evaluate the role of prehistoric contact networks, we reconstruct spheres of connectivity across Neolithic Europe by integrating recent raw material provenance studies with archaeological evidence and reanalysed radiocarbon chronologies.

### The Early Neolithic in northern France

The neolithisation of northern France began around 7100 BP with the arrival of the LBK culture, which by the appearance of the first megaliths (∼6700 BP) had diverged into a series of distinct ‘Post-LBK’ cultures. Although there is no evidence of Cardial pottery in northern France, there are indications of some influence from the Mediterranean region, including the appearance of naked wheat and barley by ∼6700 BP, crops previously restricted to areas further south (Extended Data Fig. 1). Consistent with these lines of evidence is genetic influence from both early Neolithic groups (LBK and Cardial) in northern France^33^. Resolving the relative contributions of these two ancestries is therefore essential for establishing the ancestral background of the first megalith builders. In addition, understanding the genetic framework of the region more broadly allows for subsequent population movements to be identified.

We first model the ancestry of early Neolithic LBK- and Cardial-associated individuals across Europe using distal source ancestry groups. This allows us to establish the genetic background of more proximal ancestry sources that contributed directly to the population of the first megalith builders. We applied supervised mixture modelling (methods) based on the sharing of Identity-by-descent (IBD) segments between all individuals in the dataset, as described elsewhere^28,34^. For every individual, we created a ‘painting palette’ by calculating the total amount of IBD sharing between that individual and all genetic clusters in the dataset, normalised by the number of individuals in each cluster. By selecting individuals from specific clusters as sources, the palettes of target individuals can be best modelled as a combination of the source palettes, resulting in admixture proportions.

The initial set of ancestry sources (Set 1) we used to model Neolithic Europeans included representatives of known groups relevant to western Eurasian prehistory, including early Anatolian Farmers, various Hunter-Gatherer groups and more distantly related outgroups (Table S3.5, Extended Data Fig. 2). As targets, we included all ancient European individuals from 12000-6300 BP (n=2909), covering the Mesolithic-Neolithic transition in the Near East and Europe. As has previously been established^35^, we observe that the transition to farming in Europe is accompanied by the arrival of ancestry from Anatolia (Extended Data Fig. 3a). Based on archaeology, the genetically distinct LBK and Cardial can be traced back to southeast European Starčevo-Körös Culture and Impressed Ware Cultures respectively^36,37^. However, this has not explicitly been tested genetically. For the next source set (Set 2), we therefore added early Neolithic individuals associated with the supposed source cultures of the two distinct Neolithic movements into Europe: the Impressed Ware Culture (Zemunica Cave, Croatia, ∼7800 BP) and Starčevo-Körös Culture (Hungary, Serbia, Romania and Croatia, ∼8000-7500 BP). We find that the distribution of Starčevo-Körös-related ancestry (Fig. 2a) corresponds closely to the LBK archaeological regions (Fig. 1a). Conversely, we find Impressed Ware-related ancestry has a close correspondence with regions archaeologically assigned to the Cardial culture. Corresponding with the mixture modelling results, there are smaller residuals for Neolithic individuals from across Europe when using the sources included in Set 2, compared to Set 1 (Extended Data Fig. 4a), indicating the additional sources improve the fit. While previous studies have shown Cardial and LBK populations were genetically distinct^19,23–28^, we now have sufficient resolution to trace the farming ancestry of the northern LBK groups and southwestern Cardial groups back to the Starčevo-Körös and Impressed Ware groups, respectively.

**Fig. 2.**
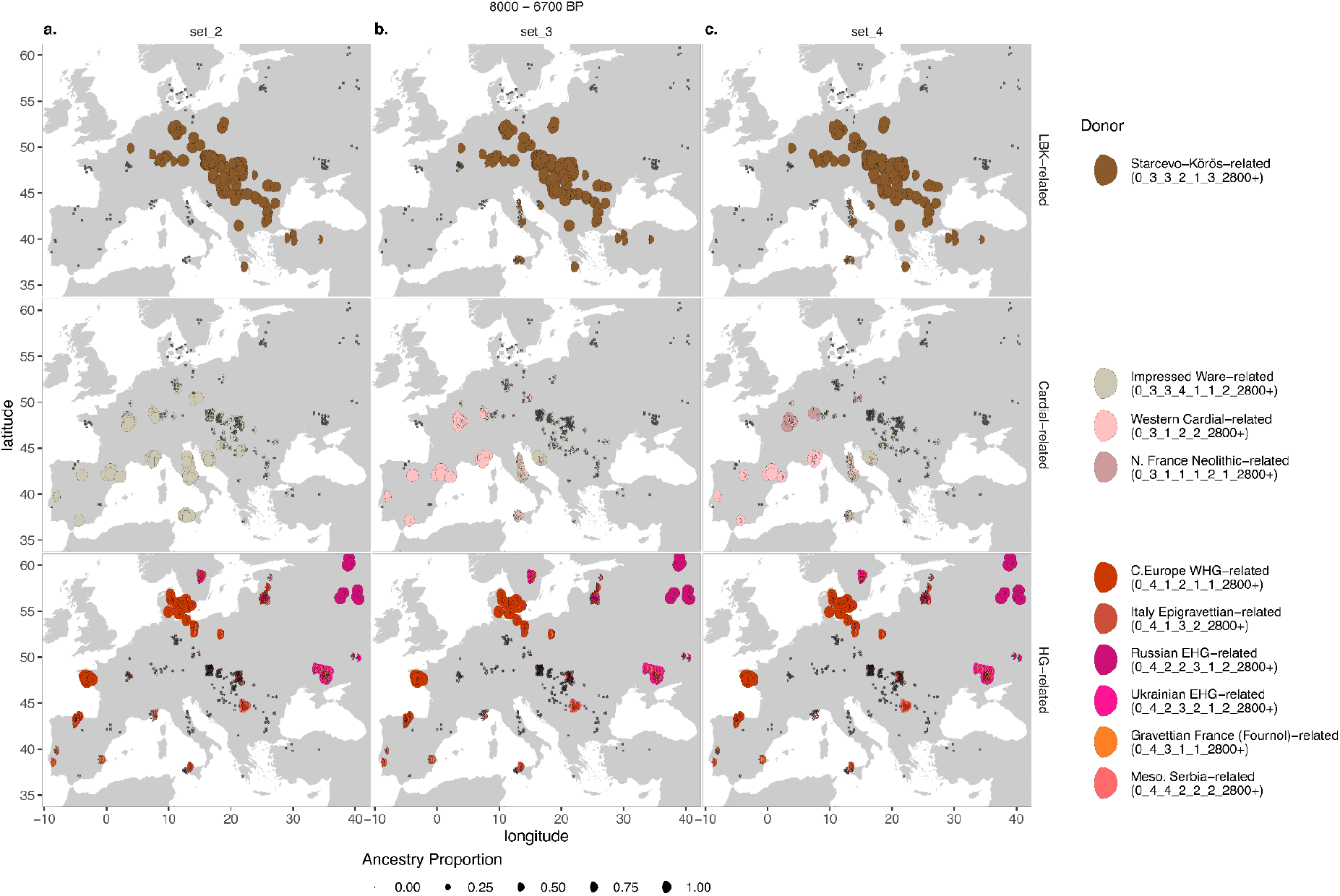
IBD mixture modelling results for all ancient Europeans dated from 8000–6300 BP for various LBK-related, Cardial-related and Hunter-Gatherer-related ancestry sources. The size of the ‘moon’ represents the proportion of ancestry modelled for each target. For each target individual, the results for three mixture modelling sets of sources (Sets 2-4) are shown. Ancestry sources are grouped either as LBK-related, Cardial-related or Hunter-Gatherer-related. **(a)** Set 2: Targets modelled with a single LBK-associated (Starčevo-Körös-related) and Cardial-associated (Impressed Ware-related) source, together with multiple Hunter-Gatherer sources. **(b)** Set 3: Same sources as Set 2, but with an additional source from Iberia (Western Cardial-related), which was previously modelled with high Cardial-related ancestry **(c)** Set 4: Same sources as Set 3, but with an additional source from Northern France (N. France-related), which was previously modelled with high Western Cardial-related ancestry. The location of all ancient target individuals is represented by an ‘x’. The full set of mixture modelling results (sets 1-6) for all individuals are plotted as bar charts in Supplementary Fig. S3.16

By modelling the northern French individuals directly using LBK- and Cardial-related sources, rather than using admixed Iberian hunter-gatherer ancestry as a proxy for Mediterranean influence^33^, we can quantify the relative contribution of each Neolithic source. We note that Cardial-related ancestry and LBK-related (Starčevo-Körös-related) ancestry meet at a boundary in Early Neolithic northern France (Fig. 2a). To identify the origin of the Cardial-related ancestry in northern France more precisely, we included a second Cardial-related ancestry source from Iberia (Western Cardial-related) in the modelling. With both Cardial sources included (Western Cardial from Iberia and Impressed Ware from Croatia, see Set 3), we find a clear geographical distinction in the targets from 8000–6300 BP: individuals from Italy and around the Adriatic Sea are modelled with Impressed Ware-related ancestry, while Cardial-related individuals from Iberia and France are now modelled with Western Cardial-related ancestry (Fig. 2b). Consistent with these results, we observe a decrease in residuals for target individuals from southwest Europe at this time, when the Western Cardial-related source is included (Extended Data Fig. 4b). The presence of Western Cardial ancestry in France suggests an influx of ancestry to the region via Iberia or southern France, rather than from Italy. The LBK-related ancestry stream in northern France was observed in three individuals (7100–7016 BP) who were modelled primarily as being Starčevo-Körös-related, consistent with their archaeological assignment (Set 3, Extended Data Fig. 5a).

However, a fourth individual (7036 BP) whose burial is also characteristic of a local variation of the LBK culture (the Haute-Alsace Rubané Culture)^33^ was modelled primarily with Western Cardial-related ancestry. This individual being modelled with ancestry from Iberia is consistent with inferences based on the presence of some Goyet2 Hunter-Gatherer ancestry^33^, a genetic signature of Iberian Neolithic groups. This discordance between ancestry and burial could suggest the adoption of foreign cultural practices by Western Cardial-related individuals. The genetic and cultural blending of these groups in northern France thus sets the stage for the subsequent emergence of megaliths in the region.

### The first megaliths in Northwestern France

The Post-LBK period in northern France saw the emergence of the first monumental graves around 6700 BP. In the Paris Basin, Cerny culture communities built elongated earthen tumuli known as ‘Passy’ graves (Supplementary Note S2.4), while to the west in Brittany, the first megalithic chambers emerged within similar monumental graves, probably as part of the same broader phenomenon^1^ (Supplementary Note S2.4.2-S2.4.3). Further northeast in France, a contemporaneous formation of other Post-LBK cultures not associated with monumental graves occurred: Großgartach, Planig-Friedberg and Rössen, and the subsequent Michelsberg (Extended Data Fig. 1a). These groups derive cultural and technological elements from both the Cardial and LBK groups, including agricultural practices from the south, as well as certain ornaments and pottery styles previously associated with the more northern LBK culture^38^. By applying the high-resolution mixture modelling method on published genomes from this region, we estimate the ancestry of these groups and test if the apparent cultural integration of Cardial and LBK elements was accompanied by genetic admixture.

From the Cerny Culture, we analysed 32 genomes of unrelated individuals from Fleury-sur-Orne, Gurgy and Buchères in the Paris Basin. While traditionally viewed as a culture derived from the LBK via the Villeneuve-St.Germain-Blicquy Culture^39^, our analyses reveal their ancestry to be modelled primarily as Western Cardial-related (Fig. 2b), consistent with previous genetic studies showing an affinity to the south^22,33^. Despite their Cardial ancestry, all Cerny individuals from the Paris basin are modelled with small proportions of LBK-related ancestry (Set 3, Extended Data Fig. 5a) providing a possible genetic vector for the spread of LBK cultural elements into this population.

Although the Cerny Culture-associated individuals cover nearly the entire Paris Basin, we found a relatively uniform ancestry profile across all 32 individuals (Set 3, Extended Data Fig. 5a). In contrast, the other Post-LBK individuals from northeast France, associated with the Großgartach, Planig-Friedberg and Rössen Cultures, show highly variable proportions of Western Cardial- and LBK-associated Starčevo-Körös-related ancestry. Despite their primary cultural association with the earlier LBK culture across the Paris Basin and northeast France, these groups are clearly a result of admixture between the two major lineages of ancestry. In contrast to northern France, we see the ancestry of individuals from southern France and Iberia continues to be modelled as Western Cardial-related throughout the Neolithic (Set 3, Extended Data Fig. 5a, b). We thus conclude that while the people of LBK culture had a lasting genetic and cultural impact across northern France, we see limited evidence of genetic and cultural impact further south.

Having established that the admixed people of the Paris Basin in northern France from 6700 BP are genetically distinct from southern France and Iberia, we can now address to what extent movements occurred during the two phases of megalithic spread. To test this directly, we added to the previous set of sources (Set 3) two early Cerny individuals (∼6700 BP) as a ‘N. France Neolithic-related’ source (Set 4). In northern France until 6300 BP, we find that the ancestry modelled as Western Cardial-related in Set 3 is now primarily modelled by the N. France Neolithic-related ancestry in Set 4 (Fig. 2c, Fig. 3a, Extended Data Fig. 5a). Ancestry of individuals associated with the Cerny Culture is modelled almost exclusively as N. France Neolithic-related, whereas the ancestry of the other Post-LBK groups is complemented by additional LBK-related ancestry. We note that even the earliest individual with Cardial-related ancestry in set 3 (Mor6, 7036 BP, Extended Data Fig. 5a) is modelled entirely by the N. France Neolithic-related ancestry in set 4, suggestive of genetic continuity from at least 7000 to 6300 BP in northern France. By contrast, further south, in Iberia and southern France, ancestry remains primarily modelled as Western Cardial-related. Thus, across the broad region where Cardial-related ancestry is found, we can distinguish the eastern Mediterranean (‘Impressed Ware-related ancestry’) from the Western Mediterranean (‘Western Cardial-related ancestry’) from northern France (‘N. France Neolithic-related ancestry’), allowing later migrations between the regions to be detected. The persistence of these ancestries in their respective regions indicates that no substantial secondary movement between the regions occurred at the time of the emergence of megaliths (6700-6300 BP).

**Fig. 3.**
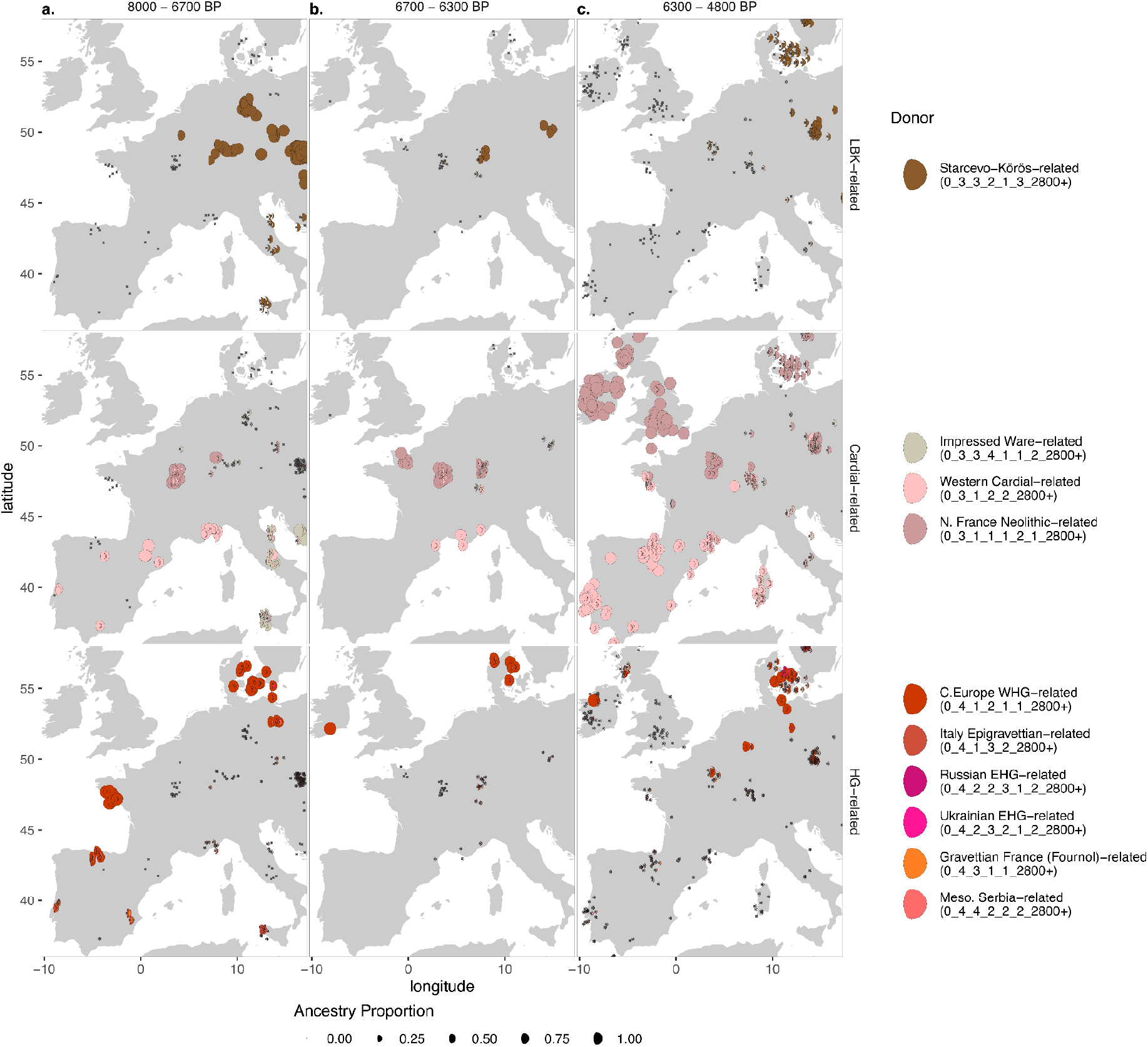
IBD mixture modelling results for all ancient Europeans dated from 8000–4800 BP for Set 4, binned by time. The size of the ‘moon’ represents the proportion of ancestry modelled for each target. **(a)** Targets from 8000–6300 BP. **(b)** Targets from 6300–5700 BP. **(c)** Targets from 5700–4800 BP. The full set of mixture modelling results (sets 1-6) for all individuals are plotted as bar charts in Supplementary Fig. S3.16

In Brittany, monumental earthen tombs associated with stone alignments emerged contemporaneously with the pre-megalithic burials of the Paris Basin (∼6700 BP), marking the transition from the Villeneuve-Saint-Germain Culture to the Neolithic Castellic Culture. Small dolmens, representing the earliest megalithic graves in Europe, subsequently appeared within these earthen monuments (Supplementary Note S2.4.2). It has been suggested that the stone alignments and early monumental graves are rooted in Mesolithic traditions^40,41^, and ancient genomes from ∼7000 BP have indeed revealed that Mesolithic ancestry persisted in Brittany alongside the Neolithic ancestry in northern France until at least ∼6900 BP^34^. However, poor preservation of skeletal remains^43^ means no genomic data are available from this time until ∼5800 BP, by which point both N. France Neolithic- and Western Cardial-related ancestries are present (Set 2, Extended Data Fig. 5a). The emergence of monumental tumuli in Brittany at the same time as the Paris Basin (∼6700 BP), where N. France Neolithic-related ancestry is widespread, suggests this ancestry also may have been present in Brittany at this time. Similarly, the subsequent emergence of passage graves and a new material culture with links to the south (∼6300 BP, Supplementary Note S2.4.3) may coincide with the arrival of the Western Cardial-related ancestry. However, testing these hypotheses requires ancient genomes spanning this entire period.

### Major Transitions and cascading effects from 6300 BP

The second phase of megalith building saw the emergence and spread of passage graves (∼6300 BP) (Fig. 1), coinciding with a major reshaping of Neolithic Europe (∼6000 BP). New rituals and technologies spread from northern France across much of Europe^1^, during a period of climatic cooling evidenced in some palaeoclimate records^44^. This period also saw the transition from hunter-gathering to farming in regions also impacted by the second phase of expansion: Scandinavia, the British Isles and Ireland. While the neolithisation of these regions has been the focus of previous genetic studies^25,28,45–47^, we are now able to explore the formation of these populations within the newly discovered genetic structure.

It is during the second phase of megalithic expansion around 5900 BP that megaliths emerged in the British Isles and Ireland. It is well established that the neolithisation of the British Isles and Ireland around 6100-6000 BP saw a demographic shift to Neolithic Farming ancestry^19,48^. However, it remains debated whether the migrations had origins in northern France, where passage graves first emerged, or elsewhere along the Atlantic coast^49,50^. Mixture modelling Set 4, which includes the ancestry source groups from northern France (‘N. France Neolithic-related’) and Iberia/southern France (‘Western Cardial-related’), allows for migrations from the two regions to be distinguished (Fig. 3). We find the ancestry in all British Isles and Irish individuals to be modelled by the N. France Neolithic-related source and additional Hunter-Gatherer ancestry, with little or no evidence of Western Cardial-related ancestry. Accordingly, we note a decrease in residuals in individuals from this region when the N. France Neolithic-related source is included (Extended Data Fig. 4c). This suggests that the ancestry found across the Paris Basin is a better proxy for migrations contributing genetically to Neolithic farmers of the British Isles and Ireland than the Western Cardial-related ancestry of southern France and Iberia. As discussed above, Brittany cannot yet be excluded as a potential source region, as the ancestry of individuals from this region between ∼6900 and ∼5800 remains unknown.

At the same time as farming reached the British Isles and Ireland, it also reached Scandinavia^51^, a region also impacted by the second phase of megalith building. Archaeologically, the expansion into northern Europe was associated with the TRB Culture, which plausibly emerged around 6100-6000 BP from the Michelsberg Culture of northeast France and western Germany^52^, rapidly spreading as far as Scandinavia, Poland and the Czech Republic. To explore the timing of potential genetic shifts associated with the appearance of TRB material culture, we plot the mixture modelling results from Set 4 as timelines (Fig. 4). We focus here on the Czech Republic, where dense sampling exists prior to and throughout the TRB period. Between ∼7200 and ∼6100 BP, almost all individuals are modelled primarily as LBK-associated Starčevo-Körös-related ancestry. However, from ∼6100 to ∼5400 BP, corresponding to the TRB period, almost all individuals are modelled with roughly half LBK-related ancestry and half N. France Neolithic-related ancestry (Fig. 4). Whether this transition represents an influx of unadmixed N. France Neolithic-related ancestry, or a more complete turnover in already-admixed form is unclear. In Scandinavia, however, where TRB represents the earliest Neolithic culture, we find all TRB Scandinavian individuals are also modelled with similar proportions of LBK-related and N. France Neolithic-related ancestry (Fig. 4). As neither of these Neolithic ancestries are found in Scandinavia in unadmixed form, it is likely that the admixture occurred prior to the neolithisation of Scandinavia. A PCA of all western Eurasian individuals predominantly of Mesolithic and Neolithic ancestry shows TRB-associated individuals in a cluster between LBK- and Cardial-associated individuals, consistent with them being modelled as an admixed group (Supplementary Note S3.3.1, Supplementary Fig. S3.7). In addition, the presence of some N. France Neolithic-related ancestry, which itself is predominantly Western Cardial-related derived, explains the previously confounding results showing Neolithic populations of Scandinavia having genetic links with the Cardial-related populations of the Neolithic British Isles and Ireland^19,25^ but also with the LBK-related populations of Central Europe^28^.

**Fig. 4.**
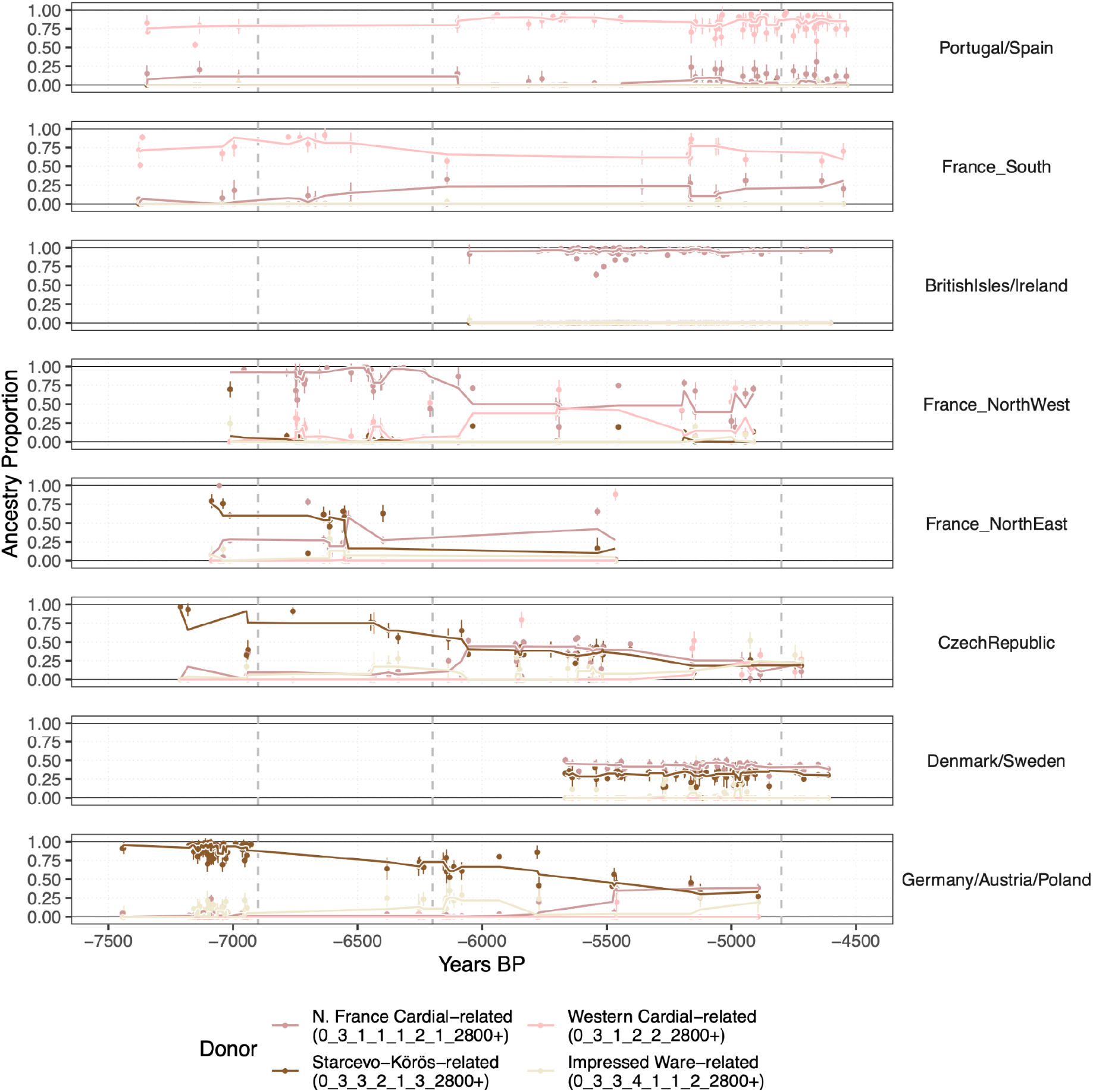
Mixture modelling Proportions for European Neolithic Farming Related ancestry sources (Set 4) from Iberia, western, central and northern Europe. For each individual, the proportion of each of the four Neolithic ancestries modelled is plotted. To visualise changing trends through time, a trendline (rolling median with a window of 5) for each ancestry is plotted. The full set of mixture modelling results (sets 1-6) for all individuals are plotted as bar charts in Supplementary Fig. S3.16

While the similar proportions of the N. France- and LBK-related ancestries across the TRB region are suggestive of a single admixture event associated with the formation of the TRB Culture, we can test how uniform they are by including TRB individuals as a source. We included individuals from the Czech Republic as the additional source (‘TRB-related’) in Set 5, representing some of the earliest TRB-associated individuals in the dataset. We note that a target individual modelled with Czech source ancestry should not be inferred as a migration from the Czech Republic. The result instead indicates that the source and target are genetically indistinguishable relative to the other sources in the set, possibly resulting from a migration in either direction, or from a third location into both. Consistent with the TRB populations deriving from a single admixture event, we find this ancestry modelled in high proportions across the regions archaeologically associated with the TRB culture, including Germany, Scandinavia, the Czech Republic, Poland and Ukraine, with only minor local contributions (Extended Data Fig. 6). This result is further supported by residuals improving in Scandinavia when including the Czech TRB source (Set 4 to Set 5, Extended Data Fig. 4), indicating that the mixture modelling proportions are not the result of overfitting. Thus, individuals from across the entire TRB range can trace part of their ancestry back to northern France, where megaliths emerged.

It was not only Scandinavia, the British Isles and Ireland that were impacted by the second phase of megalith building, but also western Iberia (5900 BP). Unlike the simultaneous movements to the north, we find that our newly sequenced genomes from western Iberian individuals associated with megalithic graves were primarily of local Western Cardial-related ancestry (Fig. 4), rather than the N. France Neolithic-related ancestry. Thus, we see the second phase of the spread of megaliths impacting people of distinct ancestry profiles - Western Cardial-related ancestry in western Iberia, N. France Neolithic-related ancestry in the British Isles and Ireland, and TRB-related ancestry in Scandinavia. Despite this distinction, all groups can trace some of their ancestry back to the Western Mediterranean: TRB-related ancestry is largely derived from N. France Neolithic-related ancestry, which in turn is primarily derived from Western Cardial-related ancestry. It should also be noted that while megaliths emerge exclusively in regions impacted by the expansion of Cardial-related ancestry, they do not appear across the entire range in which the ancestry is found but instead are found initially in coastal regions.

To formally test the association between changes in the geographical distribution of all ancestry components and the emergence of megalithic monuments, we used a discrete-time survival model, treating megalith emergence as a region-level, irreversible (absorbing) event (methods; Supplementary Note S3.3.2). We found the first significant association in mixture modelling set 3 (FWER-corrected P-value = 0.038), with Western Cardial-related ancestry being strongly associated with the emergence of megaliths (Supplementary Note S3.3.2), aligning with the observation of this ancestry spreading to all regions that megaliths are found. In sets 4-6, the N. France Neolithic ancestry is the most strongly associated ancestry (FWER-corrected P-values = 0.006, 0.010, 0.009). Consistent with the association shown for Set 3, we note that this ancestry is primarily derived from Western Cardial-related ancestry. While N. France Neolithic-related ancestry is the primary ancestry in megalith building regions of northern France, Britain, Ireland, and Scandinavia, it does not explain the southward spread of megaliths to Iberia where Western Cardial-related ancestry remains dominant.

After 6,300 BP, the geographic distribution of Y-chromosome lineages associated with Cardial and LBK ancestry also shifted, with Cardial-associated lineages expanding into the British Isles, Scandinavia and Eastern Europe, whereas the distribution and frequency of LBK-associated lineages declined (Extended Data Fig. 6). We defined Y-chromosome lineages as Mesolithic, Cardial- or LBK-associated based on their occurrence among multiple individuals carrying the corresponding autosomal ancestry profiles and their age (Supplementary Note S3.3.3; Methods). The persistence of LBK-related autosomal ancestry in TRB populations and apparent absence of LBK-associated Y-chromosome lineages could indicate a sex-biased contribution to the formation of these populations. This contrasts with the earlier Neolithic migrations, when multiple Y-chromosome lineages associated with European Mesolithic groups were incorporated into populations carrying Neolithic farmer-related autosomal ancestry.

### The spread of megaliths by Neolithic seafarers

While the spread of megaliths across Europe has been linked to seafaring^1^, the first evidence of Neolithic seafaring can be traced further back to the initial Cardial expansions. In the Mediterranean, early Neolithic seafaring was developed by Cardial groups, as reflected in the coastal spread of farming, obsidian exchange, and shared Cardial-related ancestry in Iberia and Morocco^53^. Similarly, after 6300 BP, megaliths spread along maritime routes across the Western Mediterranean coast and the central Mediterranean islands reaching Corsica by 6250 BP and Sardinia between 6700-5900 BP^1^ (Supplementary Note S2.2.3). The spread of megaliths throughout the Mediterranean, over a thousand years after the initial neolithisation of the region, presents two possible scenarios: cultural diffusion between populations who trace their ancestry back to the initial Cardial migrations, or a spread facilitated by secondary seafaring migrations.

With the newly established genetic framework distinguishing northern France from the Western Mediterranean, it is possible to explore whether migrations accompanied each of the phases of megalithic spread from northern France. We first set out to establish how populations in the Western Mediterranean region were related prior to the spread of megaliths. When including LBK, Croatian Impressed Ware, and Western Cardial sources (Set 3), we find individuals in the Western Mediterranean modelled primarily by Western Cardial-related ancestry, whereas individuals in Italy (∼8000-7200) are modelled with Impressed Ware-related ancestry. This genetic distinction allows us to test for movements eastwards into Italy during the spread of megaliths. Such a movement is first evident from an unadmixed outlier individual (R6^54^) from the Italian Peninsula dated to 7160 BP, who is modelled exclusively with Western Cardial ancestry, indicative of an ancestral origin from Iberia or southern France. While this single individual predates the spread of megaliths, we find all individuals from ∼5400 BP until the arrival of Steppe Ancestry in this region (∼4000 BP), are modelled with more Western Cardial-related ancestry than the early Neolithic samples (Fig. 5). The lack of samples from mainland Italy dated between 7160 and 5390 BP and precise dates for the emergence of megaliths in this region means it is not yet possible to identify whether Western Cardial ancestry became widespread at the exact time that megaliths emerged.

**Fig. 5.**
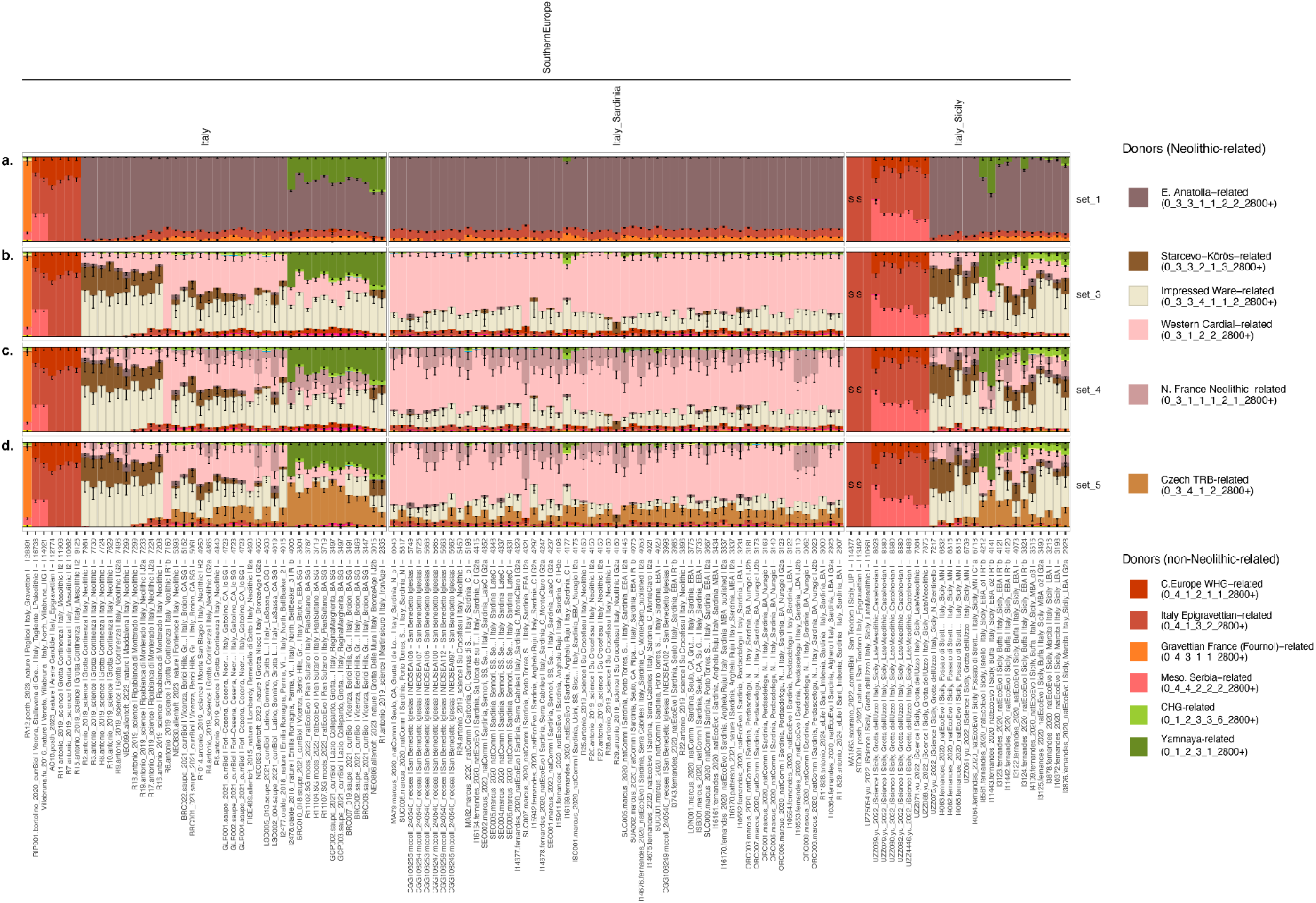
Mixture modelling results for mainland Italy, Sardinia and Sicily, highlighting (a) the transition from Hunter-Gatherer to Farming ancestry, (b) LBK and Western Cardial-related ancestry, (c) N. France Neolithic and Western Cardial, and (d) transitions when including the TRB-related source. The full set of mixture modelling results (sets 1-6) for all individuals are plotted as bar charts in Supplementary Fig. S3.16

In Sardinia, the earliest individual in the dataset (6045 BP) postdates the arrival of the first megaliths (6700–5900 BP). Like the other new genomes from Sardinia sequenced for this study, this individual derives from the hypogeum of San Benedetto, a grave form carved into the limestone and contemporaneous with the megaliths. While we lack the pre-megalithic samples required to test if a demographic shift occurred, we note a number of characteristics indicative of a migration from the mainland: 1) In Set 1 (Fig. 5a) individuals are modelled with small amounts of Fournol-related Hunter-Gatherer ancestry, typical only of individuals carrying Cardial-related ancestry from Iberia; 2) In Set 4 (Fig. 5c), many individuals are modelled with some N. France Neolithic ancestry, in higher proportions than found in Iberia, suggesting a link with northern France; and 3) in Set 5 (Fig. 5d), many individuals are modelled with some TRB-related ancestry. While suggestive of a link to continental Europe, with the current sampling it is difficult to assess exactly what continental populations and cultural groups are driving this signal. In addition, Neolithic samples from Sardinia prior to 6300 BP are required to confirm that the arrival of this ancestry coincides with the spread of megaliths. Both regions are among the few in Europe characterised by early megaliths and stone alignments (Supplementary Note S2.2). Finally, around 4200 BP, we see a subtle shift in Sardinia indicative of interactions with Bronze Age populations from the mainland and evidence of increasing TRB- (Fig. 5d) and Steppe-related ancestry (Fig. 5) from this point onwards.

The spread of megalith building expertise to islands in the Mediterranean and along the Atlantic coast would have necessitated seafaring technology, and the shift towards mainland Cardial-related ancestry coinciding with the spread of megaliths is consistent with this notion. Nevertheless, denser sampling from immediately before and after the arrival of megaliths in the western and Central Mediterranean is required to unequivocally establish this link.

### Neolithic trading networks contemporaneous to the emergence of megaliths

Under models of cultural diffusion, megaliths are assumed to have spread through exchange networks linking distinct communities. Understanding where raw materials were sourced, reworked into their later form, and ultimately deposited provides insight into how artefacts were exchanged. To reconstruct these networks, we synthesised recent raw material provenance studies with radiocarbon chronologies (see Supplementary Notes S2.1–S2.11), providing an overview of Neolithic connectivity between 8000 and 6000 BP^32,55^. The reconstruction of these networks allows for the evaluation of how changes in connectivity relate to genomic evidence for population movements and the spread of megalithic traditions.

In the Western Mediterranean, the seaborne Impressed Ware/Cardial expansion is reflected in the circulation of obsidian from Sardinia and the Tyrrhenian Islands by ∼7700–7500 BP^56,57^, while LBK groups distributed Spondylus ornaments and lithic raw materials across continental Europe by ∼7500–7000 BP^58,59^ (Fig. 6b, d). Atlantic shells in the Paris Basin and its eastern periphery further indicate connections between continental and Atlantic regions^60–62^.

**Fig. 6.**
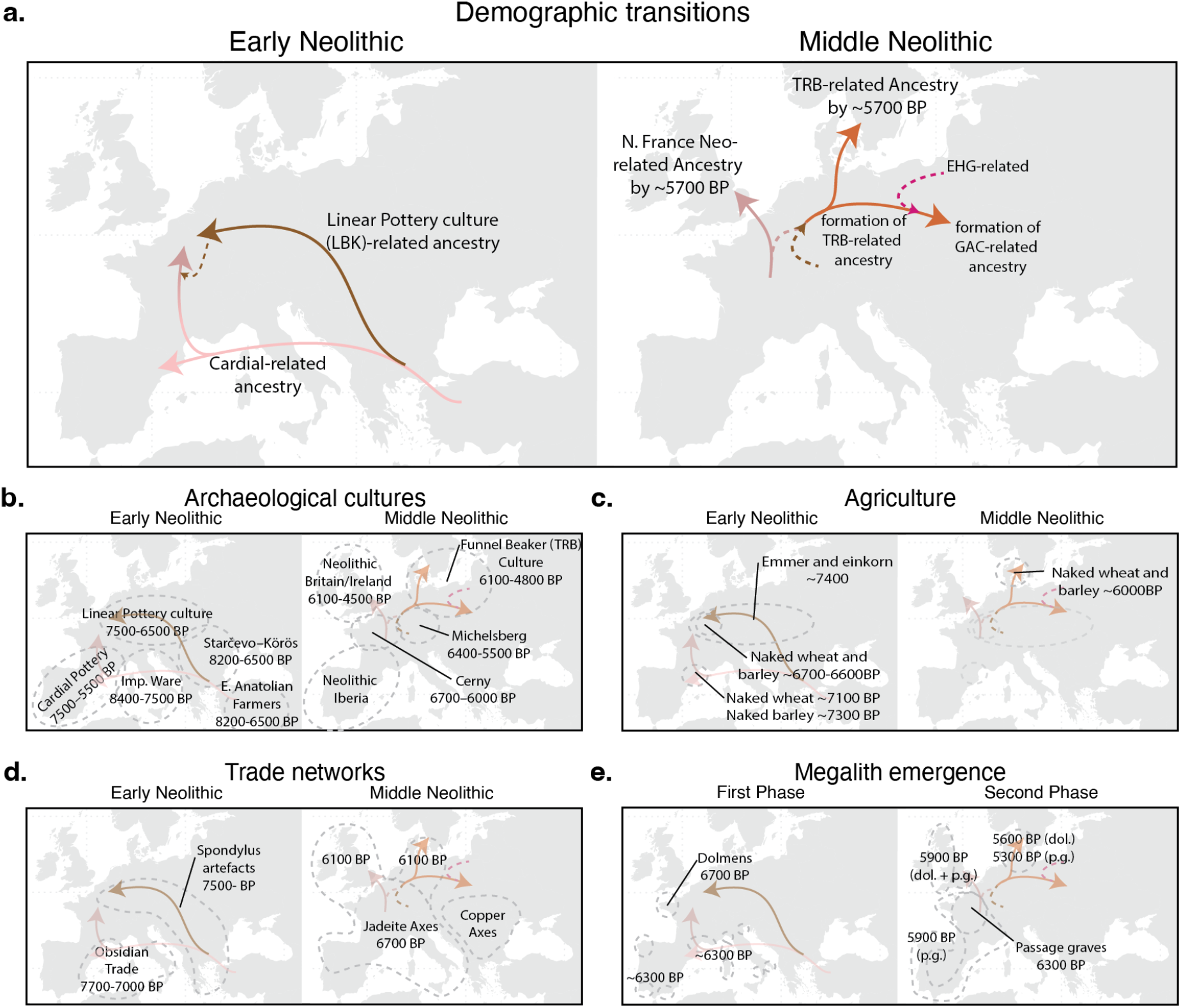
A summary of the major Neolithic demographic events discussed in this study, overlaid on established archaeological regions. **(a)** Major demographic events. **(b)** Archaeological cultures. **(c)** Agriculture. **(d)** Trade Networks. **(e)** Regional emergence of megalithic types (‘dol.’ Dolmens, ‘p.g.’ Passage Graves) across the two phases. Additional detail for panels **(b-e)** can be found in Extended Data Fig. 1.

From ∼6700 BP we see new archaeological connections linking the Mediterranean and northern France that parallel the northward Cardial expansion documented by the genetics. Provenance analyses of west Alpine jadeite and Iberian variscite ‘greenstones’ demonstrate large-scale exchange networks spanning much of western Europe connecting the Atlantic coast and the Mediterranean, with a major redistribution centre emerging in southern Brittany (Supplementary Note S2.11) (Fig. 6d). Coinciding with the construction of the earliest Breton megalithic monuments^1,32,55,63^, there is evidence of imports in northern France of Alpine jadeite from northern Italy and variscite from southwestern Iberia^1,32,55,63^. These ‘greenstone’ networks primarily followed maritime and riverine corridors. Jadeite axes reworked in Brittany reached Iberia, while engraved symbols associated with Breton megalithic societies appeared along Atlantic exchange routes^64,65^. Other raw materials also circulated within these networks, for instance, fibrolite from the French Atlantic coast to the central Iberian Peninsula^66^.

After 6300 BP interregional exchange declined along these networks: jadeite axes and variscite beads were no longer deposited as prestige grave goods in Brittany, and the quarrying of northern Italian jadeite (e.g., at Monte Viso) had ceased by ∼6300 BP and ended at the latest by 5700 BP (Supplementary Note S2.11.5). Despite this shift towards more intraregional trade, the distribution of jadeite axes expanded from the Paris Basin to southern Scandinavia and the British Isles, which continued in circulation for several centuries, until ∼5500 BP^32^ (Supplementary Note S2.11.2). In Iberia, variscite exchange shifted toward more local Galician mining sources and increasingly regional networks ^1,43^ (Supplementary Note S2.11.3). Similarly, in the British Isles after ∼6000-5800 BP, exchange networks were largely insular, reflected in megalithic distributions, shared symbolic repertoires, and, much later, in henge and Grooved Ware traditions. Scandinavia maintained exchange networks along the continental coast. Collared bottles for example, centred in southern Scandinavia and predominantly found in megalithic tombs, also occur along the North Sea and Baltic coasts to Poland and sporadically in Normandy and Brittany^67^. In the Western Mediterranean, new regional exchange systems emerged during ∼6000-5000 BP, including the circulation of Sicilian amber^68^, and a revival of earlier Mediterranean obsidian exchange networks (Supplementary Note S2.11.4).

Thus, based on the current evidence for Neolithic Europe, we find that prior to 6300 BP, major trade networks were restricted to either the southwest or northeast of Europe (Fig. 6d), mirroring the regions in which Cardial-related or LBK-related ancestry were found, respectively (Fig. 6a). It was not until 6100 BP that we see trade, in the form of Jadeite axes, extend beyond these borders to include Scandinavia and the British Isles (Fig. 6d). We see this expansion coinciding with the northern expansion of ancestry (Fig. 6a), forms of agriculture (Fig. 6c) and cultural influence associated with the formation of the TRB (Fig. 6b). While the specific traded artefacts and source regions change through time, the border between networks broadly reflects the genetic borders of the time.

## Discussion

Despite more than a century of archaeological research, the relative roles of migration and cultural diffusion in the spread of megalithic traditions have remained unresolved. The results of this study provide important new chronological and population genomic context for resolving this debate.

It is within the bounds of populations with Western Cardial-related genetic ancestry that megaliths emerge, in northern France. However, the spread of Western Cardial-related ancestry from southwestern Europe results in three distinct ancestry profiles: unadmixed in Iberia and southern France by 7300 BP, admixed with a small degree of LBK-related ancestry in N. France Neolithic-related populations by 7000 BP, and further admixed in TRB-related populations by 6000 BP (Fig. 6a). As a result of this fine-scale ancestry resolution, it is now possible to infer subsequent movements from these regions which can in turn be compared to the emergence of megaliths.

We have shown that the first monumental structures in western Europe, emerging around 6700 BP, were associated with people of N. France Neolithic-related ancestry (Extended Data Fig. 6). Following a migration model, it would therefore be expected that the first phase of the spread of megaliths would be accompanied by N. France Neolithic-related ancestry. In Iberia, which is impacted by the first (6300-6000 BP) and second (∼5900 BP) phase of megalith expansion and where dense sampling exists before and after the transition periods, we find no evidence of an increase in N. France Neolithic-related ancestry, inconsistent with migration. In Sardinia, where megaliths emerge around 6700-5900 BP, a small proportion of N. France Neolithic-related ancestry is present by at least ∼6000 BP, suggestive of a migration. However, the lack of genomes from earlier pre-megalithic burials and wide date range for the megalithic emergence means assessing when and if the transitions occurred is not possible (Extended Data Table 1). We note, however, that northern France and the Mediterranean were connected through jadeite exchange, while northern France and western Iberia were linked by variscite exchange. The emergence of megaliths during the first phase does not occur beyond the bounds of these networks, consistent with the spread of megaliths to Iberia via cultural diffusion.

The second phase of the expansion of megaliths occurred following the emergence of passage graves in northern France around 6300 BP, reaching the densely sampled regions of the British Isles and Ireland (6000-5900 BP), western Iberia (6000-5900 BP), and Scandinavia (5600 BP) (Extended Data Fig. 6). Thus, under the migration model, the emergence of megalithic graves in these regions should again be accompanied by N. France Neolithic-related ancestry. By 6100-6000 BP, we can show that people of N. France Neolithic-related ancestry colonised the British Isles and Ireland, a date that closely aligns with the emergence of megaliths in the region (6000-5900 BP). By contrast, in western Iberia where megaliths indistinguishable from those of the British Isles arrive contemporaneously (6000-5900 BP), there is no evidence of an influx of N. France Neolithic-related ancestry, as was the case for the initial phase. This suggests that their spread to western Iberia is more consistent with cultural diffusion, further supported by the existence of trade networks uniting the regions.

In Scandinavia, the alignment is less clear. Consistent with a spread facilitated by a migration, we find the TRB-related populations of Scandinavia ultimately carry ancestry from northern France, where megaliths emerged. However, the neolithisation of Scandinavia by TRB-related populations (∼6100-6000 BP) predates the arrival of dolmens (5600 BP) and passage graves (5400 BP) by hundreds of years. Prior to their emergence in northern Europe, dolmens and passage graves were found exclusively in regions in which N. France Neolithic- and Western Cardial-related ancestry were found. In Scandinavia we find no evidence of secondary influxes of these types of ancestry into the local TRB that could explain the late emergence of either type of megalith. The most obvious contradiction to megalith building expertise spreading with the people associated with the initial formation of TRB-related ancestry (6100-6000 BP) can be seen in the late first appearance of the passage graves in Scandinavia (5300 BP). This emergence represents the arrival of a specific megalith type already found in northern France, the British Isles or Western Mediterranean, rather than a local divergence from within the region of TRB-related ancestry. The chronological delay in Scandinavia, together with the highly technical and engineered nature of megalith construction, is therefore more consistent with a spread occurring later through specialised groups or networks of builders comparable to technical guilds, rather than the knowledge being introduced with the first TRB-related farming populations and then remaining dormant for centuries. We detail to what extent the British Isles, Iberia, Scandinavia and Sardinia are consistent with models of migration and cultural diffusion in Extended Data Table 1.

A simple model in which megaliths spread together with migrants is not consistent with the results presented here (Extended Data Fig. 8). However, we note that there are other cultural changes that do align with the migrations. Distinct pottery styles and farming technologies predict the spread of ancestry from northern France, along with changes in trade networks which saw the expansion of the use of Alpine jadeite axes from Iberia to Scandinavia^69^. Thus, the mid-Neolithic movements from the southwest were associated with a major reshaping of much of Europe (Fig. 6a, b, c, d), resulting in contact networks encompassing the entire region in which megaliths subsequently emerged (Fig. 6e).

The connectedness that arose in the wake of the population movements links populations from the Mediterranean to Scandinavia. Further, it provides an explanation for the intriguing emergence of megaliths exclusively within regions reached by Cardial-related ancestry, which spread northward in the form of N. France Neolithic-related ancestry. We see that the initial phase of megalith building in southwest Europe was not associated with large-scale migrations, but instead spread occurred within trade networks that formed in the wake of the initial Cardial-related migrations into the region. Despite existing in close proximity to northern France, megaliths do not initially emerge in regions where Mesolithic populations persisted (the Iberian Peninsula, British Isles, Ireland, Scandinavia, the Netherlands or northern Germany), nor do they emerge to the east, in regions where LBK-related ancestry was present. It is not until the second phase of megalith building, following the spread of ancestry from northern France, that megaliths emerge in these regions. With the exception of Britain and Ireland, the time of emergence thus tends not to be accompanied by a specific ancestry shift, but instead occurred across regions linked by long-distance trade networks: long-standing in the south, and newly formed in the north.

The lack of an associated demographic impact, combined with a scattered appearance of the two megalith types across the entire region and the emergence with insular and coastal distributions points to the spread of megalith building technology by a small number of seafaring people, rather than a large-scale migration. Thus, while the spread of Cardial-related ancestry, in various forms, was not the immediate mechanism behind the spread of megaliths in most cases, the spread of this ancestry does align with the formation of trade networks, which in turn facilitated the cultural diffusion of megaliths. It is, however, plausible that this spread of ancestry was associated with the spread of beliefs and ritualistic practices associated with Cardial and Cardial-derived populations. Under this scenario, the building of megaliths in Scandinavia, for example, would represent the adoption of a tradition practiced by their ancestors in northern France, rather than a completely foreign practice, as would be the case for people of Mesolithic or LBK cultures. While the drivers behind the population movements that shaped the later networks remain unclear, we note that 6300-6000 BP was a period of climatic instability and sea level change reshaping the coastal landscapes, particularly in northern France, that altered the distribution of land suitable for settlement and cultivation^70–75^. By contrast, the Holocene Thermal Maximum, characterised by warmer and wetter conditions, favoured the expansion of agriculture^76^ into regions such as southern Britain^77^ and southern Scandinavia^78,79^.

Finally, we note that the long-debated scenarios discussed here are simplified representations of complex dynamics underlying the spread of people and the transfer of ideologies, material culture, and technological innovations across time and space. However, the results obtained provide a much more refined demographic framework that is inconsistent with simple binary narratives. In turn, this will pave the way for more complex models to understand the intertwined roles of migrations and cultural diffusion in shaping human culture.

## Extended Data Figures and Tables

**Extended Data Fig. 1.**
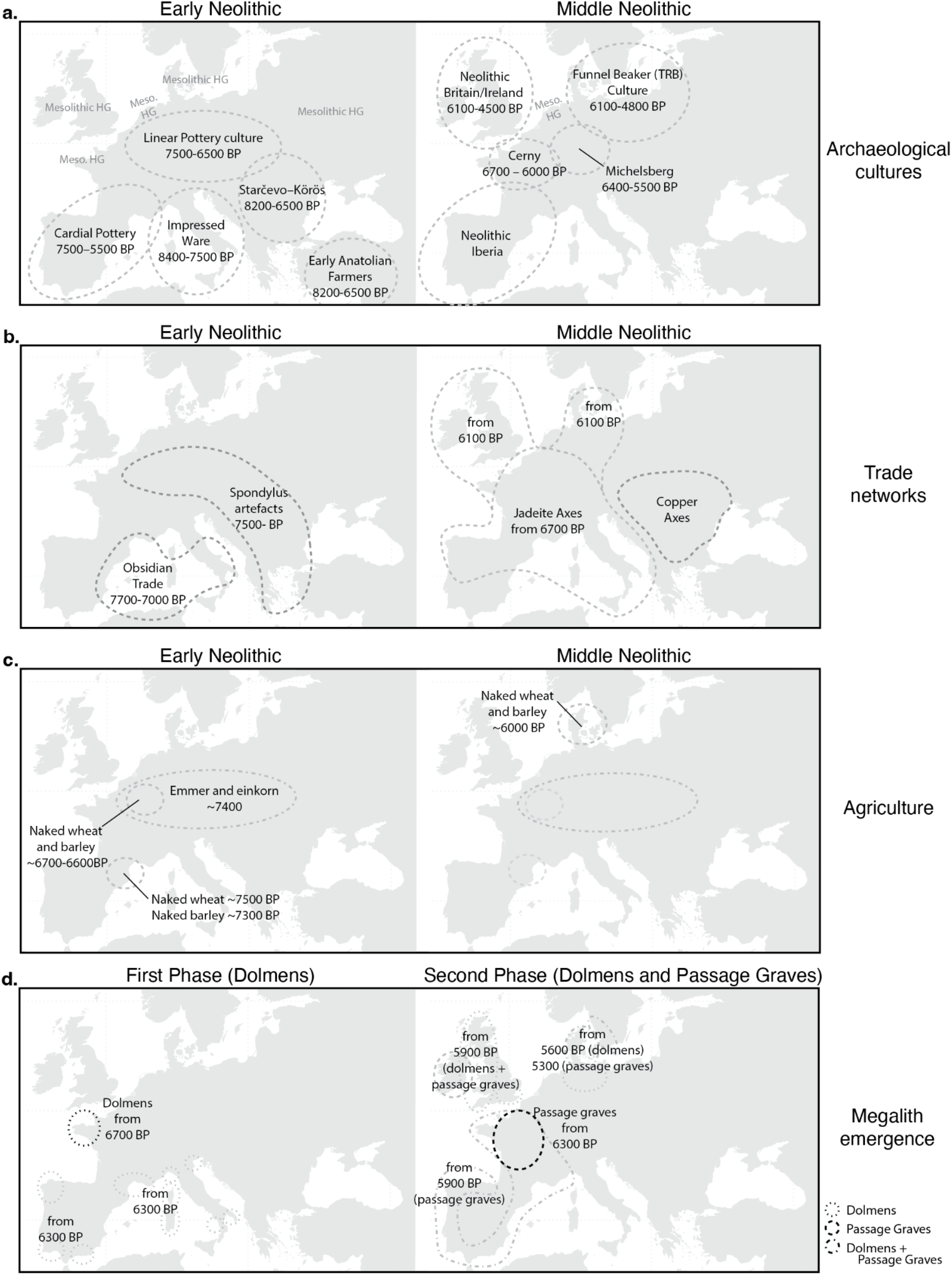
An overview of major transitions defining the early and middle Neolithic relevant to this study. **(a)** Major archaeological cultural groups. **(b)** Long-distance exchange networks. **(c)** Agriculture. **(d)** the emergence and spread of megaliths. The first phase saw the spread of dolmens. During the second phase, both dolmens and passage graves spread.

**Extended Data Fig. 2.**
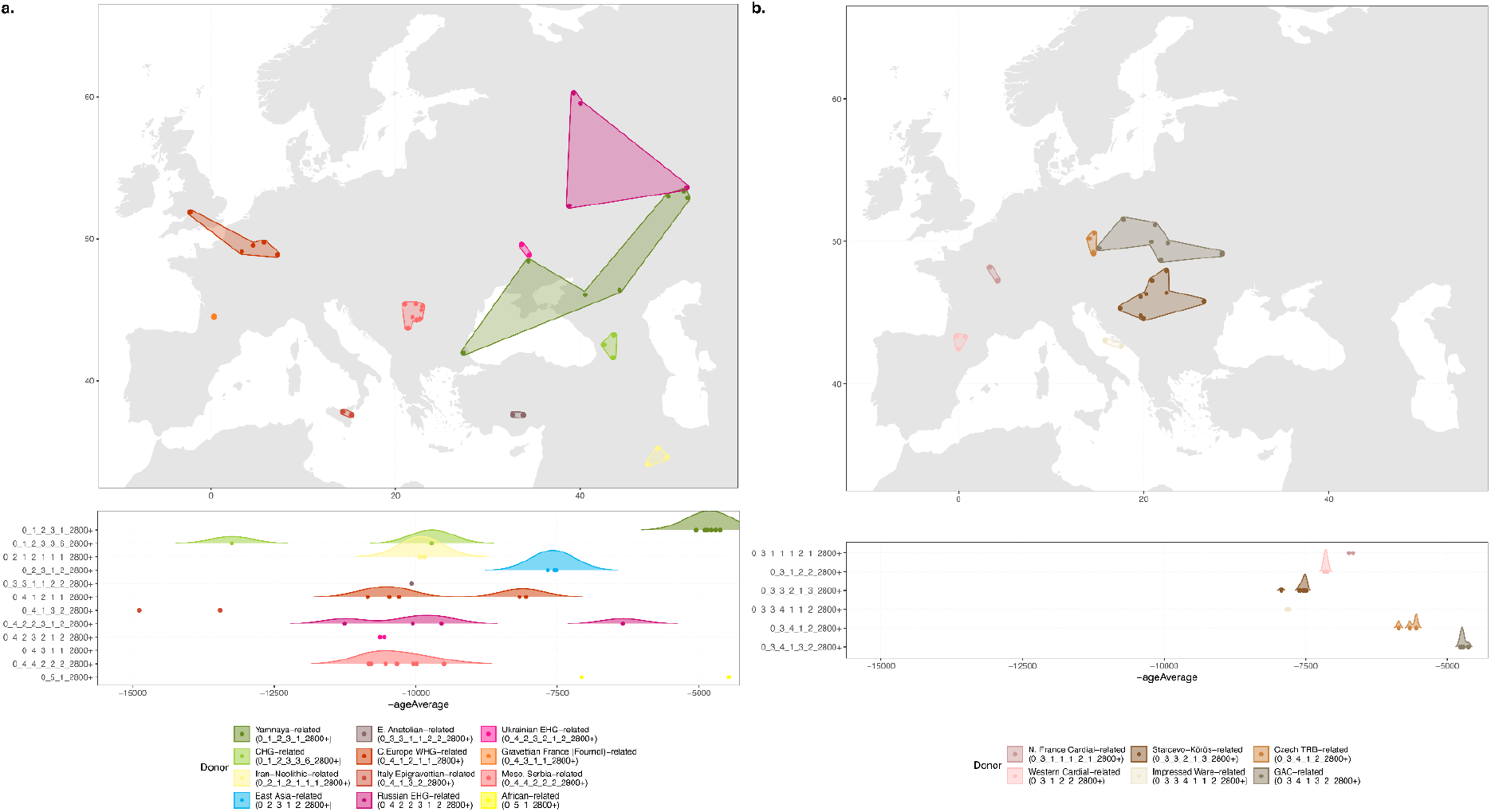
Geographical and temporal distribution of individuals used as sources in the IBD mixture modelling. **(a)** Non-European farming sources. **(b)** European farming sources.

**Extended Data Fig. 3.**
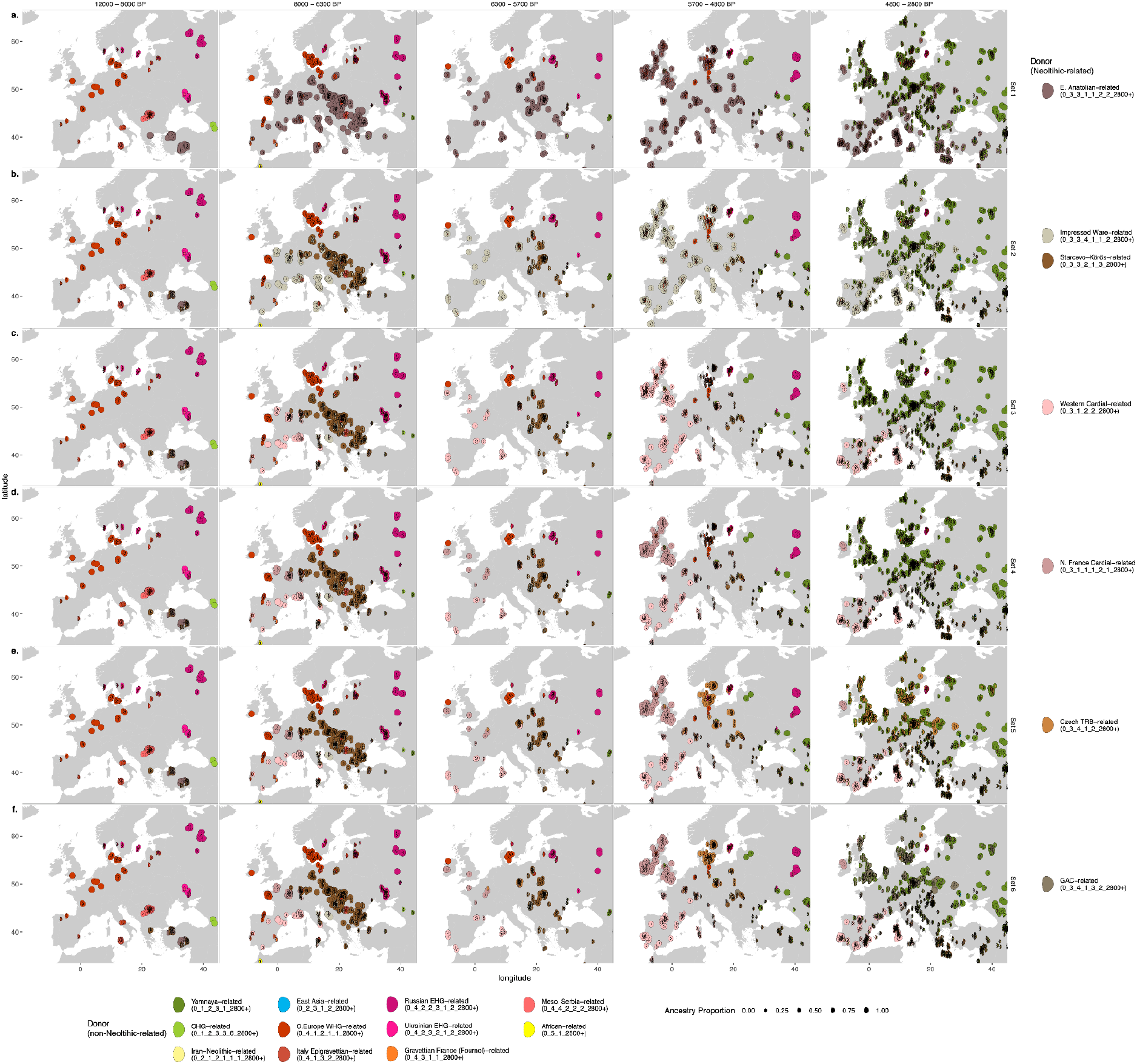
IBD mixture modelling results for all European ancient individuals from 12000 to 2800 BP. **(a)** Set 1, including a single Neolithic-related source: E. Anatolia-related. **(b)** Set 2, including two additional Neolithic-related sources: Impressed Ware- and Starčevo-Körös-related. **(c)** Set 3, including an additional Neolithic-related source: Western Cardial-related. **(d)** Set 4, including an additional Neolithic-related source: N. France Neolithic-related. **(e)** Set 5, including an additional Neolithic-related source: Czech TRB-related. **(f)** Set 6, including an additional Neolithic-related source: GAC-related. All sources from earlier sets are included in later sets. Non-Neolithic sources included in all sets are shown in the legend below. The full set of mixture modelling results (sets 1-6) for all individuals are plotted as bar charts in Supplementary Fig. S3.16

**Extended Data Fig. 4.**
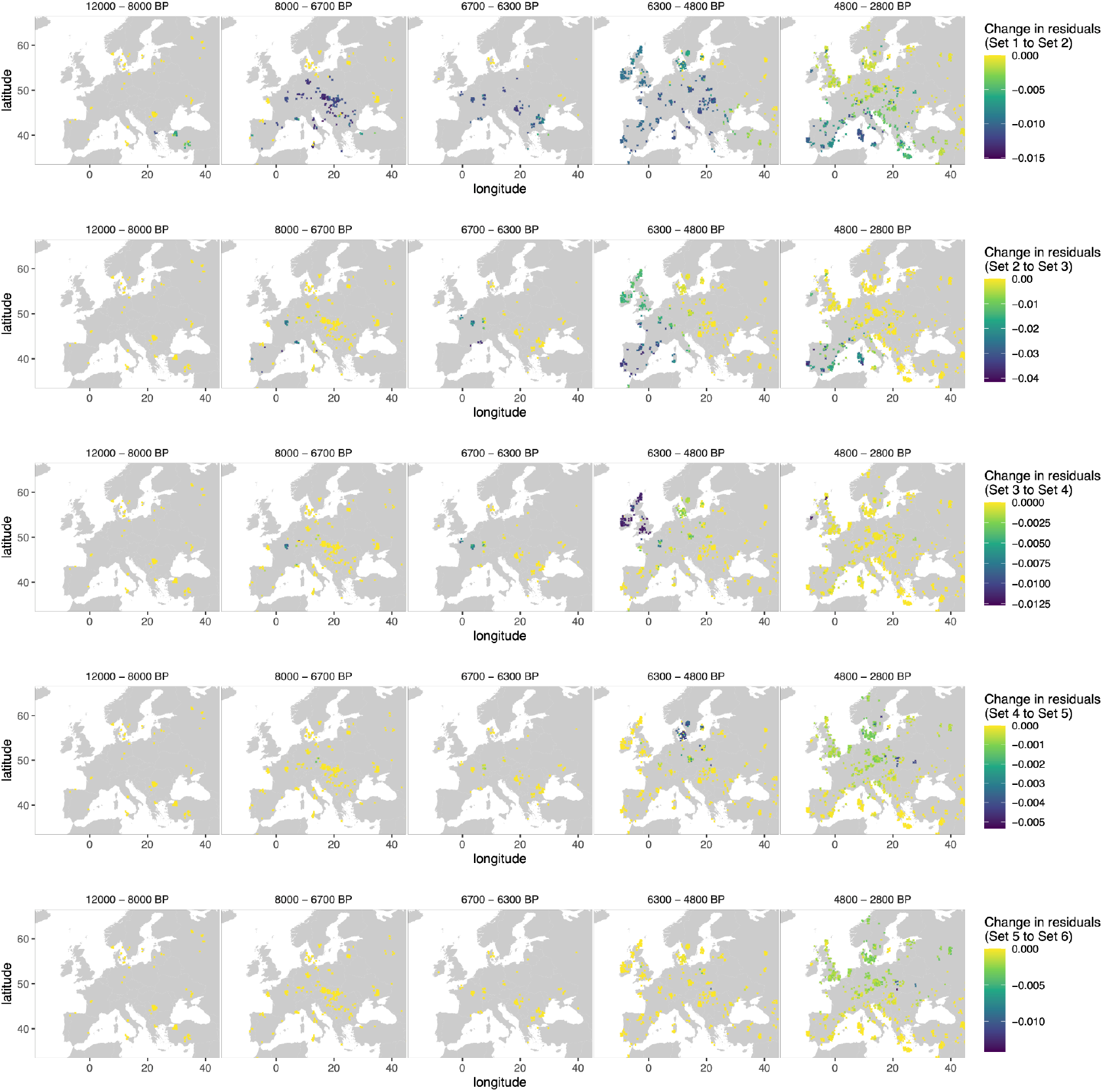
The change in residuals for IBD mixture modelling results for all European ancient individuals from 12000 to 2800 BP. **(a)** Set 1 to Set 2, representing the inclusion of the Impressed Ware- and Starčevo-Körös-related sources. **(b)** Set 2 to Set 3, representing the inclusion of the Western Cardial-related source. **(c)** Set 3 to Set 4, representing the inclusion of the N. France Neolithic-related source. **(d)** Set 4 to Set 5, representing the inclusion of the TRB-related source. **(e)** Set 5 to Set 6, representing the inclusion of the GAC-related source. All sets contain all source individuals from previous sets.

**Extended Data Fig. 5.**
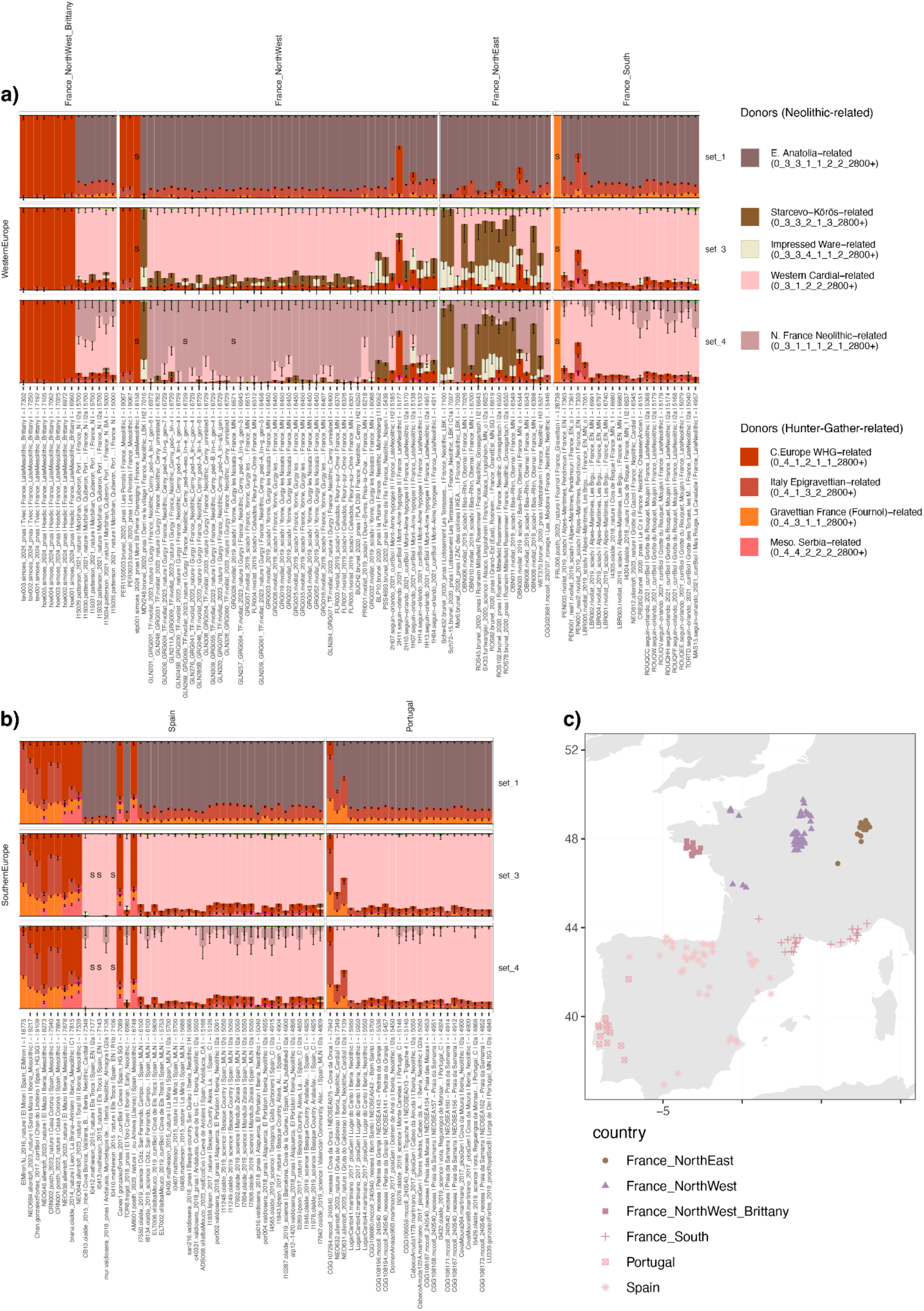
Mixture modelling results for France and Iberia, highlighting changes through time when including additional Neolithic-related sources. **(a)** Mixture modelling results for all individuals older than 4800 BP from various regions of France **(b)** Mixture modelling results for all individuals older than 4800 BP from Iberia. For **(a)** and **(b)**, Set 1 illustrates the transition from Hunter-Gatherer to Farming ancestry using an early Anatolian Farming source, Set 2 illustrates the distinction in proportions of LBK-related and Western Cardial-related ancestry, when the relevant sources are included, and Set 3 illustrates the distinction in proportions of N. French Neolithic-related and Western Cardial-related, when the relevant source is included. Ancient individuals are ordered from oldest to youngest. **(c)** The location of samples from Iberia and France included in **(a)** and **(b)**. The full set of mixture modelling results (sets 1-6) for all individuals are plotted as bar charts in Supplementary Fig. S3.16

**Extended Data Fig. 6.**
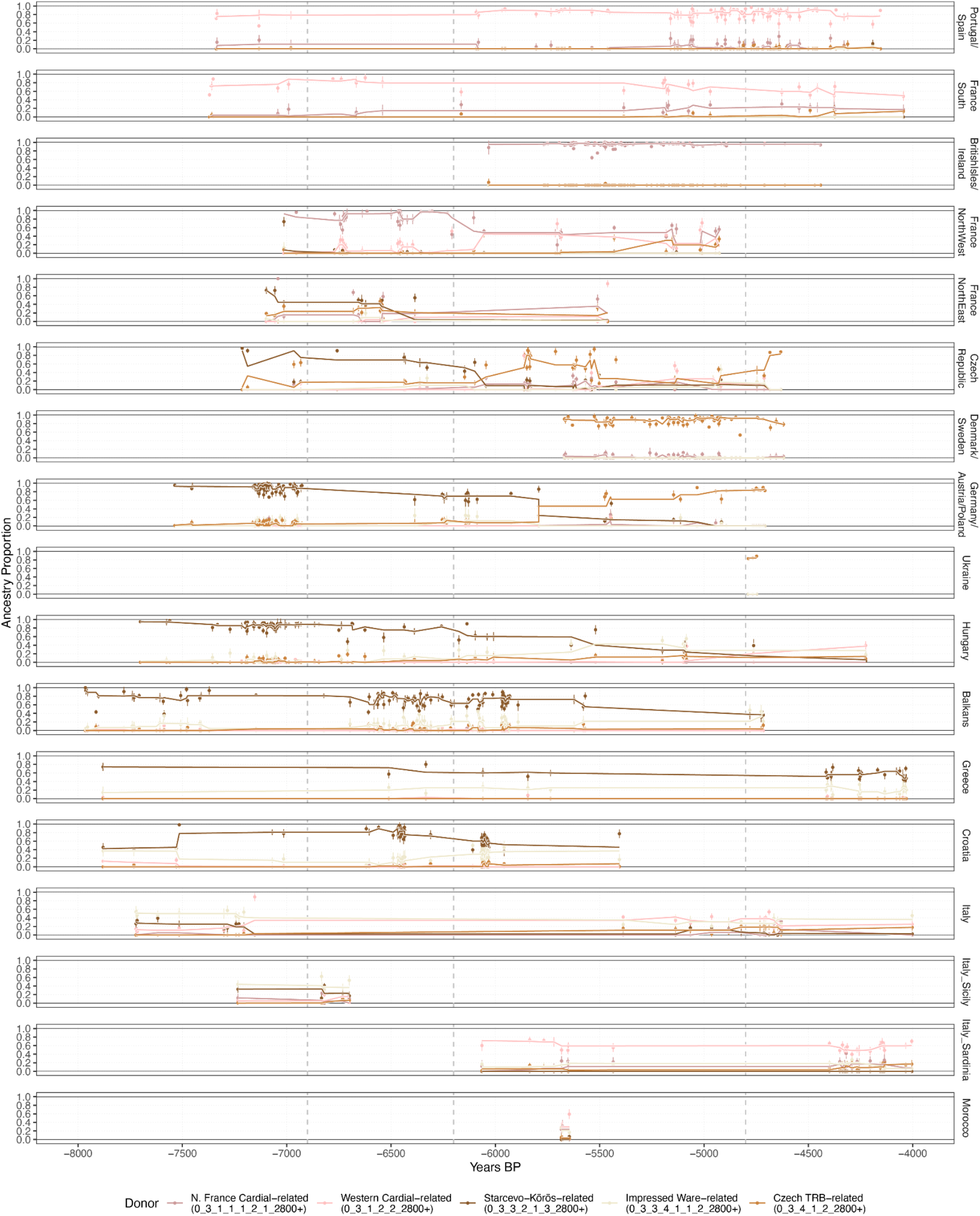
Mixture modelling proportions for European Neolithic Farming-related ancestry sources (Set 5) from across Europe. For each individual, the point estimate for the proportion of each of the five Neolithic ancestries is plotted. Standard errors were calculated using a weighted block jackknife, sequentially leaving out one of the 22 chromosomes at a time. To visualise changing trends through time, a trendline (rolling median of the point estimates with a window of 5) for each ancestry is plotted. The full set of mixture modelling results (sets 1-6) for all individuals are plotted as bar charts in Supplementary Fig. S3.16

**Extended Data Fig. 7.**
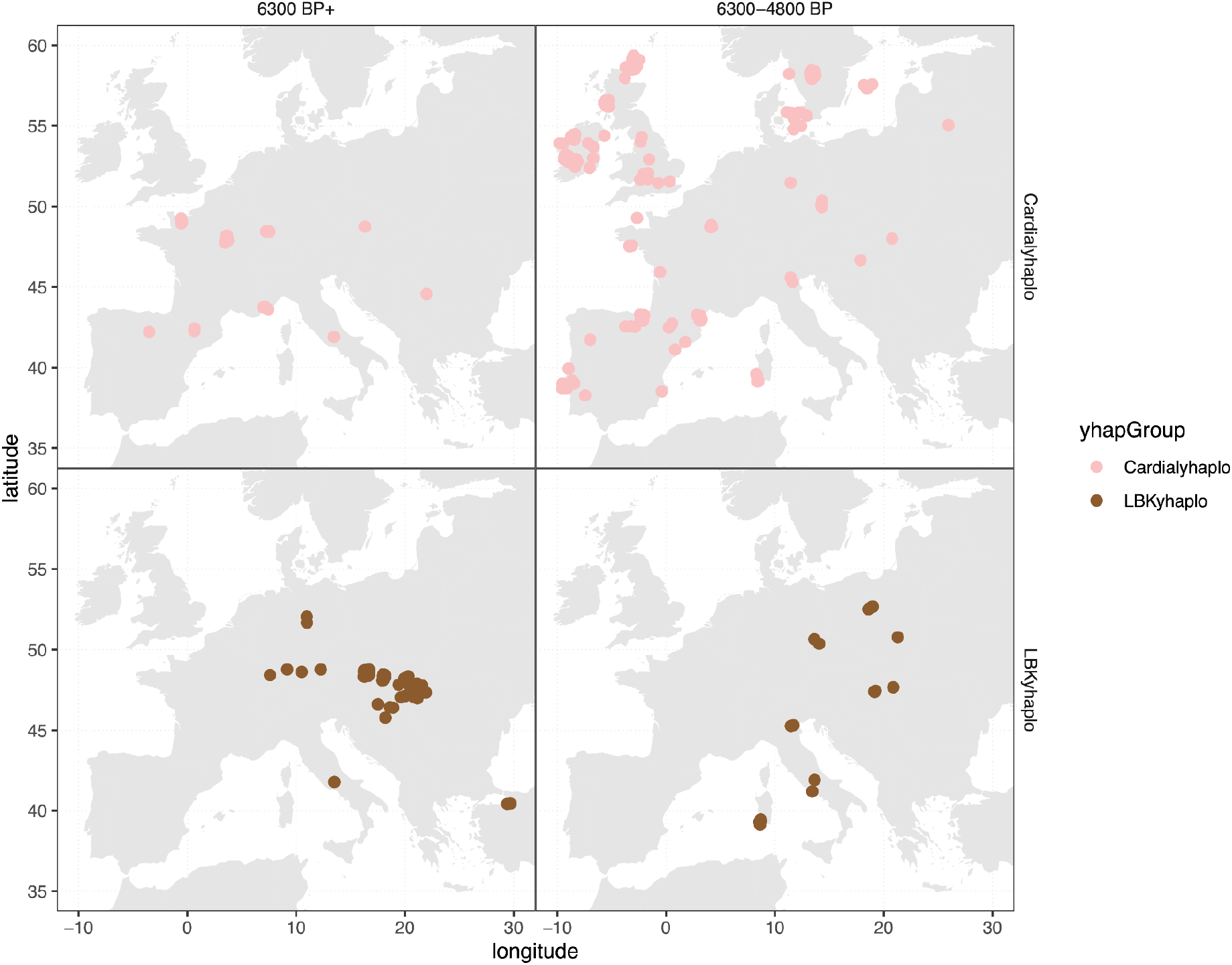
Distribution of Y-haplogroups associated with Cardial and LBK ancestry before 6300 BP (left panels) and between 6300 and 4800 BP (right panels). Y-haplogroups were classified as Cardial- or LBK-associated based on their age and their occurrence among multiple individuals carrying the corresponding autosomal ancestry profiles in the IBD mixture modelling analyses. The distribution of each subhaplogroup is plotted in Supplementary Fig. S3.15.

**Extended Data Fig. 8.**
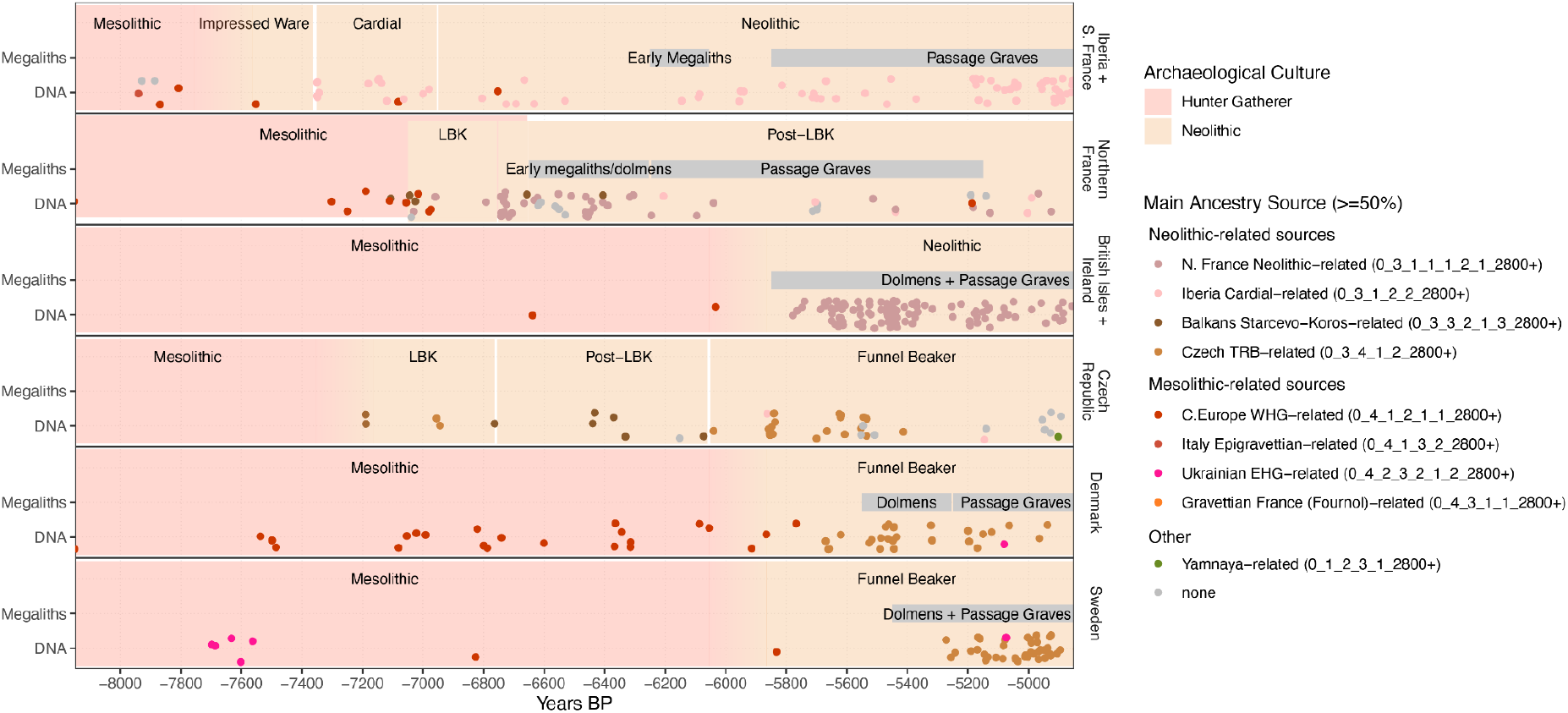
A timeline showing the relevant changes in archaeological cultures, emergence of megaliths, and genetic transitions occurring throughout the Mesolithic and Neolithic. Ancient individuals are coloured when they are modelled with >50% ancestry with one of the sources (Set 5). The full set of mixture modelling results (sets 1-6) for all individuals are plotted as bar charts in Supplementary Fig. S3.16

**Extended Data Table 1.** A summary of the expectations for the spread of megaliths with migrations.

| Region | Migration model |  |  | Cultural diffusion model |  | Conclusion |
| --- | --- | --- | --- | --- | --- | --- |
|  | Expectation 1:<br>Migration from region with megaliths | Expectation 2:<br>Migration occurs contemporaneous with megaliths | Expectation 3:<br>Divergence of megalithic types mirrors population divergences | Expectation 4:<br>Diffusion not restricted to any specific population | Expectation 5:<br>contact and communication between populations |  |
| British Isles | **Consistent:**<br>Migration from northern France | **Consistent:**<br>Migration 6100-6000 BP, megaliths 6000-5900 BP | **Consistent:**<br>Passage graves and megaliths building practiced by migrants | NA:<br>Co-occurs with migration | NA:<br>Co-occurs with migration | **Consistent with migration.** |
| Iberia | **Inconsistent:**<br>No evidence of migration from northern France | **Inconsistent:**<br>No evidence of migration from northern France during first (~6500 BP) or second phase (6000-5900 BP to W. Iberia) | **Inconsistent:**<br>Passage graves/dolmens represent arrival of megalithic style with origins in N. France | **Consistent:**<br>Spreads from people of N. France ancestry to W. Cardial ancestry | **Consistent:**<br>Spreads within Jadeite trade network | **Consistent with cultural diffusion.** No evidence of migration from regions in which passage graves/dolmens were built |
| Scandinavia | **Consistent:**<br>Migration from northern France (formation of TRB) | **Inconsistent:**<br>Migration 6100-6000 Dolmens 5600 BP, Passage graves 5400 BP | **Inconsistent:**<br>Dolmens represent arrival of megalithic style practiced elsewhere for 100s years, rather than local development | **Consistent:**<br>Spreads from people of N. France/W. Cardial ancestry to TRB ancestry | **Consistent:**<br>Spreads within Jadeite trade network | **Consistent with cultural diffusion.** Spreading with migration requires retaining of technical ability for hundreds of years (Passage Graves ~300 years, Dolmens ~600 years) |
| Sardinia | **Consistent:**<br>evidence of N. France Neolithic-related ancestry in small proportions | **Unclear:**<br>Unable to assess time of migration due to absence of pre-megalithic samples | **Consistent:**<br>Passage graves built by people of similar ancestry | **Unclear:**<br>Unable to assess time of migration due to absence of pre-megalithic samples | **Consistent:**<br>Spreads within Jadeite trade network | **Potentially consistent with migration.** Not currently possible to determine when N. France Neolithic ancestry arrived due to an absence of pre-megalithic samples. |

## Methods

### Data generation and authentication

For each of the 46 ancient individuals included in this study, tooth and/or petrous bone material were drilled under sterile conditions in dedicated ancient DNA labs at the Centre for GeoGenetics in Copenhagen. DNA was extracted and non-USER and USER treated libraries were built for all samples, either as part of an automated workflow to build double-stranded libraries as described in 34 or an updated workflow to generate single-stranded libraries based on ^34^. Libraries were amplified using either real-time PCR to determine the number of cycles, or using a pre-set number of 11, 12, 14, 16 or 18 cycles.

Sequencing reads were mapped to the human reference genome as described in ^34^. In brief, adapters were removed and paired reads collapsed using Adapter Removal v2 ^80^ and reads were aligned to the human reference genome build 37 (hs37d5, md5sum 12a0bed94078e2d9e8c00da793bbc84e) using bwa aln (*0.*7.17)^81^. Duplicates were marked using Picard MarkDuplicates (2.25.0).

Libraries were authenticated as ancient based on short fragment lengths and high damage patterns typical of ancient DNA. All non-USER-treated and USER treated data was merged for downstream analysis. Contamination estimates were assessed using contamix^82^, and can be found in Supplementary Table S3.3.

### Dataset preparation

Genomes above 0.1× were imputed with GLIMPSE (v1.1.1)^83^, as described in Allentoft et al.^28^ and Sousa da Mota, B. et al.^84^ against the 30× coverage version of the 1000 Genomes (1000G) v5 phase 3 dataset^85^ that was phased using the TOPMed panel^86^, before being lifted over to hg19. The imputed genomes were then merged with a reference panel of unrelated individuals suitable for imputation (coverage of at least 0.1× for shotgun sequenced data, 1× for 1240K capture data). The panel includes samples from published studies ^6,8–10,16–20,23–26,28,33,35,42,47,48,53,54,82,87–218^. Only individuals with an average genotype probability greater than 0.9 on the 1240K capture sites were included. Only sites that were part of the 1240K capture set, with an INFO score >0.5, and passing the 1000G mappability mask were included. The resulting dataset contained 730,235 sites considered for analyses. IBD segments were called between all pairs of individuals using IBDseq ^219^ on the imputed panel. Segments with a LOD score less than 3, or from hotspot regions of sharing were removed as described in ^28^. Sample metadata was compiled from ^34,220^ as well as the respective papers, and is included in Supplementary Table S3.4.

### IBD clustering and mixture modelling

To prepare for clustering, the summed length of shared segments and number of segments were calculated for each pair of ancient individuals. For closely related individuals (over 1500 cM shared in less than 150 segments), the lower-coverage individual was excluded, leaving 36 new individuals.

The summed length of shared segments was again calculated for all segments greater than 2 cM. Sharing between all pairs in which the summed length was greater than 5 cM was used as input for a network-based hierarchical clustering ^221^ as described in ^28^. Finally, a manual curation step was applied, in which samples dated to less than 2800 BP were excluded, leaving 2954 (2917 published + 37/46 newly sequenced) individuals. The final cluster ID and metadata for all clustered individuals can be found in Supplementary Table S3.4. The metadata was manually compiled and curated from the respective publications and the Allen Ancient DNA Resource^222^.

To understand the relationship between individuals within the dataset, we performed IBD mixture modelling as described in ^28,34^. To prepare the data, the summed length of shared segments was again calculated for all segments greater than 1cM. For each individual, we created a ‘painting palette’, representing the proportion of IBD shared with all clusters of individuals in the dataset. The amount of sharing was then normalised by the number of individuals in the respective cluster. To account for the fact that there are n-1 individuals to share with within-cluster, one individual from all other clusters was excluded from the calculation at random. Mixture modelling was then performed by setting some individuals as ‘sources’ and others as ‘targets’, and modelling the target palettes as combinations of the ‘source’ palettes, using a non-negative least squares approach, similar to chromosome painting^223^. To ensure a decrease in residuals is not due to overfitting, we recalculated mixture modelling proportions using the palettes of the even-numbered chromosomes, and then used those proportions to calculate the residuals for palettes of the odd-numbered chromosomes

### Y-haplogroup calling

To assign Y-chromosome haplogroups, we used ybyra v0.5^224^ using customised ancient DNA parameters (--step-size 5; --damage-filter: true; --damage-model: bidirectional) and using FamilyTreeDNA Haplotree v.2026.06.01. We included in the analysis all ancient males with >=50% Neolithic farming ancestry in the mixture modelling analyses, as well as Mesolithic and Palaeolithic individuals from Europe, totalling 611 males. Results can be found in Table S3.7 and Figures S10-S15.

### Testing the association between ancestry components and the emergence of megalithic monuments

We tested whether the regional proportion of any ancestry component was associated with the appearance of megalithic monuments using a discrete-time survival model, treating megalith emergence as a region-level, irreversible (absorbing) event. Calendar time was binned (250 yr), and for each region and bin we summarised ancestry as the mean proportion of each source across the individuals sampled in that bin; a region contributed at-risk person-periods until the bin of its emergence date (the event) and was right-censored if it never developed megaliths. Each ancestry source was screened individually — jointly fitting all sources is infeasible given only six events — using the Rao score statistic for adding that source to a baseline hazard, with effect sizes estimated by Firth’s penalised likelihood to guard against separation. We fitted two baselines, one constant (marginal association) and one linear in calendar time (association conditional on the secular ancestry trend), and assessed significance by permuting the emergence dates across regions (5000 permutations), controlling the family-wise error rate across sources within each source set. Full model specification, the permutation scheme, and complete per-source results are given in Supplement Note S3.3.2.

## Data availability

Sequence data for the 46 ancient genomes generated for this study have been deposited on the European Nucleotide Archive (ENA) under accession xxxxxxx. The public ancient genomes included in the dataset are listed in Supplementary Table S1.1 and referenced in the methods section.

All maps were generated in R (v4.2.3), using ‘map_data’ from ggplot2 (v3.5.0.9000)

## Code availability

Analyses were done using freely available software, which have been fully referenced throughout the manuscript and supplemental materials and methods. Custom code can be found at https://github.com/ras-nielsen/megalith-hazard-model.

## Acknowledgements

We would like to thank Johannes Müller, Kathrin Sachsenberg and Thomas Frey for helpful discussions over the years, and the Museu Geológico in Lisbon, the Museu Arqueológico de São Miguel de Odrinhas, the Dipartimento di Scienze della Vita e dell’Ambiente, Sezione di Neuroscienze e Antropologia, Monserrato, Cagliari, the Musée de Préhistoire de Carnac and the Musée de la Préhistoire Finistérienne in Penmarc’h for their assistance with sample collection. We thank Illumina for collaboration.

HMC and BSP were supported by ERC-StG NEOSEA 949424. SS was supported by the ERC Synergy project COREX, funded by the European Research Council (ERC) under the European Union’s Horizon 2020 research and innovation programme (Grant Agreement No. 951385). MS and GK were supported by Riksbankens Jubileumsfond (grant M 21-0018 for the 6-year programme MARITIME ENCOUNTERS: a counterpoint to the dominant terrestrial narrative of European prehistory). KK was supported by Riksbankens Jubileumsfond (M16-0455:1, Rise II) and the Synergy project “COREX” funded by the European Research Council (ERC) under the European Union’s Horizon 2020 research and innovation programme (Grant agreement no. 951385). The Lundbeck Foundation GeoGenetics Centre is supported by grants from the Lundbeck Foundation (R302-2018-2155, R155-2013-16338), the Novo Nordisk Foundation (NNF18SA0035006), the Wellcome Trust (WT214300), Carlsberg Foundation (CF18-0024), the Danish National Research Foundation (DNRF94, DNRF174), the University of Copenhagen (KU2016 programme) and Ferring Pharmaceuticals A/S.

## Author contributions

BSP and EW initiated the study. HMC, BSP and EW led the study. HMC, BSP and EW conceptualised the study. BSP, LV, KK and EW supervised the research. BSP and EW acquired funding for research. BSP, MGM, RPS, VS, RP, CC, EP, AMS, AFC, AB, JP, JNG, BI, VD and JM were involved in sample collection. HMC, BSP, MGM, RPS, VS, CC, AMS, AFC and BI curated bioarchaeological data. HMC, RPS, VS, RP, BP, MH, ABP, JS, CG and LV were involved in data generation. HMC, BSP, RN, TP, RP and MS were involved in data analysis. HMC, BSP, SS, RN, JW, MGM, CI, MEA, KK and EW drafted the main text. HMC, BSP, SS, RN, TP, JW, RF, MGM, CI, RPS, VS, RP, CC, EP, BP, AMS and AFC drafted the Supplementary Notes and materials. HMC, BSP, SS, RN, TP, JW, RF, MGM, CI, RPS, RP, AMS, AFC, GK, ABP, JS, CG, LV, MS, MEA, KK and EW were involved in reviewing and editing drafts.

## References

1. Schulz Paulsson, B. Radiocarbon dates and Bayesian modeling support maritime diffusion model for megaliths in Europe. Proc. Natl. Acad. Sci. 116, 3460–3465 (2019).

2. Arnold, B. ‘Arierdämmerung’: race and archaeology in Nazi Germany. World Archaeol. 38, 8–31 (2006).

3. Price, T. D., Knipper, C., Grupe, G. & Smrcka, V. Strontium Isotopes and Prehistoric Human Migration: The Bell Beaker Period in Central Europe. Eur. J. Archaeol. 7, 9–40 (2004).

4. Bentley, R. A. et al. Prehistoric Migration in Europe: Strontium Isotope Analysis of Early Neolithic Skeletons. Curr. Anthropol. 43, 799–804 (2002).

5. Collard, M., Edinborough, K., Shennan, S. & Thomas, M. G. Radiocarbon evidence indicates that migrants introduced farming to Britain. J. Archaeol. Sci. 37, 866–870 (2010).

6. Allentoft, M. E. et al. Population genomics of Bronze Age Eurasia. Nature 522, 167–172 (2015).

7. Haak, W. et al. Massive migration from the steppe was a source for Indo-European languages in Europe. Nature 522, 207–211 (2015).

8. de Barros Damgaard, P., et al. The first horse herders and the impact of early Bronze Age steppe expansions into Asia. Science 360, eaar7711 (2018).

9. Narasimhan, V. M. et al. The formation of human populations in South and Central Asia. Science 365, eaat7487 (2019).

10. Saag, L. et al. The Arrival of Siberian Ancestry Connecting the Eastern Baltic to Uralic Speakers further East. Curr. Biol. 29, 1701–1711.e16 (2019).

11. The Indo-European Puzzle Revisited: Integrating Archaeology, Genetics, and Linguistics. (Cambridge University Press, 2023).

12. Frieman, C. J. & Hofmann, D. Present pasts in the archaeology of genetics, identity, and migration in Europe: a critical essay. World Archaeol. 51, 528–545 (2019).

13. Childe, V. G. Prehistoric Communities of the British Isles. (W. & R. Chambers, London, 1940).

14. Renfrew, C. Before Civilisation: The Radiocarbon Revolution. (Penguin, London, 1973).

15. Müller, J., Kruckenberg, C., Großmann, R. & Luckner, J. A Map of European Megaliths. J. Neolit. Archaeol. 165–173 (2023) doi:10.12766/jna.2023.6.

16. Cassidy, L. M. et al. A dynastic elite in monumental Neolithic society. Nature 582, 384– 388 (2020).

17. Seersholm, F. V. et al. Repeated plague infections across six generations of Neolithic Farmers. Nature 632, 114–121 (2024).

18. Skoglund, P. et al. Separating endogenous ancient DNA from modern day contamination in a Siberian Neandertal. Proc. Natl. Acad. Sci. 111, 2229–2234 (2014).

19. Cassidy, L. M. et al. Neolithic and Bronze Age migration to Ireland and establishment of the insular Atlantic genome. Proc. Natl. Acad. Sci. 113, 368–373 (2016).

20. Fowler, C. et al. A high-resolution picture of kinship practices in an Early Neolithic tomb. Nature 601, 584–587 (2022).

21. Mathieson, I. et al. Genome-wide patterns of selection in 230 ancient Eurasians. Nature 528, 499–503 (2015).

22. Rivollat, M., et al. Ancient DNA gives new insights into a Norman Neolithic monumental cemetery dedicated to male elites. Proc. Natl. Acad. Sci. 119, e2120786119 (2022).

23. Valdiosera, C. et al. Four millennia of Iberian biomolecular prehistory illustrate the impact of prehistoric migrations at the far end of Eurasia. Proc. Natl. Acad. Sci. 115, 3428–3433 (2018).

24. Brace, S. et al. Ancient genomes indicate population replacement in Early Neolithic Britain. Nat. Ecol. Evol. 3, 765–771 (2019).

25. Sánchez-Quinto, F. et al. Megalithic tombs in western and northern Neolithic Europe were linked to a kindred society. Proc. Natl. Acad. Sci. 116, 9469–9474 (2019).

26. Villalba-Mouco, V. et al. Survival of Late Pleistocene Hunter-Gatherer Ancestry in the Iberian Peninsula. Curr. Biol. 29, 1169–1177.e7 (2019).

27. Betti, L. et al. Climate shaped how Neolithic farmers and European hunter-gatherers interacted after a major slowdown from 6,100 bce to 4,500 bce. *Nat*. Hum. Behav. 4, 1004–1010 (2020).

28. Allentoft, M. E. et al. Population genomics of post-glacial western Eurasia. Nature 625, 301–311 (2024).

29. Taylor, R., García-Rivero, D., Gonçalves, C., Cascalheira, J. & Bicho, N. The Early Neolithic at the Muge Shellmiddens (Portugal): Analysis and Review of the Ceramic Evidence from Cabeço da Amoreira. Eur. J. Archaeol. 24, 156–179 (2021).

30. Bouby, L. et al. Early Neolithic (ca. 5850-4500 cal BC) agricultural diffusion in the Western Mediterranean: An update of archaeobotanical data in SW France. PLOS ONE 15, e0230731 (2020).

31. Kirleis, W. & Fischer, E. Neolithic cultivation of tetraploid free threshing wheat in Denmark and Northern Germany: implications for crop diversity and societal dynamics of the Funnel Beaker Culture. Veg. Hist. Archaeobotany 23, 81–96 (2014).

32. Pétrequin, P. & Pétrequin, A.-M. Du Jade Pour Les Dieux: Un Système Religieux de l’Europe Néolithique, Ve et IVe Millénaires Av. J.-C. Tomes 5 et 6. (Presses universitaires de Franche-Comté, 2025).

33. Brunel, S. et al. Ancient genomes from present-day France unveil 7,000 years of its demographic history. Proc. Natl. Acad. Sci. 117, 12791–12798 (2020).

34. McColl, H. et al. Steppe Ancestry in Western Eurasia and the Spread of the Germanic Languages. 2024.03.13.584607 Preprint at 10.1101/2024.03.13.584607 (2025).

35. Lazaridis, I. et al. Ancient human genomes suggest three ancestral populations for present-day Europeans. Nature 513, 409–413 (2014).

36. Binder, D. et al. Le complexe de la Céramique Imprimée en Méditerranée centrale et nord-occidentale : une synthèse chronoculturelle (VIIe et VIe millénaires AEC). in Céramiques imprimées de Méditerranée occidentale (VIe millénaire AEC) : données, approches et enjeux nouveaux / Western Mediterranean Impressed Wares (6th millennium BCE): New data, approaches and challenges (eds Binder, D. & Manen, C.) vol. 18 27–124 (Société préhistorique française, Paris, 2022).

37. Bánffy, E. The 6th Millennium BC Boundary in Western Transdanubia and Its Role in the Central European Neolithic Transition : The Szentgyörgyvölgy-Pityerdomb Settlement. (Archaeological Institute of the Hungarian Academy of Sciences, Budapest, 2004).

38. van Willigen, S. Between Cardial and Linearbandkeramik: From no-man’s-land to communication sphere. Quat. Int. 470, 333–352 (2018).

39. Prost, D., Biard, M. & Théron, V. La culture de Cerny en Haute-Normandie (Ve millénaire avant J.-C.). in Journées archéologiques de Haute-Normandie. Rouen, 3-5 avril 2009 (eds Erlenbach, F., Lequoy, M.-C. & Legris, M.) 45–62 (Presses universitaires de Rouen et du Havre, Mont-Saint-Aignan, 2010). doi:10.4000/books.purh.6851.

40. Cassen, S. et al. Measuring distinction in the megalithic architecture of the Carnac region: from sign to material. J. Neolit. Archaeol. 10.12766/jna.2010.45 (2010) doi:10.12766/jna.2010.45.

41. Blanchard, A., Guyodo, J.-N., Paulsson, B. S. & Montassier, F. Le Plasker in Plouharnel (fifth millennium cal BC): a newly discovered section of the megalithic complex of Carnac. Antiquity 99, 915–934 (2025).

42. Simões, L. G. et al. Genomic ancestry and social dynamics of the last hunter-gatherers of Atlantic France. Proc. Natl. Acad. Sci. 121, e2310545121 (2024).

43. Paulsson, B. S. Time and Stone: The Emergence and Development of Megaliths and Megalithic Societies in Europe. (Archaeopress Archaeology, Oxford, 2017).

44. Wanner, H., Solomina, O., Grosjean, M., Ritz, S. P. & Jetel, M. Structure and origin of Holocene cold events. Quat. Sci. Rev. 30, 3109–3123 (2011).

45. Allentoft, M. E. et al. 100 ancient genomes show repeated population turnovers in Neolithic Denmark. Nature 625, 329–337 (2024).

46. Skoglund, P. et al. Genomic diversity and admixture differs for Stone-Age Scandinavian foragers and farmers. Science 344, 747–750 (2014).

47. Malmström, H. et al. The genomic ancestry of the Scandinavian Battle Axe Culture people and their relation to the broader Corded Ware horizon. Proc. Biol. Sci. 286, 20191528 (2019).

48. Olalde, I. et al. The Beaker phenomenon and the genomic transformation of northwest Europe. Nature 555, 190–196 (2018).

49. Whittle, A., et al. Gathering Time: Dating the Early Neolithic Enclosures of Southern Britain and Ireland, Volumes 1 and 2. (Oxbow Books, 2011). doi:10.2307/j.ctvh1dwp2.

50. Sheridan, J. A. The Neolithisation of Britain and Ireland: the big picture. in Landscapes in Transition (eds Finlayson, B. & Warren, G.) vol. 8 89–105 (Oxbow, Oxford, 2010).

51. Gebauer, A. B. & Price, T. D. First farmers in Northern Europe. Quat. Sci. Rev. 390, 110170 (2026).

52. Sørensen, L. From hunter to farmer in Northern Europe: Migration and adaptation during the Neolithic and Bronze Age. Acta Archaeol. 85, 73–114 (2014).

53. Fregel, R. et al. Ancient genomes from North Africa evidence prehistoric migrations to the Maghreb from both the Levant and Europe. Proc. Natl. Acad. Sci. 115, 6774–6779 (2018).

54. Antonio, M. L. et al. Ancient Rome: A genetic crossroads of Europe and the Mediterranean. Science 366, 708–714 (2019).

55. Querré, G., Vigier, E. & Cassen, S. La Parure En Callaïs Du Néolithique Européen. (Archaeopress, Oxford, 2019).

56. Tykot, R. H. Obsidian Studies in the Prehistoric Central Mediterranean: After 50 Years, What Have We Learned and What Still Needs to Be Done? Open Archaeol. 3, (2017).

57. Manen, C. Spatial, chronological and cultural dynamics of the Neolithization in the Western Mediterranean. Transit. Néolithique En Méditerranée 405–418 (2014).

58. Kerig, T. et al. Interlinking research: the Big Exchange project. Antiquity : a review of world archaeology vol. 97 (2023).

59. Windler, A. Der Austausch von Spondylus Gaederopus in Europa Zwischen 5500 Und 5000 v. Chr. (Deutsches Bergbau-Museum Bochum, 2018). doi:10.46586/DBM.180.

60. Jeunesse, C. Pratiques funéraires au néolithique ancien. (Editions Errance, Paris, 1997).

61. Whittle, A. & Bickle, P. The First Farmers of Central Europe : Diversity in LBK Lifeways. (Oxbow Books, Oxford (GB), 2013).

62. Gehlen, B. Foragers and Farmers during the Neolithic transition in Western Central Europe: Searching for evidence of mobility and intercultural networks. in Mobility in Prehistoric Sedentary Societies (eds Scharl, S. & Gehlen, B.) 39–60 (Welt und Erde Verlag, 2017).

63. Cassen, S. et al. Carnac. Récit Pour Un Imagier. (Laboratoire de recherche archéologie et architectures / CreAAH, 2021).

64. Cassen, S. et al. Real and ideal European maritime transfers along the Atlantic coast during the Neolithic. Doc. Praehist. 46, 308–325 (2019).

65. Schulz Paulsson, B. Transmission symbolique et la mer dans l’Europe mégalithique (4700–2500 cal av. J.-C.). L’Anthropologie 130, 103459 (2026).

66. Pétrequin, P. et al. Au royaume de Carnac (Morbihan) au milieu du Ve millénaire : les haches polies en fibrolite ibérique. Bull. Société Préhistorique Fr. 122, 405–457 (2025).

67. Sherratt, A. The genesis of megaliths: Monumentality, ethnicity and social complexity in Neolithic north-west Europe. World Archaeol. 22, 147–167 (1990).

68. Murillo-Barroso, M. et al. Amber in prehistoric Iberia: New data and a review. PLOS ONE 13, e0202235 (2018).

69. Pétrequin, P. et al. The value of things: the production and circulation of Alpine jade axes during the 5th–4th millennia in a European perspective. in Economic archaeology: from structure to performance in European archaeology (eds Kerig, T. & Zimmermann, A.) 65–82 (R. Habelt, Bonn, 2013).

70. Goslin, J. et al. A new Holocene relative sea-level curve for western Brittany (France): Insights on isostatic dynamics along the Atlantic coasts of north-western Europe. Quat. Sci. Rev. 129, 341–365 (2015).

71. Waller, M. P. & Long, A. J. Holocene coastal evolution and sea-level change on the southern coast of England: a review. J. Quat. Sci. 18, 351–359 (2003).

72. Joly, C. & Visset, L. Evolution of vegetation landscapes since the Late Mesolithic on the French West Atlantic coast. Rev. Palaeobot. Palynol. 154, 124–179 (2009).

73. Regnauld, H., Jennings, S., Delaney, C. & Lemasson, L. Holocene sea-level variations and geomorphological response: An example from northern Brittany (France). Quat. Sci. Rev. 15, 781–787 (1996).

74. Cyprien, A.-L., Visset, L. & Carcaud, N. Evolution of vegetation landscapes during the Holocene in the central and downstream Loire basin (Western France). Veg. Hist. Archaeobotany 13, 181–196 (2004).

75. Fernane, A. et al. Climate variability and storm impacts as major drivers for human coastal marsh withdrawal over the Neolithic period (Southern Brittany, NW France). Palaeogeogr. Palaeoclimatol. Palaeoecol. 435, 136–144 (2015).

76. Blaschek, M. & Renssen, H. The Holocene thermal maximum in the Nordic Seas: the impact of Greenland Ice Sheet melt and other forcings in a coupled atmosphere–sea-ice– ocean model. Clim Past 9, 1629–1643 (2013).

77. Fyfe, R. M. et al. The Holocene vegetation cover of Britain and Ireland: overcoming problems of scale and discerning patterns of openness. Quat. Sci. Rev. 73, 132–148 (2013).

78. Sørensen, L. & Karg, S. The expansion of agrarian societies towards the north – new evidence for agriculture during the Mesolithic/Neolithic transition in Southern Scandinavia. J. Archaeol. Sci. 51, 98–114 (2014).

79. Warden, L. et al. Climate induced human demographic and cultural change in northern Europe during the mid-Holocene. Sci. Rep. 7, 15251 (2017).

## Methods References

80. Schubert, M., Lindgreen, S. & Orlando, L. AdapterRemoval v2: rapid adapter trimming, identification, and read merging. BMC Res Notes vol. 9 88 (2016).

81. Li, H. & Durbin, R. Fast and accurate short read alignment with Burrows-Wheeler transform. Bioinformatics vol. 25 1754–60 (2009).

82. Fu, Q. et al. Genome sequence of a 45,000-year-old modern human from western Siberia. Nature 514, 445–449 (2014).

83. Rubinacci, S., Ribeiro, D. M., Hofmeister, R. J. & Delaneau, O. Efficient phasing and imputation of low-coverage sequencing data using large reference panels. Nat. Genet. 53, 120–126 (2021).

84. Sousa da Mota, B., et al. Imputation of ancient human genomes. Nat. Commun. 14, 3660 (2023).

85. Auton, A. et al. A global reference for human genetic variation. Nature 526, 68–74 (2015).

86. Taliun, D. et al. Sequencing of 53,831 diverse genomes from the NHLBI TOPMed Program. Nature 590, 290–299 (2021).

87. Rasmussen, M. et al. Ancient human genome sequence of an extinct Palaeo-Eskimo. Nature 463, 757–762 (2010).

88. Rasmussen, M. et al. The genome of a Late Pleistocene human from a Clovis burial site in western Montana. Nature 506, 225–229 (2014).

89. Rasmussen, M. et al. The ancestry and affiliations of Kennewick Man. Nature vol. 523 455–458 (2015).

90. Fu, Q. et al. The genetic history of Ice Age Europe. Nature vol. 534 200–5 (2016).

91. Gamba, C. et al. Genome flux and stasis in a five millennium transect of European prehistory. Nat. Commun. 5, 5257 (2014).

92. Lazaridis, I. et al. Genomic insights into the origin of farming in the ancient Near East. Nature 536, 419–424 (2016).

93. Lazaridis, I. et al. Genetic origins of the Minoans and Mycenaeans. Nature 548, 214–218 (2017).

94. Lazaridis, I. et al. The genetic history of the Southern Arc: A bridge between West Asia and Europe. Science 377, eabm4247 (2022).

95. Olalde, I. et al. Derived immune and ancestral pigmentation alleles in a 7,000-year-old Mesolithic European. Nature 507, 225–228 (2014).

96. Olalde, I. et al. A Common Genetic Origin for Early Farmers from Mediterranean Cardial and Central European LBK Cultures. Mol. Biol. Evol. 32, 3132–3142 (2015).

97. Olalde, I. et al. The genomic history of the Iberian Peninsula over the past 8000 years. Science 363, 1230–1234 (2019).

98. Raghavan, M. et al. The genetic prehistory of the New World Arctic. Science 345, 1255832 (2014).

99. Raghavan, M. et al. Genomic evidence for the Pleistocene and recent population history of Native Americans. Science 349, aab3884 (2015).

100. Seguin-Orlando, A. et al. Genomic structure in Europeans dating back at least 36,200 years. Science 346, 1113–1118 (2014).

101. Seguin-Orlando, A. et al. Heterogeneous Hunter-Gatherer and Steppe-Related Ancestries in Late Neolithic and Bell Beaker Genomes from Present-Day France. Curr. Biol. 31, 1072–1083.e10 (2021).

102. Skoglund, P. et al. Reconstructing Prehistoric African Population Structure. Cell 171, 59–71.e21 (2017).

103. Günther, T. et al. Ancient genomes link early farmers from Atapuerca in Spain to modern-day Basques. Proc. Natl. Acad. Sci. 112, 11917–11922 (2015).

104. Günther, T. et al. Population genomics of Mesolithic Scandinavia: Investigating early postglacial migration routes and high-latitude adaptation. PLOS Biol. 16, e2003703 (2018).

105. Jones, E. R. et al. Upper Palaeolithic genomes reveal deep roots of modern Eurasians. Nat. Commun. 6, 8912 (2015).

106. Jones, E. R. et al. The Neolithic Transition in the Baltic Was Not Driven by Admixture with Early European Farmers. Curr. Biol. 27, 576–582 (2017).

107. Llorente, M. G. et al. Ancient Ethiopian genome reveals extensive Eurasian admixture in Eastern Africa. Science 350, 820–822 (2015).

108. Broushaki, F. et al. Early Neolithic genomes from the eastern Fertile Crescent. Science 353, 499–503 (2016).

109. Gallego-Llorente, M., et al. The genetics of an early Neolithic pastoralist from the Zagros, Iran. Sci. Rep. 6, 31326 (2016).

110. Hofmanová, Z. et al. Early farmers from across Europe directly descended from Neolithic Aegeans. Proc. Natl. Acad. Sci. 113, 6886–6891 (2016).

111. Kılınç, G. M. et al. The Demographic Development of the First Farmers in Anatolia. Curr. Biol. 26, 2659–2666 (2016).

112. Kılınç, G. M. et al. Human population dynamics and Yersinia pestis in ancient northeast Asia. Sci. Adv. 7, eabc4587 (2021).

113. Martiniano, R. et al. Genomic signals of migration and continuity in Britain before the Anglo-Saxons. Nat. Commun. 7, 10326 (2016).

114. Martiniano, R. et al. The population genomics of archaeological transition in west Iberia: Investigation of ancient substructure using imputation and haplotype-based methods. PLOS Genet. 13, e1006852 (2017).

115. Schiffels, S. et al. Iron Age and Anglo-Saxon genomes from East England reveal British migration history. Nat. Commun. 7, 10408 (2016).

116. González-Fortes, G. et al. Paleogenomic Evidence for Multi-generational Mixing between Neolithic Farmers and Mesolithic Hunter-Gatherers in the Lower Danube Basin. Curr. Biol. 27, 1801–1810.e10 (2017).

117. González-Fortes, G. et al. A western route of prehistoric human migration from Africa into the Iberian Peninsula. Proc. R. Soc. B Biol. Sci. 286, 20182288 (2019).

118. Haber, M. et al. Continuity and Admixture in the Last Five Millennia of Levantine History from Ancient Canaanite and Present-Day Lebanese Genome Sequences. Am. J. Hum. Genet. 101, 274–282 (2017).

119. Haber, M. et al. A Transient Pulse of Genetic Admixture from the Crusaders in the Near East Identified from Ancient Genome Sequences. Am. J. Hum. Genet. 104, 977–984 (2019).

120. Hansen, H. B. et al. Comparing Ancient DNA Preservation in Petrous Bone and Tooth Cementum. PLoS One vol. 12 e0170940 (2017).

121. Lipson, M. & Reich, D. A working model of the deep relationships of diverse modern human genetic lineages outside of Africa. Mol. Biol. Evol. msw293 (2017) doi:10.1093/molbev/msw293.

122. Rodríguez-Varela, R. et al. Genomic Analyses of Pre-European Conquest Human Remains from the Canary Islands Reveal Close Affinity to Modern North Africans. Curr. Biol. CB 27, 3396–3402.e5 (2017).

123. Rodríguez-Varela, R. et al. The genetic history of Scandinavia from the Roman Iron Age to the present. Cell 186, 32–46.e19 (2023).

124. Saag, L. et al. Extensive Farming in Estonia Started through a Sex-Biased Migration from the Steppe. Curr. Biol. 27, 2185–2193.e6 (2017).

125. Saag, L. et al. Genetic ancestry changes in Stone to Bronze Age transition in the East European plain. Sci. Adv. 7, eabd6535 (2021).

126. Schlebusch, C. M. et al. Southern African ancient genomes estimate modern human divergence to 350,000 to 260,000 years ago. Science 358, 652–655 (2017).

127. Sikora, M. et al. Ancient genomes show social and reproductive behavior of early Upper Paleolithic foragers. Science 358, 659–662 (2017).

128. Sikora, M. et al. The population history of northeastern Siberia since the Pleistocene. Nature 570, 182–188 (2019).

129. Unterländer, M. et al. Ancestry and demography and descendants of Iron Age nomads of the Eurasian Steppe. Nat. Commun. 8, 14615 (2017).

130. Amorim, C. E. G. et al. Understanding 6th-century barbarian social organization and migration through paleogenomics. Nat. Commun. 9, 3547 (2018).

131. Damgaard, P. de B., et al. 137 ancient human genomes from across the Eurasian steppes. Nature 557, 369–374 (2018).

132. Ebenesersdóttir, S. S. et al. Ancient genomes from Iceland reveal the making of a human population. Science 360, 1028–1032 (2018).

133. Fernandes, D. M. et al. A genomic Neolithic time transect of hunter-farmer admixture in central Poland. Sci. Rep. 8, 14879 (2018).

134. Fernandes, D. M. et al. The spread of steppe and Iranian-related ancestry in the islands of the western Mediterranean. Nat. Ecol. Evol. 4, 334–345 (2020).

135. Krzewińska, M. et al. Genomic and Strontium Isotope Variation Reveal Immigration Patterns in a Viking Age Town. Curr. Biol. 28, 2730–2738.e10 (2018).

136. Krzewińska, M. et al. Ancient genomes suggest the eastern Pontic-Caspian steppe as the source of western Iron Age nomads. Sci. Adv. 4, eaat4457 (2018).

137. Lindo, J. et al. The genetic prehistory of the Andean highlands 7000 years BP though European contact. Sci. Adv. 4, eaau4921 (2018).

138. McColl, H. et al. The prehistoric peopling of Southeast Asia. Science 361, 88–92 (2018).

139. Mittnik, A. et al. The genetic prehistory of the Baltic Sea region. Nat. Commun. 9, 442 (2018).

140. Moreno-Mayar, J. V. et al. Terminal Pleistocene Alaskan genome reveals first founding population of Native Americans. Nature 553, 203–207 (2018).

141. Moreno-Mayar, J. V. et al. Early human dispersals within the Americas. Science 362, eaav2621 (2018).

142. Scheib, C. L. et al. Ancient human parallel lineages within North America contributed to a coastal expansion. Science 360, 1024–1027 (2018).

143. Veeramah, K. R. et al. Population genomic analysis of elongated skulls reveals extensive female-biased immigration in Early Medieval Bavaria. Proc. Natl. Acad. Sci. 115, 3494–3499 (2018).

144. Feldman, M. et al. Ancient DNA sheds light on the genetic origins of early Iron Age Philistines. Sci. Adv. 5, eaax0061 (2019).

145. Harney, É. et al. Ancient DNA from the skeletons of Roopkund Lake reveals Mediterranean migrants in India. Nat. Commun. 10, 3670 (2019).

146. Harney, É., Patterson, N., Reich, D. & Wakeley, J. Assessing the performance of qpAdm: a statistical tool for studying population admixture. Genetics 217, iyaa045 (2021).

147. Jensen, T. Z. T. et al. A 5700 year-old human genome and oral microbiome from chewed birch pitch. Nat. Commun. 10, 5520 (2019).

148. Järve, M. et al. Shifts in the Genetic Landscape of the Western Eurasian Steppe Associated with the Beginning and End of the Scythian Dominance. Curr. Biol. 29, 2430–2441.e10 (2019).

149. Kanzawa-Kiriyama, H. et al. Late Jomon male and female genome sequences from the Funadomari site in Hokkaido, Japan. Anthropol. Sci. 127, 83–108 (2019).

150. Kashuba, N. et al. Ancient DNA from mastics solidifies connection between material culture and genetics of mesolithic hunter–gatherers in Scandinavia. Commun. Biol. 2, 1– 10 (2019).

151. Ning, C. et al. Ancient Genomes Reveal Yamnaya-Related Ancestry and a Potential Source of Indo-European Speakers in Iron Age Tianshan. Curr. Biol. 29, 2526–2532.e4 (2019).

152. Ning, C. et al. Ancient genomes from northern China suggest links between subsistence changes and human migration. Nat. Commun. 11, 2700 (2020).

153. Schroeder, H. et al. Unraveling ancestry, kinship, and violence in a Late Neolithic mass grave. Proc. Natl. Acad. Sci. 116, 10705–10710 (2019).

154. Agranat-Tamir, L. et al. The Genomic History of the Bronze Age Southern Levant. Cell 181, 1146–1157.e11 (2020).

155. Coutinho, A. et al. The Neolithic Pitted Ware culture foragers were culturally but not genetically influenced by the Battle Axe culture herders. Am. J. Phys. Anthropol. 172, 638–649 (2020).

156. Furtwängler, A. et al. Comparison of target enrichment strategies for ancient pathogen DNA. BioTechniques 69, 455–459 (2020).

157. Jeong, C. et al. A Dynamic 6,000-Year Genetic History of Eurasia’s Eastern Steppe. Cell 183, 890–904.e29 (2020).

158. Linderholm, A. et al. Corded Ware cultural complexity uncovered using genomic and isotopic analysis from south-eastern Poland. Sci. Rep. 10, 6885 (2020).

159. Marcus, J. H. et al. Genetic history from the Middle Neolithic to present on the Mediterranean island of Sardinia. Nat. Commun. 11, 939 (2020).

160. Margaryan, A. et al. Population genomics of the Viking world. Nature 585, 390–396 (2020).

161. Bortolini, E. et al. Early Alpine occupation backdates westward human migration in Late Glacial Europe. Curr. Biol. 31, 2484–2493.e7 (2021).

162. Clemente, F. et al. The genomic history of the Aegean palatial civilizations. Cell 184, 2565–2586.e21 (2021).

163. Egfjord, A. F.-H. et al. Genomic Steppe ancestry in skeletons from the Neolithic Single Grave Culture in Denmark. PLOS ONE 16, e0244872 (2021).

164. Gnecchi-Ruscone, G. A. et al. Ancient genomic time transect from the Central Asian Steppe unravels the history of the Scythians. Science Advances vol. 7 eabe4414 (2021).

165. Hajdinjak, M. et al. Initial Upper Palaeolithic humans in Europe had recent Neanderthal ancestry. Nature 592, 253–257 (2021).

166. Mao, X. et al. The deep population history of northern East Asia from the Late Pleistocene to the Holocene. Cell 184, 3256–3266.e13 (2021).

167. Papac, L. et al. Dynamic changes in genomic and social structures in third millennium BCE central Europe. Sci. Adv. 7, eabi6941 (2021).

168. Posth, C. et al. The origin and legacy of the Etruscans through a 2000-year archeogenomic time transect. Sci. Adv. 7, eabi7673 (2021).

169. Charlton, S. et al. Dual ancestries and ecologies of the Late Glacial Palaeolithic in Britain. *Nat*. Ecol. Evol. 6, 1658–1668 (2022).

170. Childebayeva, A. et al. Population Genetics and Signatures of Selection in Early Neolithic European Farmers. Mol. Biol. Evol. 39, msac108 (2022).

171. Dulias, K., et al. Ancient DNA at the edge of the world: Continental immigration and the persistence of Neolithic male lineages in Bronze Age Orkney. Proc. Natl. Acad. Sci. 119, e2108001119 (2022).

172. Fischer, C.-E., et al. Origin and mobility of Iron Age Gaulish groups in present-day France revealed through archaeogenomics. iScience 25, (2022).

173. Gretzinger, J. et al. The Anglo-Saxon migration and the formation of the early English gene pool. Nature 610, 112–119 (2022).

174. Maróti, Z. et al. The genetic origin of Huns, Avars, and conquering Hungarians. Curr. Biol. 32, 2858–2870.e7 (2022).

175. Patterson, N. et al. Large-scale migration into Britain during the Middle to Late Bronze Age. Nature 601, 588–594 (2022).

176. Blöcher, J. et al. Descent, marriage, and residence practices of a 3,800-year-old pastoral community in Central Eurasia. Proc. Natl. Acad. Sci. 120, e2303574120 (2023).

177. Koptekin, D. et al. Spatial and temporal heterogeneity in human mobility patterns in Holocene Southwest Asia and the East Mediterranean. Curr. Biol. 33, 41–57.e15 (2023).

178. Moots, H. M. et al. A genetic history of continuity and mobility in the Iron Age central Mediterranean. Nat. Ecol. Evol. 1–10 (2023) doi:10.1038/s41559-023-02143-4.

179. Barrie, W. et al. Elevated genetic risk for multiple sclerosis emerged in steppe pastoralist populations. Nature 625, 321–328 (2024).

180. Gelabert, P. et al. Social and genetic diversity in first farmers of central Europe. Nat. Hum. Behav. 1–12 (2024) doi:10.1038/s41562-024-02034-z.

181. Omrak, A. et al. Genomic Evidence Establishes Anatolia as the Source of the European Neolithic Gene Pool. Curr. Biol. 26, 270–275 (2016).

182. Yang, M. A. et al. 40,000-Year-Old Individual from Asia Provides Insight into Early Population Structure in Eurasia. Curr. Biol. 27, 3202–3208.e9 (2017).

183. Jeong, C., et al. Bronze Age population dynamics and the rise of dairy pastoralism on the eastern Eurasian steppe. Proc. Natl. Acad. Sci. 115, E11248–E11255 (2018).

184. Lamnidis, T. C. et al. Ancient Fennoscandian genomes reveal origin and spread of Siberian ancestry in Europe. Nat. Commun. 9, 5018 (2018).

185. Mathieson, I. et al. The genomic history of southeastern Europe. Nature vol. 555 197-+ (2018).

186. Van De Loosdrecht, M. et al. Pleistocene North African genomes link Near Eastern and sub-Saharan African human populations. Science 360, 548–552 (2018).

187. Flegontov, P. et al. Paleo-Eskimo genetic ancestry and the peopling of Chukotka and North America. Nature 570, 236–240 (2019).

188. Mittnik, A. et al. Kinship-based social inequality in Bronze Age Europe. Science 366, 731–734 (2019).

189. Prendergast, M. E. et al. Ancient DNA reveals a multistep spread of the first herders into sub-Saharan Africa. Science 365, eaaw6275 (2019).

190. Villalba-Mouco, V. et al. Genomic transformation and social organization during the Copper Age–Bronze Age transition in southern Iberia. Sci. Adv. 7, eabi7038 (2021).

191. Villalba-Mouco, V. et al. A 23,000-year-old southern Iberian individual links human groups that lived in Western Europe before and after the Last Glacial Maximum. Nat. Ecol. Evol. 1–13 (2023) doi:10.1038/s41559-023-01987-0.

192. Wang, C.-C. et al. Ancient human genome-wide data from a 3000-year interval in the Caucasus corresponds with eco-geographic regions. Nat. Commun. 10, 590 (2019).

193. Wang, C.-C. et al. Genomic insights into the formation of human populations in East Asia. Nature 591, 413–419 (2021).

194. Wang, K. et al. Middle Holocene Siberian genomes reveal highly connected gene pools throughout North Asia. Curr. Biol. 33, 423–433.e5 (2023).

195. Lipson, M. et al. Ancient West African foragers in the context of African population history. Nature 577, 665–670 (2020).

196. Lipson, M. et al. Ancient DNA and deep population structure in sub-Saharan African foragers. Nature 603, 290–296 (2022).

197. Massilani, D. et al. Denisovan ancestry and population history of early East Asians. Science 370, 579–583 (2020).

198. Nägele, K. et al. Genomic insights into the early peopling of the Caribbean. Science 369, 456–460 (2020).

199. Rivollat, M. et al. Ancient genome-wide DNA from France highlights the complexity of interactions between Mesolithic hunter-gatherers and Neolithic farmers. Sci. Adv. 6, eaaz5344 (2020).

200. Rivollat, M. et al. Extensive pedigrees reveal the social organization of a Neolithic community. Nature 620, 600–606 (2023).

201. Skourtanioti, E. et al. Genomic History of Neolithic to Bronze Age Anatolia, Northern Levant, and Southern Caucasus. Cell 181, 1158–1175.e28 (2020).

202. Skourtanioti, E. et al. Ancient DNA reveals admixture history and endogamy in the prehistoric Aegean. Nat. Ecol. Evol. 7, 290–303 (2023).

203. Teschler-Nicola, M. et al. Ancient DNA reveals monozygotic newborn twins from the Upper Palaeolithic. Commun. Biol. 3, 650 (2020).

204. Yu, H. et al. Paleolithic to Bronze Age Siberians Reveal Connections with First Americans and across Eurasia. Cell 181, 1232–1245.e20 (2020).

205. Yu, H. et al. Genomic and dietary discontinuities during the Mesolithic and Neolithic in Sicily. iScience 25, 104244 (2022).

206. Freilich, S. et al. Reconstructing genetic histories and social organisation in Neolithic and Bronze Age Croatia. Sci. Rep. 11, 16729 (2021).

207. Novak, M. et al. Genome-wide analysis of nearly all the victims of a 6200 year old massacre. PLOS ONE 16, e0247332 (2021).

208. Prüfer, K. et al. A genome sequence from a modern human skull over 45,000 years old from Zlatý kůň in Czechia. Nat. Ecol. Evol. 5, 820–825 (2021).

209. Saupe, T. et al. Ancient genomes reveal structural shifts after the arrival of Steppe-related ancestry in the Italian Peninsula. Curr. Biol. 31, 2576–2591.e12 (2021).

210. Žegarac, A., et al. Ancient genomes provide insights into family structure and the heredity of social status in the early Bronze Age of southeastern Europe. Sci. Rep. 11, 10072 (2021).

211. Yaka, R. et al. Variable kinship patterns in Neolithic Anatolia revealed by ancient genomes. Curr. Biol. 31, 2455–2468.e18 (2021).

212. Zhang, F. et al. The genomic origins of the Bronze Age Tarim Basin mummies. Nature 599, 256–261 (2021).

213. Marchi, N. et al. The genomic origins of the world’s first farmers. Cell 185, 1842–1859.e18 (2022).

214. Scorrano, G. et al. Genomic ancestry, diet and microbiomes of Upper Palaeolithic hunter-gatherers from San Teodoro cave. Commun. Biol. 5, 1262 (2022).

215. Wohns, A. W. et al. A unified genealogy of modern and ancient genomes. Science 375, eabi8264 (2022).

216. Mattila, T. M. et al. Genetic continuity, isolation, and gene flow in Stone Age Central and Eastern Europe. *Commun*. Biol. 6, 1–13 (2023).

217. Penske, S. et al. Early contact between late farming and pastoralist societies in southeastern Europe. Nature 620, 358–365 (2023).

218. Posth, C. et al. Palaeogenomics of Upper Palaeolithic to Neolithic European hunter-gatherers. Nature 615, 117–126 (2023).

219. Browning, B. L. & Browning, S. R. Detecting Identity by Descent and Estimating Genotype Error Rates in Sequence Data. Am. J. Hum. Genet. 93, 840–851 (2013).

220. Mallick, S. et al. The Allen Ancient DNA Resource (AADR): A curated compendium of ancient human genomes. 2023.04.06.535797 Preprint at 10.1101/2023.04.06.535797 (2023).

221. Greenbaum, G., Rubin, A., Templeton, A. R. & Rosenberg, N. A. Network-based hierarchical population structure analysis for large genomic data sets. Genome Res. 29, 2020–2033 (2019).

222. Mallick, S. et al. The Allen Ancient DNA Resource (AADR) a curated compendium of ancient human genomes. Sci. Data 11, 182 (2024).

223. Lawson, D. J., Hellenthal, G., Myers, S. & Falush, D. Inference of Population Structure using Dense Haplotype Data. PLOS Genet. 8, e1002453 (2012).

224. Pinotti, T., McColl, H., Sikora, M. & Czech, L. ybyra: Y-chromosome haplogroup calling using a tree-based scoring method. 2025.11.20.689455 Preprint at 10.1101/2025.11.20.689455 (2025).

