## Supplementary figures and images for "Migrations facilitated the cultural diffusion of megaliths in Neolithic Europe"

### Supplementary Fig. S3.10

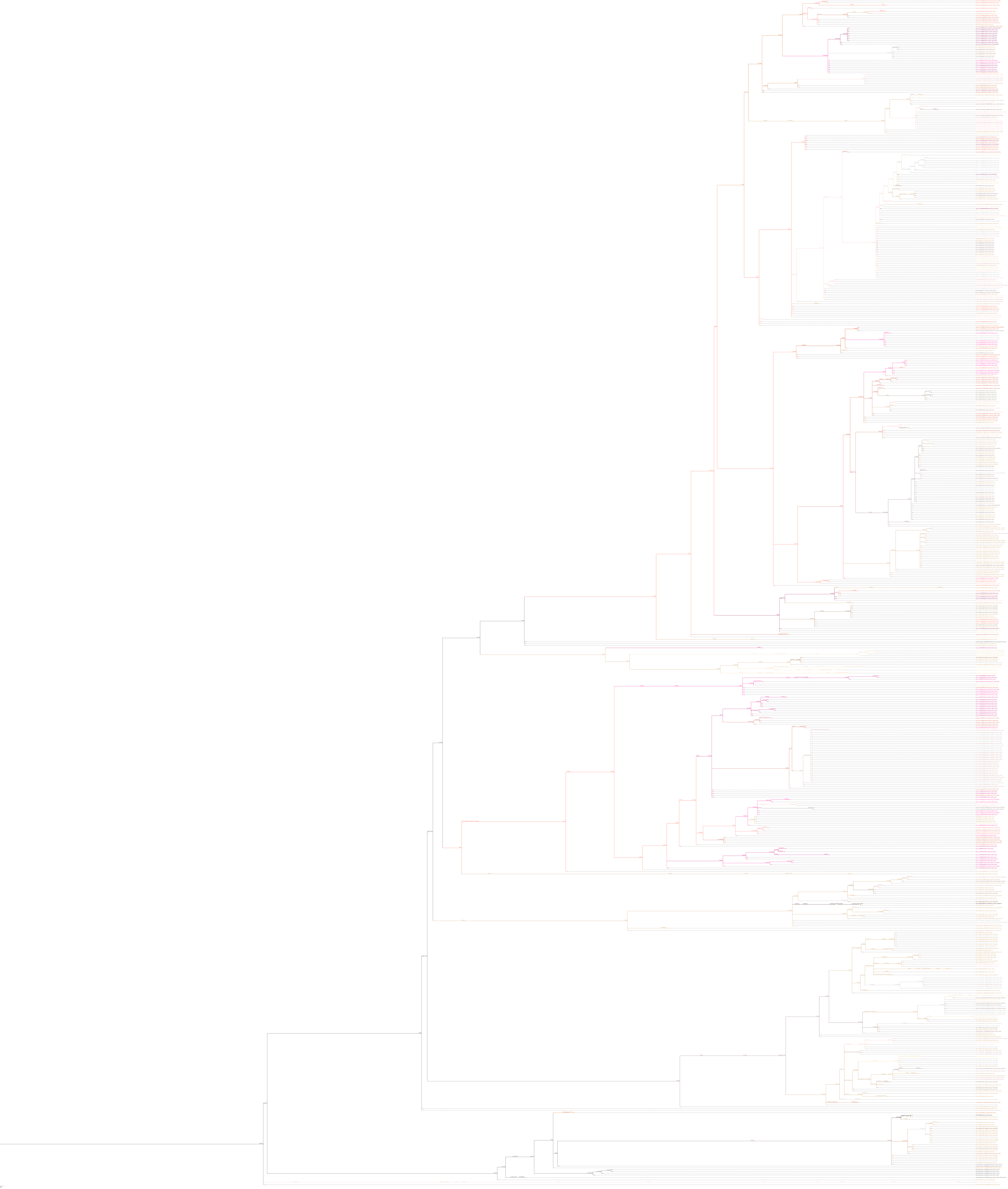

### Supplementary Fig. S3.12

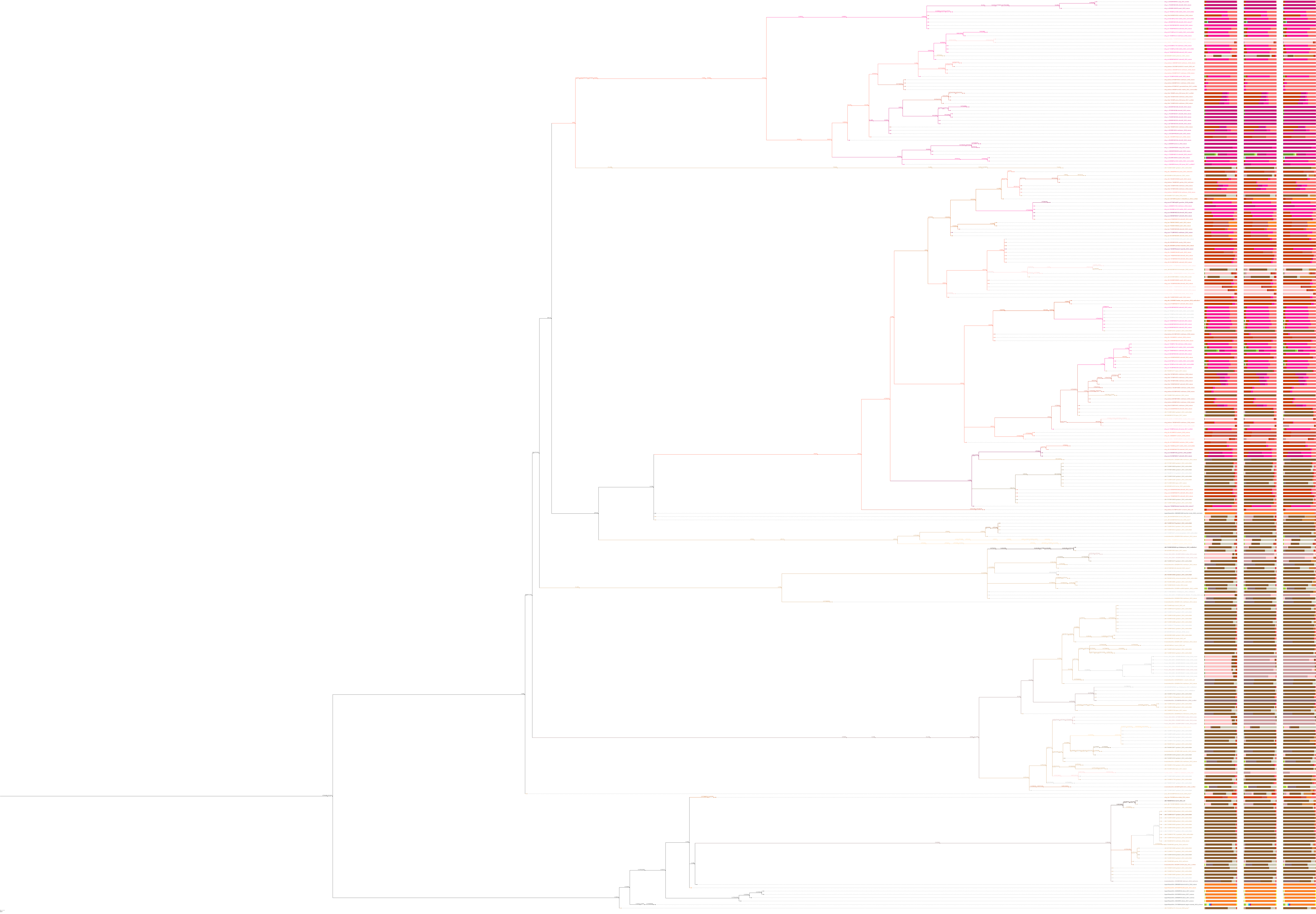

### Supplementary Fig. S3.14

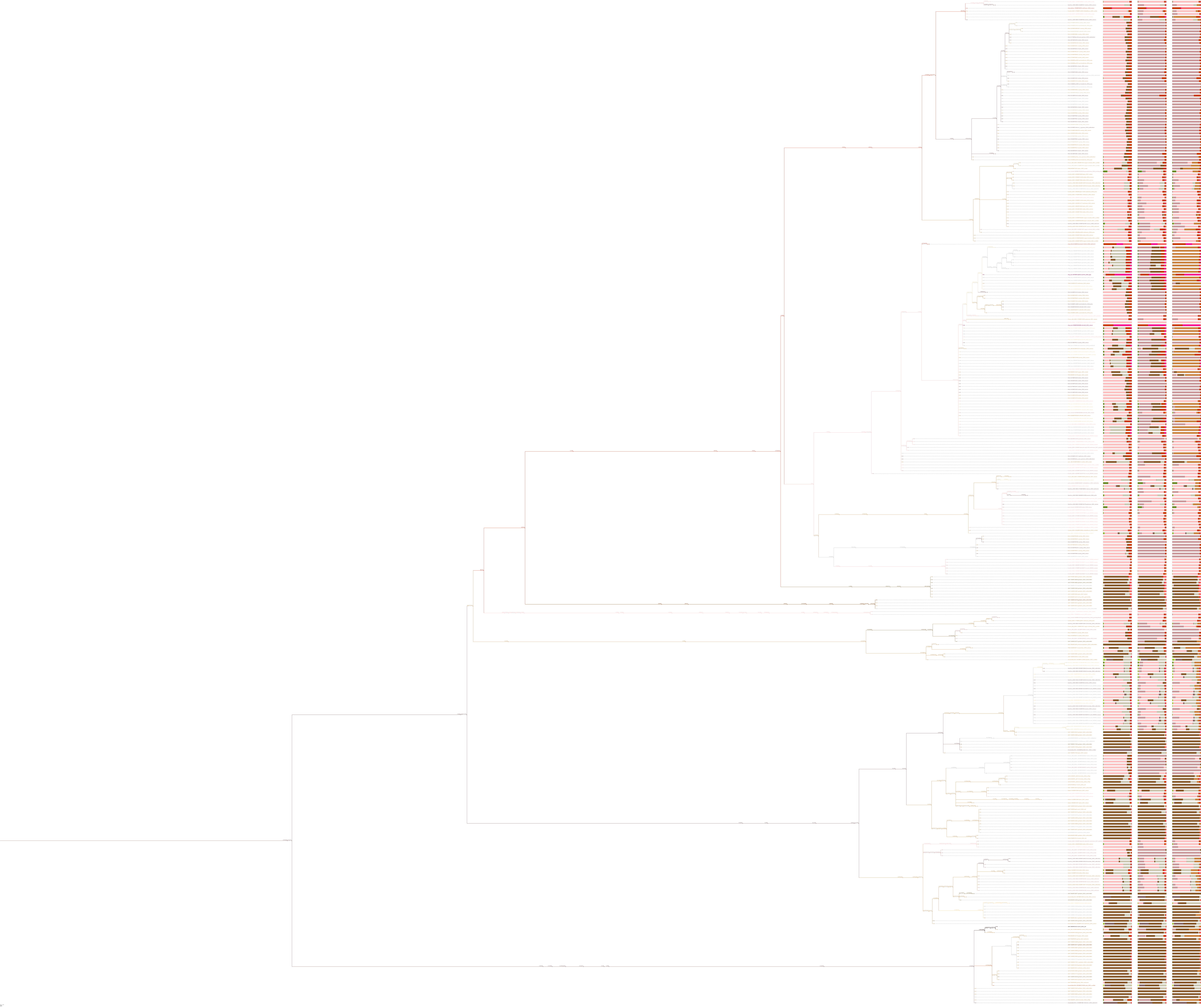

### Supplementary Fig. S3.16

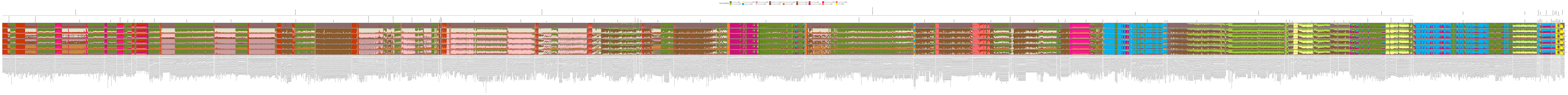
