## Supplementary Notes S1-4 for "Migrations facilitated the cultural diffusion of megaliths in Neolithic Europe"

#### Supplementary Notes S1-S4

Supplementary Note S1. Introduction 4
1.1 Migration or cultural diffusion? The emergence and spread of European megalithic traditions 4
1.2 Structure and technical notes 9
Supplementary Note S2. The archaeological context of the Cardial Neolithic expansion in Europe 10
2.1 The Neolithic expansion from Anatolia and the origin of the LBK and Cardial expansions into Central and Mediterranean Europe 10
2.1.1 Initial expansion from Anatolia 10
2.1.2 The origins of the LBK 12
2.1.3 The origins of Impressed Ware: the Adriatic and Italy 14
2.2. The Neolithic transition in the Central Mediterranean and the islands of Sardinia and Corsica 8000-6000 BP 19
2.2.1 Early Neolithic and pioneering phase 20
2.2.2 The Middle Neolithic and earliest funeral architecture 20
2.2.3 The earliest megaliths and the passage grave tradition 23
2.3. The Neolithic sequence for southern France and farming expansion northwards: from Early to Middle Neolithic 25
2.3.2 Genomic evidence 28
2.3.3 The Chasséen 30
2.4 The Neolithic transition in Northern and Northwest France 35
2.4.1 Pioneering settlers in the Early Neolithic (7000-6700 cal BP) 36
2.4.2 Monumental graves, stone alignments, earliest megaliths and long distance green stone trade in the Early Middle Neolithic (6700-6300 cal BP) 38
2.4.3 The Recent Middle Neolithic (6300-5800 cal BP) Societal and cultural change and innovations 43
2.4.4 Genetic results and societal change 46
2.5 The Iberian peninsula and the Neolithic transition 8000-6000 BP 49
2.5.1 The earliest Cardial movements along the Mediterranean coast and inland to the Northeast 49
2.5.2 The first Cardial groups in Western Iberia 52
2.5.3 Interaction between Cardial and hunter-gatherer groups 53
2.5.4 Neolithisation processes in the Northern regions and the Atlantic Iberian coast 53

2.6 The farming colonisation of Britain and Ireland 61
2.6.1 Early Neolithic subsistence patterns 65
2.6.2 Social and cultural change 67
2.7 The Michelsberg Culture 69
2.7.1 Genetics 72
2.7.2 Settlement and subsistence 73
2.8 The Neolithic settling of Scandinavia 76
2.9 Globular Amphora Communities (GAC) 82
2.9.1 Ancient DNA results 84
2.10 Neolithic expansion and megaliths in Northwestern Africa 85
2.10.1 Pioneering phase and evidence for early Neolithic settlers 86
2.10.2 Megaliths 87
2.10.3 Genetic results 88
2.11 Circulation and prospection of green stone and obsidian in Western Europe 8000-6000 BP 89
2.11.1 Introduction: the green stone trade in Europe 89
2.11.2 Jadeite and jadeite trade 8000 BP-5000 BP 90
2.11.3 Variscite and variscite trade in Europe 8000-5000 BP 93
2.11.3.1 The early variscite network in Western Europe 8000-6000 BP 94
2.11.3.2 The later variscite network in Western Europe, 6000 to 4000 BP 95
2.11.4 Obsidian in the West Mediterranean 96
2.11.5 A Bayesian statistical framework applied on the radiocarbon dates for the greenstone exchange networks in Europe 104
2.11.5.1 Introduction 104
2.11.5.2 A Bayesian statistical framework for the jadeite exploitation at the Monte Viso 104
2.11.5.3 Monumental tombs in Northwestern France with a radiocarbon sequence and jadeite and variscite deposits 111
2.11.5.4 Further Variscite contexts with radiocarbon dates from Northwest France 116
2.11.5.5 Mediterranean contexts with radiocarbon dates and variscite beads 120
2.11.5.6 West Iberian contexts with radiocarbon dates and variscite beads 130
Supplementary Note S3. The newly sampled human remains 136
3.1 Introduction 136
3.2 The sampling strategy 137
3.2.1 Sardinia 137
3.2.2 Portugal 141
3.2.2.1 Cova da Onça shell midden and the Sintra region (Tholos of Praia das Maçãs, the Dolmen of Pedras da Granja, and the prehistoric burial site of Samarra) 141
3.2.2.2 The Northern bank of the Tagus River in the Lisbon region 142
3.2.3 The sampling documentation with 3-D modelling 142
3.3. Methods 150
3.3.1 Population structure of Mesolithic and Neolithic Europe 150

3.3.2 Testing the association between ancestry components and the emergence of megalithic monuments 151
3.3.3 Y-chromosome haplogroup assignment 161
3.4 The archaeological context of the Sardinian samples 168
3.4.1 The Domu de janas II necropolis at San Benedetto -Iglesias, Cagliari 168
3.4.1.1 Archaeological data 168
3.4.1.2 Previous genetic studies on Sardinian material 172
3.4.1.3 List of sites included and brief description: 173
3.4.1.4 Chronological position of San Benedetto relative to previous Neolithic aDNA data 177
3.5 The archaeological context of the Portuguese samples 179
3.5.1 The Tholos of Praia das Maçãs 179
3.5.2 Dolmen Pedras da Granja 183
3.5.3 Samarra 185
3.5.4 Bom Santo Cave, Lisbon, Portugal. 188
3.5.6 Trigache 4, Lisbon, Portugal 195
3.5.7 Monte Abraão, Lisbon, Portugal 197
3.5.8 Cova da Onça (Tagus Valley, Portugal) 199
3.6. Radiocarbon Dates for the newly obtained genetic results 201
3.6.1 AMS dating of human bones 201
3.6.2 Results 202
3.6.3 The calibrated radiocarbon ages 203
Supplementary Note S4. Vegetation, sea-level and climatic change in Iberia and Northern France 208
4.1 Vegetation 208
4.2 Sea level 212
4.3 Climate 213
4.4 Summary 216
References 221

### Supplementary Note S1. Introduction

*Bettina Schulz Paulsson*

#### 1.1 Migration or cultural diffusion? The emergence and spread of European megalithic traditions

Megaliths (from the Greek *mégas*, large, and *líthos*, stone) are a global prehistoric phenomenon comprising tombs and standing stones, the latter occurring either as isolated monuments or in stone circles and alignments, such as Stonehenge and Carnac. In Europe, the focus of this study, megaliths were constructed primarily during the Neolithic and Copper Age (6700–4500 BP), particularly along or in relative proximity to the Atlantic and Mediterranean coasts. Approximately 35,000 monuments are commonly cited[1] but this is plausibly a conservative estimate and no comprehensive inventory yet exists. The actual number of megaliths may exceed 100,000.

Despite their vast geographical distribution, the range of types and the architecture of European megaliths are remarkably similar. Dolmens, passage graves, gallery graves, standing stones, stone circles, and stone alignments occur across both the Atlantic façade and the Mediterranean region, and many burial monuments share an east or southeast orientation (Supplementary Fig. S1.1).

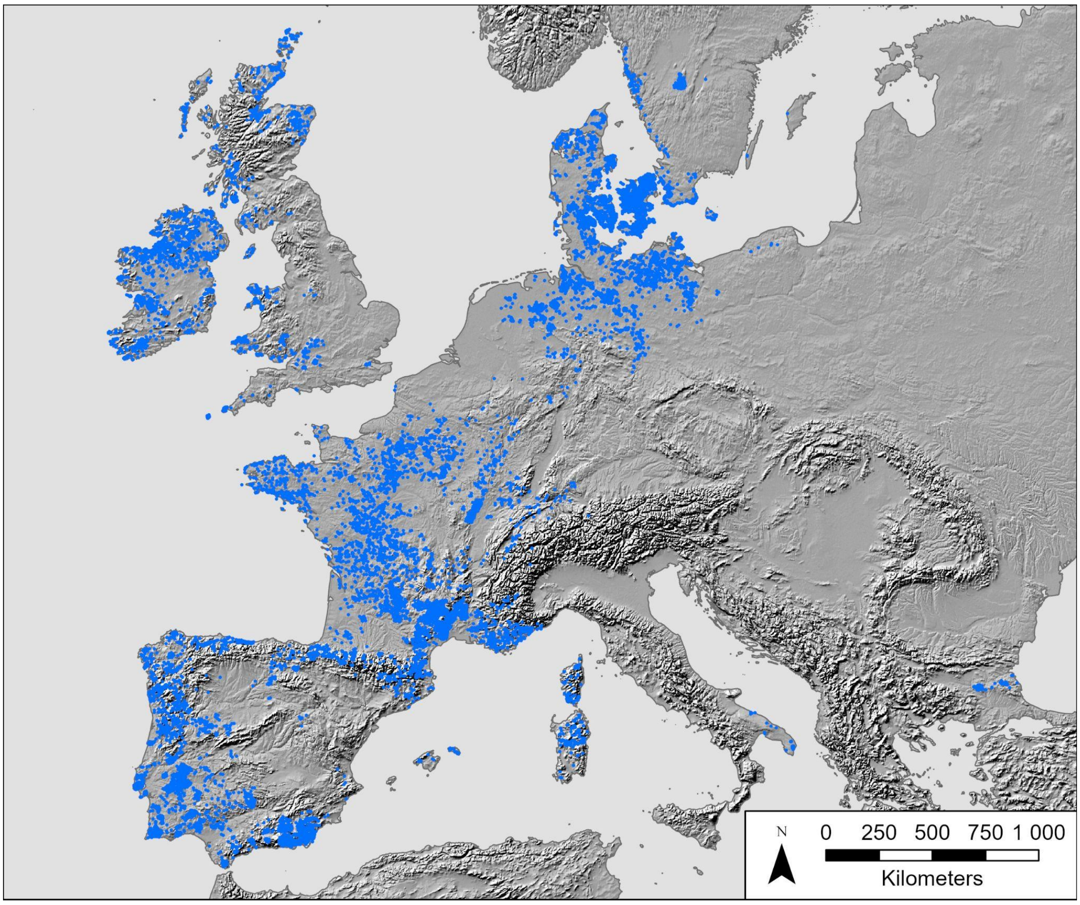

**Supplementary Fig. S1.1. The spread of megaliths in Europe[2].** The map shows approximately 17,000 published megalithic sites, representing only a fraction of the total number of monuments but broadly reflecting their overall distribution.

These similarities have long raised the question of whether megalithic traditions developed independently in different regions or spread through the movement of people, ideas, institutions, and technologies.

Early studies, based largely on monument typology, generally favoured diffusion or migration models. Montelius[3,4] proposed the spread of a megalithic concept from the eastern Mediterranean to Atlantic and northern Europe, whereas Kossinna[5] and Sprockhoff[6] argued for migration from northern Europe. Childe proposed that shared religious ideas and megalithic traditions spread by partial maritime exchange through small groups of pioneers or "missionaries"[7–10], an idea that Daniel[11,12] later extended by linking megalithic expansion to maritime colonisation and the search for copper.

The introduction of radiocarbon dating fundamentally changed the debate. Renfrew[13,14] rejected single-origin models and proposed several independent centres of megalithic emergence and development. He interpreted similar monuments as convergent responses to comparable social and economic challenges associated with the introduction of farming as the dominant subsistence strategy. This perspective was subsequently reinforced by models emphasising regional developments in social organisation, territoriality, and competition over resources rather than long-distance migration or cultural transmission[15,16].

Radiocarbon syntheses later identified north-western France, western Iberia, and possibly Ireland as the earliest centres of megalith construction followed by expansion into western, northern, and central Europe[17,18]. Subsequent research focused largely on identifying regional variability and describing and explaining local monument traditions (e.g. refs [5,19–22]).

Bayesian modelling of a substantially expanded radiocarbon dataset provided a new more detailed and robust chronological framework for European megalith construction. The results identified north-western France as the epicentre of megalith construction and supported a rapid maritime expansion of megalithic traditions along the Atlantic façade during the second half of the fifth millennium BC, reopening the debate over the relative roles of demographic processes and cultural transmission as the mechanism explaining the spatio-temporal pattern of megalith distributions[1,23].

Recent archaeological, chronological, archaeometric, and genomic studies increasingly favour interconnected interaction networks rather than independent regional trajectories, including both long-distance maritime connectivity and regional exchange systems[24–30]. However, the relative contributions of migration, cultural transmission, and the local continuity of megalithic traditions remain unresolved.

Ancient genomics provides an independent means of testing these competing hypotheses. Large-scale migration predicts substantial genetic continuity among geographically distant megalithic communities, whereas cultural diffusion predicts the spread of monument traditions between genetically distinct populations. Intermediate scenarios involving limited movement of maritime pioneers or specialist groups would be expected to leave more subtle genetic signatures.

The present study provides the first comprehensive, continent-wide genomic analysis of Neolithic populations, encompassing 2,963 ancient individuals, including 46 unpublished genomes, and comparing Neolithic megalithic communities with contemporaneous non-megalithic populations. By integrating genomic and archaeological evidence, we directly evaluate the demographic dynamics underlying the emergence and spread of megalithic traditions across Europe between 6700 and 4000 BP.

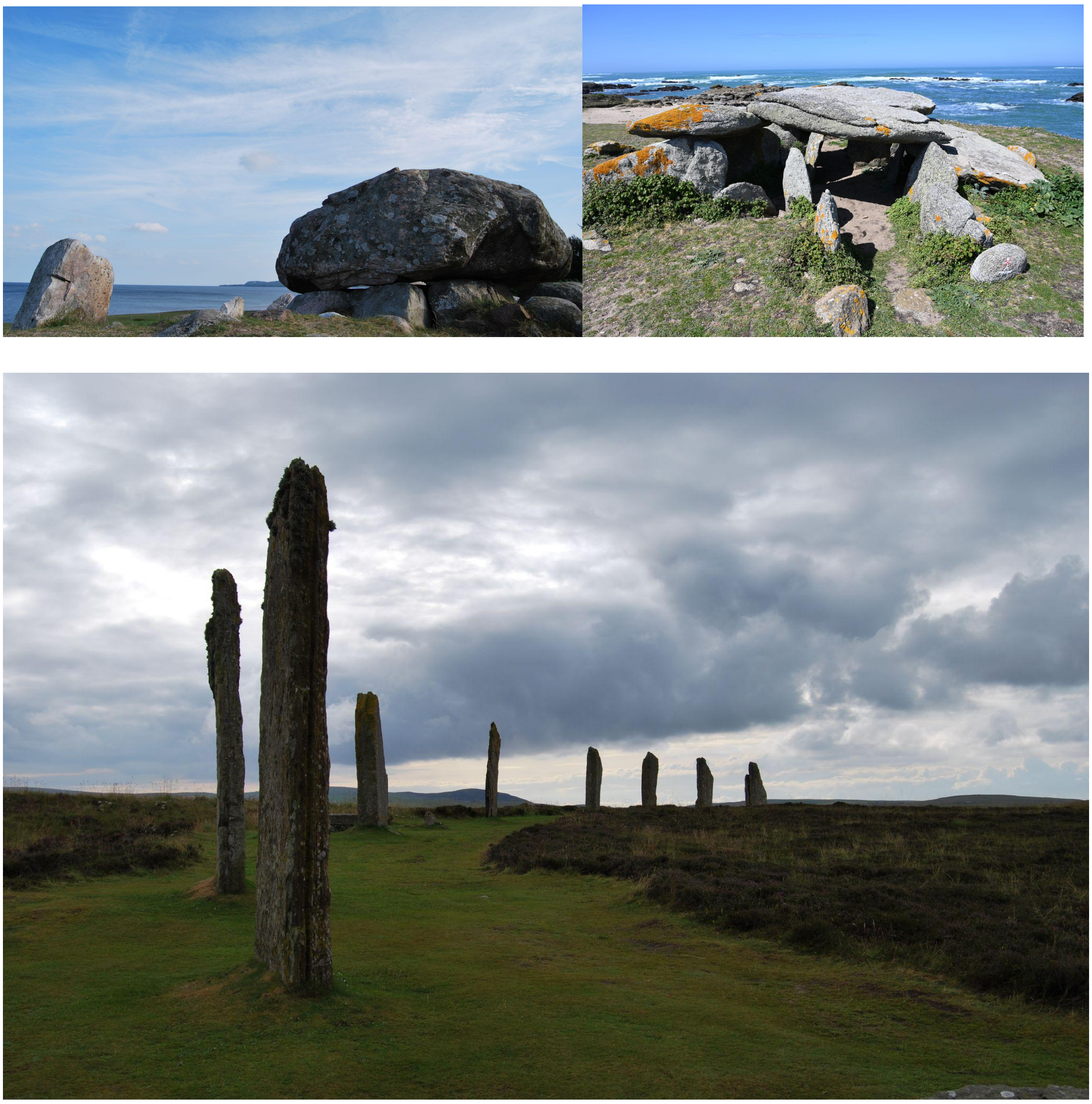

**Supplementary Fig. S1.2. Megalith types.** Above: The dolmen Havangdösen, at the sea in Southeastern Sweden (left) and a passage grave at La Planche à Puare on the French Atlantic island Ile d'Yeu (right). Below: Standing stones arranged in a circle at Ring of Brodgar, Orkney, Scotland. All over Europe we find similar varieties of megalith architecture. Photos: B. Schulz Paulsson.

#### 1.2 Structure and technical notes

This Supplementary Information is organised into three complementary sections.

1) The first section (2.1–2.11) synthesises the archaeological and chronological context of all individuals included in the genetic analyses through an extensive review of recent literature on the Neolithic in Europe and by recalibrating relevant radiocarbon dates. The radiocarbon results are reported in BP and rounded to the nearest century. For selected key sites we obtained higher temporal resolution either by recalibration or from Bayesian chronological modelling (see 2.11.5). These results are partly reported in cal BC. The models and the calibrated data presented in the analysis were generated with the OxCal v4.4 software[31,32] and the Intcal19 calibration data[33].

2) The second section (3.1–3.6) presents the newly generated ancient genetic data for this study, including the sampling strategy, genetic analyses, archaeological contexts, and radiocarbon dating.

3) The third part (4.1–4.4) consists of an outline of the environmental history of Iberia and Northern France, including vegetation history, sea-level changes, and climate history.

### Supplementary Note S2. The archaeological context of the Cardial Neolithic expansion in Europe

*Bettina Schulz Paulsson & Stephen Shennan*

#### 2.1 The Neolithic expansion from Anatolia and the origin of the LBK and Cardial expansions into Central and Mediterranean Europe

*Stephen Shennan*

##### 2.1.1 Initial expansion from Anatolia

It is now well established that farmers and farming spread into southeastern, central and Mediterranean Europe by a process of demographic expansion from the Anatolian-Aegean region[34–39] but how and where this population formed has remained unclear, with some scholars pointing to a movement along the southern Anatolian coast from the Levant and others to a central Anatolian origin[40]. Recent aDNA results[41] have thrown new light on this. It is now apparent that this gene pool formed in western Anatolia after 9000 BP as a result of expansion from further east, possibly Upper Mesopotamia, and admixture with local hunter-gatherer groups, some of whom had already adopted Neolithic cultural elements.

The broad chronological pattern is clearly seen by examining the relevant radiocarbon dates[42], which indicate that expansion out of Central Anatolia to western Anatolia, Greece and the Aegean began around 9000 BP. It reached the coastal zone of northern Greece by ~8500 BP[43]. It appears likely that the expansion of farmers into the Balkans took place via northern Greece rather than NW Anatolia[44]. It is in any case now apparent that this expansion was significantly later than in Anatolia and the Aegean. The site of Vlaho in northern

Macedonia dates to ~8300 BP and the farming frontier had reached the Danube by ~8200 BP, expanding further to the southern end of the Carpathian Basin 100-200 years later[45].

The genomic picture for the Balkan Neolithic overall indicates an average of 98% Anatolian ancestry with very little reproductive inter-mixing with local hunter-gatherers, except for the site of Lepenski Vir[38,46,47].

Studies of early 9$^{th}$ millennium BP individuals from the Starčevo and Kőrös cultures at the northern end of the Balkan farming expansion in Hungary[37] showed that five out of six individuals were essentially no different from Neolithic Anatolians though one had ~8% Western Hunter Gatherer genetic ancestry, resulting from an admixture event in the previous three to five generations.

In terms of the animal economy the picture is very clear. While in Greece the faunal spectra are largely dominated by sheep/goat, in the Balkans to the north they are much more diverse[48,49]. Ovicaprines are far less dominant and there is a stronger emphasis in many places on cattle, a pattern explicable at least in part by lower temperatures and higher rainfall[50]; there is also a greater terrestrial wild component. As regards the domesticated crops, the founder cereals einkorn, emmer and hulled barley are present in the majority of sites in SE Europe, as in SW Asia, while free-threshing wheat and naked barley are also present in at least 50% of the sites in the two regions, based on the data analysed by Colledge et al.[51]. SE Europe also has the full set of founder crop domestic pulses that are found in SW Asia: pea, lentil, bitter vetch, chick pea and grass pea (an early addition to the founder crops).

Traditionally, the main feature characterising the first farming communities of the Balkans and linking them to their Aegean and Anatolian ancestors has been their pottery, which has long provided the basis for defining the material culture patterns of the EN: the Starčevo culture in Serbia and adjacent areas, the Körös culture in Hungary, the Criș in Romania and the pre-Karanovo and Karanovo I in Bulgaria. Despite the local variations, for example in the frequency of painted pottery and the style of the painting, the pottery shows strong patterns of stylistic similarity linking the region from the southern Carpathian Basin in the north to Macedonia and Thrace in the south.

In summary, just as there was a rapid spread of farming from Central Anatolia and adjacent areas to western Anatolia and the northern Aegean, in less than 500 years, by 8500 BP, ~300-400 years later there was another spread from the northern Aegean zone to north of the Danube by ~8100 BP, visible in summed radiocarbon probabilities as a population increase (e.g. refs [45,52,53]). This Balkan EN shows a set of common features in material culture, domestic plant and animal subsistence and domestic animal genetics that links it closely to the Aegean-Anatolian region and demonstrates continuing links well into the 6$^{th}$ millennium BC (6$^{th}$ millennium BP)

##### 2.1.2 The origins of the LBK

However, the northward demographic expansion was not sustained. The central Balkans summed radiocarbon probability proxy sees a shift from boom to bust ~8000 BP and then another boom-bust sequence from ~7800-7500 BP[52], paralleling a similar boom-bust pattern in the Carpathian Basin with the arrival there of farming and the characteristic first farmer cultural traditions[54]. In fact, all the evidence points to a massive drop in population, if not an occupation gap, in the central Balkans and southern Carpathian Basin after ~7500 BP[55–57].

Thus, after arriving at a latitude of ~45° north at ~8000 BP, the northern farming frontier remained largely static in the Carpathian Basin[58]. A new phase of northwestward expansion only began after 7500 BP, probably ~7350 BP[56]. It is likely that this frontier corresponds to the limit of the Mediterranean seasonality conditions to which the wild ancestors of the crops and their domestic descendants were genetically adapted, and the ensuing delay reflected the time required for the evolution of crop varieties that would be successful at latitudes with a lower number of growing-degree days[59].

It is during this static phase of apparently low population that the Linear Pottery Culture (LBK, from its German name), which was responsible for the spread of farming into Central Europe, began to develop in the western Carpathian Basin. Work in recent years has shown that Starčevo occupation reached here, the northwest limit of its distribution, ~7800 BP [60] and on its northern fringes the sites of Brunn/Wolfholz on the outskirts of Vienna[61] and Szentgyörgyvölgy-Pityerdomb[62] in the Lake Balaton area seem to represent formative sites of the LBK, overlapping with the local late Starčevo, ~7500 BP. The site of Alsónyék in southern Transdanubia, just outside the formative area, has a Starčevo occupation beginning just after 7800 BP and continuing until ~7500 BP. It is then abandoned and a new settlement lasting over 300 years is founded on the same site ~5300 BP by people using early (but not formative) LBK pottery and building LBK longhouses[60].

Recent formal modelling suggests that the rapid LBK expansion that ensued did not begin until the 54$^{th}$ century BC (~7300 BP)[60,63]. From ~7350 BP we see another massive demographic expansion. On the basis of an extrapolation from local survey data Oross et al.[56] estimate that there were already ~285 LBK sites in the western Carpathian Basin in the 54$^{th}$ century BC (~7300 BP), increasing to nearly 2400 by the late 6$^{th}$ millennium BC (7500-7000 BP). But this was not just a local increase. At the same time there was a very rapid demographic expansion carrying farming westwards across Central Europe as far as the river Rhine, over 1000 km away, and northwards into north-central Poland, on the edge of the north European plain, in less than 150 years. By 7000 BP it had reached Normandy in the west and Ukraine, nearing the coast of the Black Sea, in the east. Across the whole area its archaeological record is remarkably uniform, not just in the architecture of the houses but also in the pottery, the crop-based agriculture and in its preferences for very specific conditions of soil, temperature and precipitation[39].

The genomic evidence shows that the LBK expansion was another population dispersal of genetic descendants of Aegean-Anatolian farmers with continuing very low levels of hunter-gatherer genetic admixture, which had probably already occurred not long after 8000 BP[37,47]. However, in the obscure phase in the northern Balkans and the Carpathian Basin in the mid-6$^{th}$ millennium BC (~7500 BP) a major cultural transformation had taken place. In fact, there were two. The Early Neolithic Starčevo tradition split into the LBK to the north and the Vinča culture to the south, in a process of cultural phylogenesis[56,63,64], with very distinct traditions in ceramics and figurines but the common appearance of longhouses.

##### 2.1.3 The origins of Impressed Ware: the Adriatic and Italy

As we have seen above, even though farming communities had reached northern Greece and NW Anatolia before 8500 BP it was the best part of 400 years later before they reached the southern Carpathian Basin. The same is true of the spread west into Adriatic Italy and beyond, indicated by the occurrence of Impressa pottery, which began around 8000 BP. It is important to note though that this is at least 500 years earlier than the spread of the LBK into Central Europe described above, presumably because Mediterranean environments were similar to those in which the initial plant domesticates evolved, so they did not face the adaptive problems posed by conditions north of 45° latitude.

The best evidence so far for its starting point comes from the site of Sidari, on Corfu in NW Greece. Berger et al.[65] show that Mesolithic contexts on the site are dubious or mixed and that its earliest Neolithic occupation dates to ~8400-8200 BP. Its ceramic assemblage is connected to the Albanian-Macedonian ceramic complex characterised by the presence of impressed and painted pottery found, for example, at the site of Mavropigi, one of the earliest Neolithic sites in northern Greece, dated to the final third of the 9$^{th}$ millennium BP[66]. In fact, in a major review of the origins and chronology of Mediterranean Impressed Ware, Binder et al. propose that these mixed painted and impressed ceramic assemblages formed the common foundation of the Impressa and Starčevo traditions that went on to develop in parallel in the following centuries, the former in the Adriatic area, the latter in the Danube basin, with evidence of occasional interactions, indicated, for example, by the presence of both Carpathian and Lipari obsidian at some sites in Croatia (pg. 62 in ref [66]).

Three sites with Archaic Impressa pottery most probably dating to the interval 8050-7900 BP have been identified on the southern Dalmatian coast – Zemunica, Rašinovac and Pokrovnik – and there are others that are not well dated. The earliest Impressa layer at Sidari also dates to this interval and follows a hiatus, with indications of erosion associated with the 8.2 ka event. On the other side of the Adriatic, in south-east Italy, only the site of Pulo Fondo Azzollini (Pulo di Molfetta) is well dated to this phase, though in Calabria the site of Favello may begin at the end of it and the same may true of the Grotta Continenza on the Adriatic side of Central Italy, which raises doubts about the conventional view that this region was settled in a second phase of colonisation (pg. 63-64 in ref [66]), though north of Apulia the majority of sites begin no earlier than 7900-7750 BP.

At Rašinovac and Pokrovnik, as well as Pulo Fondo Azzollini and Coppa Nevigata in Apulia, Archaic Impressa is found in association with so-called figulina pottery, an extremely fine ware of varying colours, sometimes slipped and with painted decoration. Its occurrence at these early sites provides further support for the argument that the Adriatic Impressa derives from the Albanian Macedonian assemblages of the preceding period discussed above (pg. 64 in ref [66]). Similarities in vessel construction techniques point in the same direction[67]. In the following phase, 7900-7750 BP, Archaic Impressa continues in central and southern Dalmatia and expands further north, with ongoing occupation at sites such as Pokrovnik and Zemunica and new sites such as Konjevrate and Crno Vrilo (pg. 64 in ref[66]). In the northeast Adriatic the earliest farming dates are ~7500 BP and by this time the style of the pottery had changed to that recognised as the Danilo culture, the local successor to the Impressed Ware.

On the east coast of the Adriatic, the earliest sites with pottery and domestic animals, overwhelmingly caprines, and with very little hunting[68], are caves, and there is generally a time gap between Mesolithic and Neolithic layers, reflected in a very abrupt change from layers with wild fauna to domestic assemblages. So far, the earliest open farming settlements, with evidence of cereals and the same faunal assemblages as the cave sites[68], date to ~7900-7800 BP. In the light of this evidence and given that the karst limestone terrain of much of coastal Dalmatia is more suitable for herding than cereal agriculture, it has been suggested that the first farming colonists were indeed mobile herders. Comparisons of carbon and nitrogen isotope values between Mesolithic and Neolithic skeletons show that there is a much lower marine component in Neolithic diets[69].

Neolithic occupation is most strikingly documented in the Tavoliere plain of Apulia, where it is characterised by the presence of a large number of ditched settlements, over 500 in total (see e.g. fig. 8 in ref [70]); this settlement pattern lasts around 1000 years and is also found in the Matera plain immediately to the south in Italy's instep. A review of the radiocarbon evidence in the Tavoliere and adjacent areas of SE Italy[71] shows a common pattern, with the number of settlements rising to a peak ~7600-7500 BP then gradually declining, and the Tavoliere apparently being abandoned by 7000 BP. The population pattern seems to reflect climatic factors: the region experienced relatively wet conditions for the first half of the 8th millennium BP but after that drier conditions became increasingly marked in the second half, continuing until ~6600 BP.

The crop pattern is dominated by emmer, einkorn, barley and legume species, which vary slightly in their frequency over time[71,72]. There are not many well-documented faunal assemblages but the Impressed Ware site of Torre Sabea on the coast of the Salento hills area of SE Italy has produced data of good quality, indicating an overwhelming emphasis on domestic animals, which make up 94% of the mammal fauna. The majority are caprines (62%), with 23% cattle and 6% pig. The age distribution suggests that both caprines and cattle were used for milk production(pg. 162 in ref [73]) and this suggestion has been confirmed by the finding of dairy fat residues in vessels from the site of Pulo Fondo Azzolini (Pulo di Molfetta) in Apulia at the very beginning of the Impressed Ware sequence ~8050-7930 BP[74]. Stable carbon and nitrogen isotope studies of human skeletons from SE Italy have produced evidence of a minor marine component in the diet of individuals buried near the coast but otherwise diets were based overwhelmingly on terrestrial, it is safe to say agricultural, resources[75].

North of Apulia, the Impressed Ware site of Colle Santo Stefano in the Abruzzo region of central Italy, with radiocarbon dates between 7840 and 7460 BP, has produced a faunal assemblage dominated by domestic animals, largely sheep/goat. Ceramic residue analyses showed that the majority of vessels tested contained ruminant adipose fats, from meat, while pork fats were also found, but dairy fats were detected as well in a number of vessels, confirming that here too dairying and milk processing were part of the subsistence practices of the earliest Impressed Ware farmers[76].

The waterlogged Impressed Ware site of La Marmotta, near Rome, with tree-ring dates between ~7690 and 7230 BP has provided evidence of the technology that made the West Mediterranean farming expansion possible, a dugout canoe more than 10 m in length and a metre wide, made from a large oak, which would have been capable of carrying animals as well as people. As Robb (pg. 255 in [77]) points out, the fact that it does not have a keel would have made it unstable in open sea conditions, but seagoing versions with outriggers could have existed and an experimental replica successfully made the voyage along the coast from central Italy to Portugal. In any event, it is now clear that the initial expansion of farming and farmers into the west Mediterranean Basin was based on maritime movement.

In material culture terms, the Impressed Ware that begins the sequence develops more complex forms of geometric impressed and incised decoration, in particular the Guadone tradition, joined in the mid 8$^{th}$ millennium by painted wares, though recent Bayesian radiocarbon modelling(pg. 72 in [66]) suggests that the traditionally defined ceramic phases actually overlap. In any case, over the period 7750-7450 BP most sites in Apulia include a painted component and there is considerable diversity in the assemblages from one site to the next. However, recent work has revealed a major difference in ceramic technology between Impressa assemblages from sites east and west of the Appennines that is already present early in the sequence and continues down to ~7600 BP[67,78]. On the Adriatic side of Italy, as already noted, the ceramic technology uses a coil-based method similar to that of the Albanian-Macedonian Early Neolithic sites that are believed to be ancestral to the Impressa. On the western side the technology uses a Spiralled Patchwork technique, for example at the early site of Saracena - San Michele in Calabria (pg. 129 in ref [78]), which is also found at the earliest sites in southern France and Liguria, as well as Corsica (see relevant section). The contrast is true for all Impressa sites in their respective areas, regardless of their date.

Moreover, there is no evidence of any transitional assemblages. In addition to the contrast in vessel-forming techniques there are also differences in vessel shape and decoration between the two traditions. Gomart et al.[78] propose that these represent two distinct communities of practice or social networks but the explanation for this difference remains unclear.

The genomics results that have become available[38,79–82] indicate that the spread of Impressed Ware is associated with the spread of people of Anatolian farmer ancestry. This is true of the three individuals from Zemunica in Croatia, dated to ~8050-7750 BP, four individuals from Grotta Continenza in the Abruzzo dating from ~8050 to 7450 BP; and two individuals associated from the site of Kargadur in Istria dated to ~7750-7450 BP. Seven individuals from the site of Uzzo in Sicily dated to ~5,460-5,220 cal BC (7,410–7,170 BP) have substantial Anatolian Farmer ancestry and are similar to early farmers from the Balkans, Hungary, and Anatolia, but not Iberia, with a small hunter-gatherer component, probably of local Sicilian origin[80]. A further four Middle Neolithic individuals from Sicily also have overwhelmingly Anatolian farmer ancestry with a small WHG contribution[79]. Finally, the Middle Neolithic individuals from the central Italian site of Ripabianca di Monterado (5465-5214 cal BC, 7415–7164 BP) reported by Antonio et al.[81] are also mainly of Anatolian farmer ancestry with a small proportion of local hunter-gatherer.

#### 2.2. The Neolithic transition in the Central Mediterranean and the islands of Sardinia and Corsica 8000-6000 BP

*Bettina Schulz Paulsson*

##### 2.2.1 Early Neolithic and pioneering phase

The Cardial Neolithic reached Corsica and Sardinia no later than 7400–7300 BP; radiocarbon-dated charcoal suggests the possibility of even earlier settlement (table 3 in ref [83]). This marks the beginning of distinctive local developments and innovations influenced by both western and eastern Mediterranean traditions. Evidence for a Mesolithic occupation of the islands, however, remains scarce and indicates the presence of hunter-gatherer groups from around 11,000 BP. Similarly relatively few sites are dated to the Early Neolithic horizon on both islands. On Corsica, sites are concentrated in the northwest and the southern part of the island, whereas in Sardinia early Neolithic evidence clusters mainly along the west coast. The only burial securely dated to the Early Neolithic on Corsica is at L'Araguina-Sennola, where an individual in crouched position was interred in a pit, possibly with ochre applied to the skeleton. Next to the skull, a small Early Neolithic vessel containing a red substance was deposited. Additional inhumations are documented in rock shelters, such as the poorly preserved skeleton from the valley of Saint Julien. This individual was accompanied by oyster shells, fragmented ceramics, obsidian flakes, and burnt ovicaprid bones that yielded a radiocarbon date in the first half of the 7th millennium BP[84,85].

Evidence of burials in the Sardinian Early Neolithic Cardial Horizon is also limited[86]. Human remains from the cave of Rifugio have been dated to 7400–7300 BP (table 3 in ref [83]). At the rock shelter of Su Carropu, skull fragments of one male and one female found together with sea mussels, schist pendants, and a grinder are interpreted as Early Neolithic burials[86,87].

##### 2.2.2 The Middle Neolithic and earliest funeral architecture

With the emergence of the Early–Middle Neolithic Bonu Ighinu horizon, the Neolithic became firmly established, introducing the first more sophisticated funerary architectures, dating to the second and third quarters of the 5th millennium BC (6750–6250 BP, fig. 9 in ref [88]). Burials from this period are documented in pits and natural caves, as well as in the earliest hypogean necropolises, which are artificial caves used for burials[89,90] (pg. 54 in ref [88]).

The necropolis of Cuccuru S'Arriu at Cabras represents the earliest known site of this kind in Sardinia[90,91]. This cemetery comprises 19 graves from the Middle Neolithic Bonu Ighinu horizon, 13 of which are hypogea while the rest are pit burials (pg. 372 in ref [90]). Most were designed as single interments with one chamber and a single access, although two graves contained double burials. The deceased were deposited in crouched position on their left side, in some cases partially covered with ochre. Grave goods include female figurines, lithic and bone tools, obsidian geometric microliths, as well as ceramic vessels such as lugged goblets, keeled pots, and flat bowls (figs. 2 and 6 in ref [90]).

In Corsica, burials of this horizon occurred in natural caves, rock shelters, single stone cists, and cist necropoleis. To date, around 75 stone cists are recorded from 41 sites, though not all can be securely assigned to the earliest phases (pg. 43 in ref [92]). These closed subterranean or semi-subterranean structures, often covered by small tumuli, are typically located on hilltops or mountain slopes, measure 1.5–3 m in length, and are mostly oriented to the southeast. Clusters of cists are found at Vasculacciu, Tivulagju, Poggiaredda, Cardiccia-Foce Cumpra, Palavese, Ciutulaghja, Capu di Locu, and Monte Revincu. Bone preservation is generally poor, and traces of calcined remains suggest cremation. While usually regarded as single graves (pg. 92 in ref [93]), conclusive evidence remains lacking since there are no bones preserved.

At Vasculacciu, at least eight cists have been documented, some marked by standing stones and tumuli. Structures A and B are more elaborate, yielding several hundred obsidian and flint fragments, as well as discoid beads, and are dated typologically to the later 7th millennium BP[94]. Poggiaredda, also in the Figari–Porto Vecchio depression, includes a southwest-oriented cist or dolmen within a stone circle and two standing stones, associated with obsidian flakes but lacking precise dating. Tivulagju has two substantial cists, one beneath a tumulus of ca. 10 m diameter [95]. Other sites include Cardiccia-Foce-Cumpra with six cists and standing stones; Palavese with a cist under a cairn containing an arrowhead from later phases; Ciutulaghja, a plundered cairn with a cist or dolmen; and Capu di Locu, a disturbed plateau site with one cist and scattered obsidian[92].

Finally, at Monte Revincu a necropolis has been identified in close association with a settlement, comprising at least nine cists and dolmens[92,96]. This site may represent the transition from stone cists to the earliest megalithic graves on Corsica, dated to around 6250 BP.

Monte Revincu is the only stone cist necropolis with radiocarbon results. One of the radiocarbon results (Ly-8396, 5405±55 BP, 4332−4261 cal BC, 68.2% and 4351−4190 cal BC, 95.4%) is from cist D in the main sector Cima di Suarella with 34 houses and grave structures. Cist D had two layers. The layer on the basement of the grave, where the sample originated, was rich in charcoal. Some small ceramic fragments, quartz, obsidian, and a grinder fragment in the cist verify the dating of this grave to the end of the 5th millennium BC (6200–6000 BP, page 49-52 in ref [92]).

2.2.3 The earliest megaliths and the passage grave tradition

Corsica represents one of the earliest megalithic regions in Europe. At Monte Revincu, a transition from subterranean cists to accessible aboveground monuments and passage graves occurred around 4335–4264 cal BC/6285–6214 cal BP (95.4%), possibly earlier. The Dolmen Somnital (Cellucia) at Monte Revincu is the earliest known passage grave in the Mediterranean and together with the dolmen Casa di L'Urcu represent the only Corsican monuments with radiocarbon dates directly tied to their construction or first use. In total, only 18 dolmens and passage graves are known on the island, mostly clustered in the southwest. Little is known about contemporaneous non-megalithic burials such as the cave burials at Calanchi. The megalithic monuments were frequently erected in remote mountain locations, sometimes associated with settlements, as at Monte Revincu.

On Sardinia, the earliest megalithic graves are the non-accessible dolmens of the Li Muri necropolis at Arzachena in the northeast. No radiocarbon results exist for these tombs, but based on grave goods attributed to the Late Neolithic San Ciriaco horizon, and on radiocarbon dates from San Ciriaco layers at the settlement of Contraguda, their construction can be placed between ca. 6400 BP (4472–4099 cal BC, 68.2%; 4728–4070 cal BC, 95.4%) and 6100 BP (4116–4005 cal BC, 68.2%; 4218–3986 cal BC, 95.4%).

These tombs are a singular phenomenon of the period, concentrated in the Arzachena region. Their architecture suggests influences from the western Mediterranean continental coast or Corsica, and they may represent a megalithic colony on Sardinia, where hypogea, pit burials, and cave burials were otherwise the dominant funerary practices. Material culture further supports links between Li Muri and the contemporaneous or slightly earlier necropolis at Monte Revincu[1,23].

During the Final Neolithic Ozieri horizon from ~ 6000 BP onwards (4218–3433 cal BC, 95.4%; 4115–3501 cal BC, 68.2%), the first accessible megalithic structures appear on Sardinia. Ozieri pottery has been documented at the dolmens Alzoledda and Ciuledda, the passage grave Dolmen de Motorra, and at the *tomba a circoli* of Pranu Muttedu. Hypogea remained the predominant burial form, with around 3500 known examples, compared with ca. 227 megalithic tombs distributed mainly across central and northern Sardinia (chapter 6 in ref [1]). Many hypogea are highly elaborate and labor-intensive, suggesting that megalithic graves represent an alternative funerary practice.

In contrast to megalithic tombs, standing stones are far more numerous: Corsica alone has 1078 recorded examples, many set in front of cists or megalithic tombs. Some are anthropomorphic or sculpted into statue-menhirs, plausibly representing individuals, mythical figures, or supernatural beings. The traditional typological sequence from simple standing stones to anthropomorphic forms and statue-menhirs remains debated. However, the spatial organisation of the Cauria alignments, from simple standing stones to statue-menhirs, is consistent with this interpretation, as are the association of single standing stones with Neolithic cists and Bronze Age weapon depictions on statue-menhirs. Sardinia follows a similar trajectory, from simple stones, associated with the megalithic monuments of Li Muri, through anthropomorphic forms to statues linked to Ozieri contexts. From ca. 5780 BP (4004–3550 cal BC, 95.4%), a centralizing tendency is evident in northwestern Sardinia with the Monte d'Accoddi settlement and megalithic pyramidal-like monument. This site, associated with the Ozieri and subsequent cultures, may have served as a communal monument[97] or an elite political/religious centre. It is unparalleled in the Mediterranean, and linked with contemporaneous hypogea, such as the San Benedetto hypogea in the municipality of Iglesias.

***In summary,*** Neolithic groups reached Corsica and Sardinia around 7400–7300 BP, initially establishing only sparse settlements concentrated along the northern and southern coasts of Corsica and the western coast of Sardinia. Between 6800 and 6400 BP, the first hypogea, artificial caves used for burial, appeared, marking the emergence of more formalised funerary practices. The earliest megalithic monuments were constructed between 6400 and 6200 BP in small clusters or colonies and display strong architectural affinities with contemporaneous monuments in Brittany, southern France, and other parts of the western Mediterranean. Later monuments, including passage graves, are comparatively rare and reflect multiple waves of migration, interaction, and cultural exchange across the western Mediterranean. On Sardinia, the highly elaborate and labor-intensive hypogea suggest that megalithic tombs represented an alternative funerary tradition that developed through repeated cultural transmission and contacts with other megalithic regions rather than serving a distinct social group. Remarkably, standing stones are far more numerous than megalithic tombs on both islands and are frequently arranged in alignments, a monument form otherwise best known from Brittany. The construction of the megalithic pyramid of Monte d'Accodi marks the beginning of a distinctly insular trajectory of monumentality, developing at the crossroads of the western and central Mediterranean.

#### 2.3. The Neolithic sequence for southern France and farming expansion northwards: from Early to Middle Neolithic

*Stephen Shennan*

In western mediterranean Europe, Iberia and southern France, the initial arrival of farming occurs ~78/7600 BP via a process of rapid maritime colonisation by small groups with a subsistence based on sheep/goat rearing and emmer wheat[98–100], associated with Impressa pottery. The presence of obsidian indicates long-distance maritime links from southern France to Sardinia and especially to the source on the island of Palmarola, off the Italian coast between Rome and Naples.

The earliest dated Impressa sites in the northwest Mediterranean date to ~7900-7750 BP (stage 3B in Binder et al.'s [66] Mediterranean Impressed Ware chronology): the sites of Arene Candide in northwest Italy and Peiro-Signado and Pont de Roque Haute on the Languedoc coast. The site of Pendimoun starts slightly later, ~7750-7600 BP. Their ceramic assemblages are very diverse (pg. 66 in ref [66]) but they are all linked by the use of the Spiral Patchwork technique of vessel construction, which also occurs at the site of Saracena - San Michele in Calabria[101] (pg. 129 in ref [78]). The subsequent Franco-Iberian Cardial style is characterised by shell impressions in horizontal and vertical bands or panels. It is found along the northern Mediterranean coast from NW Italy across southern France to Iberia but also occurs in Tuscany, Sardinia and Corsica. Its origin probably lies in the Tyrrhenian Cardial though an alternative proposes that it originated in Iberia and then diffused back eastwards. Ibañez-Estévez et al.[102] suggest that the Cardial may represent a second wave of Neolithic expansion, mainly terrestrial, that would have started in NE Italy and then spread around the north Mediterranean region from Liguria to Catalonia and inner Iberia. As one indicator they draw attention to the distribution of the distinctive La Draga type sickles, which, together with abrasion traces on some blades with sickle gloss suggested to result from cutting stems on the ground, are found in NE Italy in the mid-6$^{th}$ millennium and then spread westwards through the coastal regions of southern France as far as Catalonia, where they are found at the site of La Draga itself, and the interior of Iberia. This led, they suggest, to the gradual substitution of the La Marmotta sickle type by La Draga ones among the pre-existing Impressa farming communities. Cardial exchange links reflect this terrestrial pattern. The obsidian coming from

Palmarola and Sardinia found at the early pioneer maritime Impressa sites from Liguria to Languedoc disappears with the Cardial.

The earliest dated Cardial phases occur at Pendimoun and Arene Candide, early in Binder et al's phase 4A, ~7450-7300 BP (pg. 75 in ref [66]). Two Cardial sites in the middle Rhone valley, Ronze and Oullins, also begin at this time. In NW Italy and Provence the Cardial continues through the two following phases, 4B and 4C (7300-7000 BP) but further west, in Languedoc, the early Epicardial is found at the sites of Taï and Camprafaud from ~7300 to ~7000 BP. In phase 4C, 7150-7000 BP, many sites in the middle Rhone valley and eastern Languedoc adopt this style (pg. 87 in ref [66]).

Settlement was initially in the coastal zone but had reached the middle Rhone valley by ~7400 BP and by ~73/7200 BP had already expanded into mountainous areas in NE Iberia[103] and the south of France[104]. In fact, van Willigen (pg. 340 and fig. 1 in ref [105]) points to evidence that there was Franco-Iberian Cardial occupation as far north as the upper Loire valley and along the Saône already by 7250 BP, with evidence of an Epicardial spread west towards the Atlantic coast at the same time. He goes on to suggest that the rapid northward Cardial expansion can be accounted for by the farmers' preference for easily cultivable soils. For the Rhone valley Berger[106] suggests that favourable soil conditions were associated with increasing settlement densities from ~7200 BP and consequent northern expansion. However, there was no stereotyped settlement formula of the kind that characterised the LBK.

Further to the north, in the upper Rhone valley in the Jura, well beyond the Mediterranean zone, the Gardon cave has elements pointing to interactions between the Cardial and the LBK-derived Neolithic. The earliest Neolithic there dates to ~7150 BP and contains lithics with southern connections and sherds reminiscent of northern Limburg pottery. This is in keeping with Limburg and La Hoguette sherds at the western Languedoc site of Gazel in an Epicardial context. The Gardon site is also important because it has evidence of a Mesolithic occupation succeeding a Neolithic one, suggesting shared territories[107]. Further north again, the Franco-Iberian Cardial seems to be the inspiration for the enigmatic La Hoguette and Limburg pottery that is widespread in the Rhine valley and across France in the later 6th millennium (fig. 11 in ref [105]). Moreover, there is evidence of contact between the Cardial and the LBK of northern France and its successor, the Blicquy/ Villeneuve-St Germain (BVSG) culture (~6850-6600 BP), for example bracelets of limestone, serpentinite and jadeite from Provence and northwest Italy found in LBK and BVSG contexts in the Paris Basin (e.g. pg. 345 and fig. 14 in ref [105]). Subsequently, the Augy-Sainte Pallaye group develops in the SW Paris Basin from ~6900 BP as a synthesis of elements from both the Mediterranean and Central Europe, with slate bracelets and trapezoid house plans paralleled in contexts but with ceramics whose closest parallel is in the Franco-Iberian Cardial (pg. 347 and figs 16-17 in ref [105]). Hamon and Manen (pg. 731 in ref [100]) suggest that these regions should be seen as 'melting pots' where novel cultural and economic constellations took shape under a variety of influences.

##### 2.3.2 Genomic evidence

A growing number of ancient genomes are now available for southern France (pg. 85-96 in ref [66])[108]. The earliest burial at Pendimoun, a female, dated to ~7600 BP has a Western Hunter Gatherer (WHG) component of ~23%. A female and a male dating to ~7300 BP have WHG components of 56% and 30% respectively. At the site of Bréguières, also in eastern Provence, three individuals dated to 7300-7000 BP also have high WHG percentages, from 23% to 38%. Further Cardial-Epicardial samples with high levels of WHG admixture come from the Baume Bourbon and Gazel caves. Such high levels of admixture are in marked contrast to LBK individuals in the Central European farming stream but also to individuals from Italy and Croatia, as well as Iberia, which is perhaps more surprising. With one exception the HG component in early Neolithic individuals from southern France is associated with Villabruna ancestry. The exception is an individual from Baume Bourbon, with GoyetQ2 Magdalenian-related ancestry, which is more frequent in Iberia, in Andalusia and Catalonia, which may point to social networks linking these regions. The early end of the estimated admixture date ranges is ~8200 BP, before the arrival of the Impressa in the NW Mediterranean but the later end coincides with the date of their arrival ~7700 BP.

Radiocarbon dating suggests that in Italy there was a 200 year gap between the end of Mesolithic occupation and the arrival of farmers. In Catalonia and Languedoc, Mesolithic and Neolithic groups immediately succeed one another. In NW Italy they may have overlapped but there is ‘no evidence of direct, permanent contact between them’ [107]. In fact, the latter conclude that what they call the ‘no man’s land’ and ‘musical chairs’ models represent the generally prevalent model of Neolithic colonisation (fig. 7 in ref [107]). All this makes the high levels of HG admixture puzzling. Unsurprisingly, in the light of the rapid coastal maritime expansion process, the evidence indicates that West Mediterranean farming groups had smaller effective population sizes than those in Central Europe, and only gradually increased [108], which could have a bearing on the high level of admixture.

The Epicardial in southern France is succeeded ~6800 BP by a variety of groups under the heading of Middle Neolithic 1 which are in turn followed by the beginning of the Southern Chasséen ~6400 BP. Genomes from these periods, as well as those from post-Chasséen groups are similar to those from other regions in France and from the Iberian peninsula, and indeed more generally to others in western Europe in the Middle Neolithic, with levels of HG ancestry between 10% and 30%, and sometimes higher. Tests showed these individuals ‘were best modelled as a two-way mixture of Villabruna WHG and Anatolian farmers', with no evidence of GoyetQ2 ancestry[108]. These high levels of HG ancestry continue into the Late Neolithic. One of the individuals from a Late Neolithic collective burial at Aven de la Boucle in the Gard region of southern France had an estimated HG component of 36% with an estimated admixture date ~7000-6200 BP; the others had values of 13-29% with estimated admixture dates ~7800-7100 BP.

##### 2.3.3 The Chasséen

As just noted, after a period when a number of different groups emerge from the Epicardial, the Chasséen cultural complex develops ~6400 BP. Its nature has been the subject of extensive and inconclusive discussion[109] but it seems to be essentially a set of ceramic forms and styles, with varying distributions, and common technological features in their manufacture. Forms include dishes with flat decorated rims, dishes with decorated bases (so-called vase-supports) which may have had a ritual significance, and carinated bowls, sometimes with single or multi-perforated horizontal lugs[110,111]. It seems to have originated as a result of the interaction between local Cardial successor styles and elements of the Vasi a Bocca Quadrata (VBQ) ceramic tradition of northern Italy, driven by interaction in exchange networks involving the distribution of Alpine jadeite axes and Bédoulian flint from southern France.

During this period the upland cave and rock shelter sites of southern France and Liguria have evidence of intensive use as sheep-folds[112]. On the basis of the seasonality of animal slaughter at different types of sites Helmer et al. [113] argue that summer transhumance was occurring at least from the end of the Cardial, associated with specialised milk production, and that different communities, and different groups within the same community, had complementary specialisations, especially pastoralism, and were linked by exchanges.

This pattern is very different from that of the lowland sites, for example in the Rhone valley, which have a much more even balance between cattle and caprines and the sites include large earthwork enclosures, though the extent of cereal agriculture is unclear. Bréhard et al. [114] have carried out faunal analyses that integrate the large lowland river terrace sites of the middle Rhone valley and the role of cattle into the picture. For the caprines (mainly sheep) they confirm the picture of herd movement and complementarity between lowland and upland sites and show that a similar complementarity between different sites is implied by the ages at death of the cattle, though in this case not all the parts of the system have been discovered. The fact that the annual movement patterns of sheep/goats and cattle were not the same suggests that there must have been specialist cowherds as well as shepherds and also leads them to the conclusion that the large lowland sites which appear at this time must have been seasonal gathering places rather than permanent settlements (pg. 186 in ref [114]), which corresponds to the marked lack of evidence for structures that might be permanent houses[115] .

The context of these developments is one of environmental stability. Geoarchaeological work has demonstrated the existence of a characteristic soil horizon found over broad areas in the Rhone valley and elsewhere, dating to ~6500-5500 BP indicating continuous grassland cover, over the period of the Chasséen culture (pg. 52 in ref [112]). Since this was also the period when charcoal evidence indicated that oak cover was at its most extensive they suggest the existence of ‘a particular type of agro-sylvo-pastoral management’, analogous to the ‘Dehesa’ system still found in certain environments in Iberia. These have a parkland appearance and involve grazed grassland with areas of cereals interspersed with managed trees, especially oak, exploited for a variety of products including leaf-fodder (pg. 56-59 in ref [112]). The vegetation beneath the trees would have been maintained as grassland both by the grazing and by regular controlled fire-setting.

The impact of the Chasséen can also be seen In Liguria where the characteristic pottery and lithics occur in the late 7th and early 6th millennium BP, marking a sharp break with the material of the preceding Square Mouthed Pottery culture[116]. At Arene Candide, 250 km from the source on Mont Ventoux, Bedoulian flint makes up over 50% of the lithic assemblage; it appears that at this time the site was a sheepfold like those examined by Helmer and colleagues[113]. Indeed, the economic picture generally is very similar to that further west, with a greater human impact on the environment than previously, including increased use of fire, and evidence for summer transhumance and woodland management, including the use of leaf fodder.

A new feature of the Chasséen is the emergence of enclosures, for example Villeneuve-Tolosane[117]. A number of these are known from southern France, though not nearly as many as in the Chasséen of northern France. At Villeneuve-Tolosane the earliest occupation may have taken place as early as 6500 BP; a ditched and palisaded enclosure was constructed after 6200 and abandoned before 5950; a new enclosure was then built, larger than the previous one, that was itself abandoned c.5800 BP. The final enclosure may have covered as much as ~28 ha. Gandelin (pg. 172-173 in [117]) sees the ditches and palisades as defensive and as indications of 'competition for conquest of territory' in a pioneer zone of farming occupation, and notes increasing evidence of violence in southern France from the MN.

Nevertheless, any warfare or inter-group competition certainly did not interfere with an enormous intensity of exchange during the southern Chasséen MN since there is evidence of extensive long-distance contacts. Most striking is the large quantity of pale Bedoulian flint from Mont Ventoux in Provence over 300 km away, which represents well over 50% of some of the assemblages at Villeneuve-Tolosane. Significant quantities of flint also came from Aquitaine to the west. Variscite beads are also found, which most probably came from the mine at Can Tintorer near Barcelona.

As noted above, after the appearance of obsidian from the islands of Sardinia and Palmarola in association with the first settlements of incoming farmers in Liguria and southern France it is not found in Cardial contexts of the second half of the 6$^{th}$ millennium BC (7500–7000 BP). It appears that maritime contacts and exchanges were not maintained after the initial settlement. Obsidian, a naturally occurring volcanic glass, then occurs in small quantities in southern France through the 5$^{th}$ millennium, including the earlier Chasséen phases, when most of it comes from Lipari, as it does in Italy during the same period[118]. In the late 5$^{th}$ millennium BC (6500-6000 BP) and the first half of the 4$^{th}$, the classic and late phases of the Chasséen, however, there is a major increase in the number of obsidian finds, now dominated by the Monte Arci source in Sardinia[118]. At this time the distribution even extends as far as Catalonia to the west, where it occurs in a number of graves with other materials obtained by long-distance exchange, including Bedoulian flint, as well as local variscite beads from the Can Tintorer/Gavá mine.

According to Lemercier[119,120], the Chasséen MN represents ‘the end of the process of neolithisation of continental western Europe and the completion of the process of establishing an agro-pastoral system, and at the same time a major territorial expansion of Neolithic societies, that reach their furthest limits. This world was now full and suggestions of social transformation were already very apparent.’ This claim is confirmed by the peak in the summed radiocarbon calibration curve at just after 6000 BP, after which the curve shows a steep decline to a trough in the middle of the 6$^{th}$ millennium. Lemercier suggests that problems may have been exacerbated by a shift to wetter and cooler climate conditions from ~5700 BP for ~400 years and also proposes that a population decrease in the mid-6$^{th}$ millennium would make sense of the scale of the changes and of the scarcity of sites in this period. Delhon et al. (pg. 62 in ref [112]) also see evidence that the rather fragile 'agro-sylvo-pastoral' system that they reconstruct for the Chasséen MN in the Rhone valley comes to an end at this time, when the wetter conditions lead to the erosion and illuviation of the black soils characteristic of this system, which had lasted for the best part of a thousand years. They also note a resulting change in settlement pattern that involves leaving locations in the lower plains. Similarly, in the Franche-Comté region in eastern France, just outside the Mediterranean zone, Pétrequin and Weller (fig. 10 in ref [121]) show that the Néolithique Moyen II, from 6300 BP, corresponding to the Chasséen shows a gradual rise in population up to c.5700 BP, followed by a very rapid crash. The result of the demise of the Chasséen is not its replacement by a new large-scale uniformity but the appearance of a number of new localised cultural groups in different parts of Mediterranean France.

While the main domain of the Chasséen is the southern half of France, a version of it also appears in the north, the so-called 'northern Chasséen'[Chasséen septentrionale], whose precise definition remains under discussion[122]. It occupies the western half of the Paris Basin with the Michelsberg in the eastern half. Its appearance in northern France by ~6300 BP indicates that its spread was extremely rapid, presumably based on the links created by the exchange of jadeite axes which had already existed for several hundred years (pg. 545 in ref [123]), but the mechanism responsible for its appearance remains unclear. It characterises the Middle Neolithic II phase, following the Cerny culture of Middle Neolithic I. In the northern Chasséen the characteristic vessel forms such as carinated bowls and the well-known decorated 'vases-supports' are frequent in the early phase but rare in subsequent phases, suggesting that the impact of the southern Chasséen was short-lived. Moreover, the technical quality of the pottery, in terms of finish, firing and tempering, is also inferior to that found in the south, while the carinated bowl forms become more loosely defined and no longer follow the rules that applied elsewhere, resulting in a much freer development of both forms and techniques[110,122]. In fact, Michelsberg vessels are technically closer to the southern Chasséen than northern Chasséen ones are, suggesting that there were two different influence routes from the south, an eastern and a western one, the latter being associated with the northern Chasséen.

However, the ceramics are not the only domain in which the northern Chasséen is distinctive. Another specific feature is the occurrence of female figurines (fig. 3 in ref [122]). There are also differences in the lithics between the northern Chasséen and other cultures in the Paris Basin, not least in its participation in the networks that circulated axes of bartonian flint from mines such as Jablines. In terms of animal economy it is also distinctive in the higher proportions of cattle and ovicaprids compared with the Michelsberg and the Noyen Group, which have greater emphasis on pig.

Overall, Colas et al. (pg. 138 in ref [122]) see a short Chasséen impulse on a strong Cerny foundation in the west and a combination of Cerny and Rössen in the east, visible not just in the presence of enclosures, which already existed in both, but also in particular features of the pottery, such as bone temper, and in the flake-based lithic industry as well as the use of tranchet arrowheads.

#### 2.4 The Neolithic transition in Northern and Northwest France

*Bettina Schulz Paulsson*

##### 2.4.1 Pioneering settlers in the Early Neolithic (7000-6700 cal BP)

The Neolithisation process in northern France was relatively late and slow compared to neighbouring regions and characterised by longer alternating phases of stabilisation and expansion. While in Belgium, particularly the Hainaut region, the oldest LBK sites are dated to around 7400–7300 BP, the earliest LBK settlements in the Paris Basin appear around 7100 BP and are clustered along major waterways, a characteristic of pioneering settling processes[124] .

The Paris Basin thus emerges as a transitional zone and a cultural melting pot, where LBK and Blicquy/Villeneuve-Saint-Germain groups coexisted for several hundred years. Understanding the relationship between these different groups has proven challenging, as they share many cultural traits, including a similar typological development of pottery, comparable compositions of lithic assemblages and house structures[125–127].

Further to the north and towards northwestern France, the Early Neolithic archaeological record is characterised primarily by the Blicquy/Villeneuve-Saint-Germain (BVSG) horizon. Rescue excavations in recent years have uncovered several Early Neolithic sites in the far north of France, confirming that these regions were not settled before 6950 cal BP while the absence of LBK groups in this area cannot be explained solely by soil conditions or geological factors[128].

Three of these northern sites have recently been radiocarbon dated, indicating a time frame consistent with the generally accepted duration of the BVSG horizon of approximately 300 years, spanning from 7000/6950 BP to 6700/6650 BP[128,129], the transition into the Middle Neolithic horizon and the Cerny and, further to the west, Castellic horizons. One possible explanation for this delayed settlement is the continued strong presence of hunter-gatherer groups, although archaeological evidence for this remains limited in the region.

Subsistence at these BVSG sites is based on emmer and the new naked wheat, with the earliest evidence found at Loison-sous-Lens. In contrast, in the Paris Basin, naked wheat is only observed in the later BVSG horizons. The introduction of this new cereal coincided with another early technological innovation: the tranchet technique for flint suggesting a shift in agricultural practices, particularly in field maintenance and hoeing techniques, required for cultivating naked wheat, which is more delicate than hardier cereal varieties[128,130] .

The situation is different in northwestern France and Brittany. Here, only a handful of early Neolithic settlements associated with the BVSG horizon have been documented. A stronger presence of hunter-gatherer groups, potentially leading to a stagnation or halt in Neolithic expansion, has been the subject of ongoing discussion for this region for several years and recently studies confirmed that Hunter-Gatherers co-existed with the earliest farmer and pioneer settlements in Northwestern France[128]. It is noteworthy that the acidic soils of the region restrict in general the preservation of organic and in particular of bone materials. Only a few sites in sandy and chalky environments at the coasts provide good preservation conditions for bone material, such as the two Late Mesolithic cemetaries on the islands of Hoedic and Teviec.

One of the individuals at Hoedic (B1, see table 1 in ref[131]) suggests the reservoir effect corrected time interval of 7050-6670 cal BP, corresponding to the presence of the earliest farmers; four more individuals from Hoedic (C2, C3, J7, J11) and one from Teviec (B2) show possible overlaps to the earliest farming populations.

##### 2.4.2 Monumental graves, stone alignments, earliest megaliths and long distance green stone trade in the Early Middle Neolithic (6700-6300 cal BP)

Around 6700 cal BP, with the onset of the Middle Neolithic, a cultural break is observable across Northern and Northwestern France. This period is marked by monumental, though pre-megalithic, grave structures in the whole area and more to the Northwest, in Brittany, also by megalithic stone alignments, particularly in the Carnac region, as well as the earliest megalithic tombs and long-distance intercultural exchange of rare raw materials and green stone such as jadeite and variscite.

In the Paris Basin we meet, contemporaneous to grave fields with pit burials, a monumental but non-megalithic grave architecture, which starts contemporaneously with, or even slightly earlier than, in the rest of the Northern region, at the beginning of the Middle Neolithic and the Cerny horizon 6700 BP. These so-called Passy-type graves emerge in the central Paris Basin and the Seine and Yonne river valleys and the plain of Caen[119,132–136]. The graves in the eponymous Passy necropolis itself are impressive, labor-intensive structures with a length of up to 280 m, built as round earthen tumuli covering single or a few burials and enhanced with long palisade systems. 19 monuments are known from the Passy necropolis covering altogether 26 burials. For 17 of these individuals, radiocarbon results are available. The earliest individual died in ~4960 cal BC/6910 BP (5029–4946 cal BC, 68.2%; 5061–4858 cal BC, 95.4%). The measurement of the last burial yields a time span ranging from 4356−4238 cal BC (95.4%; 4342−4283 cal BC, 68.2%). These results indicate that the individuals buried in the Passy monuments do not represent the population as a whole, as the estimated burial rate corresponds to, at most, a single individual per generation. This suggests that the Passy monuments served as burial places for a selected segment of society, potentially including emerging religious, social, or political elites. However, this interpretation is not unequivocal:

more than half of the individuals are children with few or no grave goods, and only four of the 26 graves contain prestige objects, including the so-called "Eiffel Tower" bone spearheads[119,136].

Altogether 13 necropolises with these types of monuments have been detected in the Seine-Yonne region from aerial photography. From the same region somewhat later results from similar monuments are available with the dates for the necropolis of Les Réaudins Balloy from 6420 BP onwards (4442–4356 cal BC, 68.2%, 4489–4341 cal BC, 95.4%).

Monuments comparable to those of the Cerny horizon also occur in the Caen Plain, including the necropolises of Rots, Mathieu, Blainville-sur-Orne and Fleury-sur-Orne. The latter comprises 32 earthen long barrows, some extending up to 300 m in length, containing the remains of 19 individuals attributed to the Middle Neolithic Cerny culture. Two phases have been identified: an earlier phase dated to 6630–6275 cal BP and a later phase dated to 5950–5540 cal BP, represented by a small number of secondary burials inserted into pre-existing monuments[137].

Two burial types can be distinguished: simple pit burials and stone-lined pit burials. The simple pits, measuring up to 3 × 1 m, contained individuals buried in crouched or supine positions, closely resembling the burial practices of the Passy groups in the Yonne–Seine Valley. Grave goods are generally sparse and include bone awls, strike-a-lights consisting of a flint blade and an iron pyrite nodule, wild boar tusks, and arrowheads. Animal offerings are also present, including sheep and goat remains; notably, one grave at Fleury-sur-Orne contained the remains of five complete sheep.

The second burial type consists of subterranean chambers constructed from limestone slabs. These were probably roofed with wooden beams before being covered with stone slabs and enclosed within the earthen monument (pg. 131 in ref [138]).

The situation is different to the Northwest, particularly in Brittany. The Middle Neolithic and the Ancient Castellic horizon here is characterised not only by pre-megalithic monumental tombs but also by the emergence of megalithic structures in Europe, standing stones arranged into alignments, and large deposits of green stone such as jadeite axes and variscite beads indicating long-distance trade (cf. section 2.11).

The first monumental, but non-megalithic, graves in Brittany were built as long or round tumuli (tertre), covering pit burials, stone cists and dry wall chambers as the first phases of the tumuli Carnacéens, large monumental tombs in the Bay of the Morbihan, which are up to 120 m long. The earliest construction phase of the Carnacéen tumulus of St. Michel, consisting of a small above-ground dry-stone chamber subsequently covered by a low earthen mound, is dated by a cremated human bone to c. 6680 BP (4780–4590 cal BC; 95.4% probability). A recently discovered and substantially smaller monumental tomb at Le Plasker exhibits a closely comparable architectural concept, comprising a small above-ground dry-stone chamber covered by a low mound. Bayesian modelling places its construction to c. 6700 BP (4720-4680 cal BC, 95.4%)[139].

This early monumental phase is associated in Brittany partly with remarkable grave inventories, with an accumulation of prestigious grave goods fabricated from exotic raw materials, such as jadeite axes and variscite beads (e.g. refs [140–149]). The green stone was traded over long distances via the waterways; thus, the jadeite from Northern Italy imported to Carnac over the Rhone Valley and reworked into the typical Breton-style jadeite axes, which were then exported to Catalonia. The variscite was even traded from as far as a mine in Southern Spain, Encinansola in Huelva [150,151].

The first megalithic grave structures in Europe are known from Brittany and possibly from the Channel Islands[1,23]. Since there are far too few reliable radiocarbon results or sequences available, it is not possible to untangle the nuances of the emergence of megaliths in Northwest France and it is thus only possible to formulate tendencies. The earliest megalithic graves in Brittany are built within the *tertre* - round and long tumuli - and are represented by ancient non-accessible dolmens such as Tumiac, Kervinio, Castellic, St. Germain, Manio 5, Mané Hui, and the first phases of Kerlescan[152] (pg. 551– 554 in ref [153], pg. 124–125 in ref [154]). For these early megaliths, no radiocarbon determinations are available. It is only possible to limit the time interval of construction, based on the typo-chronological considerations of the grave goods, to the Ancient Castellic horizon and on the basis of Ancient Castellic contexts with associated radiocarbon results ranging from 6800-6250 cal BP. Among these early megalithic graves, the Carnacéen tumulus of Tumiac stands out for its exceptionally rich assemblage, including 251 variscite beads and pendants together with fifteen axe heads of jadeite, eclogite and nephrite. The chamber exhibits megalithic elements and could be dated to ~6400 cal BP

Remarkable grave goods are similarly registered from smaller early megalithic graves or dolmens such as St. Germain, Erdeven, in the Morbihan. While the early stage of St. Germain is connected to small ceramic pieces and a transversal arrowhead, from a later grave three golden clasps/hooks are recorded beside diagnostic Ancient Castellic ceramics. Other small structures, such as Kervinio in Trinité-sur-Mer and also Castellic in Carnac, contained more modest grave goods, for example, vessels with an oval opening (pg. 420-421 in ref [153]).

Beyond Brittany, an early megalithic tomb is documented at Les Fouaillages on the Channel Islands, where the associated ceramic assemblage combines Cerny and Ancient Castellic traditions, indicating close cultural connections across the western Channel during the earliest phase of megalithic construction.

The grave type spread from Northwestern France in the second half of the 5th millennium cal BC (~6500-6000 BP) along the waterways to Catalonia, West and Central Southern France, Northern Italy, to Corsica and on Sardinia. Moreover, the Algarve, Andalusia and Galicia presumably belong to these early megalithic regions, although the available data base does not yet permit a definite classification.

While single standing stones are a characteristic feature of the megalithic landscapes of northwestern France, they are considerably rarer farther northeast, reflecting differences in local geology and the availability of suitable stone. The earliest standing stones were erected as early as the Mesolithic and are associated with both grave and domestic contexts..

Evidence from the Mesolithic cemeteries on the island of Hoedic shows that several burials were marked by small standing stones placed in front of the graves (graves J and H, see pg. 4 in ref [143]). These graves are dated to 7490-6880 cal BP (table 1 in ref[131]). Even earlier evidence for the erection of small standing stones has recently been uncovered at Le Plasker[139,155], where a Mesolithic hut dated to ~7660 BP (5700–5620 cal BC, 95.4%) was surrounded by a series of small erected monoliths, one of which is anthropomorphic.

However, standing stones arranged in alignments seem so far to be associated with the early Middle Neolithic. The alignments in the Carnac region comprise thousands of standing stones stretching over 10 km. Their function is unclear, as was the dating until quite recently. The Le Plasker site's radiocarbon sequence—comprising nearly 50 dates—has made it possible to place the Carnac stone alignments in this sector between 6670 and 6270 cal BP. In the centuries that followed, there was only limited activity at the site, with a few additional hearths. This chronology is supported by radiocarbon dates from the stone alignment belonging to the Grand Menhir at Locmariaquer[156]. The menhir, originally 21 meters long and weighing 330 tons, was transported from the Vannes region, 12 km away, and bears the engraving of a sperm whale. Radiocarbon dates from the Grand Menhir alignment likewise suggest a main period of activity between 6600 and 6300 BP.

Typical for this time horizon are engravings of boats, crooks and jadeite axes on the standing stones, with whales as the most common motif[24].

There are nearly no settlement structures documented so far.

##### 2.4.3 The Recent Middle Neolithic (6300-5800 cal BP) Societal and cultural change and innovations

The Recent Middle Neolithic marks a sharp cultural break, particularly evident in the North, Central West, and Brittany. This shift is characterised by changes in burial rites and architecture, the reuse of destroyed engraved standing stones to build new grave forms, the decline of the greenstone trade and associated deposits.

Burial practices transitioned at the beginning of the Middle Neolithic from single or double interments to collective burial rites. Architecturally, there was a shift from small, non-accessible dolmens, also referred to as megalithic cists or coffres, to passage graves. These new grave forms consisted of a megalithic chamber and a passage or, depending on the availability of suitable building stone, were constructed using dry-stone walls or organic materials, as more commonly observed in Normandy and the Paris Basin. This new architectural concept enabled graves to be reopened repeatedly for successive and collective burials, often over several generations, while also supporting the construction of increasingly large and monumental tumuli enclosing the megalithic chambers.

The introduction of passage graves may indicate a new tendency in megalithic constructions that spread rapidly along the Atlantic and Mediterranean coasts. Access to the chambers provided the possibility to perform regular rituals within the chambers or the access corridor, either for burial activities, *rites de passage* or for ancestor worship. Accordingly, the emergence of passage graves might have been a result of complex social processes within and between the megalithic societies and the construction of increasingly more elaborated and prestigious graves, an expression of communal undertakings and an important factor for group identity.

Throughout Europe, most accessible tombs are collective graves characterised by the sequential deposition of bodies and the repeated disturbance of earlier burials by subsequent interments. Earlier skeletal remains were either carefully sorted and rearranged or simply displaced to accommodate new burials, resulting in assemblages containing up to several hundred individuals. Additional practices, including commemorative rituals, sporadic reuse, exhumation, and excarnation, have also been proposed. The earliest archaeological evidence for accessible burial monuments, although not yet megalithic, comes from the Paris Basin in northern France. It is represented by passage grave A at Vierville, Butte de Luzerne, a dry-stone-built monument with a circular chamber[1]. Radiocarbon results were gained from human bones dating to ~6450 cal BP onwards (OxA-11395, 5690±45 BP; 4536–4368 cal BC, 68.2%; 4596–4356 cal BC, 95.4%). Nevertheless, stable isotope values of this bone material are unknown and the location close to the sea indicates a possible marine diet of the population buried in the grave. Accordingly, we cannot exclude a reservoir effect, as also suggested by evidence for the region with the passage grave Le Déhus on the Channel Islands.

The earliest accessible megalithic grave in Northern and Western France is located in the central west area with the dolmen of the first phase of tumulus C in the necropolis of Prissé-la-Charrière, Deux-Sèvres. The beginning of burial activities at that grave is calculated to ~6320 BP (4357–4276 cal BC, 68.2%; 4370–4263 cal BC, 95.4%). Transitional structures to the passage graves are documented for Brittany, and for the long tumulus or *tertre* Lannec er Gadouer radiocarbon results are available which pinpoint this transition to the end of the Ancient Castellic horizon and so to ~4300 cal BC/6250 cal BP (4431–4232 cal BC, 68.2%; 4503–4104 cal BC, 95.4%). The first fully developed megalithic passage graves with simple round or quadratic chambers are not documented in Brittany before the Recent Castellic horizon (4250–3880 cal BC, 95.4%)[1].

Jadeite axes are not known from the passage graves in Northwest France[153] (pg. 515–516 in [150]) and the peak of exploitation activities for jadeite in North Italy ended at ~6300 cal BP (see SI 2.11.5.2), coinciding with the proposed emergence of the passage graves, which underlines arguments for their late chronological position (pg. 173 in ref [149]). It appears that jadeite lost its cultural esteem at that time in Northwest France and the value and social system possibly changed.

Interesting is the last stage of the already mentioned tumulus St. Michel in Carnac which was successively extended over centuries and reached its final form 120 m in length not before the Recent Castellic horizon from ~6270 cal BP onwards. The last stage incorporates a passage grave suggesting the occupation of ritual space through communities performing the new passage grave religion[1].

##### 2.4.4 Genetic results and societal change

*Gurgy*

Recent genetic studies on individual sites in northern France have provided new insights into social systems associated with the different burial traditions during the 7th millennium cal BP. Genome-wide data from 94 out of 128 individuals buried at the Gurgy "Les Noisats" cemetery in the Paris Basin (France) enabled the reconstruction of kinship patterns, demographic structures, and community contact networks between 6850 and 6500 cal BP[157].

The individuals were interred in pit graves in crouched, flexed, or extended positions, sometimes covered with ochre and accompanied by only a few grave goods. It is likely the largest cemetery in the region and the strontium isotope ratios ($^{87}Sr/^{86}Sr$) for 57 individuals allowed the identification of regional mobility and contact networks.

The burial community was organised around two main pedigrees spanning seven generations, reflecting a patrilineal and patrilocal system. The absence of half-siblings and the high number of adult full siblings point to small nuclear family units and a lack of polygamous unions or serial monogamy, including levirate and sororate marriage practices (where a woman remarries her deceased husband's brother or a man his wife's sister).

The strontium isotope ratios ($^{87}Sr/^{86}Sr$) for 57 individuals suggest regional mobility and contact networks and there is evidence for female exogamy and exchange with genetically related neighboring groups.

***Fleury sur Orne***

The cemetery of Fleury-sur-Orne provides the first genomic evidence from a monumental Passy-type necropolis. It comprises 32 monuments of varying size containing the burials of 19 individuals from the Middle Neolithic Cerny culture. Two chronological phases have been identified: 6630–6275 cal BP and 5950–5540 cal BP, the latter represented by a small number of secondary burials inserted into existing monuments.

Among the 14 genetically analysed individuals, 13 were male. The results of uniparentally inherited markers and the overall low number of long runs of homozygosity suggest a patrilineal group. Two father–son pairs were identified, both buried together in the same monument. However, no biological relationships were found between individuals buried in different monuments, suggesting that each monument may represent a genetically distinct lineage. One female individual, buried with arrowheads among the males, challenges a strictly biological or gender-based interpretation. The Fleury-sur-Orne cemetery appears to reflect a social or political elite within the Cerny culture, rather than representing the broader population.

If this is the case, the genetic evidence suggests a non-hereditary elite or chiefdom system where each generation built a monument for their leaders, who were chosen through ritual or social selection among the members of the societies.

***Port Blanc***

The two passage graves at Port Blanc, located in the sandy dunes on the west coast of the Quiberon Peninsula, are among the very few megalithic tombs in northwestern France that have preserved human skeletal remains suitable for ancient DNA analysis, owing to the exceptional preservation provided by the calcareous dune sand[158]. The monuments can be reconstructed from the excavation report by F. Gaillard[159] (pg. 123 in ref [160]), which describes two burial layers separated by stone slabs. The upper layer contained at least 12 skulls and disarticulated human remains, whereas the lower layer contained 10 skulls, including two articulated individuals. Additional articulated skeletons were recorded in the eastern tomb, which comprised a quadrangular chamber measuring 2 × 2.5 m (pg. 123 in ref [160]).

Genomic data were obtained from four individuals dated to 5950–5500 cal BP[158]. These individuals show admixture between French and Western Cardial ancestry, together with a high proportion of Western Hunter-Gatherer ancestry, suggesting admixture between French Cardial and local hunter-gatherer groups before the arrival of Southern Cardial populations in Brittany prior to 6300 cal BP.

*To summarise,* archaeological and genetic evidence suggests that a French Cardial group emerged in the southern Paris Basin around 7100 cal BP as part of the Cardial expansion into northern France. After 6700 cal BP, during the Early Middle Neolithic, this population was associated with the Cerny and Castellic horizons, characterised by the appearance of monumental graves, the circulation of prestigious greenstone artefacts, long-distance exchange networks, stone alignments, and some of the earliest megalithic monuments in Europe. The Carnac region appears to have been a major centre within this emerging megalithic world.

Around 6300 cal BP, a marked cultural transformation took place. This phase is associated with the arrival of Western Cardial influences and the emergence of passage graves, which combined collective burial traditions with local megalithic practices. In Brittany, these communities reused fragments of earlier engraved stelae in the construction of new funerary monuments, while jadeite deposition and possibly also the construction of large stone alignments came to an end. Occupying established ritual landscapes such as Saint-Michel, they introduced a new funerary and ritual tradition that spread rapidly along the Atlantic façade and beyond during the following centuries

#### 2.5 The Iberian peninsula and the Neolithic transition 8000-6000 BP

*Bettina Schulz Paulsson*

##### 2.5.1 The earliest Cardial movements along the Mediterranean coast and inland to the Northeast

The Neolithic transition followed a different course along the Mediterranean coast of Iberia than in the Atlantic regions, both in its timing and in the cultural processes involved. Debate has focused on the arrival of farming communities in coastal areas and their interactions with indigenous hunter-gatherers[98,161–163]. Improved sampling and more critical assessments of older radiocarbon dates have recently provided a more reliable chronology for this process (e.g. refs [98,163]). The earliest well-dated Neolithic evidence comes from the eastern Mediterranean coast, particularly the Valencia–Alicante region. From there, farming communities expanded southwards into Andalusia and Portugal and later into northern Iberia.

The earliest Neolithic pottery in Iberia is generally classified as *Impressa*. It is mainly found in the Alicante region at sites such as Mas d'Is and El Barranquet. Evidence for this phase remains sparse, especially when compared with southern France, where *Impressa* pottery is well documented[101,164,165]. *Impressa* was followed by Cardial pottery, which spread along the Mediterranean coast and later reached the Atlantic façade (e.g. refs [163,165,166]). Cardial farming communities initially expanded along the Mediterranean coastline. Their rapid westward movement is widely explained by the maritime pioneer colonisation model first proposed by Zilhão[98]. According to this model, small groups of farmers travelled by sea and established new settlements in favourable coastal areas. This expansion began shortly after the first

Neolithic settlement of southern France and reached eastern Iberia during the late sixth millennium cal BC (~7500-7000 BP)[98,167]. Early evidence comes from phase A of Guixeres de Vilobí in Catalonia, where domesticated sheep have been radiocarbon dated to 5643–5481 cal BC/7593–7431 cal BP at 95.4% probability (table 5 in ref [168]).

Seafaring allowed these communities to settle the Mediterranean coast rapidly while maintaining long-distance contacts and exchanging objects and ideas between regions. Cardial pottery appears from the earliest horizons. It usually has smoothed or polished surfaces decorated with impressions made using shells, including *Cerastoderma edule* and *C. glaucum*, or comb-like tools. Common vessel forms include bowls, cylindrical jars and globular bottles.

The incoming communities also introduced domesticated animals and cultivated plants. Their livestock consisted mainly of caprines, accompanied by smaller numbers of cattle and pigs. Wild animals, birds and fish are less common in the faunal assemblages. Crops included cereals, pulses and oil plants, although their frequency varied between regions and their archaeological visibility is affected by poor preservation. Barley (*Hordeum vulgare*) and wheat (*Triticum* spp.) are frequently identified, despite the often-fragmentary condition of the remains[103,169]. Pulses are more common in Andalusia and Portugal than in central and northern Iberia, whereas free-threshing wheats predominate in eastern and southern Iberia[167,170].

The lithic industry included regionally sourced flint and geometric microliths, macrolithic tools consist of adzes, grinding stones, and ochre-stained pebbles. The whole Mediterranean shared common harvesting technologies, which spread rapidly from Greece to Portugal [167] evidenced by 'glossy tools' across the Mediterranean. In Iberia, complete, parallel-hafted blades appear after 7300 BP at both coastal (e.g., La Draga, Sant Pau del Camp) and inland sites (e.g., Cueva Chaves, La Vaquera). At Los Cascajos, a transition from diagonal to parallel hafting is documented, while L-shaped sickles at La Draga and Costamar reflect local innovation. At sites such as Cova del Frare (Catalonia), domestic faunal assemblages are dominated by ovine and caprine remains, with additional evidence for cattle, dated to between 7200 and 6850 BP[166,171].

Epicardial pottery, although broadly similar in form to Cardial ware, belongs to a later stage of the Early Neolithic[164,166,171,172]. This horizon is associated with the completion of the Neolithic expansion across the Iberian Peninsula, including the interior highlands, from around 7200 BP onwards.

There is, however, some evidence for the early presence of Cardial pottery in the interior and northern sectors of the peninsula. Nevertheless, the clearest evidence for the Neolithic occupation of these regions derives from the Epicardial horizon. This is particularly apparent in the upper Ebro valley, on both banks of the river and in several adjacent valleys to the north, as demonstrated by the consistent radiocarbon chronologies from sites such as Atxoste, Los Husos and Los Cascajos. These sites have yielded Epicardial pottery together with faunal assemblages dominated by domestic animals. This northern evidence may reflect a second, inland expansion of Neolithic communities moving northwards from southern regions. At approximately the same time, Neolithic settlements also appeared in the interior of the peninsula, particularly on the Northern Meseta, at sites such as La Lámpara and La Revilla del Campo[173].

Early Neolithic funerary practices also included collective burial sites in caves and a remarkable example is the Cueva de los Murciélagos in Albuñol (Granada), where early reports describe numerous human interments. According to 19th-century accounts and subsequent evaluations (pg. 125 in ref [174], pg. 29 and 50 in ref [175]), the partially mummified remains of over sixty individuals were recovered from the cave. Notably, twelve individuals were reportedly arranged in a semicircle around a woman dressed in leather clothing and adorned with a necklace made of esparto grass, seashells, and a carved boar tusk. Other chambers contained further individuals with leather garments, esparto hats and sandals, and woven baskets. Radiocarbon dates obtained from esparto artefacts range from 7200 to 6050 BP (pg. 221 in ref [1]).

##### 2.5.2 The first Cardial groups in Western Iberia

In western Iberia, comprising present-day Portugal and Galicia, the Neolithic followed the broader Iberian pattern: farming communities arrived along the southern coast and subsequently expanded inland and northwards. This process involved both interaction with local hunter-gatherers, particularly around the Muge and Sado shell middens and coastal Alentejo, and the colonisation of sparsely populated or abandoned landscapes north of the Mondego River[170,176,177].

Owing to regional preservation conditions, Early Neolithic human remains and Cardial-associated grave goods are concentrated in the limestone karst of Estremadura in central Portugal. At sites such as Galeria da Cisterna and Gruta do Caldeirão, the dead were deposited on cave floors or beside cave walls rather than in excavated graves[177]. Recent rescue excavations in Lisbon, however, uncovered individual burial pits containing bodies in flexed positions, documenting a previously unknown funerary practice in Portugal[178].

Radiocarbon dates place these funerary contexts between c. 7500 and 7000 BP[177,178]. At Galeria da Cisterna and Gruta do Caldeirão, they span an earlier Cardial phase with richly decorated shell- and comb-impressed pottery and a later phase with typical Late Cardial ceramics. Ancient DNA identifies the individuals as members of the first farming populations in western Iberia, genetically related to contemporary Early Neolithic groups across the western Mediterranean and consistent with the broader Cardial expansion[179].

##### 2.5.3 Interaction between Cardial and hunter-gatherer groups

A long standing debate concerns the relationship between the incoming Cardial groups and the local Hunther-Gatherer societies and the dynamics of their co-existence[177,180].

Bayesian modelling of early Neolithic coastal movements supports a migration-based model, suggesting rapid assimilation or replacement of indigenous hunter-gatherer groups in northeastern Iberia, while indicating a longer period of coexistence in the Ebro Valley[180]. In the case of Mendandia, located to the northeast in Navarra at the foothills of the Pyrenees, Arias[162] proposes that Mesolithic communities either adapted pottery technology or had direct exchanges with Neolithic groups during the sixth millennium cal BC (~8000-7000 BP).

A markedly different scenario has emerged also for the Western Iberian Peninsula, particularly in central Portugal. Recent radiocarbon analyses, including marine reservoir corrections, suggest that Mesolithic shell midden sites in the Tagus estuary (e.g., Muge) and farther south in the Sado basin were occupied by hunter-gatherer groups who persisted for several centuries after the arrival of farming communities [176]. This prolonged coexistence may have spanned the entire duration of the region's Classic Cardial phase, approximately 7500–7250 BP, or up to 250 years[177].

##### 2.5.4 Neolithisation processes in the Northern regions and the Atlantic Iberian coast

Galicia, on the northern Iberian coast, was only marginally affected by the initial Cardial expansion. Why farming communities apparently halted north of the Mondego remains uncertain, but environmental and maritime constraints may have contributed. Early farming practices developed under Mediterranean conditions required adaptation to the mild, wet oceanic climate of the Euro-Siberian zone, which lacks a pronounced summer drought[181]. Although the Cardial expansion was largely maritime, Atlantic navigation presented different challenges from the Mediterranean. Strong swell, surf and currents, particularly along the Galician coast, together with prevailing northerly and north-westerly winds such as the *Nortada*, may have hindered northward movement[182].

The evidence for the Mesolithic–Neolithic transition in north-western Iberia remains sparse and difficult to interpret, partly because acidic soils hinder the preservation of organic remains[162]. Mesolithic occupation along the northern coast is represented by relatively few radiocarbon dates between c. 10,300 and 7000 BP, with the most reliable evidence clustering around 7500 BP (table 1 in ref [183]). Paradero de Reiro remains the only clearly identified Mesolithic site on the Galician coast, while poorly documented inland sites, including Serra do Xistral and O Bocelo, and coastal lithic scatters may also belong to this period[162].

Neolithic settlement evidence is similarly limited, despite the thousands of megalithic tombs recorded across Galicia. A shell-impressed Cardial-style vessel from Cova Eirós in eastern Galicia, tentatively attributed to the end of the sixth millennium cal BC (7250-7000 BP), indicates earlier contacts or small-scale Neolithic penetration than previously recognised[184]. However, this remains an isolated find rather than evidence for widespread settlement. More secure evidence combining pottery with domesticated plants or animals appears north of the Douro around 6750 cal BP, notably at Buraco da Pala and Prazo[162]. The earliest well-dated pottery-bearing sites in Galicia, including Porto dos Valos and A Gándara, belong to the second half of the fifth millennium cal BC (~6500-6000 BP) and produced mainly undecorated ceramics[185].

Impressed pottery has also been reported from the southern Galician coastal sites of A Cunchosa, O Regueiriño and Lavapés. These assemblages have been interpreted as possible evidence of maritime contacts with central Portugal and the French Atlantic coast[162,186]. The material from O Regueiriño has more specifically been compared with the Blicquy/Villeneuve-Saint-Germain tradition of northern France, potentially reflecting settlement or exchange during the second half of the fifth millennium cal BC (~6500-6000 BP).

##### 2.5.5 Intercultural maritime exchange and the emergence of megaliths on the Iberian Peninsula

In addition to the material from O Regueiriño, several finds indicate maritime contacts between Galicia and northern France, particularly the Morbihan region. Galicia was one of the few areas beyond northwestern France where megalithic monuments appeared at an early stage along the Atlantic façade.

One of the clearest indications of contact is the jadeite axe from Vilapedra. Its Alpine origin and long distance circulation have recently been confirmed[187]. The axe was made from jadeite originating in the northern Alps and has the distinctive Carnacean or Tumiac form associated with the Morbihan region of Brittany. No comparable intermediate finds are known from the northern Iberian coast, leading the authors to propose a direct maritime connection between Brittany and Galicia. Carnacean style axes are also known from Catalonia, indicating the extent of the networks maintained by the Carnac and Morbihan communities between approximately 6700 and 6200 BP. Jadeite was imported into Morbihan, worked into axes of the Tumiac type, and subsequently circulated to Galicia and Catalonia, either through exchange networks or as prestigious objects carried by individuals. The Vilapedra axe may therefore support the existence of an early phase of megalithic activity in Galicia.

Additional material evidence supports the notion of sustained interactions between Brittany and Galicia. These include the concentration of perforated axes of the so-called *Cangas type* in Galicia[29], which may reflect localised transformations of imported or shared axe traditions. Likewise, Castellic-style pottery, diagnostic of Brittany's passage grave culture, has been found in the chamber of the Dombate tomb in Galicia[187].

The large quantities of variscite beads found in the early monumental graves of the Morbihan provide further evidence of long-distance exchange. Some of the variscite originated in the Encinasola mining district in the Huelva Pyrite Belt of southern Spain[151]. Galicia may therefore have served as a transit region within maritime networks connecting southern Iberia with Brittany, although the precise routes remain uncertain[23,29].

One of the most significant indicators of shared ideology across this maritime axis is the appearance of whale depictions on reused standing stones in megalithic tombs, an iconographic motif found exclusively in Brittany and Galicia [29,188].

Monument construction starts in the second half of the 5th millennium cal BC (6300-6000 BP), and based on architectural features, Galicia has been identified as one of the earliest megalithic regions in Europe[1,23]. With at least 4,000 recorded monuments, it also represents one of also the highest regional densities of megaliths in Europe.

The northern part of the Western Iberian Peninsula, encompassing Beira, Northern Portugal and Galicia is a region with poor bone preservation. Therefore, the available radiocarbon stock from these regions is nearly exclusively limited to results gained from charcoal samples, whereas many of them represent terminus post quem values. As in Beira and Northern Portugal megaliths emerge in these regions within tumuli[189–194] from the final Neolithic and ~6200–6000 cal BP onwards or even slightly earlier the available data do not yet permit a more detailed scenario. As in Brittany, the first megalithic stage is here represented by non-accessible or accessible dolmens covered by round small earthen tumuli. These tumuli likewise cover central graves, wooden constructions, pits, stone cists, and para-megalithic structures.

The first dolmens were built with a rectangular or mostly polygonal ground plan covered by a more modest to average size earthen tumuli compared to later monuments with passage graves and exclusively stone structures (pg. 508–509 in ref [195]). The available radiocarbon dates from the chambers are gained from charcoal and represent terminus post quem values for the construction of the graves. The samples stem from pre-megalithic horizons in the 6th and the first half of the fifth millennium cal BC (7000-6500 BP, pg. 251 in ref [1]).

Cotogrande 1 is a less classical structure with a chamber consisting of five orthostats in a half round circle and one big stone slab leaning against those and therefore building a cap and a side stone[196]. This chamber, which is documented as an individual grave, was finally covered by a round tumulus. Based on the context, one of the radiocarbon results originates from the central structure with possible usage data (GrN-18420, 4940±80 BP, 3888–3644 cal BC, 68.2%; 3951–3538 cal BC, 95.4%). A further possible usage or construction date is available from Chan de Cruz 1 (CSIC-642, 5210±50, 4144–3961 cal BC, 68.2%; 4230–3947 cal BC, 95.4%).

Beside Galicia there are other Iberian and Central Mediterranean regions with small occurrences of early megalithic graves. The radiocarbon dates suggest that the first megalithic graves in Europe were closed small structures or dolmens built above ground with stone slabs and covered by a round or long mound of earth or stone. These graves emerge in the second half of the fifth millennium calibrated years BC within a time interval of c. 6700-6000 BP (4794 cal BC to 3986 cal BC, 95.4%; 4770 cal BC to 4005 cal BC, 68.2%), which can be reduced most probably to 200 y to 300 y, in northwest France, the Channel Islands, Catalonia, southwestern France, Corsica, and Sardinia and taking the architecture and the associated cultural material into consideration, megalithic graves from Andalusia, the Algarve, and northern Italy presumably belong to this first stage

In Catalonia, in the Tavertet region, early megalithic graves emerged during the same time interval, even contemporaneous with the graves in Brittany. A re-evaluation of the available radiocarbon results yielded a dating of the construction of these graves not before 6700 BP to 6100 BP (4722 cal BC to 4068 cal BC (95.4%; 4581 cal BC to 4267 cal BC, 68.2%). A part of these data exhibit an inbuilt age.

From the southwest Iberian Peninsula in Andalusia, the Algarve, and the Alentejo, we find more of these possible early megaliths. While Northwestern France is discussed as the region of origin for the megalithic phenomenon the remaining regions with an early megalithic proliferation in the fifth millennium cal BC (~7000-6000 BP) on the Iberian Penninsula megaliths are found occurring in small clusters. These are exceptional grave forms for this period in their respective regions, at a time when subterranean cists, pit burials and hypogea (dug-out subterranean burial chambers) were still the most common burial rites[1,23].

A new expansion began during the first half of the fourth millennium cal BC (6000-55000 BP), when thousands of passage graves were constructed along the Atlantic coasts of Iberia, France, Ireland and Britain. Their distribution reflects close maritime connections and the rapid transmission of the passage grave tradition along coastal routes. These monuments marked a major transformation in burial practices, accompanied by broader social and economic changes. Passage graves first appeared in central and north-western France around 6300 BP and spread rapidly along the Atlantic façade shortly after 6000 BP to Western Iberia and the British Isles. From the West Iberian Peninsula, there is best evidence for accessible structures and passage graves. For the Portuguese Estremadura, the outset of the accessible structures is calculated to 3838−3384 cal BC/5788–5334 cal BP (95.4%; 3658−3426cal BC, 68.2%), for the Alentejo to 3746−3516 cal BC/5696–5466 cal BP (95.4%; 3671−3566 cal BC, 68.2%), for the northern regions somewhat earlier, and for Beira to 3865−3784 cal BC/5815–5734 cal BP (95.4%; 3833−3797 cal BC, 68.2%)[1,197] .

For the western part of the working area in Andalusia, there is evidence for a few passage graves within the initial phase of the Almeria horizon, dating within the time interval of ~6200−5600 BP.

In Catalonia, small passage graves with subcircular chambers are not documented in the Alt Empordà before the first centuries of the fourth millennium cal BC (6000–5700 cal BP), and may be even later. A partial hiatus of several centuries can be observed between the earliest megalithic phase, characterised by small clusters of closed dolmens, and the emergence of accessible structures and passage graves.

It is also noteworthy that the predominant contemporaneous burial practices on the Iberian Peninsula during the emergence of megaliths involved natural caves, rock shelters, hypogea or rock-cut tombs, and pit burials. These burial forms originated within pre-megalithic traditions and continued alongside the construction of megalithic monuments. Their parallel existence suggests that the megalithic tradition was adopted only by a subset of communities and may support the interpretation that it was introduced by incoming groups who became integrated into local populations.

*To summarise*, the Neolithic arrived in northeastern Iberia (Catalonia) through maritime pioneer colonisation around 7630–7550 BP, following earlier Neolithic developments in southern France associated with the Cardial and later Epicardial traditions. Groups of early settlers also followed the Rhône and Ebro river valleys into northeastern Iberia. These early farming communities buried their dead primarily in collective cave burials and pit graves. While archaeological and genetic evidence suggests a rapid assimilation or replacement of hunter-gatherer populations in northeastern Iberia, prolonged coexistence between foragers and farmers is documented in central Portugal, the Tagus and Sado estuaries, and parts of the Upper Ebro and Navarra regions. The Cardial settlers reached the western Iberian Peninsula c 734O BP. Northern Iberia, including Galicia and the Cantabrian coast, was not part of the initial Cardial expansion. The spread of the Neolithic into these regions appears to have been delayed by several centuries, possibly due to environmental constraints and the challenges of Atlantic navigation, including the Nortada wind system.

Although Galicia exhibits a comparatively sparse Early Neolithic record, the region provides evidence for maritime contacts with northwestern France and an early proliferation of megalithic monuments. Outside Brittany, it represents one of the earliest Atlantic regions with megalithic architecture, with small non-accessible dolmens appearing by approximately 6200–6000 BP. Architectural similarities and radiocarbon chronologies suggest an early megalithic horizon shared across northwestern France, Catalonia, Galicia, and parts of southern Iberia, characterised by small non-accessible dolmens and regional concentrations of monuments between approximately 6500 and 6100 BP. A second major expansion of megalithic traditions followed after c. 6000 BP, marked by the widespread adoption of passage graves and the rapid diffusion of related architectural and ideological traditions along both the Atlantic and Mediterranean coasts.

During the first half of the fourth millennium cal BC (6000-5500 BP), a second phase of expansion transformed this initial pattern with the rapid spread of accessible dolmens and passage graves along the Atlantic façade, from Portugal to Brittany and the British Isles. At the same time, older funerary traditions—including cave burials, pit burials and, subsequently, rock-cut hypogea—continued to coexist with megalithic monuments throughout much of Iberia. Rather than replacing earlier mortuary customs, here megalithic architecture became integrated into already diverse funerary practices, reflecting the emergence of new forms of monumentality while preserving substantial regional variability.

#### 2.6 The farming colonisation of Britain and Ireland

*Stephen Shennan*

Although farming groups in the LBK tradition had reached the English Channel coast by 7000 BP, there is no evidence of contact until the later $5^{th}$ millennium and even much of this is open to question[198,199]. The earliest definite evidence of farming communities in Britain occurs at ~6100 BP at a time when there is evidence of much more extensive Neolithic settlement than in the LBK phase in northern France and Belgium, associated with the Chasséen and Michelsberg cultures of the Middle Neolithic 2 period, beginning ~63/6200 BP (see section 2.3.3 and 2.7). As is now well known (summarised in Brace and Booth[200] and Cassidy[201]), the introduction of farming was associated with an almost complete population turnover between ~6000 and ~5500 BP, in which the 'Western Hunter Gatherer' (WHG) ancestry of preceding Mesolithic populations was almost entirely replaced by that of populations from adjacent areas of the continent, whose own ancestry was dominated by that of descendants of Anatolian farmers (Early European Farmers, EEF) together with a lesser element of local European hunter gatherer ancestry with whose bearers they had reproduced as they spread across Europe.

However, there remain differing views of how farming and farmers spread into Britain and Ireland. Thomas[202] maintains that the ancient DNA evidence does not exclude a possible role for local Mesolithic groups in the initial uptake of farming. Sheridan (e.g. ref [203]) suggests that there were three different routes and movements, one starting from Brittany in the local MN II 6300-6000 BP and going north via the Irish Sea; the second, referred to as the 'Cross-Channel West' route, dating to the first quarter of the 6$^{th}$ millennium BP, and linking Normandy with south and southwest England; and finally, her 'Cross-Channel East' route, represented by the so-called Carinated Bowl Neolithic. This has much more extensive evidence, not just the specific pottery with its Michelsberg and Chasséen antecedents but also settlement sites with domestic plants and animals; evidence of its arrival in large parts of Britain and Ireland dates to the period 6100-5800 BP. Whittle[198] takes the view that only the 'Cross-Channel East' route is strongly supported, that farmer colonisation of Ireland was largely via Britain and that similarities in material culture and monuments indicate continued close interaction between the two in the early Neolithic. This was in contrast with the apparent loss of contact, both genetic and cultural, with continental Europe.

The speed of the spread is worth emphasising[198]. The analysis by Bocquet-Appel et al.[204] of the speed of expansion of the different regional cultures associated with the spread of farming shows that the rate of expansion of the Carinated Bowl tradition is only matched by the seaborne spread of the western Mediterranean Impressa, thus supporting the idea of a coastal expansion; the average rate of spread is three times as fast as the LBK. By ~5800 BP, farming and pottery, of the Carinated Bowl Tradition, had reached Ireland and northern Scotland. Clearly this implies that the same sort of leapfrog or long-distance movement pattern that is apparent in the West Mediterranean was operating in the case of the colonisation of Britain and Ireland as well, with rapid movements into new territories by fissioning groups long before there was any question of local carrying capacity limits being reached.

In the last few years virtually all the argument about origins and processes has focussed on the increasing quantities of genomic evidence available and its analysis by increasingly sensitive methods. It has been shown that individuals from Britain and Ireland have greater affinities with individuals from southern France descended from the Mediterranean farmer expansion than with descendants of the Central European LBK stream[200,205]. Ariano et al. [206], using the more sensitive method of analysing IBD-sharing found greater sharing with west Mediterranean and northern French groups, while Cassidy[201] specifically proposed the coastal Chasséen-Michelsberg groups of northern France, from Normandy in the west to Nord-Pas de Calais in the east as the most likely origin for British and Irish Neolithic populations on the basis of current evidence, coinciding with the archaeological evidence for the 'Cross-Channel East' route and the Carinated Bowl tradition.

A key question in discussions of a possible role for indigenous Mesolithic populations in the expansion of farming in Britain and Ireland has been whether higher proportions of WHG ancestry were present than in the continental populations from which the Neolithic immigrants came. Another has been whether there was regional genomic variation within Britain and between Britain and Ireland that could point to immigration from multiple sources in continental Europe, connecting to the debates based on the material culture; even in northwest continental Europe there is considerable variation from place to place. The two questions are related because higher proportions of WHG ancestry in some areas could be accounted for by the fact that the immigrants came from different sources rather than because they were reproducing with local hunter-gatherers.

The fact that estimated dates for the admixture between EEF and WHG groups in Britain and Ireland go back to the late $6^{th}$ millennium, long before the arrival of farming in Britain, combined with the evidence that some continental populations had high proportions of WHG ancestry, would argue against the local admixture claims but does not definitively exclude it[200] and there is an example of recent admixture from a megalithic tomb in Ireland[201].

As Brace and Booth (pg. 132 in ref [200]) point out, there has in any case been a tendency for the ‘population replacement’ argument to be overdrawn. If, as seems likely, the Mesolithic population of Britain was small then opportunities for admixture would be few, while immigration of small groups could have taken place over a couple of hundred years rather than a single mass migration. Even if reproductive interaction between the two was not actually avoided, a combination of higher farmer fertility and continuing immigration would have led to the minimal Mesolithic representation in subsequent generations that is apparent.

Genetically-based estimates of Neolithic population size in Britain and Ireland vary considerably (compare pg. 137 in ref [200] with pg. 157-8 in ref [201]) but they point to rapid population increase. The fact that ‘even the earliest Neolithic individuals from Ireland appear outbred, implying large community sizes’ (pg. 158 in ref [201]) indicates that the number of migrants was considerable.

These claims are supported by the demographic patterns in Britain and Ireland inferred from summed radiocarbon probabilities, which showed a very marked increase in population size from ~6000 to 5600 BP, followed by an even more rapid decrease in the period to ~5300 BP[207–209]. For Britain, Woodbridge et al. [210] using pollen analysis data showed that between 6000 and 5500 BP the proportion of woodland decreased and semi-open arboreal landscapes increased in correspondence with the population boom as primary forest was cleared. The reverse occurred between ~5500 and 5300 BP as the population decreased and woodland increased again. In Ireland too the boom and bust pattern seen in the radiocarbon demographic proxy has been confirmed by other evidence[211].

##### 2.6.1 Early Neolithic subsistence patterns

The evidence for EN subsistence patterns and their associated practices in Britain and Ireland fits entirely with the picture of initial large-scale immigration and demographic expansion, summarised in Rowley-Conwy et al [212]. With the exception of Ferriter’s Cove in Ireland[213], there are no bone assemblages with domestic animals prior to the beginning of the Neolithic at ~6000 BP and the proportion of domestic animals in the British EN (3900-3300 BC / 5850-5250 BP) is over 90%[214]. It is notable that the same high proportion, well over 90%, is seen in the faunal assemblages of the Paris Basin in the period 6400-5800 BP. Moreover, Tresset (e.g. fig. 3.2 in ref [215]) has shown that the relative proportions of the different domestic animals at EN sites in southern England are matched by those at sites in Normandy and the Paris Basin, while analyses of the sizes of domestic cattle show that these were also very similar.

This picture of an introduced set of subsistence practices has been confirmed in recent years by increasing numbers of isotope studies. Analyses of stable carbon and nitrogen isotopes in human skeletons showed that there was a very marked shift in the diets of communities at coastal sites from a heavy reliance on marine protein sources at Mesolithic sites to dependence on largely terrestrial resources at Neolithic sites, a pattern that occurs throughout Britain albeit with some minor variations (summary in ref[214]). These studies have been increasingly confirmed by ceramic residue analyses, which have also shown the fundamental importance of dairying and milk processing in EN economies and societies that we have seen elsewhere[216].

The earlier Neolithic crop systems in Britain and Ireland seem to have been very similar to one another[217,218]. The range was narrow, comprising emmer and to a lesser extent barley as the main crops, with some evidence of free-threshing wheat[219] and no evidence of pulses. Nevertheless, the farming system as a whole was not unchanged since the LBK. Even if intensive working of small fields continued, it was accompanied by a new forest-based dimension, perhaps associated with the creation of forest pasture for cattle, that led to the creation of extensive areas of secondary forest.

Regardless of the nature of the farming system, what is striking is the fact that, according to the data presented by Stevens and Fuller[220], after increasing rapidly to a peak ~57-5600 BP the numbers of directly dated wild plant foods and cereals in Britain both declined quite rapidly, with cereals showing an initial sharp fall from ~5600 BP and wild plant foods from

~5400 BP. An almost identical pattern in the chronological distribution of directly dated cereal and hazelnut remains has now been found in Ireland[218], with a rapid rise to a peak from ~5750 BP and a virtual disappearance of both from ~5300 BP.

Subsequent work comparing the population trajectories for Britain and Ireland as a whole with the chronological distribution of directly-dated plant macroremains using a massively increased dataset[209] has confirmed these patterns and the correlation between them. Thus, it is clear that the agricultural system brought to Britain and Ireland by the first farmers was initially successful, but it could not be sustained, hence the boom and bust, whether because of climatic deterioration, soil degradation or a combination of the two.

##### 2.6.2 Social and cultural change

In addition to the pottery, the appearance of the first farmers with their domestic crops and animals in Britain and Ireland saw the introduction of many other novel phenomena. These include long barrow burial mounds, megalithic tombs, rectangular post-built houses, flint mines and causewayed enclosures, which all have predecessors in continental north-west Europe.

Mines exploiting the flint in the southern English chalk in Sussex, such as Harrow Hill, are among the earliest Neolithic features, with dates going back to 6000 BP[221], contemporary with similar developments going on across the Channel, for example at the site of Spiennes. The Medway megalithic tombs in Kent may be nearly contemporary with the initial arrival[222]. The first earthen long barrows had appeared by ~5800 BP. It is clear though that the great majority of the various types of sites and monuments in both Britain and Ireland were constructed and mainly used between ~5750 and ~5500 BP. At the same time the earliest circular megalithic passage tombs in Ireland were constructed at the site of Carrowmore[223], while the earliest megalithic so-called court tombs in their trapezoid mounds begin only slightly later[224]. In Britain the construction and use of Cotswold-Severn chambered tombs, such as Ascot-under-Wychwood[225] and West Kennet[226], also took place at this time and the large rectangular timber buildings found in Scotland, such as Balbridie[227], were also built, as were the causewayed enclosures of southern Britain[228], which have antecedents in the source area for British and Irish immigration in the Chasséen-Michelsberg in northern France. The peak of construction of causewayed enclosures in southern England occurs during the period of the population peak, beginning ~5700 BP. Whittle et al (fig. 14.20 in ref [228]) show that the majority of the ditch construction for the enclosures they examine in detail occurs in the 57th century BP, with a subsidiary peak in the 56th century. In addition, there is evidence of extensive exchange. Beginning ~5800 BP[221] the quarries in Langdale in northern England were producing thousands of axes that were distributed over large areas of England and Scotland, as well as across the sea to Ireland.

By ~5600 BP the currency of Carinated Bowls as well as of the Scottish ‘halls’ had come to an end; from 5500 to 5400 BP we see the end of the Sussex flint mines, of digging new ditch circuits at causewayed enclosures, of stone axe exchange networks, and then ground-stone axes themselves. By ~5300 BP the southern British enclosures are no longer used, though long barrows and cairns, linear monuments and the miscellaneous category of diverse and small monuments continue later[228]. In Ireland the house phase is finished by ~5600 BP[229], while pit complexes seem to decline markedly in frequency shortly after ~5500 BP during MN I[230]. The construction of court tombs also appears to cease although their use continued. In contrast, however, the building of passage graves, which had begun during the EN II settlement boom not only continued but apparently peaked during the MN II phase, ~5400-5100 BP with the construction of the major tombs such as Knowth and New Grange in the

Boyne valley (see ref [231] for overall summary of developments). The rise and decline of these phenomena is closely associated with the demographic boom and bust described above .

#### 2.7 The Michelsberg Culture

*Stephen Shennan*

As noted in section 2.4, from ~64/6300 BP changes in the material culture of the Paris Basin and northern France in general, especially in pottery traditions, mark the beginning of Middle Neolithic II, with the development of the northern Chasséen culture in the more westerly part. Immediately to the east of the Chasséen the Michelsberg culture develops, with its characteristic round-bottomed undecorated pottery, including the well-known tulip beakers. In fact, the Chasséen and the Michelsberg have many similarities in their material culture and practices, including the extensive construction of enclosures (see below). It is now widely accepted, following Jeunesse[232], that the Michelsberg Culture began in the eastern Paris Basin. However, whereas the northern Chasséen represents continuity from its predecessors in settlement terms, beyond its likely area of origin the Michelsberg marks a new phase of agricultural expansion. It has been unclear whether or not this represents a demographic expansion like the LBK from a single region, or corresponds to the spread of new stylistic norms among groups across a wide area that were beginning to adopt new and expansive settlement strategies and, as a result, also interacting with the forager groups that continued to exist outside the areas of agricultural settlement (see below). Regional assemblages, for example pottery, have broad family resemblances. Where and when the Michelsberg appears, however, it seems to mark the final break with the preceding Danubian tradition inherited from the LBK, in terms of settlement and house form as well as material culture and symbols[233].

By ~6300 BP the Michelsberg had spread to Belgium, the Rhine and the Neckar valley and by 6000 BP it had reached as far as Bavaria and Central Germany and the borders of Scandinavia in the north. Moreover, whether or not it involved the expansion of a population with a single origin, the key process at the heart of the Michelsberg spread was the expansion of human populations beyond the previously favoured patches of loess, although these continued to be occupied; there was an internal expansion into zones between the old LBK settlement areas and an external one to the north. The history of settlement in the area of present-day Belgium provides an excellent example of the process. After the end of the post-LBK Blicquy phase, ~6800 BP, the occupation of the small loess islands occupied by the LBK is followed by a hiatus, or at least much scarcer occupation, until the Michelsberg at ~6300 BP[234]. Instead of small, concentrated settlement enclaves we see a much more extensive and more even distribution of settlement, together with the appearance of enclosures. By ~6000 BP small single-house sites with Michelsberg material culture appear in the sandy lowland sandy region, replacing the final Mesolithic[235].

The expansion is visible at a broad scale in Zimmermann et al.'s map of the 'Young Neolithic', ~6500-5500 BP, in Germany and adjacent areas[236]. In comparison to the preceding phases, it represents an *extensification* of settlement. In the case of the LBK, core settlement areas that include the vast majority of occupation can be defined as those where sites are no more than 4 km apart[237]. For the Michelsberg, however, the corresponding figure is 12.5 km, made up of a lower density of sites within settlement nuclei but also greater distances between those nuclei, each made up of open settlements and an enclosure. In other words, the settlement area was much bigger but, at two different levels, the sites were much more widely spaced than in the LBK.

Despite the evidence for expansion, the Michelsberg Culture has a relatively limited archaeological record. Burials are not common, though a Michelsberg long barrow over 15 m in length containing two burials is known from the site of Beaurieux in the Aisne valley[238]. A cemetery of more than twenty burials is also known from Gougenheim in Alsace[239]. The best-known sites are the enclosures already mentioned (see below); open sites are not common, though this is likely to reflect at least in part the nature of the evidence, which mainly consists of groups of pits with little evidence of houses; these were presumably of lighter construction than the post-built houses of earlier periods. According to Seidel[240], the groups represented by the open settlements were small and short-lived, which suggests that Michelsberg extensification was associated with greater settlement mobility than in earlier periods.

About 120 Michelsberg enclosures are known[241,242], most of them no bigger than 40 ha, but there are four giant ones, including Urmitz in the Rhine valley, which encloses nearly 1 sq.km. Enclosures are found throughout the Michelsberg distribution area, though more densely in some areas than others, and their construction continues from the earliest ones in the Paris Basin ~64/6300 BP, until ~57/5600, though they are both more numerous and larger in the period down to ~6000. Like the settlements, they were often relatively short-lived, such as the Klingenberg in SW Germany, ~5800 BP.

For the Rhine area, Zimmermann[243] calculates that each enclosure would have had a settlement catchment area of ~1,000 sq. km, giving ~6,000 people per enclosure area on the basis of his population density estimate for the period. It may be that they are an indicator of inter-group competition in regions where population was high relative to the requirements of the extensified subsistence system, and where competition for expansion was occurring. Zimmermann suggests that the enclosure catchments reflect the size of the largest collectively acting group, which might be linked to the scale of group needed for defensive reasons, and some authors see strong indications of violence in Michelsberg contexts. A find in a pit from the Michelsberg site of Bergheim in eastern France of the simultaneous burial of seven individuals with their left arms amputated and evidence of violent blows to the head in one case, is seen by the authors of the report as evidence of inter-group armed violence and they point to other examples in the bones from enclosure ditches as well as skull finds that have been interpreted as trophies[244]. It seems that there were concentrations of enclosures at the Michelsberg colonisation frontier, for example on the northern edge of the German Mittelgebirge[245], which might be an indication that at least on occasion colonisation movements were undertaken by relative large and organised groups as opposed to small numbers of households.

Despite these indications of higher order organisation, there is little evidence for social inequality in the Michelsberg Culture, for example in burials or differences in the material culture between enclosures and open settlements, and also little sign of the circulation of valuables**.**

##### 2.7.1 Genetics

Some clarification on whether the Michelsberg represents a demographic expansion or simply a spread of styles and practices by interaction between existing local communities has previously been given by an analysis of ancient mtDNA from individuals buried with Michelsberg material culture from the cemetery of Gougenheim[239]. Here 46 individuals were found buried in 30 circular pits. The analyses revealed a break in haplogroup frequencies and haplotype composition between the Michelsberg samples and those of the preceding LBK and LBK-descended individuals included in the analysis. The difference can be accounted for by greater similarity with individuals from the Paris Basin Middle Neolithic, including a strong presence of hunter-gatherer mtDNA lineages in both. This evidence is in keeping with the hypothesis that spread of the Michelsberg culture was connected to some degree of population movement eastwards from the Paris Basin. It also fits in with the mtDNA evidence from the site of Gurgy 'les Noisats' that Middle Neolithic populations in the Paris Basin[246] were mixing with local hunter-gatherers and that their farmer DNA came from the Mediterranean stream of the farming expansion.

##### 2.7.2 Settlement and subsistence

So what accounts for this major settlement and demographic expansion in the late 7$^{th}$ millennium BP? While some authors argue for a continuation of the intensive LBK farming system, others suggest a system based on mobile slash-and-burn farming, in which the burning of the cut-down trees provides a short period of soil fertility after which crop yields fall and it is necessary to repeat the process elsewhere. Those emphasising the continuity of the LBK system have argued that the pollen evidence showing a coincidence of clearance and burning events is not sufficiently fine-grained in its chronological resolution to be decisive on this question (pg. 26 in ref [247]). Schier[248], on the other hand, sees a change both in social patterns and in the farming system and sees the key to expansion is a shift from intensive cultivation of the best soils to extensive slash-and-burn that made it possible to exploit soils of lower quality, but only for short periods before fertility fell; it was a form of farming that would have paid off in areas that were otherwise suboptimal. It would have led to lower population densities in the occupied areas than previously but greatly increased their extent, the situation suggested by Zimmermann's maps. The argument is supported by the existence of evidence for clearance, including burning in many cases, in pollen diagrams and soil profiles from northern Germany from the late 7th millennium BP.

The resulting large areas of regenerating forest would have been available for grazing and foddering animals, and it is on the animal husbandry dimension, rather than slash-and-burn cereal agriculture, that the pollen analyst Kalis[249] places the emphasis in explaining the new developments of the Michelsberg subsistence system. He sees it as involving a new style of cattle keeping, on a larger scale, that made major inroads into the forest for forest pasture, reflected in the late 7th millennium BP elm decline and other changes (see also pg. 92 in ref [250] for examples); the existence of micro-charcoal in some pollen diagrams could also be related to burning to improve pasture. Apart from the elm decline, the pollen spectra also show an unprecedented frequency of hazel, alder, birch and ash, pioneer species that are quick to colonise areas that have previously been cleared. From ~5800 BP there was a peak in hazel pollen that lasted 300 years, resulting from the destruction of the tree layer over wide areas and the creation of a scrubby landscape.

As far as crops are concerned, emmer, einkorn, tetraploid naked wheat and naked barley are the cereals found in Michelsberg contexts, while there are only rare occurrences of the legumes pea and lentil and no evidence of oil/fibre plants (pg. 94 in ref [250]). This crop cultivation pattern is similar to that of the preceding post-LBK cultures of eastern France and western Germany, being more diverse than the LBK and with greater emphasis on naked wheats and naked barley[251]. Both these crops have their origin in Mediterranean France and Spain and are indicative of southern connections[252,253]. The tetraploid naked wheat and naked barley are also something the Michelsberg culture also has in common with the TRB Early Neolithic in southern Scandinavia[254].

The contrast between glume wheats and naked wheats is a significant one since it has implications for crop processing and storage practices, in that the former 'are usually stored in spikelets, dehusked on a daily basis and often consumed differently (e.g. as bulgur) while free-threshing cereals are usually processed in bulk, stored as grain and more often transformed into flour or semolina' (pg. 240 in ref [252]).

Why the Michelsberg expansion occurred, into areas whose agricultural potential was generally worse, in terms of temperature, rainfall and soil conditions, than the LBK regions, and why it happened at this time, is not at all clear then, but it clearly represented an overcoming of the limits of the LBK system. Given that the process did indeed start in the Paris basin and that it involved the use of crops of Mediterranean origin, it may represent the adoption of models that had already been established in the Cerny of northern France, derived from forms of subsistence that had proved successful in the northward expansion of Mediterranean farmers, which seems to have been much less closely tied to the distribution of loess soils.

As we have seen, the nature of the farming system remains a matter of debate, so it is hard to evaluate what the costs and benefits would have been in relation to the previous system. Changing household labour budgets may be relevant. Kerig (pers.comm.) has suggested that the increased use of naked as opposed to hulled cereals that one finds in the Michelsberg and related cultures would have led to a massive saving in household energy budgets because dehusking was no longer needed, and this would have opened the way to taking up other activities. What does seem clear is that the new system involved the creation of large areas of secondary forest and was more extensive than the LBK system, because settlement was at lower density and more spatially dispersed. In other words, it was a strategy that was successful despite decreased returns per unit area, perhaps because returns per unit time increased. Population increase was not upwards, increasing in density, but outward, with expanding frontiers.

This expansion had important consequences. While in the southern half of France it is likely that forager-farmer interaction was already going on by 7000 BP if not before, it was not until the Michelsberg, three-quarters of a millennium later, that farmers in the broad region of west-central Europe were brought into more intense contact with foragers, reflected in the increased evidence of genetic admixture into farmer populations as foragers joined farming communities, but also in an apparent symbiosis in which some foragers maintained their way of life but became linked to farming groups, as the Blätterhöhle evidence suggests[255]. It was this new system that provided the basis for the colonisation of southern Scandinavia, by people whose genetic background now included the results of forager-farmer interaction.

In contrast to north Germany and Scandinavia, where growth continued, a decline in the Michelsberg subsistence system is seen in the pollen diagrams, with large-scale reafforestation taking place almost everywhere in the Michelsberg distribution area in the mid-6$^{th}$ millennium BP[249]. Significant population decline is also implied by Zimmermann's settlement map for the Rhine valley and southwest Germany in the period 5500-4800 BP (fig. 12c in ref [236]), which shows very few settlement concentrations even though the region was not altogether devoid of occupation.

#### 2.8 The Neolithic settling of Scandinavia

*Bettina Schulz Paulsson*

Compared to continental Europe, the Neolithisation of Scandinavia occurred relatively late, around 6000 BP, with recent archaeogenetic studies showing the advent of farming as the result of a demographic expansion from further south[39,256–259]. These studies link the neolithic Scandinavians to the earlier, central European LBK-related populations[259], but it is unclear where these groups came from, how large they were, on what routes they moved or how encounters with the indigenous communities took place. While the LBK had reached the Baltic sea (only about 100km south of the coast) (pg. 129 in ref [39]), there was a halt of a 1000 years in Northern Germany. A long lasting research debate as to why the Neolithic expansion stopped for such a long time period before proceeding and reaching the Scandinavian countries remains unresolved. Possible explanations included resistance by hunter-gatherer populations or the shortage of loess soils accompanied by societal changes and the formation of post LBK groups[260].

However, the Michelsberg culture of central Europe is regarded archaeologically to be influential on the emerging TRB complex, see for example[241,250,261,262], reflected most in the material culture especially in the ceramic characteristics and forms such as tulip beakers or round-based flasks. Contacts between these groups are demonstrated at sites such as Flintbek in northern Germany, where both early Funnel Beaker and Michelsberg material have been found together[263].

The settling of Scandinavia must have involved several waves of migration. While the earliest known individual with Central European farmer DNA stems from around 5900 BP or even slightly later (Viksø Mose (5050 +/- 15 BP, UCIA-232706, *3934-3774 cal. BC (95.4%; 3880-3782 cal. BC, 68.2%),* earlier pioneering groups, scouts, contact zones and/or intercultural exchange have all been discussed. There is some evidence of early dated cereals, interpreted as the result of such a pioneering phase before 6000BP[264]. In the first centuries, humans were increasingly offered in the bogs or socially sanctioned. This practice seems to have developed in Southern Scandinavia, either within these groups or as a reaction to encounters and conflicts with indigenous people in these early phases[188].

Early in the neolithization process, the construction of the first enclosures in the form of ritual mega-sites and central meeting places took place. The function of these enclosures is still highly debated, as is their importance to the ritual culture. What is certain, however, is that these sites were only used for very short periods of time or single events every two or three generations. Recent data place the first enclosures like Liselund in Denmark slightly after 5800 BP[265,266]. In Scandinavia, two different forms of these sites can be distinguished: the earlier Sarup enclosures and the more recent palisade enclosures, which are documented mainly in southern Sweden, east of the Storebælt and on Bornholm. The Sarup enclosures (named after the Sarup site on Funen) are characterised by linear or circular ditches that range in size from 1.6 to 20 hectares[267–269]. At the bottom of the ditches, generally there were pottery fragments, tools, bones of animals and humans, and layers of dark organic material found. These ditches were open for short periods and then carefully refilled, partly on a relatively regular basis (pg. 12 in ref [268]), but only every 50 to 100 years[270]. Interpretations of the use of these enclosures are varied, ranging from marketplaces to fortified settlements or ritual centers. For a long time, the role of the Sarup enclosures in burial rituals has been discussed due to the finds of single human bones in some of the ditches. In this hypothesis, the ditches could have served as the primary burial place. After the soft tissue had decomposed, the bones were recovered and brought to their final resting place in a megalithic grave.

Monumental and megalithic tombs, in contrast, were built slightly later from c. 5700 BP onwards. The appearance of these graves in Scandinavia was considered to be part of the expansion of a European movement (e.g. refs [1,271], pg. 206-219 in ref [272]) and various waves of migration in which monuments such as long barrows and enclosures spread.

The first monumental graves in the form of long barrows reached Denmark around 5700 BP. They seem contemporary with the British long barrows, which were built within a short time interval of only a few decades[228,273]. More than 70 long barrows are known from Denmark and six from southern Sweden[274–276]. Only a few human remains have been preserved, but through these few cases burial rites can be reconstructed. The graves usually contain one or two individual burials. In some cases, up to four individuals are documented[277,278]. Contemporaneous with the long barrows, or possibly earlier, simple flat graves following Mesolithic burial practices are recorded[275,279].

A new megalithic era, marked by the construction of the earliest megalithic tombs, emerged in Scandinavia around 5600 cal BP[280]. Similar developments occurred contemporaneously, or at least within the same 100–200 years, across the wider Funnel Beaker region, including northern Germany suggesting that the spread of megalithic architecture formed part of a broader supra-regional phenomenon. .

Most megaliths in Scandinavia are recorded from Denmark, and more specifically from East Jutland, Southern Denmark and from Zealand. Graves are also known from the Swedish coastal regions and from a dense concentration inland in Västergötland[281,282]. A few graves are also found on the southeastern coast of Norway on the northern fringe of the Funnel Beaker Culture[283]. Today there are 7,000 remaining graves in Denmark, while 600 megalithic graves are registered in Sweden[275,284,285].

The Scandinavian megalithic graves are classified into three main groups: dolmens, passage graves, and the gallery and smaller cist graves (hällkistor/hellekisten). These graves are embedded into a primarily earthen tumulus, partly with an encompassing stone setting. Dolmens are built as to be either non-accessible or simple accessible, and in the regional terminology they are classified as round or long dolmens according to the form of the mound or the stone setting. Passage graves are partly built with dry walls between the orthostats and also, mostly in Denmark, as twin graves. The gallery and smaller cist graves are fully or partially embedded in the ground and may have a so-called porthole.

Compared to the passage graves, the emergence of the earlier Scandinavian dolmens and the associated funeral rites are not clearly defined. For Denmark there are around ten dolmens in which preserved skeletons are recorded[284]. According to a recent dating project, the earliest individuals in the Danish dolmens are from around 5600 cal BP, though some might be even earlier and contemporaneous with the first long barrows[280].

On the Swedish Baltic islands of Öland and Gotland, the earliest dated passage graves were built in the 34th century cal BC (~5300 BP)[271], and were contemporary with the building of megaliths inland in the Falbygden area, Western Sweden[286]. The Baltic graves are located on the west coasts, either clustered together or as single graves. This suggests that the megalith builders had migrated from the south over the sea, bringing the passage grave tradition with them.

Based on the radiocarbon results from birch bark, which was used as filling material in the dry stone walls between the orthostats[271,287], it is clear that the passage grave tradition in Denmark starts somewhat later. Passage graves begin with Hvalshøje and Snibhøj on North and Northeast Jutland and with Raevehøj on Zealand around 5300 cal BP and end with Olshøj around 4900 cal BP[271].

Bogs, lakes and springs served in prehistory as places for votive offerings and are interpreted as having been associated to the supernatural, fertility and seasonal abundance[288–290]. The Neolithic lifestyle in Scandinavia was accompanied with new rituals and depositions in wetlands. These depositions consisted of complete human bodies or human body parts, animals, pottery (containing food), wooden artefacts, amber beads, copper ornaments and flint axes[275,291]. The practice emerges with the first waves of farming and is associated with Funnel Beaker contexts. It continues until 5000 cal BP but then seems to lose importance[290,291].

*To summarise*, the Neolithisation of Scandinavia was not the result of LBK farming communities remaining stagnant along the Baltic coasts for nearly a millennium before expanding northwards. Rather, the available archaeological evidence points to a more complex sequence of movements and interactions associated with Funnel Beaker groups from around 6000 BP onwards. The Funnel Beaker complex emerged from developments rooted in the post-LBK societies of Central Europe, particularly within the Michelsberg sphere. This relationship is reflected not only in ceramic forms and decoration, but also in the appearance of causewayed enclosures.

The enclosure tradition reached Scandinavia during the earliest phases of the Neolithic around 5800 BP and was followed by the construction of long barrows around 5700 BP, dolmens around 5600 BP, and passage graves around 5400 BP. The close similarities in material culture, monument forms, and ritual practices indicate long-distance connections between Scandinavia and regions further south, particularly northern France and adjacent areas. Enclosures, long barrows, and megalithic tombs should therefore be understood as elements of broader cultural traditions that spread across large parts of Neolithic Europe.

The precise mechanisms through which these traditions reached Scandinavia remain difficult to reconstruct from the archaeological evidence alone. Northern Germany likely played an important role as a contact zone and intermediary region, although the archaeological record remains incomplete and human skeletal material is scarce. Sites such as Flintbek demonstrate the existence of mixed cultural contexts and underline the complexity of these processes. Rather than representing a simple northward expansion of farming populations, the Neolithisation of Scandinavia appears to have involved multiple episodes of interaction, mobility, and cultural transmission.

#### 2.9 Globular Amphora Communities (GAC)

*Stephen Shennan*

Traditionally called the Globular Amphora Culture, the GAC represents a network of inter-connected communities stretching from northwest Germany to Ukraine, an area of well over 200,000 sq. km., covering a timespan from ~5200 to ~4700 BP. It has recently been the subject of a comprehensive study by Müller[292], from which the summary below is derived.

The main basis of its definition is a set of vessel forms, especially the Globular Amphora itself, decorative techniques and decorative motifs that occur across this broad area. In keeping with earlier studies, Müller's multivariate analysis shows that the region can be divided into a western group, with its core area in the Middle Elbe-Saale region, and a marked eastern boundary at the river Oder; and central and eastern groups that are less sharply divided from one another, covering most of Poland and western forest-steppe areas of Ukraine. These are argued to correspond to relatively distinct communication networks. A distinctive feature that unites the whole GAC area is the practice of ritual double cattle burials.

Müller's chronological analysis indicates that the earliest GAC sites occur from ~5200 BP in both the Middle Elbe and lower Vistula valleys, i.e. both the western and central groups. From there they spread both east and west by ~5000 BP and by ~4700 BP GAC practices had died out over most of the area concerned, being replaced by Corded Ware traditions.

A key feature of the western GAC is that it coexists with other contemporary groups, such as the Řivnáč culture in Bohemia, the Bernburg culture in the Middle Elbe area and the Wartberg culture in western Germany. In hilltop settlements of these cultures GAC pottery occurs together with the local forms. In the Middle Elbe area Müller's detailed work has shown how Bernburg decorative motifs and their sequential occurrence are matched in the GAC (pg. 338 in ref [292]). However, there are distinct differences in burial practices: in the case of Bernburg walled chamber graves are the usual form of burial while GAC is mainly characterised by single graves.

As far as subsistence is concerned, the evidence indicates cattle herding for both meat and milk as well as a higher proportion of ovicaprids and pigs than other contemporary groups, but little in the way of cereal cultivation. Isotope studies of humans and animals together with raw material distribution analyses point to mobility ranges smaller than 50 km in radius. Settlement traces are slight, made up of pits and pit huts, suggesting that movement was frequent and group sizes small. This is in keeping with other studies cited by Müller suggesting that in areas occupied by the GAC the openness of the landscape had decreased and human impact was lower than had previously been the case.

##### 2.9.1 Ancient DNA results

It had long been assumed that GAC and Corded Ware ceramic traditions were related to one another and when it was first shown that Corded Ware could be linked with people whose ancestry lay in the steppe it was believed that this must also be true of the GAC. However, a series of aDNA analyses from the Polish sites of Koszyce and Kierzkowo[38,293,294] have shown that this was not the case. They fall within the general European Middle Neolithic pattern of a combination of Anatolian Farmer and Western Hunter-Gatherer (WHG) ancestry, corresponding to that of previous groups within Germany and Poland, albeit with exceptionally high WHG proportions. Importantly, at Kierzkowo, Tassi et al.[293] identified a component they called Western Europe Neolithic, which ‘accounts for a large share of the ancestry of individuals such as those from Iberia, La Mina and Els Trocs’.

Similarly high levels of WHG ancestry to the GAC are also found at contemporary and slightly earlier sites in western Germany, at the collective burial site of Niedertiefenbach[295] and the cave burials of the Blätterhöhle[255]. In the former case the date of the admixture is estimated at ~5710-5550 BP, in the latter at ~5940-6170 BP, which in both cases would place the admixture in the context of the Michelsberg culture. In Central Germany the same is true of the individual with ~40% WHG admixture from the Bernburg burial of Esperstedt, contemporary with GAC. Thus, Müller (pg. 328 in ref [292]) suggests that preceding regional groups with high levels of WHG admixture are likely GAC ancestors, paralleling earlier suggestions of TRB and Michelsberg connections made on archaeological grounds: ‘consequently, based on the aDNA analyses, we understand the Globular Amphora phenomenon as a newly emerging, everyday social practice developed by parts of the indigenous population of Central Europe’ (pg. 336 in ref [292]). An exception, however, is the case of Bohemia; here it could be shown that the appearance of GAC communities represented an incursion from the north and that they were genetically distinct, in terms of the amount and source of WHG ancestry, from individuals of the Řivnáč culture with whom they interacted[296].

The implications of this are that the pre-existing TRB societies underwent some sort of split, with some communities continuing the previous cereal farming and cattle-herding way of life and others becoming more mobile, but the two continued to interact and influence one another as indicated, for example, by the presence of both Bernburg and GAC material at the enclosure site of Halle-Dölauer-Heide in the Middle Elbe-Saale region and the parallels in decorative motifs between the two. At the same time new burial practices in the form of crouched inhumations in pits developed as well as the distinctive new ceramic styles epitomised by the Globular Amphora vessel itself. However, this does not imply uniformity. As innovations spread across and beyond the Middle Elbe and Lower Vistula regions considerable local variation developed.

Müller proposes that previous centralising tendencies in TRB societies and the investment of surplus in grave monuments might have led to increased social tensions which in the more pastorally oriented areas resulted in increasing economic and ritual differentiation from more agriculturally focussed groups and the development of new interaction networks. The decreased population after ~5200 BP could be a result of this more pastoral and mobile way of life.

#### 2.10 Neolithic expansion and megaliths in Northwestern Africa

*Bettina Schulz Paulsson*

##### 2.10.1 Pioneering phase and evidence for early Neolithic settlers

The Maghreb consists of a vast Saharan hinterland in combination with a Mediterranean climatic zone of approximately Iberian extent, and it provides the narrowest maritime crossings between North Africa and southern Europe. During the early Holocene, a diversity of hunter-gatherer groups, exemplified by the Capsian culture of the eastern Maghreb, is attested[297]. These were succeeded by the westward expansion of pastoralist lifeways and, in the northwest, by the introduction of a Mediterranean agro-pastoral Neolithic closely linked to contemporaneous developments across the Strait of Gibraltar[298]. In the coastal Maghreb, Cardial ceramics and early radiocarbon dates have been recovered from two sites: Kaf Taht el-Ghar and Ifri Oudadane[163]. The available evidence does not indicate that the Neolithic emerged here earlier than in Iberia. The earliest reliable date for the regional Early Neolithic remains 6350 ± 85 BP (Ly-971/OxA), obtained from a carbonised cereal grain at Kaf Taht el-Ghar[299]. At Ifri Oudadane, the earliest secure radiocarbon determination derives from a goat bone (6136 ± 34 BP; OxA-23528), establishing the presence of domesticated animals in the coastal Maghreb around, but not prior to, 7200 BP.

Subsequent to this horizon, research evidence becomes sparse. Given the scale of contemporaneous developments across the Mediterranean—from Dynastic Egypt to the Iberian coasts—this paucity of data more likely reflects a research gap than an actual absence of occupation or activity. The north-western Maghreb, situated at the confluence of Atlantic and Mediterranean climatic regimes and in close proximity to the Atlas Mountains, benefited from relatively stable precipitation. This setting buffered the region against the progressive aridification that affected much of North Africa from the fourth millennium BC (~6000 BP) onwards[300]. In addition to a few isolated and poorly dated finds, the long-recognised distribution of mid- to late-third-millennium BC (~4500-4000 BP) pottery and metalwork of south-west European Beaker types in the Maghreb, together with the presence of African ivory and ostrich eggshell in Copper Age Iberia, clearly indicates intercultural exchange around the Western Mediterranean in this time horizon[301–303].

A recently investigated site is Oued Beht in Morocco, which provides evidence for farming, aggregation, and storage. Oued Beht has been interpreted as part of the wider western Mediterranean exchange networks of the fourth millennium cal BC (~6000-5000 BP), with possible parallels to the so-called "silo culture" known from western Andalusia to the Portuguese Alentejo[297]. Situated at 170–210 m above sea level within a Mediterranean-type, semi-arid bioclimatic zone[304], the site lies c. 100 km inland. Maritime access may have been closer in prehistory. The settlement includes several pits, silos, and linear structures. Thirteen radiocarbon determinations, all on charcoal or seeds, place the occupation between 5400 and 4900 BP (95.4% probability). The material culture consists of macroliths, adzes, axes, grinding stones, and late Neolithic ceramics, some of which are decorated with painted motifs.

Macrobotanical remains are naked barley (hordeum vulgare var. nudum), wheat (*Triticum* sp), pea (p*isum sativum*), wild pistachio, (p*istacia atlantica/terebinthus)*, and oleaster/wild olive (o*lea europaea* var. sylvestris). Faunal remains attest to cattle (*Bos taurus*), sheep (*Ovis aries*), and goat (*Capra hircus*)[297].

##### 2.10.2 Megaliths

In the Maghreb there are several thousand megalithic tombs which remain poorly researched. While the numerous megalithic tombs in the eastern Maghreb date to the Late Bronze Age and the 1st millennium BC (~3000-2000 BP), there are a few indications available that some of the structures in the Western Maghreb are probably associated to the Neolithic[305–308].

South of there is the cromlech of M'zora or stone circle with 167 stones and unknown dating[309]. In the northeast of Kabylia, i.e., Algeria, there are a few gallery graves that appear to be unique in the entire Maghreb and seem to be older. However, precise datings are not known[307]. There is a general consensus that most of these monuments were constructed during the Late Bronze Age or Early Iron Age and were subsequently re-used during the Punic and Roman periods[310] (pg. 402–415 in ref [306]).

In the Sahara, earlier monuments have been recorded in regions such as Libya, northern Niger, northern Chad, southeastern Algeria, Mauritania, southern Morocco, and Western Sahara, dating between 6000 and 4500 BP[310,311]. This Saharan megalithism is regarded as an indigenous North African development among pastoral groups and, in its later phases, is associated with mobile, trans-egalitarian tribes (pg. 190 in ref [310]).

##### 2.10.3 Genetic results

Recently, the previously scarce archaeological evidence has been complemented by genetic analyses demonstrating a mixed ancestry comprising local hunter-gatherers, Neolithic groups, and Saharan pastoralists[312,313]. The Neolithic genetic component has been interpreted as deriving from the Iberian coast, supporting theories of trans-Gibraltar gene flow. Furthermore, the discovery of an individual of African descent buried at the Late Copper Age (Beaker period) site of Camino de las Yeseras in the Iberian interior provides clear evidence of long-distance mobility and intercontinental migration[314].

To summarise, the Cardial expansion reached the Maghreb by around 7200 BP. Recent genetic evidence demonstrates sustained contacts across the Strait of Gibraltar, with Neolithic ancestry in north-western Africa likely deriving in part from Iberia. While there is currently no direct evidence linking European and North African megalithic traditions, the documented genetic and cultural connections across the western Mediterranean make such a link plausible.

#### 2.11 Circulation and prospection of green stone and obsidian in Western Europe 8000-6000 BP

*Bettina Schulz Paulsson & Stephen Shennan*

##### 2.11.1 Introduction: the green stone trade in Europe

Evidence for short- and long-distance contacts, based on the dispersal of raw material, prestige artefacts, and domesticated resources is common in Western Europe 8000 BP-6000 BP. Especially research on jadeite and variscite artefacts and the provenience of these rare green stone materials has led to completely new insights into the value system of the societies of Western Europe 8000-6000 BP (among others cf. [29,141–149]). Provenance analyses have demonstrated the significant role of green stones as highly valued raw materials and prestige goods in Western Europe, while in eastern Europe a different value system existed based on copper and gold. Jadeite, sourced from Northern Italy, circulated widely across France, the Pyrenees, the Central Mediterranean, and into Central and Northern Europe. Variscite, in contrast, was extracted from southern and western Iberia and mainly distributed as beads within Iberia and Northwestern France.

By far the largest accumulation of artefacts made from jadeite and variscite in the form of prestige axes, beads, and pendants is documented from the Morbihan in Brittany, Northern France and there especially from the region of Carnac. Eleven axes made from alpine jadeite are documented from the first stage of the St. Michel tumulus. From Tumiac, 15 jade axe heads made from jadeite and eclogite and one made from nephrite are recorded. From Mané er Hroëck, 11 jade axes are documented. Artefacts made from variscite are recorded from the St. Michel tumulus, including 101 beads and nine pendants. From Tumiac, 237 beads and 12 pendants and from Mané er Hroëck 44 beads and two pendants are registered. An important fact is that the variscite pendants and the jadeite axes are only recorded and diagnostic for the early funeral contexts in Brittany described until now and they are also mostly from the 'tumuli Carnacéens'[153] (pg. 173 in ref [150]). The transfer of early megaliths in the second half of the 5th millennium cal BC (6500-6000 BP) outgoing from the Morbihan to the Mediterranean and possibly to the West Iberian Peninsula aligns with patterns of green stone exchange networks and prospection for the minerals[1].

##### 2.11.2 Jadeite and jadeite trade 8000 BP-5000 BP

Quantitatively the most important green stone objects that were exchanged and deposited in western Europe from the mid 8$^{th}$ millennium BP to the end of the 6$^{th}$ millennium were axes and adzes of jade, used here as a general term covering a number of different types of very hard green metamorphic rock, including jadeitite, omphacite and eclogite, from the Alps. Their production and distribution has been identified and presented in a remarkable series of monumental volumes by Pierre and Anne-Marie Pétrequin and their colleagues[142,315,316]. The account below largely summarises key aspects of Pétrequin and Pétrequin[142].

Axes and adzes were the fundamental practical tools of the Neolithic, for clearing forest, but the extraordinary quality of some of them, the enormous time and effort involved in making them and the long distances that many of them travelled make clear that they were also much more than that. Many different rocks had the relevant qualities for making axes (see e.g. pg. 63 in ref [142] for France) and axes made from them circulated at a regional scale, up to 100-200 km from their source. However, the circulation of the jade axes from Monte Beigua and Monte Viso in the northwest Italian Alps was of a different order. It covered the western half of Europe as far as Denmark and Scotland, defining an area where jade axes circulated that was differentiated from the eastern half of Europe where copper was the prevalent high-value material (e.g. fig. 32 in ref [142]). To the east there are only occasional jade finds, the most significant exception being their occurrence at the famous cemetery of Varna and other sites in Bulgaria[317].

The earliest evidence of the use of the Monte Beigua jade source, which was not far from the coast, comes already from sites of the earliest Impressed Ware Neolithic colonists of the western Mediterranean, for example Portiragnes, southern France, at 7700-7600 BP, but this was for utilitarian items only. Monte Viso is less accessible and the main jade sources are found between 2100 and 2400 metres (pg. 105 in ref [142]). Exploitation here, to produce large roughouts of exceptional size and quality, requiring a very high level of expertise, began at the end of the 8$^{th}$ millennium BP. These were taken down to the valleys for further working, following expeditions into the mountains to obtain the raw materials. It seems that these were subject of systematic prospecting.

The earliest large polished form identified is the Bégude-type adz (pg. 119-133 in ref [142]), which began to be produced ~7000 BP and was distributed very widely, especially across the Alps in a north-westerly direction as far as northern France and Brittany (map fig. 63 in ref

[142]), mainly from the Monte Viso source; some were further polished in the course of their journey. Two large blades were found in a burial in one of the earliest Carnac monumental burial mounds in southern Brittany, ~6600 BP, associated with four disc-rings, also of Alpine origin, which Pétrequin and Pétrequin (pg. 143 in ref [142]) see as representing symbolic copulation.

From the 66th century BP, although significant numbers occur in the Paris Basin, the key destination for large high-quality polished jade blades was southern Brittany, as witnessed by the large number of items that ended their circulation there in the extraordinary burials in the monumental Carnac mounds, such as Tumiac, and in other apparently ritual contexts, sometimes after multiple episodes of reshaping and repolishing (page 223 and 263 in ref [142]). The requirements of the sacred elites that in the Pétrequins' view created these monuments exercised a pull over the long-distance circulation of the jade axes of the finest materials, via a network of high-status groups or individuals. Even at the source, they suggest, these materials were considered sacred and not destined for normal use.

From the 63rd century BP, however, the situation in Brittany changes. Pétrequin and Pétrequin(pg. 359 in ref [142]) point to the mound of Moustoir at Carnac. The mound was still large but there were far fewer grave goods and they did not include any jade axes. On the other hand, they did include two long blades of flint from the Grand-Pressigny source 300 km to the east, and a ceramic so-called 'vase-support' or 'incense burner', a decorated vessel made of a base with a shallow dish-like top apparently used for ritual purposes and widely distributed in Brittany. This is a form associated with the Chasséen culture (see relevant section).

At the same time as jade axes were disappearing in Brittany, we see an expansion of their distribution northward from the Paris Basin(fig. 227 in ref [142]), linked in its space-time distribution to the expansion of the Michelsberg culture (see section 2.7). Somewhat later they appear in Britain and Ireland, associated with its farmer colonisation. In both regions, Pétrequin and Pétrequin suggest (pg. 375 in ref [142]), many of the axes seem to be of forms that had already long been in circulation in the Paris Basin, perhaps pointing to their being heirlooms, but they circulated alongside novel forms, in particular the Puy type. The earliest date for this type is in a Chasséen context dated 6301-6202 cal BP and it continues in circulation down to the mid-6$^{th}$ millennium BP (pg. 375 in ref [142]). An example of this type, of Monte Beigua jadeitite, even occurs in a hoard in Denmark (pg. 377 in ref [142]). The numbers in Britain are very small, perhaps a consequence of a decline in contacts between Britain and the continent after the initial period of colonisation.

In the course of the 6$^{th}$ millennium the social significance of jade axes decreased, and Puy-type axes gradually disappeared. At the same time, production quality declined and a wider range of rocks of lesser quality were used. Jade axes continued to be used but for utilitarian purposes and copper became increasingly important.

##### 2.11.3 Variscite and variscite trade in Europe 8000-5000 BP

Variscite, also called *callais*, is an extremely shiny green mineral used for jewellery in the Neolithic. It is a relatively rare phosphate mineral in Europe and at present there are not more than ten geological deposits known on this continent plus some indications for sources (pg. 120 in ref [318]). Most of the variscite sources in Europe are concentrated in the Northwest of the Iberian Peninsula at El Bostal by San Vincente de la Cabeza and at Palazuelo by Zamora (Supplementary Fig. S2.4). However, several additional variscite deposits or indications for deposits are known from this region[319,320] (pg. 120 in ref [318]; Fig. 16 in ref [321]). From the southern and the southwestern part of the Iberian Peninsula in Andalusia, Pico Centeno/Encinasola and La Encantada are recorded and in the very east of the Iberian Peninsula in Catalonia Can Tintorer by Gavà, Barcelona[322,323] (pg. 19 in ref [324]). More variscite deposits are available in Pannec´e in Northwest France and indications for sources are also registered in the Middle French Montrebras and in South Sardinian Sarrabus.

The only prehistoric mining activities so far documented are in the Iberian Peninsula at Can Tintorer/Gavà (Barcelona), Palazuelo de las Cuevas (Zamora) and Pico Centeno/Encinasola (Huelva)[151,324].

Variscite was used mainly for beads and pendants deposited in funerary contexts, including individual cave burials, collective graves, hypogea, cists and megalithic tombs. Non-destructive PIXE analyses indicate that variscite was extracted from Iberian sources from the late eighth millennium BP onwards and circulated over considerable distances[151,318].

###### 2.11.3.1 The early variscite network in Western Europe 8000-6000 BP

The results suggested that variscite has been extracted since the end of the 8th millennium cal BP from deposits located in the Iberian Peninsula. Decisive support was found for long-distance contacts and the circulation of variscite during the early time horizons. The movement of variscite has occurred across land and/or seaways to Northern France.

The earliest ornaments, dating to approximately 7300 to 6800 BP, occur in cave and collective burials in Aragón and Navarra. Their raw material may have originated from Can Tintorer/Gavà. By around 7000 BP, variscite from Pico Centeno/Encinasola in Andalusia had reached Plichancourt, Le Monts, in eastern France, providing early evidence of long-distance circulation.

In north-western France, the sequence begins with the earliest phase of the large tumuli in the Morbihan, first with the tumulus du Mane er Hroech in Locmariaquer[325] and then Saint-Michel around 6700 BP. Rich deposits continued until the end of the Ancient Castellic horizon, around 6300 BP.

The provenance analyses identified Pico Centeno/Encinasola as the source of the variscite found in these northern French contexts. As no securely identified intermediate finds are known, we need to take long distance exchange or import into consideration over maritime and riverine routes.

Between approximately 6750 and 6000 BP, large quantities of variscite from Can Tintorer/Gavà were deposited in Catalan caves. Mining at Can Tintorer/Gavà reached its peak between about 6050 and 5450 BP, when variscite and turquoise ornaments were increasingly placed in hypogea, cists and megalithic graves. In southern France, variscite circulated from the Ancient Chassean horizon, around 6300 to 6000 BP. Elsewhere in Iberia, its wider funerary deposition became visible from approximately 6200 to 6000 BP onwards.

###### 2.11.3.2 The later variscite network in Western Europe, 6000 to 4000 BP

After 6000 BP, long-distance variscite exchange with the Morbihan declined. Variscite beads continued to occur in passage graves there, but never in quantities comparable to the earlier deposits. Within the Iberian Peninsula, however, exchange networks expanded, new sources were exploited and the number of reliable contexts increased. Palazuelo may first be represented as a source in the Río Fortes mound shortly after 6000 BP. By around 5000 BP, Palazuelo and Pico Centeno/Encinasola had become the principal sources supplying Late Neolithic and Chalcolithic communities, while Can Tintorer/Gavà lost much of its earlier supraregional importance[151].

To summarize, both minerals were predominantly exchanged and imported during the 7th millennium cal BP. Primarily, the raw material or the axes were traded into Northwestern France, where it was subsequently worked or reworked into Carnac axes or variscite beads. Prehistoric variscite mining activities are documented exclusively in the Iberian Peninsula; however, the largest concentration of Neolithic artefacts made from variscite *called also callaïs* is found in Morbihan, Brittany, particularly in the Carnac region, providing strong evidence for long-distance trade and the import of the raw material. After 6000 BP, other variscite mines in the Northwestern Iberian Peninsula, such as Palazuelo, gained importance, primarily for the circulation of beads within the Iberian Peninsula.

Although jadeite axes are absent from passage graves in northwest France after ~6300 BP, production at the Alpine sources continued, but the peak of exploitation is over. We find later forms of large polished axes such as Puy type, found as far as Germany and Denmark in the final centuries of the 7th millennium BP, as noted above. Over the course of the 6th millennium BP production declined and was increasingly oriented towards utilitarian items of local distribution (pg. 393-401 in ref [142]).

##### 2.11.4 Obsidian in the West Mediterranean

There are four main island sources of the volcanic glass obsidian that was used during the Neolithic in the western and central Mediterranean: Sardinia, Lipari in the Aeolian islands, Palmarola off the west coast of central Italy and Pantelleria between Sicily and North Africa. Characterisation studies over the last fifty years and more[326] have made it possible to distinguish between them and between different sources within the same island. On this basis it has been possible to show that obsidian mainly from Lipari was already reaching the Adriatic islands and the Dalmatian coast of Croatia in the early Neolithic. Lipari was also the main source for obsidian found in mainland Italy, including the Tavoliere region, the main centre of early Neolithic settlement in Italy, whose origin lies in Croatia. As Tykot (pg. 271 in ref [326]) point out, this implies that people were crossing the Adriatic in both directions.

The maritime nature of farmer colonisation in the West Mediterranean is also reflected in the presence of obsidian from the West Mediterranean islands at the earliest Impressed Ware sites[327]. The site of Arene Candide in Liguria contained obsidian from Sardinia and Palmarola. Further west, on the far side of the river Rhone in Languedoc the adjacent Impressa sites of Pont de Roque-Haute and Peiro Signado, are the earliest known Neolithic sites, dating to ~7700 BP. Pont de Roque-Haute also has obsidian from Palmarola, albeit only in small quantities. Peiro Signado again has obsidian, from both Palmarola and Sardinia.

After this initial appearance in association with the first settlements of incoming farmers in Liguria and southern France it is not found in Cardial contexts of the second half of the 8th millennium BP. It appears that maritime contacts and exchanges were not maintained after the initial settlement. Obsidian then occurs in small quantities as pressure-flaked bladelets in southern France through the 7th millennium, including the earlier Chasséen phases, when most of it comes from Lipari, as it does in Italy during the same period[118]. A bladelet of obsidian from Lipari found at the early Chasséen site of St Michel-du-Touch, over 1200 km from the source, fits into this context.

In contrast, in southern Italy and Sicily, for example on the coast of Calabria, obsidian from Lipari made up most of the local lithic assemblages, implying that it was a utilitarian material and that people could obtain regular supplies, which were in the form of pre-form cores ( pg. 272 in ref [326]); these may have been produced during seasonal expeditions to the island (pg. 521 in ref [328]). At coastal sites further away the proportion decreases to 10-30% and then generally decreases the further you go inland. Pantelleria and Palmarola had smaller distribution areas than Lipari and by the beginning of the $6^{th}$ millennium BP this had become the predominant source in the central Mediterranean, with the exception of Sardinia, which had its own sphere of distribution in the north, visible in southern France.

In the late $7^{th}$ millennium BP and the first half of the $6^{th}$, the classic and late phases of the Chasséen there is a major increase in the number of obsidian finds, now dominated by the Monte Arci source in Sardinia[118], again for pressure-flaked bladelets. This most probably involved direct sea travel from Corsica to southern France as there is no evidence of obsidian from the Sardinian A source at north Italian coastal sites (pg. 376-7 in ref [329]). At this time the distribution even extends as far as Catalonia to the west, where it occurs in a number of graves of the *Cultura de los Sepulcros de Fosa* with other materials obtained by long-distance exchange, including Bedoulian flint, as well as variscite beads from the local Can Tintorer/Gavà mine[330]. The discovery at the late Chasséen site of Les Terres Longues at Trets, east of Aix-en-Provence, of over 4500 pieces of Sardinian obsidian, including blade cores, suggests that it may have been an intermediary distribution site, and points to the highly organised nature of obsidian exchange at this time[331]. Binder et al. (pg. 197 in ref [118]) suggest that, as well as importing obsidian, sites such as this would have been sending heat-treated Bedoulian flint towards Italy; the Grotta all'Onda site in Tuscany, for example, has large quantities of Sardinian obsidian as well as Bedoulian flint. They also point out that obsidian occurs at sites further inland in southern France at this time, especially in places that would have had access to other materials for exchange, such as hard stone for axes.

During the course of the $6^{th}$ millennium BP, however, the role of obsidian declined. It continued in use for some time at sites close to the sources but then even this use eventually faded away.

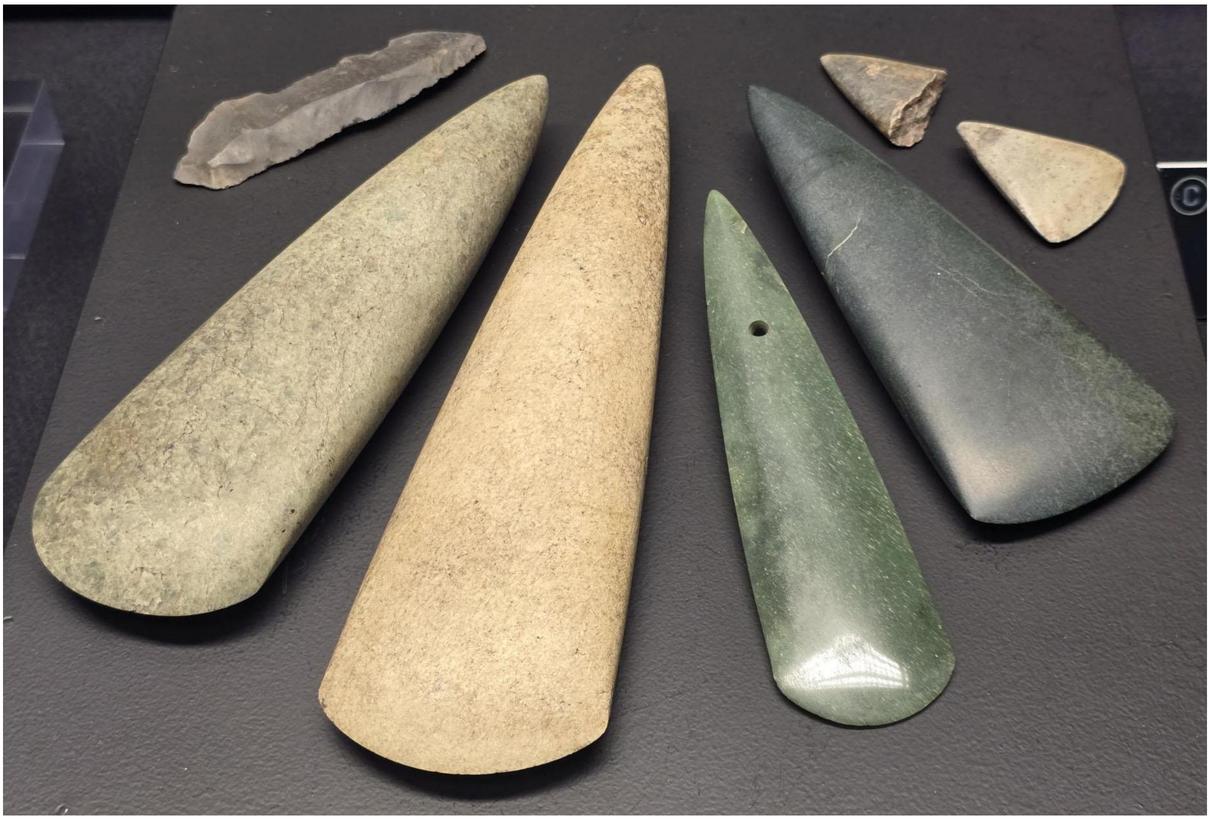

**Supplementary Fig. S2.1. Jadeite axes, museum Carnac Jadeite axes recovered from the monumental tombs of the Carnac region and from regional hoards were manufactured from raw material originating in the Monte Viso massif of the northwestern Italian Alps, providing further evidence for long-distance exchange networks during the Neolithic.** Photo courtesy of Bettina Schulz Paulsson, reproduction with the kind permission of the Carnac museum

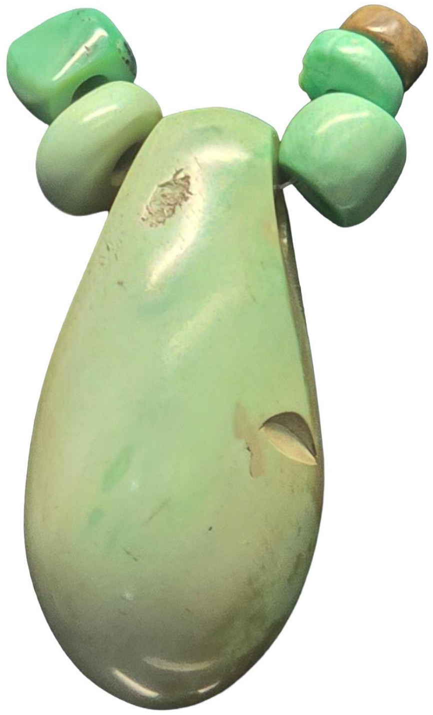

**Supplementary Fig. S2.2. Variscite (callaïs) bead from Er Grah tumulus, Carnac Museum.** The raw material used for this bead, recovered from a monumental burial in the Carnac region, is consistent with variscite sources in southwestern Iberia, providing evidence for long-distance exchange along Atlantic maritime routes. Photo courtesy of Bettina Schulz Paulsson. Reproduction with the kind permission of the Carnac museum.

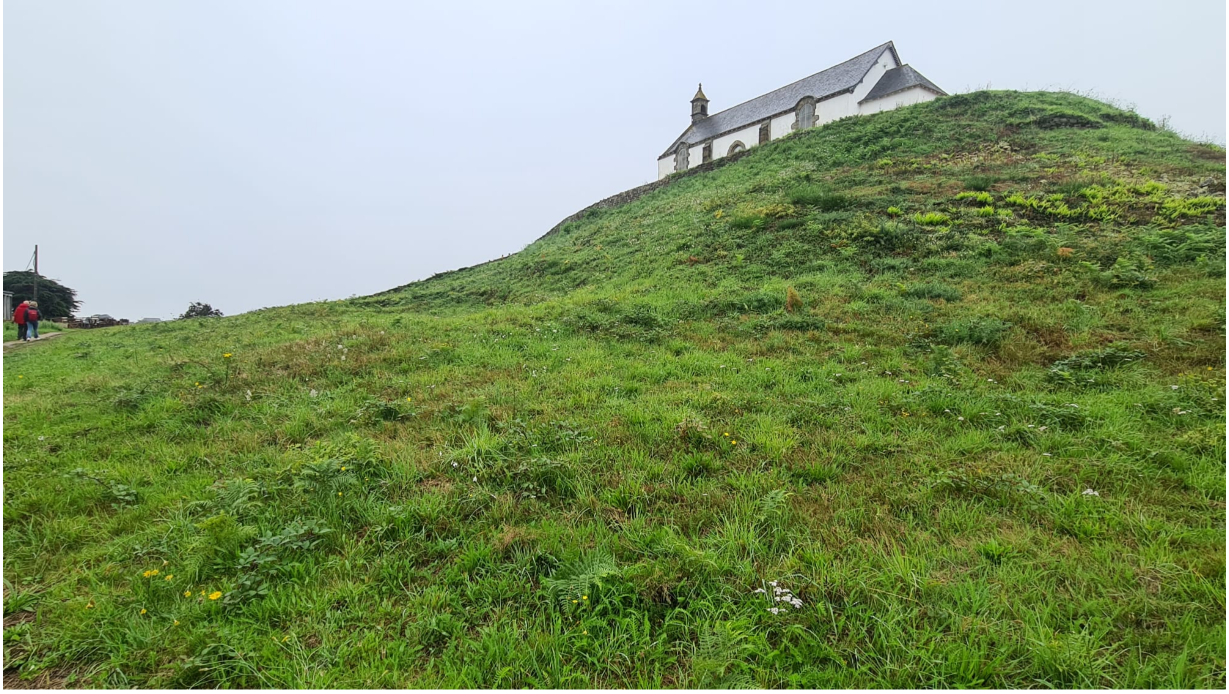

**Supplementary Fig. S2.3. The Saint-Michel tumulus, Carnac. The Saint-Michel tumulus is one of the largest Neolithic burial monuments in Europe, measuring approximately 120 m in length.** Today, it is surmounted by a chapel. The monument contains three funerary structures: a dry-stone chamber, a dolmen, and a later passage grave. Between 6700 and 6200 BP, the Carnac region witnessed the construction of monumental burial mounds (*tumulus carnacéens*), which contained abundant prestige objects manufactured from raw materials imported from the Iberian Peninsula and the northwestern Italian Alps, including variscite, fibrolite, jadeite, and nephrite. These early tombs were generally designed for the burial of only one or two individuals. Photo courtesy of Bettina Schulz Paulsson.

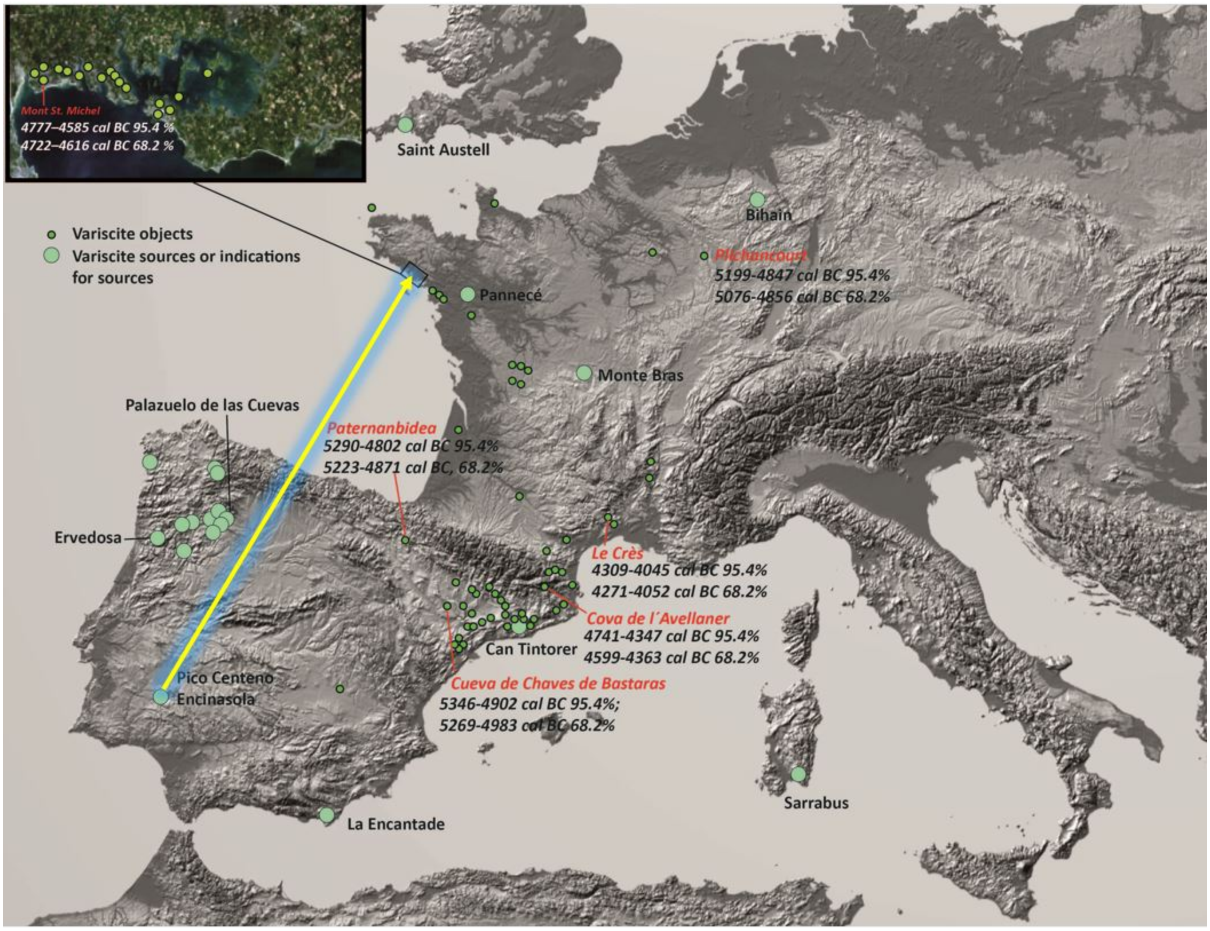

**Supplementary Fig. S2.4. Variscite trade and timeline in Europe (after [1]).**

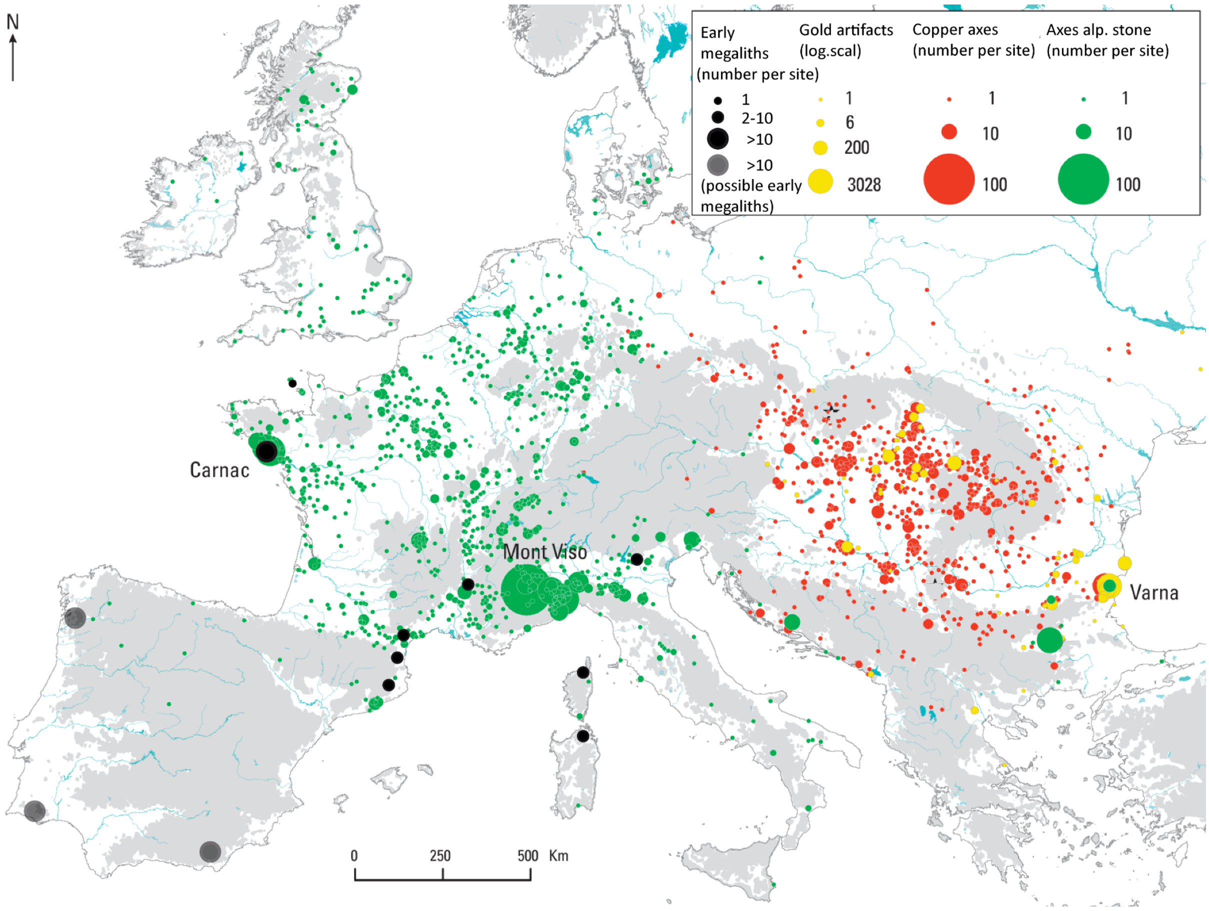

**Supplementary Fig. S2.5. Distribution of jadeite axes, copper axes, gold artefacts, and early megalithic clusters during the fifth and early fourth millennia cal BC (~7000-5500 BP).** Distribution of jadeite axes in western Europe in relation to the Alpine jadeite sources, compared with the distribution of copper axes and gold artefacts in eastern Europe and correlated with the principal clusters of early megalithic monuments[1] (after pg. 193 in [316]).

##### 2.11.5 A Bayesian statistical framework applied on the radiocarbon dates for the greenstone exchange networks in Europe

###### 2.11.5.1 Introduction

For selected key sites that provide insight into exchange networks across Neolithic Europe, Bayesian chronological modelling was used to obtain higher chronological resolution of the radiocarbon evidence. Bayesian modelling integrates radiocarbon measurements, archaeological prior information, and the calibration curve to produce more precise estimates of calendar age ranges than calibration alone. All Bayesian models and calibrated dates were generated in OxCal v4.4 or 4.1[31,32] using the IntCal20 calibration curve[33].

The radiocarbon dates were compiled from published literature and archaeological reports, with most samples derived from charcoal. To account for potential inbuilt age in unidentified charcoal samples, sites dated exclusively by charcoal were modelled using the OxCal Charcoal Outlier Model (*Outlier_Model*("Charcoal", *Exp*(1,-10,0), *U*(0,3), "t")) following Bronk Ramsey[332]. This model assumes an asymmetric exponential distribution of inbuilt age, reflecting the expectation that most charcoal samples are approximately contemporaneous with deposition but that some may derive from older wood. Sites dated by mixed sample materials were modelled using standard Bayesian sequences and phases, incorporating vertical and, where possible, horizontal stratigraphic relationships.

###### 2.11.5.2 A Bayesian statistical framework for the jadeite exploitation at the Monte Viso

The Neolithic jade quarries of Monte Viso (Oncino, Piedmont, Italy) comprise a complex of high-altitude extraction sites that provided the principal source of Alpine jadeitite exploited during the fifth and fourth millennia BC (~7000-5000 BP). Provenance studies of Pierre

Pétrequin and his team have firmly established Monte Viso as the principal source of Alpine jade axeheads distributed across western and northern Europe[142,315,316].

The archaeological investigations identified more than thirty quarrying and workshop areas distributed between approximately 1,500 and 2,400 m a.s.l. The Cercle des Blocs constituted the principal extraction and habitation area, from which surrounding jadeitite outcrops were exploited during seasonal summer expeditions. Quarrying employed fire-setting to detach large blocks of jadeitite, omphacitite, eclogite and fine amphibolite, which were subsequently shaped into roughouts before being transported throughout western Europe; the quarries provided charcoal and datable material. A total of 31 AMS radiocarbon dates are currently available from the Monte Viso quarry complex (pg. 214-257 in ref [316]).

Of these, 22 dates belong to the Neolithic, whereas nine dates fall within the Iron Age, Roman, Medieval and later historical periods, documenting renewed use of the mountain long after prehistoric quarrying had ceased. Four quarry areas yielded exclusively post-Neolithic dates, whereas five sites produced exclusively Neolithic ages. Only the Cercle des Blocs preserves evidence for both Neolithic exploitation and later reuse.

To establish a robust chronology for the principal phase of jadeite extraction, we focused on modelling the 22 Neolithic radiocarbon dates within a Bayesian framework. The model groups the individual quarry areas as unordered sub-phases within a single overarching phase of exploitation, allowing estimation of the onset, duration and end of the main quarrying episode.

Because all dated samples consist of unidentified charcoal, potential inbuilt-age effects cannot be excluded. The quarry sites are located close to or slightly above the upper conifer treeline, which during the Neolithic was dominated by Norway spruce (*Picea abies*) and silver fir (*Abies alba*) rather than by larch as today[316]. Although fuelwood was probably collected locally during seasonal summer expeditions, both spruce and silver fir may attain considerable ages. We therefore evaluated the robustness of the chronology using the OxCal Charcoal Outlier Model[332] and comparing it to a standard model.

Application of the Charcoal Outlier Model had only a minor influence on the Bayesian chronology. Both models showed excellent agreement (Charcoal model: *A*model = 99.2, *A*overall = 99.3; standard model: *A*model = 96.0, *A*overall = 96.1).

Comparison with the standard Bayesian model demonstrates that accounting for potential inbuilt-age effects produces only a slight shift towards younger ages for the end boundary, while leaving the overall chronology virtually unchanged. The close agreement between both models demonstrates that the inferred chronology of jadeite exploitation at Monte Viso is robust to the inclusion of a charcoal outlier model.

Results:

The Charcoal model estimates the start of the main exploitation phase at 5457–5103 cal BC/7407–7053 cal BP (68.3% probability) and 5580–4876 cal BC/7530–6826 cal BP (95.4% probability), with the end of exploitation estimated at 3911–3746 cal BC/5861–5696 cal BP (68.3%) and 3941–3596 cal BC/5891–5546 cal BP (95.4%). This corresponds to a modelled duration of approximately 1,273–1,661 years (68.3%) or 1,039–1,850 years (95.4%).

Despite this long modelled duration, the radiocarbon evidence shows that exploitation was concentrated in its earlier part, prior to c. 6340 BP. Twenty of the 22 Neolithic radiocarbon dates predate c. 6340 BP, and fourteen of these fall before c. 6700 BP, corresponding to the beginning of the Middle Neolithic and the emergence of the Carnac megalithic centre in north-western France. Only two radiocarbon dates postdate c. 6340 BP, one from Cercle des Blocs (4341–4059 cal BC, 95.4%) and one from Abri Puymirol (3962–3796 cal BC, 95.4%). These younger determinations indicate that jadeite quarrying continued into the later Neolithic, but only on a much more limited scale than during the principal phase of exploitation.

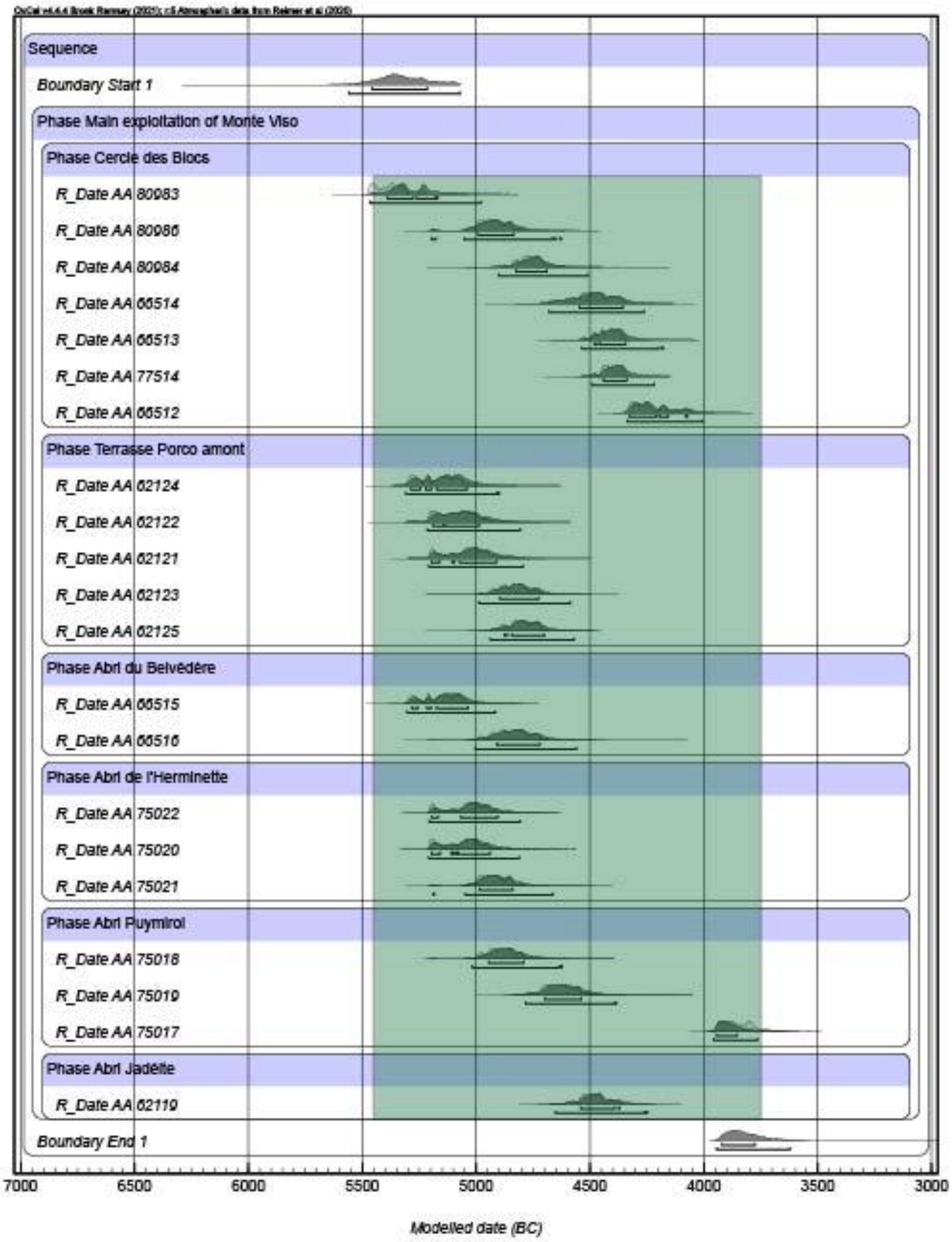

**Supplementary Fig. S2.6. Bayesian chronological model for the main exploitation phase of the Monte Viso jadeite quarries with application of the Charcoal Outlier Model [332].** Radiocarbon dates from six quarry areas were grouped into unordered site-specific subphases within a single overarching phase. *A*model = 99.2, *A*overall = 99.3.

2362

```
Plot()
{
 Outlier_Model("Charcoal",Exp(1,-10,0),U(0,3),"t");

  Sequence("Monte Viso")
  {
       Boundary("Start main exploitation");

       Phase("Main exploitation of Monte Viso")
       {
       Phase("Cercle des Blocs")
       {
       R_Date("AA 80983",6401,46){Outlier("Charcoal",1);};
       R_Date("AA 80986",6041,44){Outlier("Charcoal",1);};
       R_Date("AA 80984",5898,44){Outlier("Charcoal",1);};
       R_Date("AA 66514",5662,71){Outlier("Charcoal",1);};
       R_Date("AA 66513",5605,42){Outlier("Charcoal",1);};
       R_Date("AA 77514",5575,41){Outlier("Charcoal",1);};
       R_Date("AA 66512",5393,42){Outlier("Charcoal",1);};
       };

       Phase("Terrasse Porco amont")
       {
       R_Date("AA 62124",6231,48){Outlier("Charcoal",1);};
       R_Date("AA 62122",6146,49){Outlier("Charcoal",1);};
       R_Date("AA 62121",6110,48){Outlier("Charcoal",1);};
       R_Date("AA 62123",5959,49){Outlier("Charcoal",1);};
       R_Date("AA 62125",5931,48){Outlier("Charcoal",1);};
       };

       Phase("Abri du Belvédère")
       {
       R_Date("AA 66515",6222,44){Outlier("Charcoal",1);};
       R_Date("AA 66516",5963,61){Outlier("Charcoal",1);};
       };

       Phase("Abri de l'Herminette")
       {
       R_Date("AA 75022",6107,43){Outlier("Charcoal",1);};
       R_Date("AA 75020",6121,43){Outlier("Charcoal",1);};
       R_Date("AA 75021",6040,40){Outlier("Charcoal",1);};
       };

       Phase("Abri Puymirol")
       {
       R_Date("AA 75018",6005,43){Outlier("Charcoal",1);};
       R_Date("AA 75019",5791,50){Outlier("Charcoal",1);};
       R_Date("AA 75017",5044,37){Outlier("Charcoal",1);};
       };
       Phase("Abri Jadéite")
       {
       R_Date("AA 62119",5665,47){Outlier("Charcoal",1);};
       };
       };

       Boundary("End main exploitation");
       Span("Duration main exploitation");
  };
};
```

**Supplementary Fig. S2.7**. Code of the Bayesian model of the main exploitation phase at Monte Viso based on radiocarbon dates from six extraction areas using the OxCal Charcoal Outlier Model to account for potential inbuilt age effects.

###### 2.11.5.3 Monumental tombs in Northwestern France with a radiocarbon sequence and jadeite and variscite deposits

Reliable evidence for early variscite ornaments and their long-distance circulation has been identified in Eastern France, at Plichancourt/Marne, Le Monts. An Early Neolithic burial containing a variscite pendant has been dated to 5199-4847 cal BC/7149–6797 (95.4%; 5076-4856 cal BC, 68.2%). The source for this variscite has been identified as Pico Centeno/Encinasola in Andalusia[150,151]. The Bay of Morbihan contains not only the highest concentration of Alpine jadeitite axes outside the Monte Viso source region, but also the richest assemblages of variscite ornaments in Europe, deposited from at least 6700 BP onwards during the Middle Neolithic[29]. The securely dated contexts associated with jadeite axes and variscite (callaïs) ornaments in northwestern France are the Carnac tumuli of Mont Saint-Michel, Er Grah and Tumiac.

###### Tumulus Saint Michel

At Mont Saint-Michel, excavations indicate that the earliest monument consisted of a small dry-stone burial chamber, later enlarged by successive construction phases into the monumental tumulus visible today. The central chamber yielded 12 Alpine jadeitite axes and more than 110 variscite beads and pendants, forming one of the richest known Neolithic prestige deposits in Europe[143,333,334]. Five radiocarbon determinations constrain the construction sequence. Two dates from the same cremated human bone place the primary burial and associated jadeite and variscite deposition at 4777–4585 cal BC/6727–6535 cal BP

(95.4%), whereas a later date from the eastern passage grave indicates that the final enlargement of the monument occurred during a subsequent construction phase (4339–3631 cal BC/6289–5581 cal BP, 95.4%).

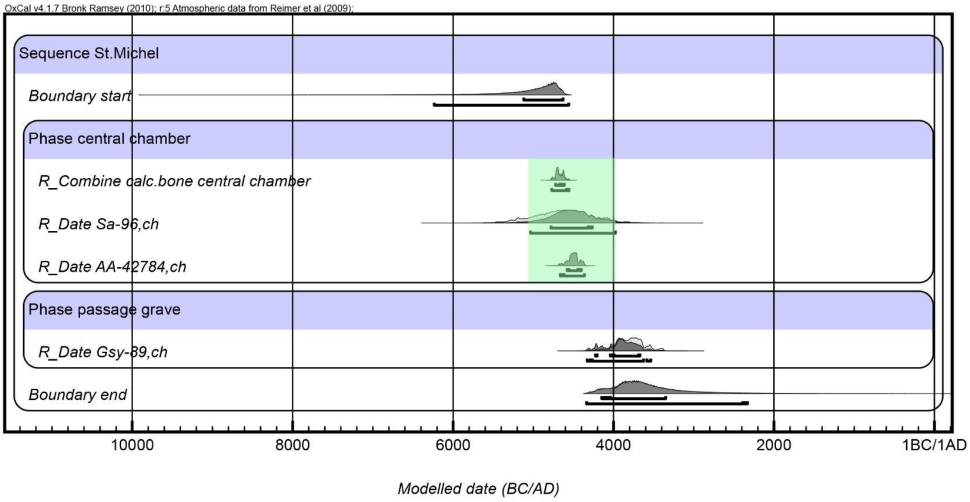

**Supplementary Fig. S2.8. The Probability of radiocarbon dates from the tumulus Carnacéen St. Michel in Carnac.** Model agreement: Amodel: 95.4, Aoverall: 96.7.

The Er Grah tumulus forms part of the major Neolithic monumental complex at Locmariaquer, together with the Table des Marchands and the Grand Menhir. Excavations indicate that the c. 140 m long monument was constructed in at least two phases above an earlier occupation horizon containing Castellic material. Eight radiocarbon determinations are available, although three unreliable or intrusive dates were excluded from the Bayesian model[156].

Pre-monument activities are dated to 5461–4721 cal BC/7411–6671 cal BP (95.4%), including an Early Neolithic hearth and associated Châtelet lithic material. A dry-stone burial chamber in the northern part of the monument, later enclosed within a round and subsequently elongated tumulus, contained a variscite pendant associated with the earliest construction phase. Two radiocarbon determinations associated with this phase, although not derived directly from the chamber, place its construction between 4692–4269 cal BC/6642–6219 cal BP (95.4%; 4603–4338 cal BC, 68.2%)

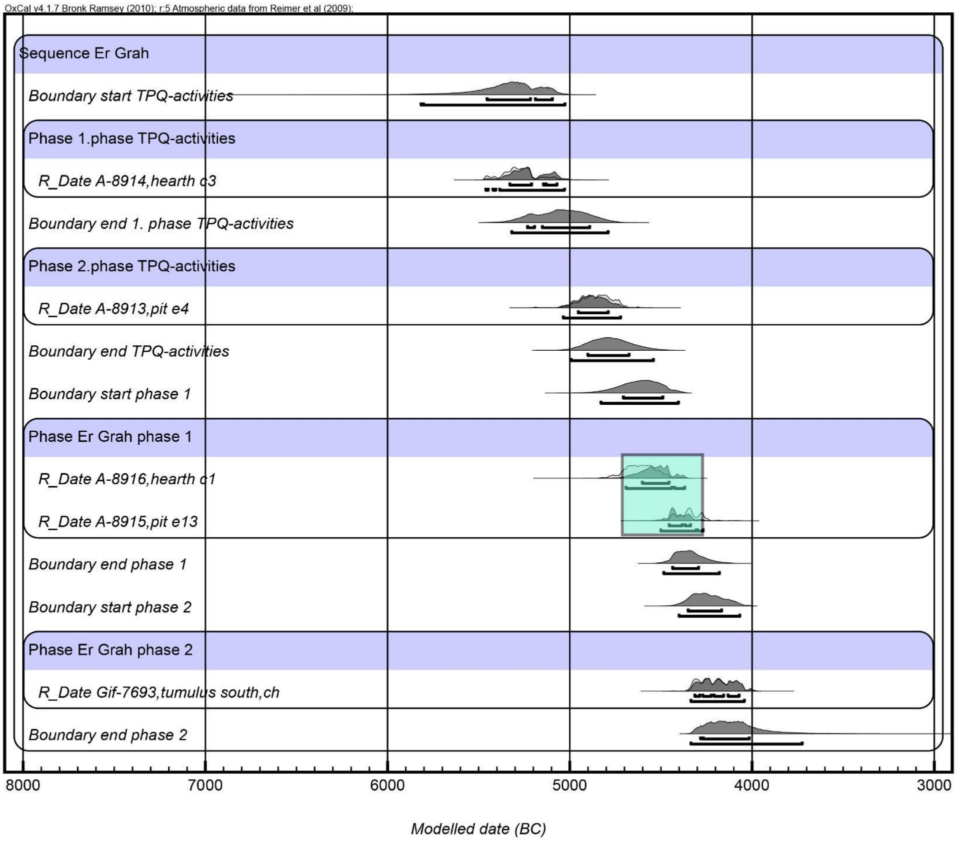

**Supplementary Fig. S2.9. Radiocarbon determinations from Er Grah.** Model agreement: $A_{model}$=79.2, $A_{overall}$=80.1.

#### Tumiac, Arzon

The construction sequence of the Tumiac (Butte de César) tumulus at Arzon is comparatively straightforward. The c. 50 m diameter monument consists of a stone-built burial chamber, with megalithic side stones and dry-walls, which was first covered by a stone cairn and subsequently enclosed within a large earthen mound[335–337] (Fouquet 1862; Galles 1878; Le Rouzic 1935). The chamber contained the remains of a single individual accompanied by an exceptional assemblage of prestige goods, including jade and fibrolite axes and 251 variscite (callaïs) beads and pendants, the richest such deposit known from the Morbihan with Encinansola in Southwestern Spain as the source for the variscite and the Monte Viso region for the jadeitite[142,151].

Four radiocarbon determinations are available, including one from the oak-plank floor of the chamber. Assuming that burial and monument construction were broadly contemporaneous, we constructed a Bayesian model comprising two phases. The first represents activity providing a *terminus post quem* for monument construction, based on sediment containing settlement material and including AA103575 (6135 ± 46 BP). The second phase represents construction of the wooden cist and mound. Charcoal determinations were modelled using the Charcoal Outlier model to account for potential inbuilt age. Because the chamber floor was constructed from large oak timbers and the dated tree rings may substantially predate timber felling and use, a relatively conservative inbuilt-age correction of 200 ± 100 years (normally distributed) was applied to AA105063; an even larger offset cannot be excluded.

The model shows good agreement (*A*model = 84.7; *A*overall = 86.3). The TPQ determination AA103575 is modelled to 5211–4846 cal BC/7161–6796 cal BP (95.4% probability; 5196–

4976 cal BC at 68.3%), with the end of this phase estimated at 5183–4551 cal BC/7133–6501 cal BP (95.4%; 5093–4766 cal BC at 68.3%). The beginning of the subsequent wooden-cist and mound-construction phase is estimated at 4899–4362 cal BC/6849–6312 cal BP (95.4%; 4685–4410 cal BC at 68.3%). After correction for potential inbuilt age, AA105063 is modelled to 4725–4336 cal BC/6675–6286 cal BP (95.4%; 4584–4385 cal BC at 68.3%) and thus to the whole range of the Ancient Castellic phase. The two further determinations associated with this construction phase are modelled to 4526–4307 cal BC/6476–6257 cal BP (AA10357) and 4490–4273 cal BC/6440–6223 cal BP (AA105064) at 95.4% probability.

These latter determinations may provide a closer indication of the timing of grave construction. Tumiac, together with its exceptionally rich greenstone deposits, may therefore belong to a later phase of the Carnac greenstone florescence. Tumiac is likewise one of the earliest full megalithic graves in the region.

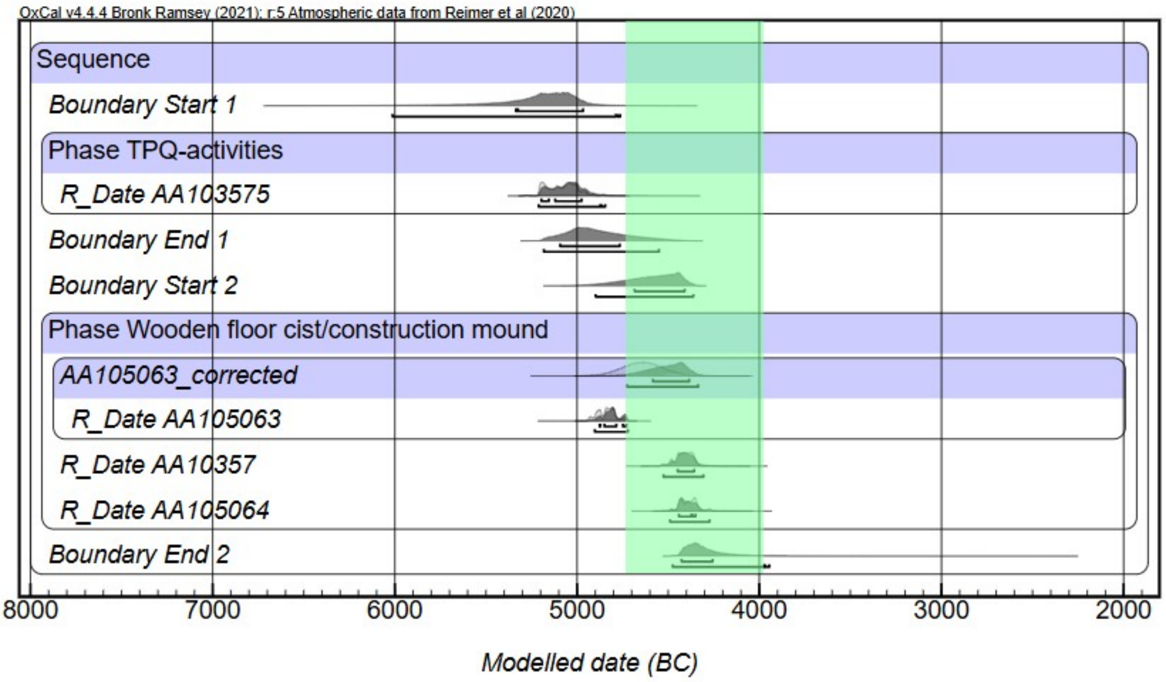

**Supplementary Fig. S2.10. Radiocarbon determinations from Tumiac.** Model agreement: $A_{model}$=84.7, $A_{overall}$=86.3.

| Name<br>Show all<br>Show structure | Unmodelled (BC/AD) | | | | | | Modelled (BC/AD) | | | | | | Indices<br>$A_{model}$=84.7<br>$A_{overall}$=86.3 | | | | | Select<br>All<br>Visible | Page break |
|---|---|---|---|---|---|---|---|---|---|---|---|---|---|---|---|---|---|---|---|
| | from | to | % | from | to | % | from | to | % | from | to | % | $A_{comb}$ | A | L | P | C | | |
| ▼ Outlier_Model Charcoal | | | | | | | -18 | 3 | 68.3 | -119 | 3 | 95.4 | | | | | 100 | 17 | |
| Exp(1,-10,0) | -1.24 | -0.05 | 68.3 | -3.19 | -0.05 | 95.4 | | | | | | | | | | | 99.8 | 2 | |
| U(0,3) | 2.21177e-17 | 3 | 68.3 | 2.21177e-17 | 3 | 95.4 | 3.59955e-17 | 1.317 | 68.3 | 3.59955e-17 | 2.145 | 95.4 | | 100 | | | 99.2 | 3 | |
| ▼ Sequence | | | | | | | | | | | | | | | | | | 4 | |
| Boundary Start 1 | | | | | | | -5336 | -4968 | 68.3 | -6013 | -4764 | 95.4 | | | | | 98 | 5 | |
| ▼ Phase TPQ-activities | | | | | | | | | | | | | | | | | | 6 | |
| R_Date AA103575 | -5207 | -4998 | 68.3 | -5212 | -4949 | 95.4 | -5196 | -4976 | 68.3 | -5211 | -4846 | 95.4 | | 98.3 | | | 99.8 | 7 | |
| Boundary End 1 | | | | | | | -5093 | -4766 | 68.3 | -5183 | -4551 | 95.4 | | | | | 99.6 | 8 | |
| Boundary Start 2 | | | | | | | -4685 | -4410 | 68.3 | -4899 | -4362 | 95.4 | | | | | 99.4 | 9 | |
| ▼ Phase Wooden floor cist/construction mound | | | | | | | | | | | | | | | | | | 10 | |
| ▶ AA105063_corrected | -4755 | -4526 | 68.3 | -4866 | -4416 | 95.4 | -4584 | -4385 | 68.3 | -4725 | -4336 | 95.4 | | | | | 99.6 | 11 | |
| R_Date AA10357 | -4449 | -4361 | 68.3 | -4533 | -4342 | 95.4 | -4449 | -4357 | 68.3 | -4526 | -4307 | 95.4 | | 100.6 | | | 99.7 | 14 | |
| R_Date AA105064 | -4444 | -4343 | 68.3 | -4489 | -4266 | 95.4 | -4442 | -4350 | 68.3 | -4490 | -4273 | 95.4 | | 99.7 | | | 99.7 | 15 | |
| Boundary End 2 | | | | | | | -4427 | -4256 | 68.3 | -4475 | -3946 | 95.4 | | | | | 96.7 | 16 | |

**Supplementary Fig. S2.11. Results for the calibration of the radiocarbon determinations from Tumiac.**

##### 2.11.5.4 Further Variscite contexts with radiocarbon dates from Northwest France

More early variscite contexts are documented in northwest France, although some are chronologically insecure. At Table des Marchands, a variscite bead was recovered from the pre-monument layer. A charcoal determination from the foundation pit of the backstone dates to 4933–4714 cal BC/6883–6664 cal BP (95.4%; Ly-13083, 5930 ± 45 BP), but cannot be directly associated with the bead. A second determination from hearth F3, contemporary with the later cairn, dates to 3971–3667 cal BC/5921–5617 cal BP (95.4%; Gif-7555, 5040 ± 70 BP). At Kercado, 156 variscite beads were recovered, but the only determination derives from unidentified charcoal and has a very large standard deviation (SA-95, 5840 ± 300 BP; 5464–4055 cal BC, 95.4%). The pre-monument layer at Dissignac also contained a variscite bead, but three charcoal determinations span 6230–3973 cal BC/8180–5923 cal BP (95.4%) and cannot securely date its deposition [156,338,339].

A reliable contemporary context outside Brittany is provided by the Replats cist at Aime in Rhône-Alpes, which contained one variscite bead. Two determinations on human bone date the burial to 4585–4343 cal BC/6535–6293 cal BP (95.4%; 4499–4368 cal BC/6449–6318 cal BP, 68.2%)[340] .

After c. 6300 BP, variscite ornaments occur more frequently in passage graves and other collective burial monuments. At Petit Mont in Arzon, 11 charcoal determinations constrain a complex sequence of successive tumuli and passage graves[156,341–343]. A determination associated with the initial long mound dates to 4515–4333 cal BC/465–6283 cal BP (95.4%; Gif-6844, 5650 ± 70 BP), although it may derive from the underlying soil, itself dated to 4679–4387 cal BC/6629–6337 cal BP (95.4%; Gif-6846, 5600 ± 70 BP). Grave II is modelled to 3944–3798 cal BC/5894–5748 cal BP (95.4%), passage IIIB to 3882–3777 cal BC/5832–5727 cal BP (95.4%), and activity at the façade of the final cairn to 3808–3536 cal BC/5758–5486 cal BP (95.4%). Seven variscite beads were recovered from chamber IIIA and the later excavations, while a pendant was found behind chamber IIIA in association with the façade of cairn I. The model has acceptable agreement (Amodel = 74.4; Aoverall = 77.8), but the exact association of the ornaments with the earliest mound remains uncertain.

At Île Carn, three variscite beads came from a mixed burial layer with secondary depositions in the northern passage grave[344] (pg. 42 in ref [341], pg. 205–209 in ref[345]). Of four available determinations, three have standard deviations of ±150–250 years. A charcoal sample from the central chamber dates to 4313–3811 cal BC/6263–5761 cal BP (95.4%; Gif-1968, 5230 ± 75 BP) (pg. 431 in ref [346]), but cannot directly date the variscite deposition.

The chronology of variscite contexts at Bougon is uncertain because several chambers functioned as ossuaries, preventing direct association between the dated human remains and the variscite beads[1,134,347,348]. The E1 determination has a large error (Ly-966, 5800 ± 230 BP). Dates associated with chamber F2 span 4224–3188 cal BC/6174–5138 cal BP (95.4%), while a nearby quarry dates to 4362–4052 cal BC/6312–6002 cal BP (95.4%) and may provide a terminus for its construction.

Later variscite contexts occur in the collective gallery graves of northern France. Tomb 1 at Porte-Joie contained one variscite bead together with stone and bone beads, axe pendants, gold plates and arrowheads[349,350] (pg. 145 in ref [351]). Four human-bone determinations indicate use from 3331–2915 cal BC (5281–4865 cal BP, 95.4%)) to 2111–1092 cal BC (4061–3042 cal BP, 95.4%), although the variscite bead cannot be assigned to a specific burial phase.

At La Butte Saint-Cyr, six variscite beads were recovered from a gallery grave containing at least 109 individuals[349]. Four human-bone determinations indicate use from 3498–3016 cal BC (5448–4966 cal BP, 95.4%) to 2881–2505 cal BC (4831–4455 cal BP, 95.4%). Given the large number of burials, however, these dates cannot be directly linked to the deposition of the variscite ornaments.

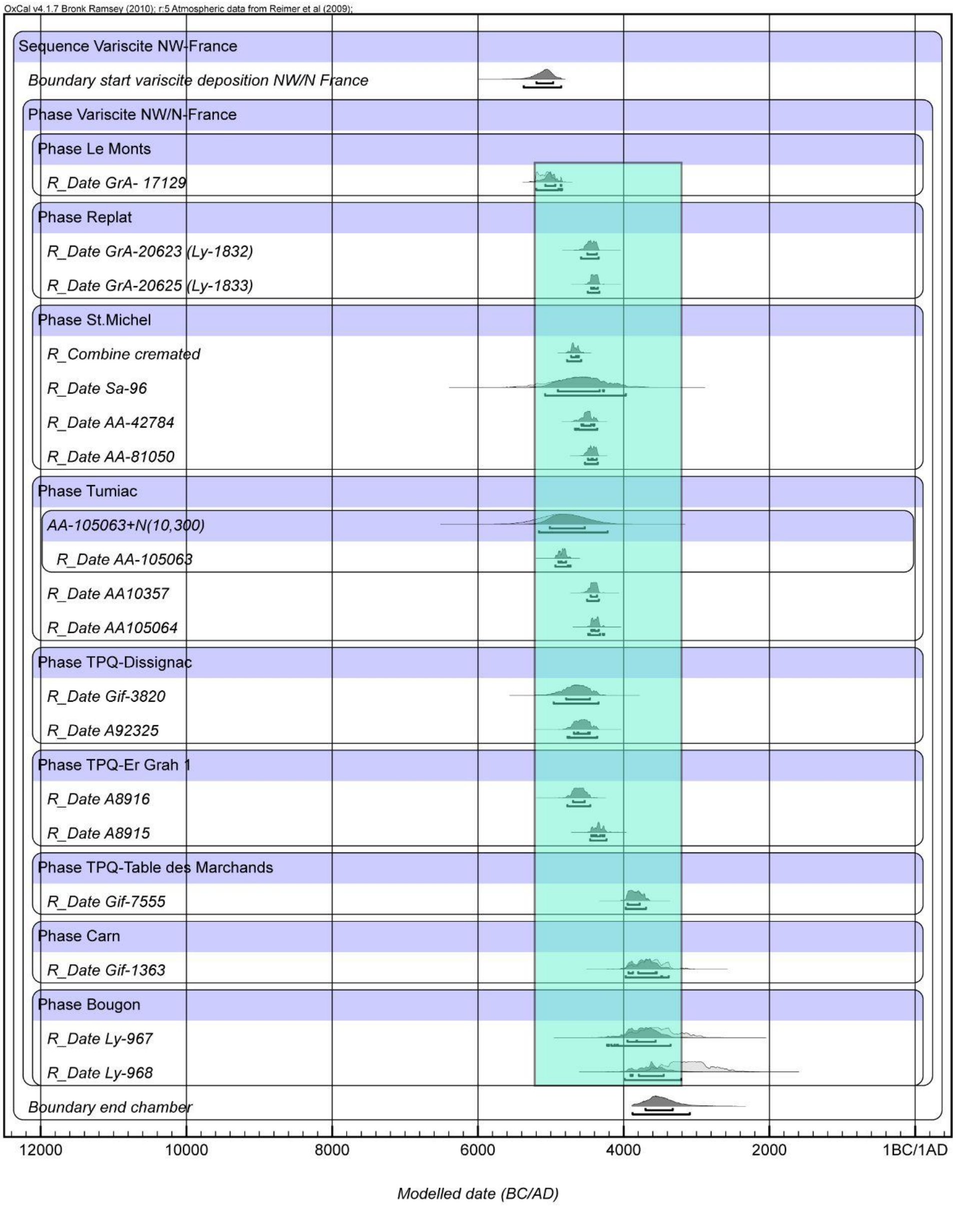

**Supplementary Fig. S2.12. Probability of all available radiocarbon dates out of reliable variscite context in Northern France**. Model agreement: $A_{model}$=86, $A_{overall}$=81.4.

##### 2.11.5.5 Mediterranean contexts with radiocarbon dates and variscite beads

In the time interval of 6750-6050 BP (4741-4048 cal BC, 95.4%; 4599-4101 cal BC, 68.2%) large variscite deposits have been recorded in natural caves in Catalonia. This variscite is obtained from the Can Tintorer/Gavà mines.

Can Tintorer/ Gavá is likewise in discussion as a source for the raw material of the very early variscite finds from funeral contexts in natural caves or collective burials from the Aragón and Navarra regions of Northeastern Spain (5346-4802 cal BC/7296–6752 cal BP, 95.4%; 5269-4871 cal BC/7219–6821 cal BP, 68.2%)[23,151]. These earliest variscite horizons are not reflected by the sequence of the excavations at the Can Tintorer/ Gavá mines, which starts later.

We modelled 21 radiocarbon dates available from the mine and associated funerals in the mine shafts[322,323]. The directly dated mining activity at Gavà begins in 4055–3976 cal BC/6005–5926 cal BP (68.3% probability; 4156–3946 cal BC/6106–5896 cal BP, 95.4%) and ends in 3619–3525 cal BC/5569–5475 cal BP (68.3%; 3632–3422 cal BC/5582–5372 cal BP, 95.4%). Its estimated duration is 351–446 years (68.3%; 320–546 years, 95.4%). Mines 5, 6, 16 and 70 appear to have been exploited from the beginning of this documented phase, whereas Mines 7 and 90 belong mainly to its later part. The funerary evidence in the mines begins considerably earlier, in 4630–4341 cal BC/6580–6291 cal BP (68.3%; 5246–4274 cal

BC/7196–6224 cal BP, 95.4%). This supports variscite extraction before the earliest directly dated mining contexts.

Variscite and turquoise beads are further deposited in the middle Neolithic in Catalonia within funeral contexts as in hypogea, cists and megalithic graves. This is the time of the peak of the documented mining activities at the Can Tintorer/Gavá mines between 6050-5450 BP (4041-3439 cal BC; 95.4%; 3998-3439, 69.2%). For Southern France the radiocarbon results indicate the circulation of variscite artefacts in the Ancient Chassean horizon, from 6300-6050 BP (4309-4053 cal BC 95.4%; 4271-4067 cal BC, 68.2%) onwards. Twenty-one radiocarbon dates were considered for the variscite deposition in southern France. The results indicate that variscite ornaments circulated from the Early Chassey period, beginning in 4309–4053 cal BC/6259–6003 cal BP (95.4% probability; 4271–4067 cal BC, 68.2%). The earliest evidence comes primarily from funerary contexts, including the child burial in Grave 33 at Béziers, Le Crès, and contemporary graves at Cugnaux.

Variscite continued to be deposited in graves, caves and settlements during the Chassey, Montbolo and Final Neolithic periods[161,352]. A further phase of use is documented in Early Chalcolithic caves and megalithic graves in 3444–3095 cal BC/5394–5045 cal BP (95.4%; 3372–3161 cal BC/5322–5111 cal BP, 68.2%). Variscite beads also occur in closed Bell Beaker contexts dating typologically to approximately 2500–2300 cal BC/4500–4300 cal BP, although these assemblages currently lack radiocarbon dates. Overall, the evidence demonstrates the long-term circulation and changing deposition of variscite in southern France from the late fifth to the third millennium BC (6500-4000 BP).

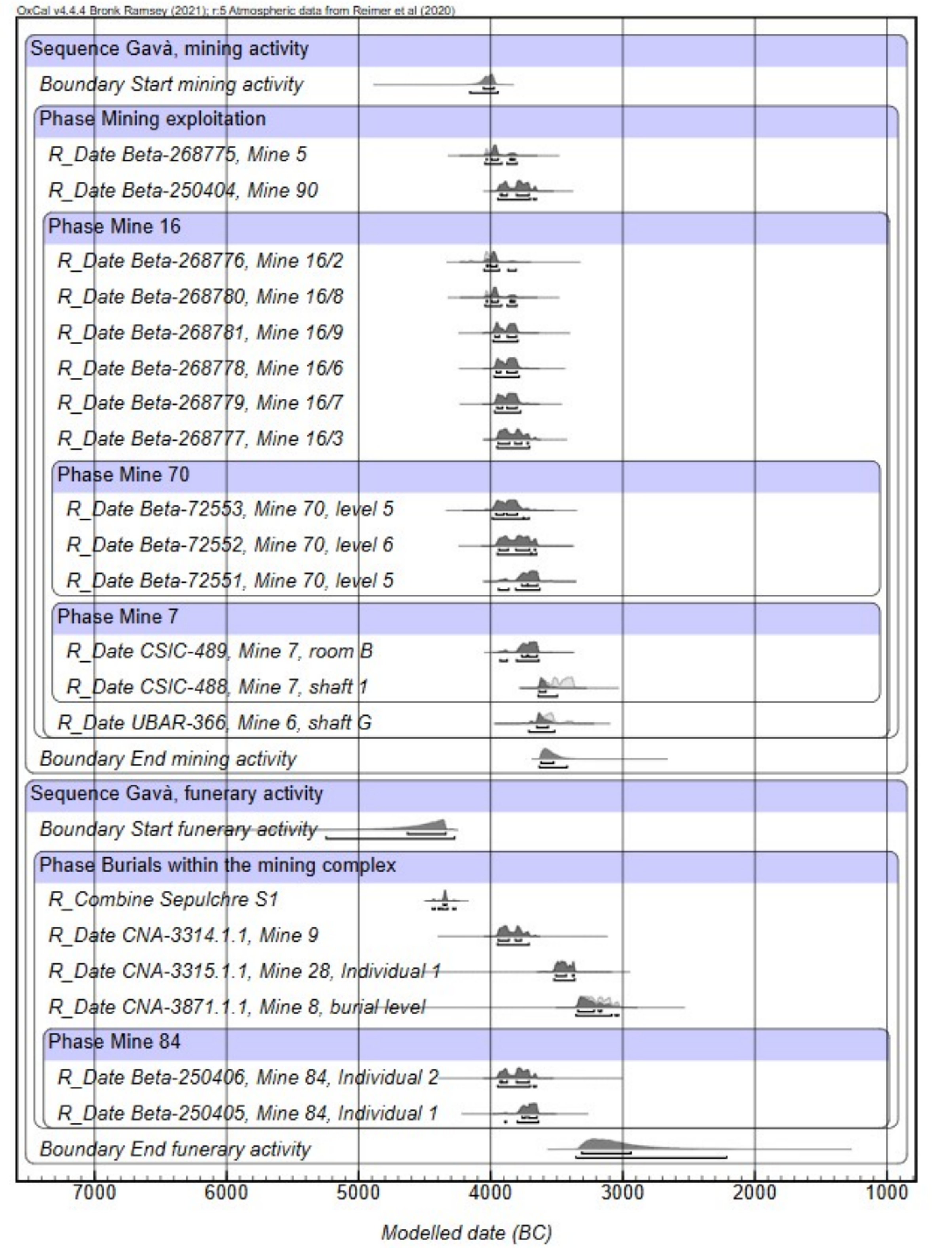

**Supplementary Fig. S2.13. Modelled probability distributions for mining and funerary activity at Can Tintorer, Gavà.** Charcoal determinations were analysed using the Charcoal

Outlier Model to account for possible inbuilt age. The model shows good overall agreement (Amodel = 102.2; Aoverall = 101.3).

```
Plot()
 {
  Outlier_Model(
       "Charcoal",
       Exp(1,-10,0),
       U(0,3),
       "t"
  );
  Outlier_Model(
       "General",
       T(5),
       U(0,4),
       "t"
  );
  Sequence("Gavà, mining activity")
  {
   Boundary("Start mining activity");
   Phase("Mining exploitation")
   {
       R_Date("Beta-268775, Mine 5",5160,40)
       {
       Outlier("Charcoal",0.05);
       };
       R_Date("Beta-250404, Mine 90",5010,40)
       {
       Outlier("Charcoal",0.05);
       };
       Phase("Mine 16")
       {
       R_Date("Beta-268776, Mine 16/2",5190,40)
       {
       Outlier("Charcoal",0.05);
       };
       R_Date("Beta-268780, Mine 16/8",5160,40)
       {
       Outlier("Charcoal",0.05);
       };
       R_Date("Beta-268781, Mine 16/9",5110,40)
       {
       Outlier("Charcoal",0.05);
       };
       R_Date("Beta-268778, Mine 16/6",5090,40)
       {
       Outlier("Charcoal",0.05);
       };
       R_Date("Beta-268779, Mine 16/7",5080,40)
       {
       Outlier("Charcoal",0.05);
       };
       R_Date("Beta-268777, Mine 16/3",5030,40)
       {
       Outlier("Charcoal",0.05);
       };
       Phase("Mine 70")
       {
       R_Date("Beta-72553, Mine 70, level 5",5090,60)
       {
       Outlier("Charcoal",0.05);
       };
       R_Date("Beta-72552, Mine 70, level 6",5010,60)
       {
       Outlier("Charcoal",0.05);
       };
       R_Date("Beta-72551, Mine 70, level 5",4930,60)
       {
       Outlier("Charcoal",0.05);
```

```
      };
      };
      Phase("Mine 7")
      {
      R_Date("CSIC-489, Mine 7, room B",4940,50)
      {
      Outlier("Charcoal",0.05);
      };
      R_Date("CSIC-488, Mine 7, shaft 1",4710,50)
      {
      Outlier("Charcoal",0.05);
      };
      };
      R_Date("UBAR-366, Mine 6, shaft G",4810,60)
      {
      Outlier("Charcoal",0.05);
      };
      };
      Span("Duration mining activity");
  };
  Boundary("End mining activity");
 };
 Sequence("Gavà, funerary activity")
 {
  Boundary("Start funerary activity");
  Phase("Burials within the mining complex")
  {
      R_Combine("Sepulchre S1")
      {
      R_Date("CNA-4566.1.1",5510,30);
      R_Date("CNA-3312.2.1",5499,31);
      };
      R_Date("CNA-3314.1.1, Mine 9",5029,33)
      {
      Outlier("General",0.05);
      };
      R_Date("CNA-3315.1.1, Mine 28, Individual 1",4658,33)
      {
      Outlier("General",0.05);
      };
      R_Date("CNA-3871.1.1, Mine 8, burial level",4478,31)
      {
      Outlier("General",0.05);
      };
      Phase("Mine 84")
      {
      R_Date("Beta-250406, Mine 84, Individual 2",5010,40)
      {
      Outlier("General",0.05);
      };
      R_Date("Beta-250405, Mine 84, Individual 1",4940,40)
      {
      Outlier("General",0.05);
      };
      };
      Span("Duration funerary activity");
  };
  Boundary("End funerary activity");
 };
};
```

**Supplementary Fig. S2.14. Code of the Bayesian model of the investigated exploitation phase at Can Tintorer, Gava based on radiocarbon dates from the mining shafts and**

**funerals in the mining shaft using the OxCal Charcoal Outlier Model to account for potential inbuilt age effects and intrusive material..**

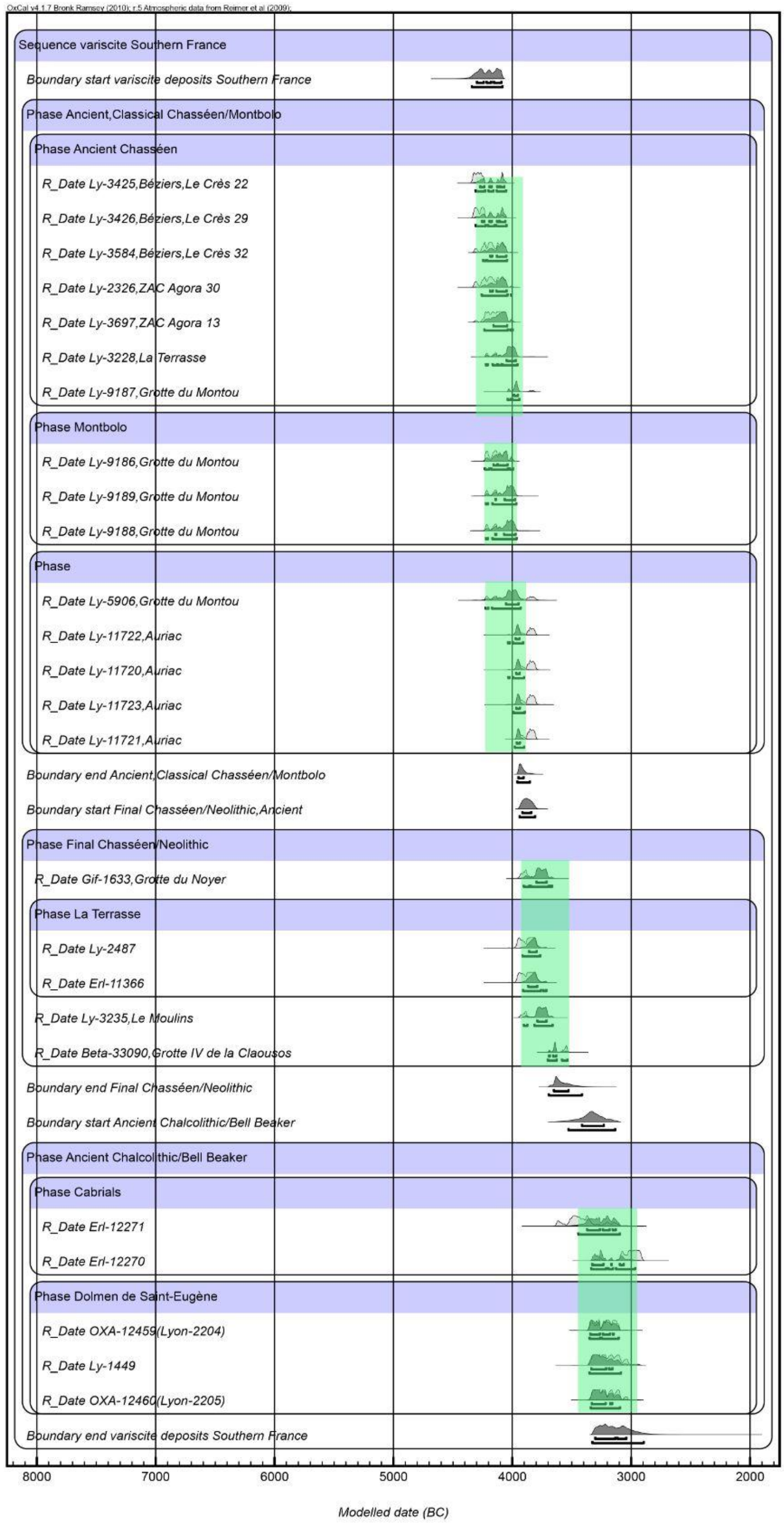

OxCal v4.1.7 Bronk Ramsey (2010); r:5 Atmospheric data from Reimer et al (2009);
Sequence variscite Southern France
Boundary start variscite deposits Southern France
Phase Ancient,Classical Chasséen/Montbolo
Phase Ancient Chasséen
R_Date Ly-3425,Béziers,Le Crès 22
R_Date Ly-3426,Béziers,Le Crès 29
R_Date Ly-3584,Béziers,Le Crès 32
R_Date Ly-2326,ZAC Agora 30
R_Date Ly-3697,ZAC Agora 13
R_Date Ly-3228,La Terrasse
R_Date Ly-9187,Grotte du Montou
Phase Montbolo
R_Date Ly-9186,Grotte du Montou
R_Date Ly-9189,Grotte du Montou
R_Date Ly-9188,Grotte du Montou
Phase
R_Date Ly-5906,Grotte du Montou
R_Date Ly-11722,Auriac
R_Date Ly-11720,Auriac
R_Date Ly-11723,Auriac
R_Date Ly-11721,Auriac
Boundary end Ancient,Classical Chasséen/Montbolo
Boundary start Final Chasséen/Neolithic,Ancient
Phase Final Chasséen/Neolithic
R_Date Gif-1633,Grotte du Noyer
Phase La Terrasse
R_Date Ly-2487
R_Date Erl-11366
R_Date Ly-3235,Le Moulins
R_Date Beta-33090,Grotte IV de la Claousos
Boundary end Final Chasséen/Neolithic
Boundary start Ancient Chalcolithic/Bell Beaker
Phase Ancient Chalcolithic/Bell Beaker
Phase Cabrials
R_Date Erl-12271
R_Date Erl-12270
Phase Dolmen de Saint-Eugène
R_Date OXA-12459(Lyon-2204)
R_Date Ly-1449
R_Date OXA-12460(Lyon-2205)
Boundary end variscite deposits Southern France
8000
7000
6000
5000
4000
3000
2000
Modelled date (BC)

2750

**Supplementary Fig. S2.15. Probability of all available radiocarbon dates out of reliable variscite context in Southern France**. Model agreement: $A_{model}$=68.8, $A_{overall}$=65.4.

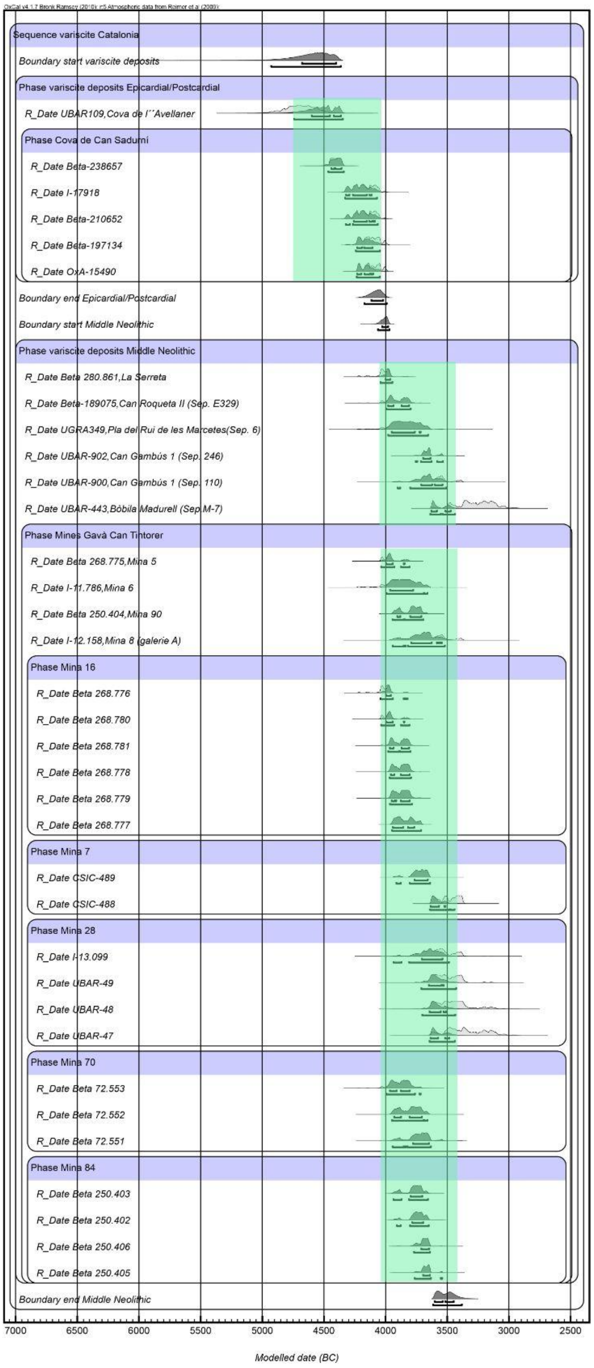

OxCal v4.1.7 Bronk Ramsey (2010); r:5 Atmospheric data from Reimer et al (2009);
Sequence variscite Catalonia
Boundary start variscite deposits
Phase variscite deposits Epicardial/Postcardial
R_Date UBAR109,Cova de l´´Avellaner
Phase Cova de Can Sadurní
R_Date Beta-238657
R_Date I-17918
R_Date Beta-210652
R_Date Beta-197134
R_Date OxA-15490
Boundary end Epicardial/Postcardial
Boundary start Middle Neolithic
Phase variscite deposits Middle Neolithic
R_Date Beta 280.861,La Serreta
R_Date Beta-189075,Can Roqueta II (Sep. E329)
R_Date UGRA349,Pla del Rui de les Marcetes(Sep. 6)
R_Date UBAR-902,Can Gambús 1 (Sep. 246)
R_Date UBAR-900,Can Gambús 1 (Sep. 110)
R_Date UBAR-443,Bòbila Madurell (Sep.M-7)
Phase Mines Gavà Can Tintorer
R_Date Beta 268.775,Mina 5
R_Date I-11.786,Mina 6
R_Date Beta 250.404,Mina 90
R_Date I-12.158,Mina 8 (galerie A)
Phase Mina 16
R_Date Beta 268.776
R_Date Beta 268.780
R_Date Beta 268.781
R_Date Beta 268.778
R_Date Beta 268.779
R_Date Beta 268.777
Phase Mina 7
R_Date CSIC-489
R_Date CSIC-488
Phase Mina 28
R_Date I-13.099
R_Date UBAR-49
R_Date UBAR-48
R_Date UBAR-47
Phase Mina 70
R_Date Beta 72.553
R_Date Beta 72.552
R_Date Beta 72.551
Phase Mina 84
R_Date Beta 250.403
R_Date Beta 250.402
R_Date Beta 250.406
R_Date Beta 250.405
Boundary end Middle Neolithic
7000
6500
6000
5500
5000
4500
4000
3500
3000
2500
Modelled date (BC)

2754

**Supplementary Fig. S2.16. Probability of all available radiocarbon dates out of reliable variscite context in Catalonia**. Model agreement: $A_{model}$=86.9, $A_{overall}$=83.6.

##### 2.11.5.6 West Iberian contexts with radiocarbon dates and variscite beads

We considered 37 Portuguese sites with variscite ornaments, for which more than 100 radiocarbon determinations are available[23]. However, only contexts of moderate or high reliability were included in the analysis, reducing the dataset to 41 dates[23].

Possible Early Neolithic evidence comes from Gruta do Caldeirão in Estremadura, where three variscite beads were recovered from a cave used for both settlement and burial. Three radiocarbon dates on two animal bones and one human bone indicate a time interval of 7000-6550 BP (4989 to 4543 cal BC 95.4%; 4846 to 4622 cal BC, 68.2%). However, the cave was occupied from the Early Neolithic to the Visigothic period, and recent reassessment suggests that the beads may belong to its Middle or Late Neolithic phases. Their Early Neolithic attribution must therefore remain provisional[353]. Variscite beads from possible proto-megalithic cists in the Monchique Mountains have been assigned typologically to the fifth millennium cal BC (7000-6000 BP), but this chronology remains unsupported by radiocarbon dates[354].

The earliest more reliable associations between variscite and megalithic graves come from the passage graves of Areita 1 and Carapito 1 in northern Portugal[355,356]. Two dates on human bone are available from the chamber of Areita 1. The earlier result dates to approximately 5960 BP (3976 to 3795 cal BC, 68.3%; 4037 to 3712 cal BC, 95.4%), while the second falls around 5660 BP (3695 to 3537 cal BC, 68.2%; 3756 to 3517 cal BC, 95.4%). The gap of approximately 300 years between them, together with the isolated position of the earlier date within the wider western Iberian dataset, suggests that it may derive from a secondary burial.

It should therefore be treated as a *terminus post quem* rather than as a construction date. Only GrN-18497, 4840 ± 60 BP (3695 to 3537 cal BC, 68.2%; 3756 to 3517 cal BC, 95.4%), can be associated more securely with the deposition of variscite, although contamination and reservoir effects cannot be entirely excluded. A closely contemporary result from Carapito 1, GrN-5110, 4850 ± 40 BP (3693 to 3540 cal BC, 68.2%; 3707 to 3530 cal BC, 95.4%), supports this chronology.

The Middle Neolithic evidence is followed by a hiatus of at least 800 years. Variscite use reached its peak during the Chalcolithic, when 35 of the 41 accepted dates fall within 4900-4220 BP (2860 to 2220 cal BC, 95.4%; 2834 to 2145 cal BC, 68.2%). Variscite ornaments occur in the hypogea of São Pedro do Estoril, Quinta do Anjo and São Paulo, the tholoi of Agualva and Perdigões, and the megalithic grave of Casainhos. Most assemblages contain between one and 37 beads, while Anta Grande do Zambujeiro produced an exceptional group of 277.

Anta Grande do Zambujeiro has a long burial history extending across the fourth and third millennia cal BC (6000-4000 BP). Its only radiocarbon determination, Beta-243693, 3910 ± 40 BP, was excluded because it cannot be related directly to the deposition of the variscite. However, the association with more than 150 idol plaques supports an attribution to the interval 4860-4220 BP (2860 to 2220 cal BC, 95.4%; 2834 to 2145 cal BC, 68.2%)[197,357,358]. Large quantities of variscite beads are also known from Late Neolithic and Chalcolithic settlements across Portugal, including Pragança, Leceia, Moita da Ladra and Buraco da Pala[359–361].

Provenance analyses identify Palazuelo and Pico Centeno as the sources of all characterised Late Neolithic and Chalcolithic variscite from Portugal[362,363]. With the exceptions of Anta

Grande do Zambujeiro and Vila Nova de São Pedro, individual sites contain material from only one source. Palazuelo de las Cuevas, approximately 400 km to the north-east, is much more strongly represented than the closer source of Pico Centeno, located only 50 to 100 km to the east. This pattern indicates structured exchange networks rather than simple reliance on the nearest source.

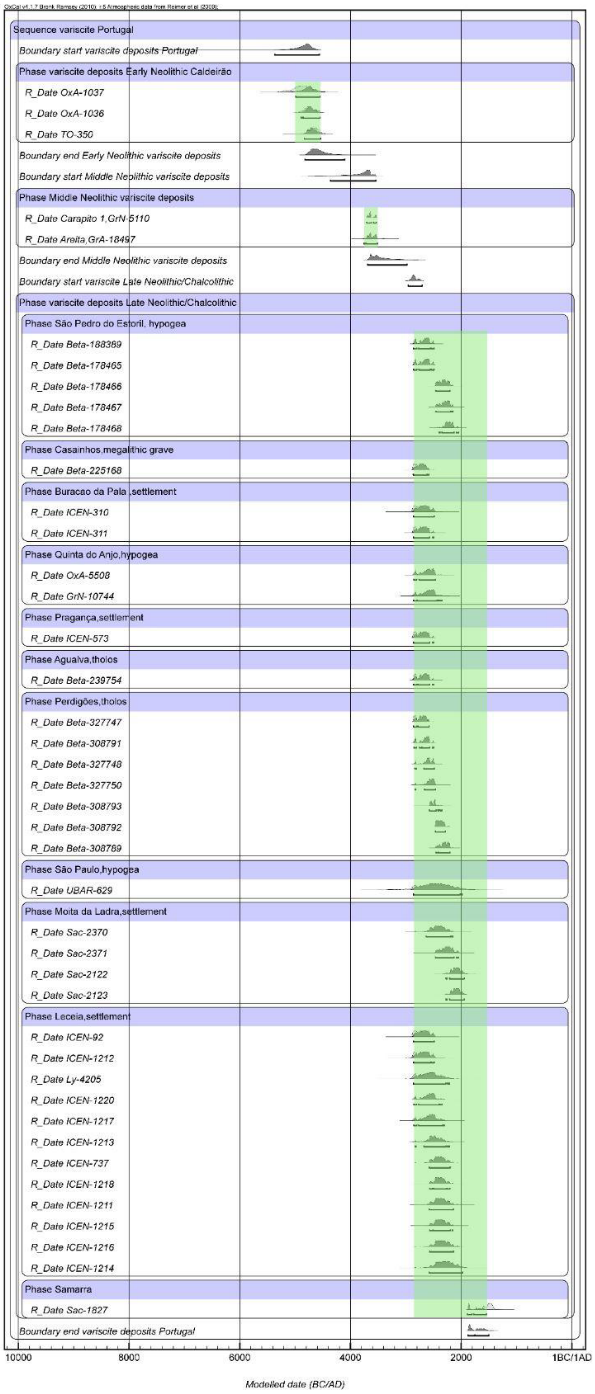
Sequence variscite Portugal
Boundary start variscite deposits Portugal
Phase variscite deposits Early Neolithic Caldeirão
R_Date OxA-1037
R_Date OxA-1036
R_Date TO-350
Boundary end Early Neolithic variscite deposits
Boundary start Middle Neolithic variscite deposits
Phase Middle Neolithic variscite deposits
R_Date Carapito 1,GrN-5110
R_Date Areita,GrA-18497
Boundary end Middle Neolithic variscite deposits
Boundary start variscite Late Neolithic/Chalcolithic
Phase variscite deposits Late Neolithic/Chalcolithic
Phase São Pedro do Estoril, hypogea
R_Date Beta-188369
R_Date Beta-178465
R_Date Beta-178466
R_Date Beta-178467
R_Date Beta-178468
Phase Casainhos,megalithic grave
R_Date Beta-225168
Phase Buracao da Pala ,settlement
R_Date ICEN-310
R_Date ICEN-311
Phase Quinta do Anjo,hypogea
R_Date OxA-5508
R_Date GrN-10744
Phase Pragança,settlement
R_Date ICEN-573
Phase Agualva,tholos
R_Date Beta-239754
Phase Perdigões,tholos
R_Date Beta-327747
R_Date Beta-308791
R_Date Beta-327748
R_Date Beta-327750
R_Date Beta-308793
R_Date Beta-308792
R_Date Beta-308789
Phase São Paulo,hypogea
R_Date UBAR-629
Phase Moita da Ladra,settlement
R_Date Sac-2370
R_Date Sac-2371
R_Date Sac-2122
R_Date Sac-2123
Phase Leceia,settlement
R_Date ICEN-92
R_Date ICEN-1212
R_Date Ly-4205
R_Date ICEN-1220
R_Date ICEN-1217
R_Date ICEN-1213
R_Date ICEN-737
R_Date ICEN-1218
R_Date ICEN-1211
R_Date ICEN-1215
R_Date ICEN-1216
R_Date ICEN-1214
Phase Samarra
R_Date Sac-1827
Boundary end variscite deposits Portugal
10000
8000
6000
4000
2000
1BC/1AD
Modelled date (BC/AD)

2807

**Supplementary Fig. S2.17. Probability of all available radiocarbon dates out of reliable variscite context in Portugal.** Model agreement: $A_{model}$=79.9, $A_{overall}$=79.8.

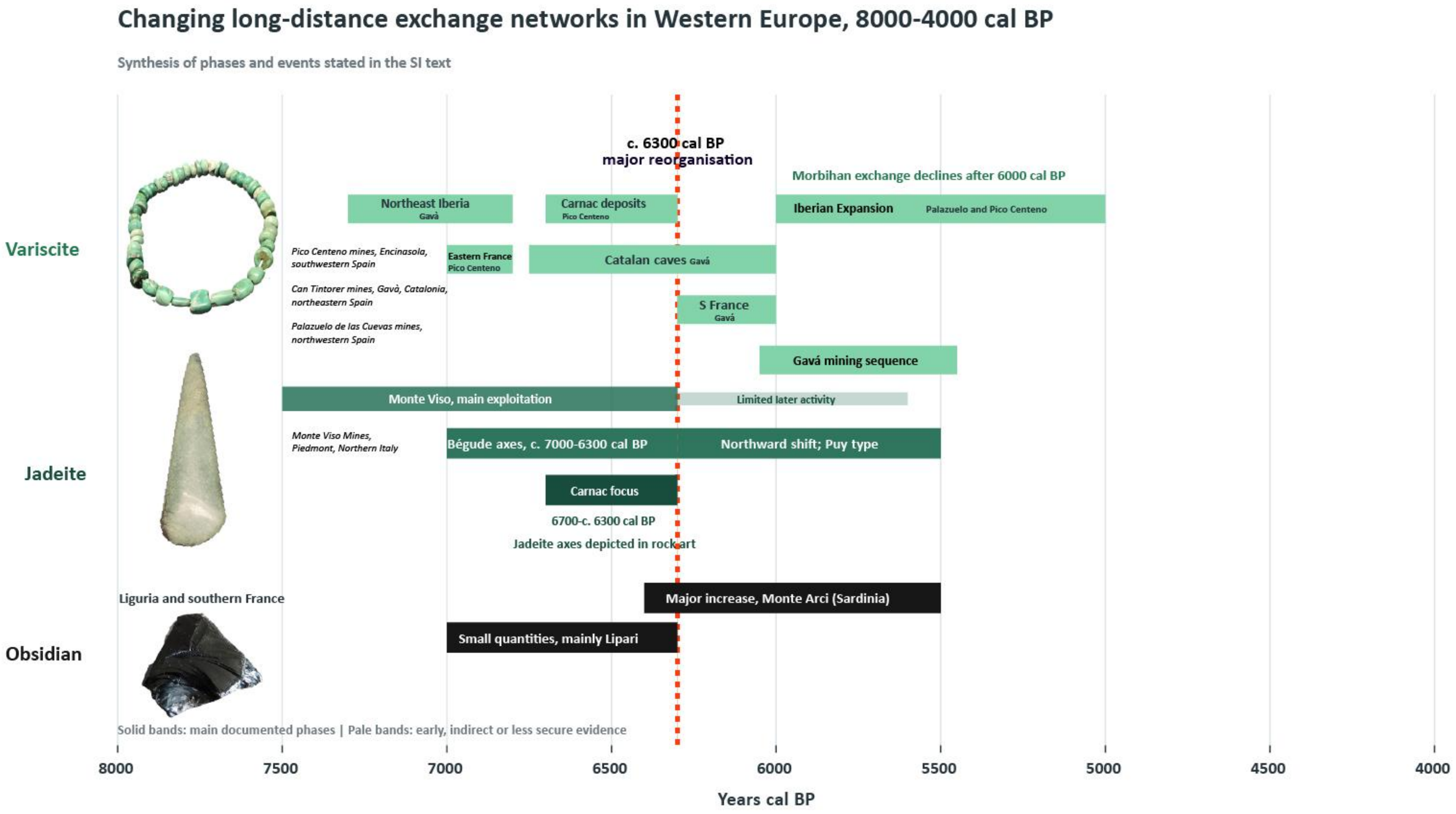

**Supplementary Fig. S2.18. Synthesis: trading networks and timeline**

### Supplementary Note S3. The newly sampled human remains

*Rita Peyroteo Stjerna, Maria Grazia Melis, Bettina Schulz Paulsson, Rich Potter, Vitale Sparacello, António Faustino Carvalho, Eduardo Porfirio, Catarina Costeira, Ana Maria Gama da Silva, Ana Bini, José Moita*

#### 3.1 Introduction

In this Supplementary Information, we provide detailed information on all newly sequenced individuals included in this study. Sampling efforts focused on regions that have remained underrepresented in ancient DNA research, either because of limited access to human remains or poor preservation conditions. One such region is Sardinia, where the availability of skeletal material for genetic analysis has been constrained by administrative factors. Another is Atlantic Iberia, where acidic soils and unfavourable depositional environments have generally resulted in poor bone preservation. As a consequence, prehistoric populations from these regions remain comparatively underrepresented in genomic datasets.

Despite these challenges, suitable human remains were identified in the Tagus estuary region of Portugal and from the coast in the Sintra region. We successfully obtained and sequenced samples from five megalithic tombs, one tholos, two burials of uncertain monument type, one cave burial context, and one Mesolithic cemetery. In total, 33 samples from Portugal are presented and discussed below.

For Sardinia, sampling focused on the hypogeum of San Benedetto. A total of 18 individuals were analysed, yielding 13 successful genetic datasets. These individuals substantially increase the currently available genomic data from prehistoric Sardinia and provide important new insights into population history in the central Mediterranean.

#### 3.2 The sampling strategy

##### 3.2.1 Sardinia

*Maria Grazia Melis, Vitale Sparacello, Ana Bini*

*San Benedetto/Iglesias*

The skeletal series from San Benedetto di Iglesias (Sardinia) included in this study consists of human remains from two collective hypogeal burials, or *Domus de Janas* (Tomba II and B). The Tomba II reportedly contained cranial and postcranial elements belonging to a MNI of 35 [364,365], while the Tomba B yielded a MNI of 5. Five published radiometric determinations place the material of tomb II within the first phase of the Ozieri Neolithic culture (ca. 4000-3500 BC/6000-5500 BP; [365–367]). The skeletal remains are currently housed in the deposits of the Museo Sardo di Antropologia ed Etnografia (MSAE), University of Cagliari.

Most of the assemblage consists of crania and calvaria, as well as several cranial fragments. The skeletal series has not been subject to a complete osteological reassessment since the first publication (ref [364]; cf. refs [365,368]), and new attempts to associate cranial fragments and loose teeth into dentitions are currently ongoing. Sampling for aDNA analysis did not undermine the ongoing osteological analysis, given that it did not significantly alter the morphology of the skeletal elements. The majority of crania and calvaria were numbered during the excavation, and some documentation is available on their positioning in the hypogeal chambers that constituted the tomb (e.g. ref [365]). For this reason, the survey for the aDNA sampling prioritised the numbered crania. However, to minimise the damage to the skeletal collection – and given the impossibility of transporting intact crania to the extraction facility – only five individuals represented by numbered crania were included in the sampling, i.e. the ones that presented a detached temporal bone (four right and one left elements) (Table 1).

Twelve additional temporal bones or fragments of petrous portions (all from the left side) that were not attributed to a numbered cranium were sampled, making sure that none of them was contralateral to elements from the numbered crania (Table 1). In addition to petrous portions of the temporal bone, two teeth (an upper left first premolar from individual 12 and an upper left second molar from individual 16) certainly associated to numbered crania were sampled (Table 3.1).

| Intact crania | | | | | |
|---|---|---|---|---|---|
| **Individual number** | **MSAE**<br><br>**Catalogue number** | **Right temporal** | **Left temporal** | **Temporal sampled** | **Tooth sampled** |
| 1 | 6230 | Separated | Attached | Right | |
| 2 | 6235 | Attached | Attached | | |
| 3 | 6231 | Attached | Attached | | |
| 4 | 6236 | Attached | | | |
| 5 | 6237 | Attached | Attached | | |
| 6 | 6234 | Attached | Attached | | |
| 7 | 6233 | Attached | | | |
| 12 | 6241 | Separated | Separated | Right | ULP1 |
| 13 | 7689 | Attached | | | |
| 14 | 6239 | Attached | | | |

| | | | | | |
|---|---|---|---|---|---|
| 15 | 6238 | | Attached | | |
| 16 | 6245 | Attached | Attached | | ULM2 |
| 18 | 6246 | Attached | Attached | Right | |
| 19 | 6240 | Attached | | Right | |
| 24 | 6248 | Attached | Attached | | |
| 25 | 6250 | Attached | Attached | | |
| 26 | 6249 | Attached | | | |
| 27 | 7641 | Attached | Attached | | |
| 28 | 6247 | Attached | Attached | | |
| B1 (Tomba B) | 6412 | Attached | Separated | Right | |
| B2 (Tomba B) | 6413 | Attached | Attached | | |
| Detached temporal bones | | | | | |
| LABEL | MSAE Catalogue number | Right temporal | Left temporal | **Temporal sampled** | **Tooth sampled** |
| A | | | Separated | Left | |
| D | | | Separated | Left | |
| 11 | | | Separated | Left | |

| | | | | |
|---|---|---|---|---|
| F | | | Separated | Left |
| M1 | | | Separated | Left |
| NONE | 6260 | | Separated | Left |
| NONE | | | Separated | Left |
| NONE | | | Separated | Left |
| NONE | | | Separated | Left |
| NONE | | | Separated | Left |
| NONE | | | Separated | Left |
| NONE | 6257 | | Separated | |
| B | | Separated | Separated | Left |
| FRGDN | 6298 | Separated | | |
| NONE | | Separated | | |
| NONE | 6270 | Separated | | |
| NONE | 6268 | Separated | | |
| NONE | | Separated | | |
| NONE | | Separated | | |

**Supplementary Table S3.1. List of crania with intact temporal bones and detached temporal elements from the skeletal series of San Benedetto di Iglesias.** All the remains derive from Tomba II, if not otherwise indicated. The elements that were sampled are indicated in the rightmost columns.

**References:** [364–368]

3.2.2 Portugal

*Rita Peyroteo Stjerna & Ana Maria Gama da Silva*

3.2.2.1 Cova da Onça shell midden and the Sintra region (Tholos of Praia das Maçãs, the Dolmen of Pedras da Granja, and the prehistoric burial site of Samarra)

Human skeletal remains were collected for ancient DNA (aDNA), radiocarbon, and isotope analysis from legacy collections curated at the Museu Geológico, Lisbon (MG) and the Museu Arqueológico de São Miguel de Odrinhas (MASMO), Sintra. R.P.S. conducted the individual selection from four distinct sites: the Cova da Onça shell midden (MG), the Tholos of Praia das Maçãs (MG, MASMO), the Dolmen of Pedras da Granja (MASMO), and the prehistoric burial site of Samarra (MASMO).

Original field documentation for these sites is scarce; typically, individuals lack unique excavation IDs or specific stratigraphic information. Consequently, our selection strategy focused on identifying repeated skeletal elements to prevent redundant sampling of the same individual while simultaneously maintaining the integrity of the osteological collections. We prioritised the sampling of the petrous portion of the temporal bone, followed by teeth as a secondary priority when the petrous portion was unavailable. In two specific instances (Praia das Maçãs, MG), bones of the feet were selected due to the nature of the available material. To avoid duplication within these now commingled collections, we preferentially selected the same element from a single anatomical side (e.g., right-side petrous portions only). Contralateral elements were bypassed to ensure non-redundancy, acknowledging that while some teeth and pedal elements may belong to the same individuals as the petrous samples.

###### 3.2.2.2 The Northern bank of the Tagus River in the Lisbon region

On the northern bank of the Tagus River in the Lisbon region lies one of the few areas in western Iberia with favourable conditions for the preservation of human bone. As a result, a small number of Neolithic burial sites have yielded human skeletal remains. Several of these remains were sampled by Rui Boaventura for radiocarbon dating[197]. With the kind permission of the Museu Geológico, we re-sampled the collection for ancient DNA analysis.

Late Neolithic funerary contexts in present-day Portugul are characterise as collective burials. When human bones were found, they were very disturbed, fragmented and commingled, with total or almost total absence of anatomical connection. Accordingly, sample strategy was adapted to this particular funerary context. All the selected burial site housed at Museu Geológico de Lisboa (MG), include no anatomical connection. So, the most representative side of petrous bone was selected, separately of adults and non-adults, to avoid duplication of sampling the same individual.

##### 3.2.3 The sampling documentation with 3-D modelling

*Rich Potter*

As part of the NEOSEA sampling protocol, skeletal material was documented, comprehensively recorded, and made accessible in its original condition prior to destructive analyses, including ancient DNA (aDNA), radiocarbon, and stable isotope analyses.

Projects using Structure from Motion (SfM) and scanning methods for the documentation of teeth and bones have previously been published: some have been based in dentistry[369,370], while others have been based within the archaeological sphere[371–373]. However, these have mostly focused on documenting and presenting bone material as reference material rather than digital preservation prior to sampling.

Within this project eleven of the teeth and bone fragments that were to be sampled were successfully documented using a standard Structure from Motion (SfM) technique which has seen wide applications in archaeology[374–379]. As the teeth and bone fragments were small objects with very small details that needed to be captured, our photographs were taken using a 100 mm macro lens. Bone fragments were propped up with foam blocks, and the teeth were either placed on a foam block or part of an eraser to hold them steady. This was then centered on a rotating plate to make photography easier. The camera used was a 45 Megapixel Canon R5, with an RF 100 mm f/2,8L Macro IS USM lens. As the teeth and bone fragments were not flat, it was necessary to use a quite high aperture, i.e. f32, to reduce the depth of field and get as much of each tooth in focus in every shot. As such, additional lighting was required. In this case a setup of three static lights and a ring flash attached to the end of the lens were used to obtain well lit photographs. A relatively fast shutter speed was used to ensure images were sharp and blur free. An average of 141 photographs were taken for each object, with more being taken for larger bone fragments. Estimated image quality is a function in Agisoft Metashape which attributes a value based on the sharpest pixel border within an image and is useful for detecting poor quality and blurred images. Due to the nature of working with macro photography the values were low, but also irrelevant. As such they are not presented in table 1. In this case, each image was evaluated individually with poor quality images removed prior to image alignment.

| Sample | Photos |
|---|---|
| NEOSEA080 | 157 |
| NEOSEA081 | 148 |
| NEOSEA082 | 177 |
| NEOSEA101 | 112 |
| NEOSEA101 | 145 |
| NEOSEA102 | 115 |
| NEOSEA107 | 187 |
| NEOSEA108 | 97 |

| | |
|---|---|
| NEOSEA109 | 160 |
| NEOSEA111 | 139 |
| NEOSEA112 | 112 |

**Supplementary Table S3.2. List of samples, with number of pictures taken for each model.**

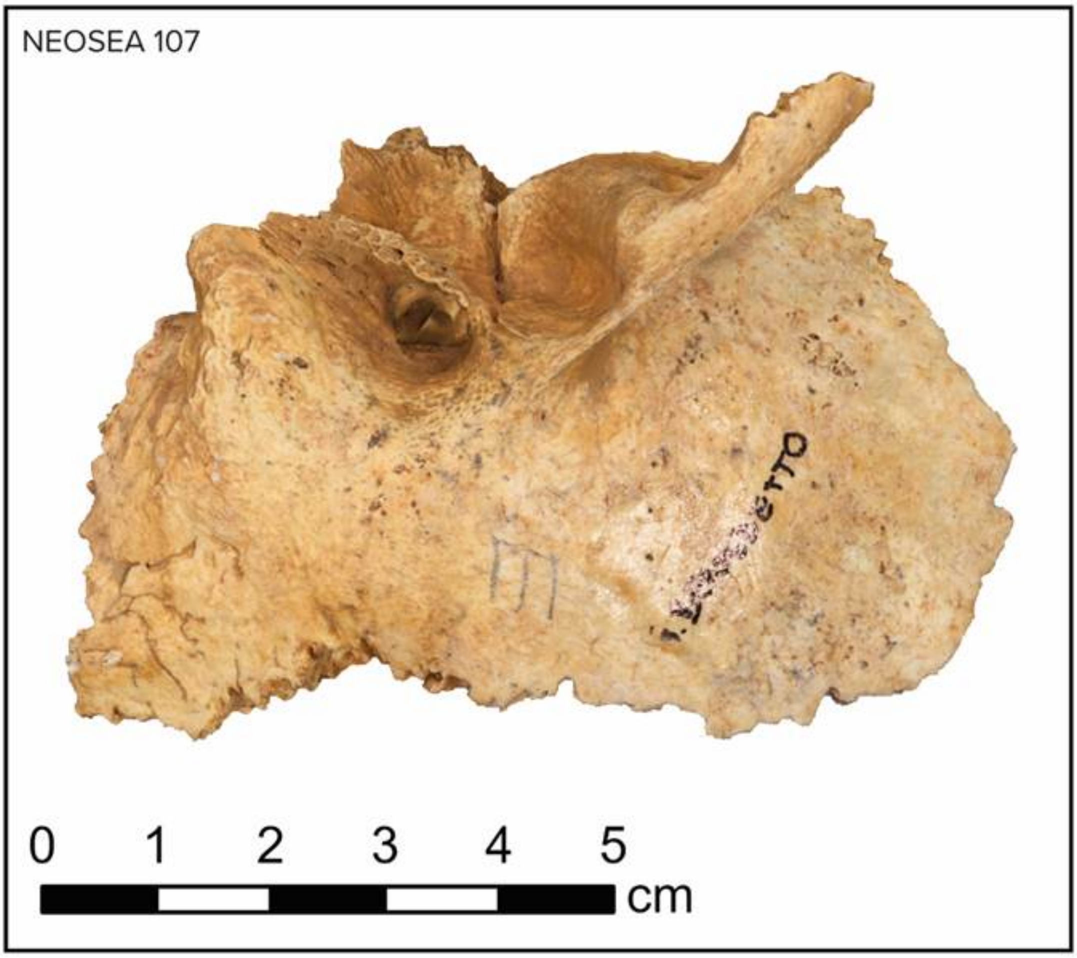

**Supplementary Fig. S3.1. Orthophoto of sample NEOSEA 107.**

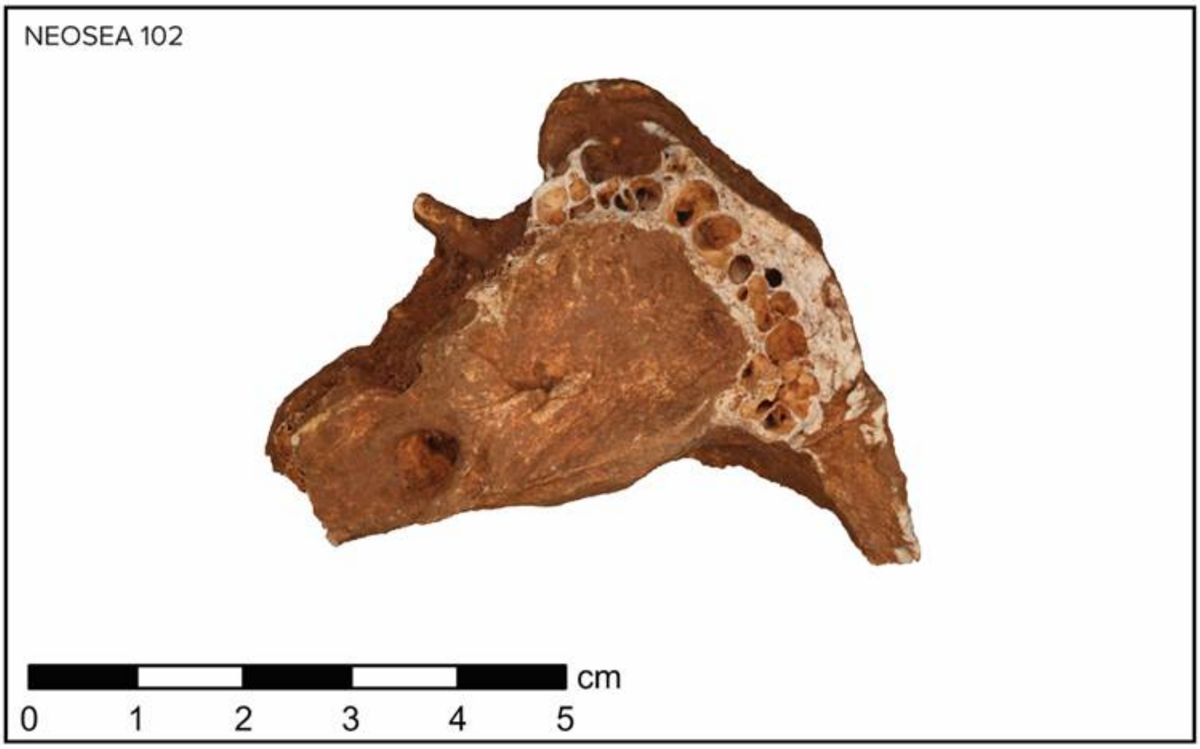

**Supplementary Fig. S3.2. Orthophoto of sample NEOSEA 102.**

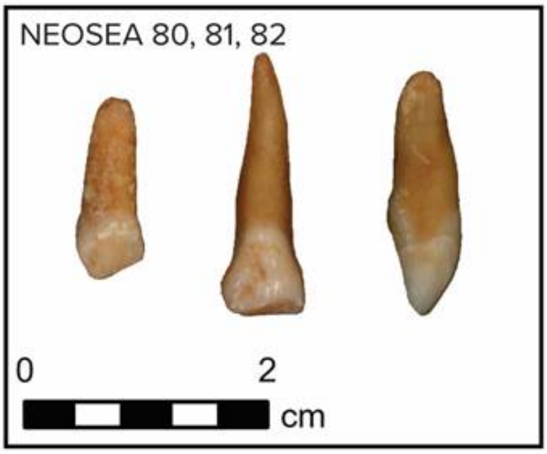

**Supplementary Fig. S3.3. Orthophoto of sample NEOSEA 80, 81 and 82.**

2947 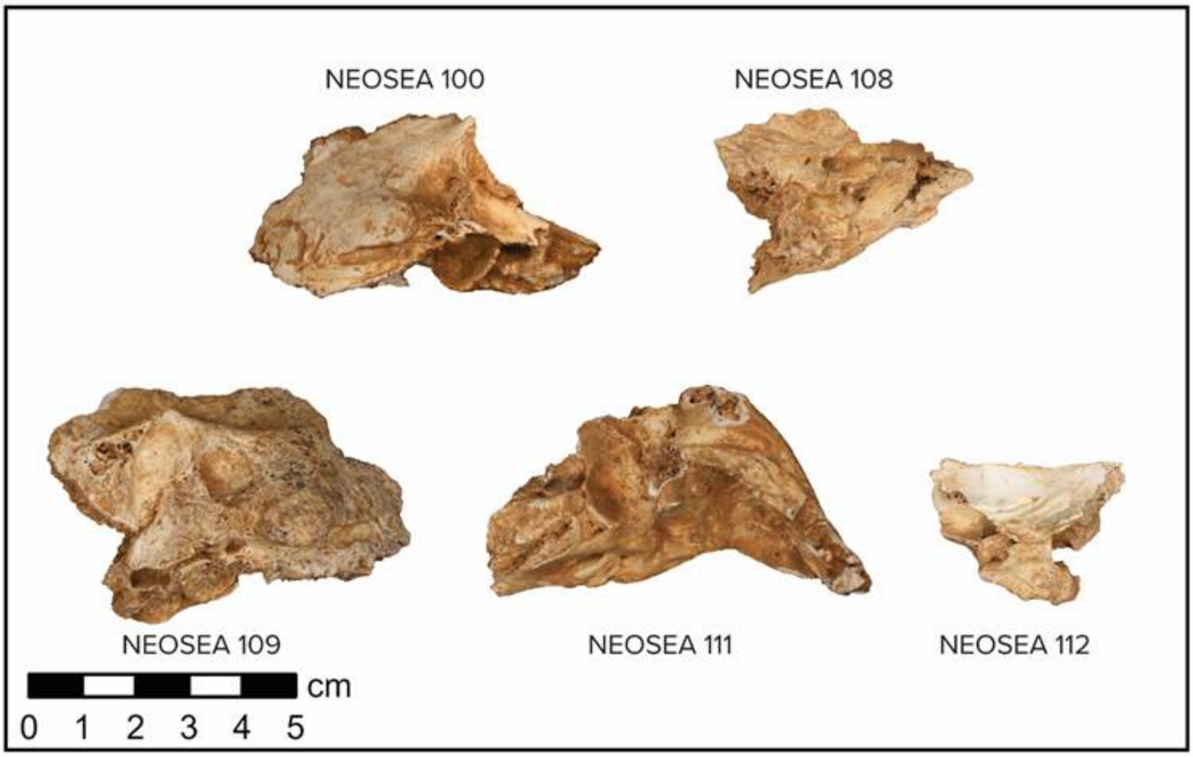

2948 **Supplementary Fig. S3.4. Orthophoto of sample NEOSEA 100, 108, 109, 111 and 112.**

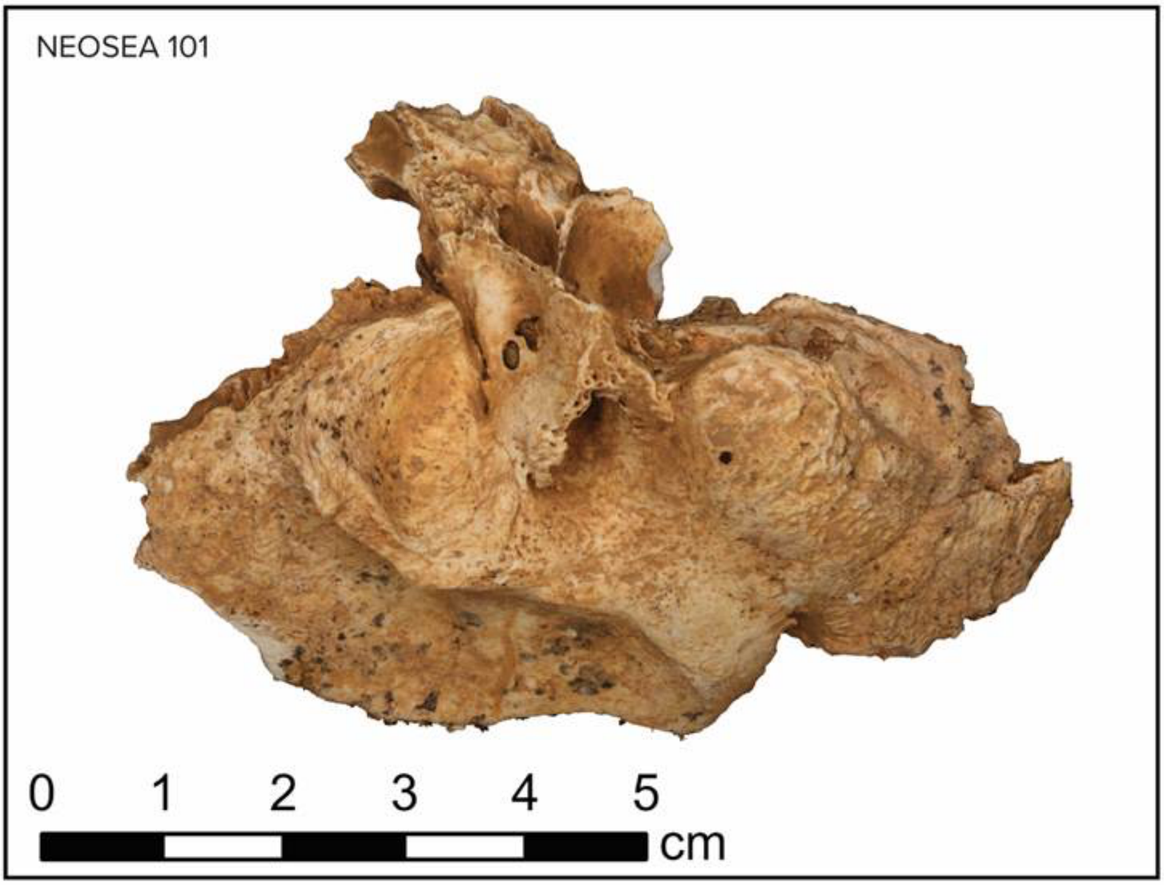

**Supplementary Fig. S3.5. Orthophoto of sample NEOSEA 101.**

The photographs were processed in Agisoft Metashape 2.2.2 (https://www.agisoft.com/),[1] with additional photographs taken when areas were seen to be lacking in resultant models. Masks, areas where the software does not create data during the image alignment process, were automatically generated using AI and corrected manually when necessary. Models were processed at highest settings, going directly from photo alignment to model building from depth maps. Texture maps were created at 8192 by 8192 pixels. A consistent issue throughout was the shiny nature of parts of the enamel on the teeth, which was mitigated by altering lighting and rephotographing as required. Where necessary, models were reprocessed with a point cloud, in order to clean up meshes with point cloud confidence filtering[380].

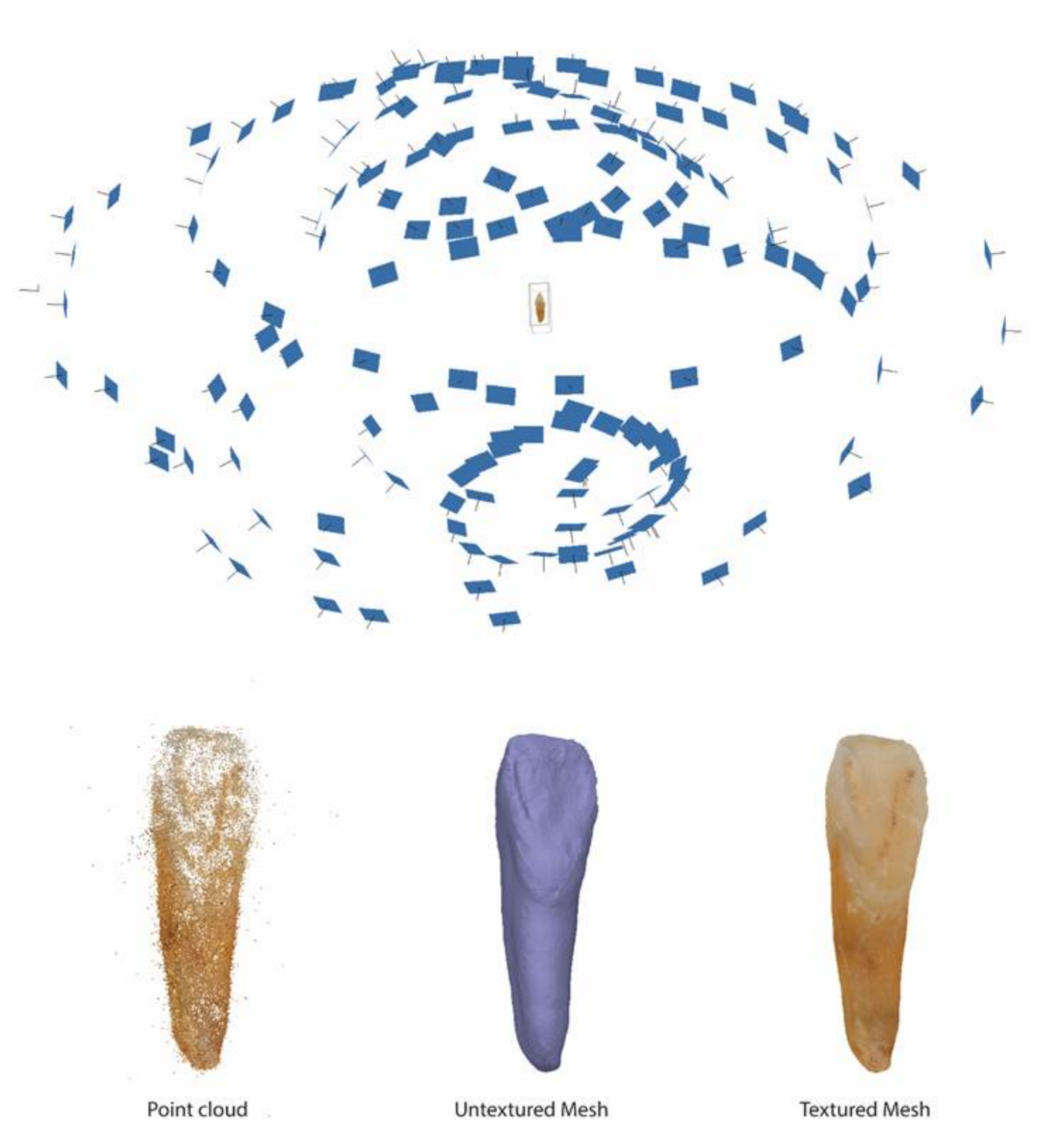

**Supplementary Fig. S3.6. The spread of photographs taken for NEOSEA 82, along with the various stages of processing of the mesh.**

In total eleven teeth were documented within this project prior to them being sampled at the GeoGenetic laboratory in Copenhagen. The output of the documentation means that we have an accurate, colour correct representation of the teeth created prior to them being damaged by sampling. In addition to being able to be presented and shared digitally, they can also be reproduced as 3D printed replicas as required[381].

**References:** [369–376,378–381]

#### 3.3. Methods

*Hugh McColl, Rasmus Nielsen, Thomaz Pinotti*

##### 3.3.1 Population structure of Mesolithic and Neolithic Europe

To explore the population structure of Mesolithic and Neolithic Europe, we performed a principal component (PCA) in plink, using the –pca option (PLINK v1.90b6.21). We included only individuals older than 2800 BP, who were modelled less than 1% Yamnaya ancestry in mixture modelleing (Set 6), and fell within clusters of predominantly Mesolithic (0_4_1_1_, 0_4_1_2_,0_4_1_3_, 0_4_1_4_, 0_4_4_1_, 0_4_4_2_) or Neolithic (0_3_1_1_, 0_3_1_2_1_, 0_3_1_2_2_, 0_3_1_2_3_, 0_3_1_3_, 0_3_1_5_, 0_3_3_1_, 0_3_3_2_, 0_3_3_3_, 0_3_3_4_, 0_3_3_5_, 0_3_4_1_) ancestry. The PCA was run on the remaining 1003 individuals, across 730,235 sites. Individuals were then plotted (Supplementary Fig. S3.7) coloured by age (see ‘ageAverage’ in Supplementary Table S3.3) and archaeological assignment (see ‘arch_group’ in Supplementary Table S3.6)

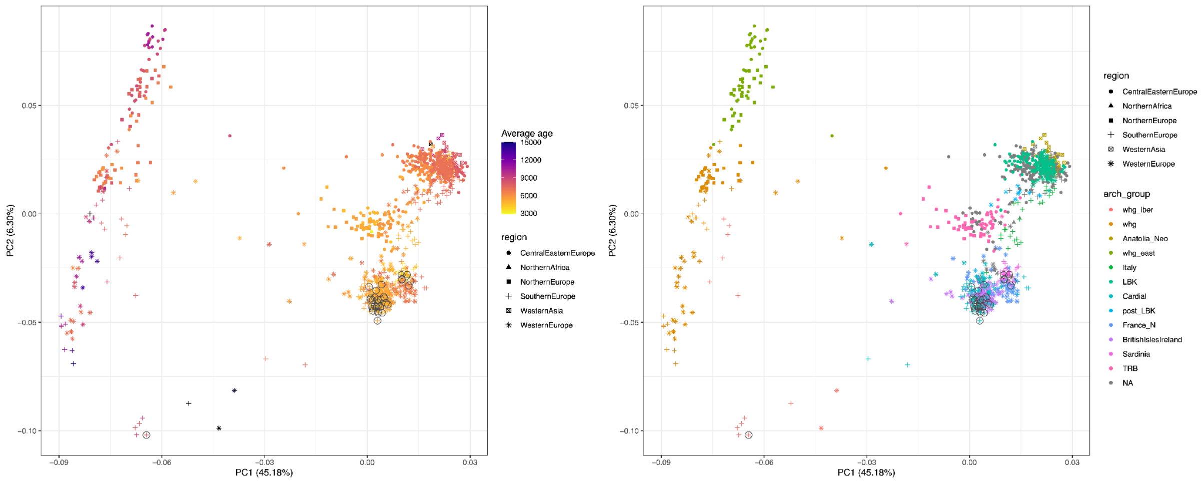

**Supplementary Fig. S3.7. PCA of Eurasian and Western Eurasian ancient populations, coloured by age and by archaeological assignment.** Circled individuals are newly generated in this project.

We find the neolithic individuals fall within three main clusters to the right of the plot. We find most LBK samples fall within the upper cluster, near the early Anatolian individuals from Turkey. Most Cardial individuals fall within the lower cluster. While the upper and lower cluster contain many of the earlier Neolithic individuals, the central cluster contains mainly later individuals, assigned to the TRB culture. The position and age of these clusters are consistent with the mixture modelling results, which suggest that TRB individuals formed as a mixture of LBK and Cardial-related groups.

##### 3.3.2 Testing the association between ancestry components and the emergence of megalithic monuments

<u>Overview</u>

We tested whether the proportion of any inferred ancestry component in a region was associated with the emergence of megalithic monuments in that region. Because emergence is a one-time, irreversible transition, we modeled it as a region-level, absorbing event and estimated its discrete-time hazard as a function of the region's time-varying ancestry composition. Each ancestry component was screened individually and assessed by a permutation test.

Data processing

We obtained ancestry-proportion estimates for the 1,148 ancient individuals and divided them into 12 European regions: Portugal/Spain, France (South), France (North), Sardinia, Britain/Ireland, Denmark/Sweden, Italy (except Sardinia), Latvia/Lithuania/Estonia, Bulgaria/North Macedonia/Romania/Albania/Serbia, Hungary, Greece/Croatia, and Austria/Poland/Czech Republic. We used the previously described six different ancestry paintings (comprising 12,14,15,16,17 and 18 sources, respectively). Each region was assigned a megalith emergence date or was coded as never developing megaliths. Six of the twelve regions were assigned emergence dates: France (North) 6700, France (South), Iberia and Sardinia 6300, Britain/Ireland 5900, and Denmark/Sweden 5600 BP; the remaining six regions were assigned no megaliths.

Discrete-time hazard model

Calendar time was discretised into bins of width $\Delta = 250$ yr. For each region and bin we computed the mean proportion of each ancestry source across the individuals sampled in that bin, giving a time-varying, region-level covariate. Let $h_{rt}$ denote the probability that megaliths first emerge in region $r$ during bin $t$, given that they have not emerged in an older bin. Time runs from older to younger bins; a region is at risk while the bin midpoint is older than its emergence date. Each region contributes a person-period for every observed at-risk bin (event $= 0$) plus the bin containing its emergence date (event $= 1$), after which it leaves the risk set; regions without megaliths remain at risk in all their observed bins and are right-censored (event $= 0$ throughout). Where an emergence bin contained no sampled individual (Sardinia, whose emergence predates its oldest sample), the covariate was imputed from the nearest observed bin and flagged. This construction yields $\approx 105$ person-periods and exactly six events.

The discrete-time hazard is modelled by logistic regression over person-periods,

$$logit\,h_{rt} \;=\; \alpha(t) \;+\; \beta\, x_{rt}^{(k)},$$

where $x_{rt}^{(k)}$ is the proportion of source $k$ in region $r$ at bin $t$ and $\alpha(t)$ is a baseline hazard. Two baselines were used: (A) $\alpha(t) = \alpha_0$ (a constant, giving the marginal association of ancestry with the hazard) and (B) $\alpha(t) = \alpha_0 + \gamma\, t$ (a linear calendar-time term, giving the association conditional on the secular ancestry trend, i.e. beyond time itself).

<u>Per-source screening.</u>

With only six events, joint estimation over all sources is infeasible and maximum-likelihood coefficients are subject to separation. Each source was therefore tested individually using the Rao score statistic and we used Firth's method for estimation of effect sizes. The Rao score statistic for adding source $k$ to the baseline model, evaluated at the baseline (null) fit, is

$$S_k = \frac{U_k^2}{V_k}, \qquad U_k = x_k^\top (y - \hat{\pi}_0), \qquad V_k = x_k^\top W_0 x_k - x_k^\top W_0 Z\,(Z^\top W_0 Z)^{-1} Z^\top W_0 x_k,$$

where $Z$ is the baseline design, $\hat{\pi}_0$ and $W_0 = diag(\hat{\pi}_0(1 - \hat{\pi}_0))$ are the fitted probabilities and weights under the baseline, and $y$ is the event indicator. The score statistic is evaluated at the null and is therefore stable under the near-separation induced by six events and invariant to the scaling of the covariate. Effect directions and magnitudes were summarised by Firth penalised-likelihood coefficients, fitted per standard deviation of the covariate to avoid infinite estimates under separation.

Permutation test

Because the outcome is exchangeable at the level of regions, significance was assessed by permuting the emergence dates (including the never-megalith label) across the twelve regions, holding each region's ancestry trajectory fixed. For each of $5000$ permutations the person-period structure and all score statistics were recomputed. The per-source $P$-value is the permutation-null exceedance probability of the observed statistic. Family-wise error across the sources within a set was controlled by referring each observed statistic to the permutation distribution of the maximum statistic over sources. The full analysis was run under both baselines for all six source sets.

Results

The marginal (A) and time-controlled (B) analyses were concordant, indicating that the associations reported below are not artefacts of the secular increase of some ancestry components through time. Table S3.8  lists, for each source set, the most strongly associated source under the time-controlled model, and Supplementary Fig. S3.8 shows the corresponding association strengths. The regional distribution and temporal trajectory of the best source in each set are shown in Supplementary Fig. S3.9. Across the six decompositions the association strengthened as the source sets became finer, and in every set the best source was one carried at high proportion in western Europe, where megaliths occur.

| Set | Best source | Ancestry | $\boldsymbol{S}$ | $\boldsymbol{\beta}$/SD | $\boldsymbol{P_{perm}}$ | $\boldsymbol{P_{FWER}}$ |
|---|---|---|---|---|---|---|
| set_1 | 0_4_1_3_2 | WHG | 6.8 | +0.62 | 0.0014 | 0.269 |
| set_2 | 0_3_3_4_1_1_2 | Impressed Ware | 8.9 | +1.10 | 0.0028 | 0.187 |
| set_3 | 0_3_1_2_2 | Western Cardial | 17.0 | +1.03 | 0.0006 | 0.038 |
| set_4 | 0_3_1_1_1_2_1 | North French | 28.4 | +0.95 | 0.0008 | 0.006 |
| set_5 | 0_3_1_1_1_2_1 | North French | 26.5 | +0.75 | 0.0016 | 0.010 |
| set_6 | 0_3_1_1_1_2_1 | North French | 26.9 | +0.76 | 0.0008 | 0.009 |

**Supplementary Table S3.8.**

The most strongly associated source is 0_4_1_3_2, a Western Hunter-Gatherer (WHG) source. Its association does not exceed the significance threshold ($P_{FWER} = 0.27$), but the component is present in its highest proportions in the west, where megaliths are found (Supplementary Fig. S3.8). The farming source in this decomposition (an Anatolian-farmer-related source, 0_3_3_1_1_2_2) is, perhaps surprisingly, not the most closely associated, likely due to the geographic distributions of the two components: the WHG component is concentrated in the west and is, therefore, markedly higher in megalith than non-megalith regions (regional means 0.09 versus 0.02), whereas the Anatolian-farmer component is high throughout Neolithic Europe and is, if anything, slightly more abundant in non-megalith regions (regional means 0.67 versus 0.72), so it does not discriminate megalith emergence.

Set 2.

The most strongly associated source is 0_3_3_4_1_1_2, a Cardial-associated Impressed Ware source. Its association likewise does not exceed the threshold ($P_{FWER} = 0.19$), but the component is again in its highest proportions in the west, where megaliths are found (Supplementary Fig. S3.9).

Set 3.

The most strongly associated source is 0_3_1_2_2, a Western Cardial-related source. This is the coarsest decomposition in which the best association exceeds the significance threshold ($P_{FWER} = 0.038$).

Sets 4–6.

The most strongly associated source is the same in all three of the finest decompositions, 0_3_1_1_1_2_1, a North-French-related source ($P_{FWER} = 0.006$–$0.010$; $S \approx 27$–$28$; Firth $\beta \approx +0.9$ per SD). A higher proportion of this component is associated with a higher hazard of megalith emergence. The association is stable to the calendar-time control (baseline B ≈ baseline A) and to exclusion of the imputed Sardinia event ($S$ rises from 28.4 to 31.3, $P_{FWER} = 0.003$), confirming that neither the temporal trend nor the imputation drives it. The component rank-orders the regions almost perfectly by megalith presence (Supplementary Fig. S3.9): it is high in five of the six megalith regions (Britain/Ireland 0.79, France North 0.57, Denmark/Sweden 0.32, France South 0.23, Sardinia 0.18) and low in all six regions that lack megaliths ($\leq 0.12$). The single exception is Iberia (Portugal/Spain), a megalith region carrying a low proportion of this component.

Conclusion

Four of the six decompositions (Sets 3–6) yield a source whose regional abundance is significantly associated with the emergence of megaliths after family-wise correction. The signal sharpens as the decompositions become finer (Supplementary Fig. S3.8): the coarser sets blend the relevant ancestry with more widely distributed farming ancestry and dilute it, whereas the finer sets isolate a North-French-related component that most cleanly separates megalith from non-megalith regions.

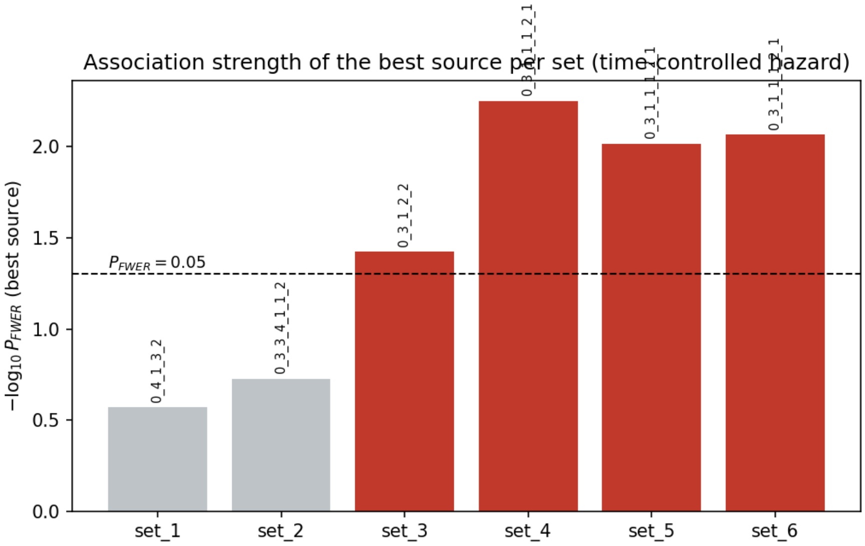

**Supplementary Fig. S3.8. Strength of the association between the most strongly associated ancestry source and megalith emergence, for each source set, under the time-controlled hazard model (permutation family-wise $\boldsymbol{P}$-value, $-\boldsymbol{log_{10}}$ scale).** Red bars exceed the $P_{FWER} = 0.05$ threshold (dashed line); labels give the source. The signal strengthens from the coarsest (set_1) to the finest (set_6) decomposition.

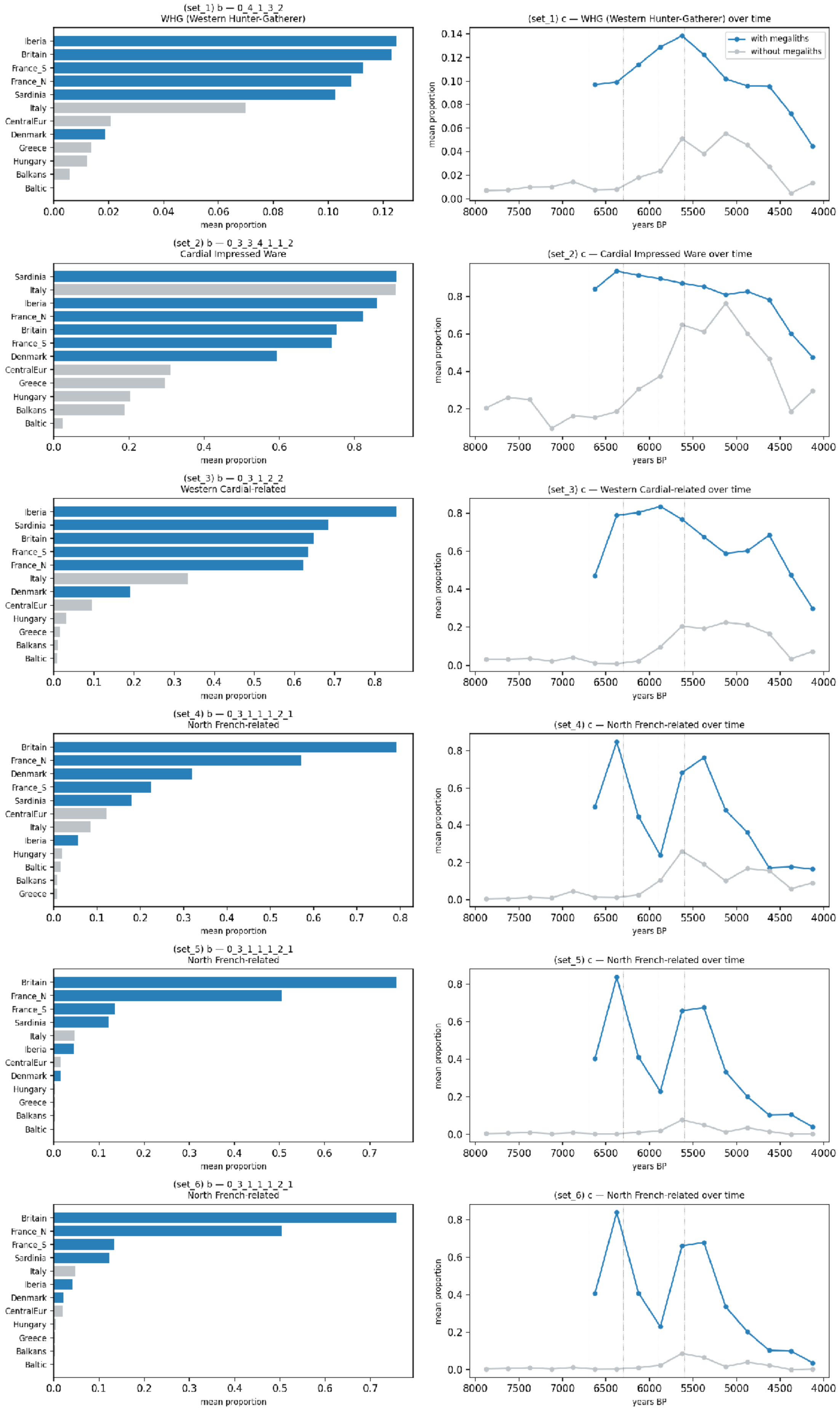

(set_1) b — 0_4_1_3_2
WHG (Western Hunter-Gatherer)
Iberia
Britain
France_S
France_N
Sardinia
Italy
CentralEur
Denmark
Greece
Hungary
Balkans
Baltic
mean proportion
(set_1) c — WHG (Western Hunter-Gatherer) over time
with megaliths
without megaliths
mean proportion
years BP
(set_2) b — 0_3_3_4_1_1_2
Cardial Impressed Ware
Sardinia
Italy
Iberia
France_N
Britain
France_S
Denmark
CentralEur
Greece
Hungary
Balkans
Baltic
mean proportion
(set_2) c — Cardial Impressed Ware over time
mean proportion
years BP
(set_3) b — 0_3_1_2_2
Western Cardial-related
Iberia
Sardinia
Britain
France_S
France_N
Italy
Denmark
CentralEur
Hungary
Greece
Balkans
Baltic
mean proportion
(set_3) c — Western Cardial-related over time
mean proportion
years BP
(set_4) b — 0_3_1_1_1_2_1
North French-related
Britain
France_N
Denmark
France_S
Sardinia
CentralEur
Italy
Iberia
Hungary
Baltic
Balkans
Greece
mean proportion
(set_4) c — North French-related over time
mean proportion
years BP
(set_5) b — 0_3_1_1_1_2_1
North French-related
Britain
France_N
France_S
Sardinia
Italy
Iberia
CentralEur
Denmark
Hungary
Greece
Balkans
Baltic
mean proportion
(set_5) c — North French-related over time
mean proportion
years BP
(set_6) b — 0_3_1_1_1_2_1
North French-related
Britain
France_N
France_S
Sardinia
Italy
Iberia
Denmark
CentralEur
Hungary
Greece
Balkans
Baltic
mean proportion
(set_6) c — North French-related over time
mean proportion
years BP

**Supplementary Fig. S3.9. For each source set (rows), the most strongly associated ancestry source.** Left column: the mean proportion of that source by region; blue = megalith regions (using post-emergence samples only), grey = regions without megaliths. Right column: the mean proportion of the source through time; the blue curve pools megalith regions after their own emergence date ("with megaliths") and the grey curve pools regions that never develop megaliths ("without megaliths"); dotted vertical lines mark the megalith emergence dates. In every set the associated source is more abundant in, and around the emergence of, megalithic populations; the contrast is sharpest for the North-French-related source of Sets 4–6.

Data and software availability

The analysis is fully reproducible from the following files. Data: the input ancestry-painting table is in Supplementary Table S3.9. Results: the complete per-source output for all six sets under the marginal and time-controlled baselines (Supplementary Table S3.10 and Supplementary Table S3.11, respectively), each listing every ancestry source with its Firth coefficient per standard deviation, Rao score statistic, asymptotic $P$-value, permutation $P$-value, and family-wise permutation $P$-value. Software: the person-period risk-set construction (prep.py), the Firth hazard model, score test and permutation procedure (model.py), and the figure-generation script (make_figures.py), written in Python 3 using numpy, pandas, scipy and matplotlib. All permutation analyses use a fixed random seed.

##### 3.3.3 Y-chromosome haplogroup assignment

Y-chromosome haplogroups were assigned using ybyra v0.5[382], with parameters optimised for the characteristics of ancient DNA. We used a step size of 5 (--step-size 5) and enabled damage filtering (--damage-filter true), with the damage model set to bidirectional (--damage-model bi). Haplogroup assignment was performed against the FamilyTreeDNA Y-DNA Haplotree release v.2026.06.01, which was used as the reference nomenclature and phylogenetic framework.

We included all ancient male individuals that had at least 50% Neolithic farming-related ancestry in our IBD-based mixture modelling analyses. To provide additional context for the early diversification of Y-chromosome lineages, we additionally included Mesolithic and Palaeolithic male individuals from Europe. The resulting dataset comprised 611 ancient males. For each individual, we recorded the most downstream Y-chromosome haplogroup that could be confidently assigned under the parameters described above. Individual Y-chromosome haplogroup assignments, together with the corresponding sample information, are provided in Supplementary Table S3.7.

We reconstructed a Y-chromosome phylogeny including all 611 individuals using the SNP-defined relationships represented in the FamilyTreeDNA Haplotree. The resulting phylogeny was calibrated according to the number of diagnostic SNPs defining each node and was used to visualise the phylogenetic relationships among the sampled individuals and to identify the position of ancient individuals within individual Y-chromosome subhaplogroups. The complete SNP-number-calibrated Y-chromosome phylogeny is provided in Supplementary Fig. S3.10. Individuals are coloured according to IBD clustering and nodes according to its oldest representative downstream.

Association of Y-chromosome lineages with autosomal ancestry

To investigate the relationship between Y-chromosome subhaplogroups and autosomal ancestry, we combined the Y-chromosome phylogeny with the IBD-based mixture modelling results. We focused on mixture-modelling sets 3, 4 and 5, which provided the ancestry components used for distinguishing individuals associated with Cardial- and LBK-related ancestry (Supplementary Fig. S3.11)

We examined the distribution of Y-chromosome lineages within the phylogeny while simultaneously considering the autosomal ancestry profiles of the individuals carrying those lineages. To assess the temporal structure of these associations, we repeated this inspection after restricting the dataset to individuals older than 6,300 years ago and to individuals older than 4,800 years ago. These time-restricted phylogenies are shown in Supplementary Fig. S3.12 and Supplementary Fig. S3.13, respectively.

We then systematically screened the resulting phylogenies to identify Y-chromosome subhaplogroups whose multiple individuals showed a consistent association with either Cardial-related or LBK-related autosomal ancestry. Using these criteria, we identified eight principal Y-chromosome subhaplogroups associated with LBK-related ancestry: lineages downstream of C-PH428, G-L91, G-PF3359, G-Z6885, G-PF3345, H-FT68618, J-FTF148 and I-BY136048. We similarly identified 15 principal subhaplogroups associated with Cardial-related ancestry: G-PF3178, G-FTA29810, G-BY201148, G-PF3378, G-BY49026, H-Z19072, I-S9403, I-FGC14213, I-BY1003, I-CTS11030, I-PF3981, I-L1298, I-FT344600, I-CTS5375 and R-FGC21027. The designation of these lineages reflects their distribution among the ancient individuals included in our dataset and their correspondence with the autosomal ancestry profiles inferred from the mixture-modelling analyses.

For visualization and further examination of these patterns, we generated a reduced SNP-calibrated Y-chromosome phylogeny containing the identified key lineages and the ancient individuals carrying them. This phylogeny is provided in Supplementary Fig. S3.14. We plot the geographical distribution of these lineages before 6300 BP, and between 6300 and 4800 BP in Supplementary Fig. S3.15.

---

LARGE FIGURE INCLUDED SEPARATELY

---

**Supplementary Fig. S3.10. Y-chromosome phylogeny of all ancient Western Eurasian males.** Complete SNP-calibrated Y-chromosome phylogeny of the 611 ancient male individuals, with individuals coloured by IBD cluster and internal nodes according to the oldest downstream representative.

---

LARGE FIGURE INCLUDED SEPARATELY

---

**Supplementary Fig. S3.11. Y-chromosome phylogeny of ancient all European males and respective mixture modelling results for set 3, 4 and 5.**

---

LARGE FIGURE INCLUDED SEPARATELY

---

**Supplementary Fig. S3.12. Y-chromosome phylogeny of ancient European males 6300 BP+ with respective mixture modelling results for sets 3, 4 and 5.**

---

LARGE FIGURE INCLUDED SEPARATELY

---

**Supplementary Fig. S3.13. Y-chromosome phylogeny of ancient European males 4800 BP+ with respective mixture modelling results for sets 3, 4 and 5.**

---

LARGE FIGURE INCLUDED SEPARATELY

---

**Supplementary Fig. S3.14. Y-chromosome phylogeny of ancient European males of Cardial/Neolithic ancestry, with respective mixture modelling results for sets 3, 4 and 5**

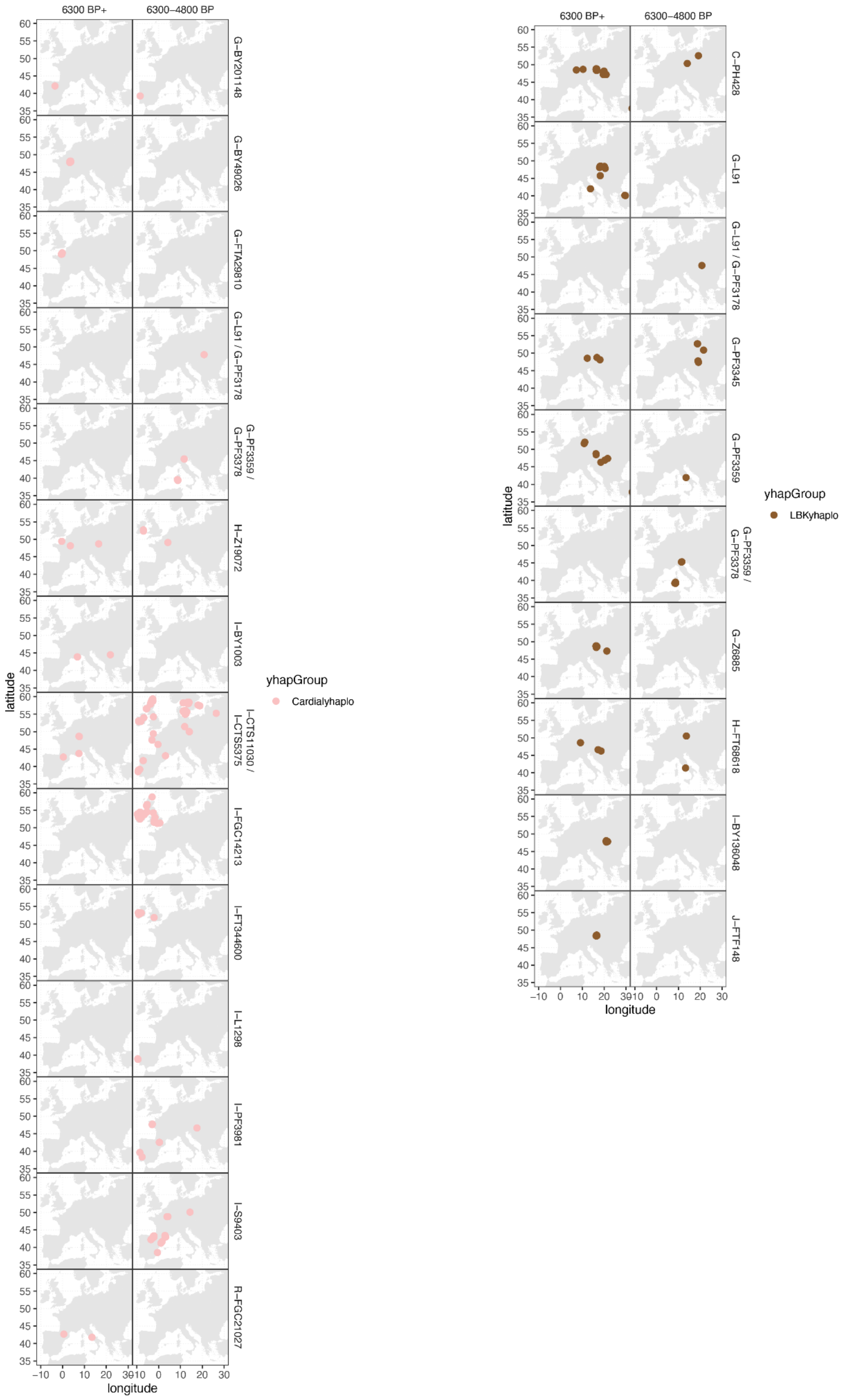

**Supplementary Fig. S3.15. Distribution of principal Y-haplogroups subgroups associated with Cardial and LBK ancestry before 6,300 BP and between 6,300 and 4,800 BP.** Y-haplogroups were classified as Cardial- or LBK-associated based on their age and their occurrence among multiple individuals carrying the corresponding autosomal ancestry profiles in the IBD mixture modelling analyses.

Phylogeographic interpretation of Y-chromosome lineages

The phylogenetic distribution of these lineages provides additional information about the relationship between Y-chromosome diversity and the spread of Neolithic ancestry. Several of the major Y-chromosome haplogroup branches represented in our dataset, including C, G, H and J, contain their oldest sampled individuals among Neolithic individuals from Anatolia. This observation is consistent with an ultimate origin of these Y-chromosome lineages in Neolithic Anatolia, despite the age of the oldest sampled individual does not by itself establish the geographic origin of the corresponding haplogroup.

At the same time, several of the lineages associated with Neolithic farmer-related autosomal ancestry belong to haplogroup I, a Y-chromosome haplogroup that is strongly represented among European hunter-gatherers. The significant occurrence of I lineages among Neolithic individuals therefore indicates that Y-chromosome diversity associated with European hunter-gatherer populations was incorporated into populations carrying Neolithic farmer-related autosomal ancestry.

---

LARGE FIGURE INCLUDED SEPARATELY

---

**Supplementary Fig. S1.16. IBD mixture modelling results (Sets 1–6) for all ancient samples in the dataset (n=2963), faceted by region and country.** Standard errors for each individual's ancestry proportions were calculated using a weighted block jackknife with n=22 chromosomes, sequentially leaving out one chromosome at a time.

#### 3.4 The archaeological context of the Sardinian samples

*Maria Grazia Melis, Vitale Sparacello, Anna Bini*

##### 3.4.1 The Domu de janas II necropolis at San Benedetto -Iglesias, Cagliari

###### 3.4.1.1 Archaeological data

The necropolis at San Benedetto is situated in the mineral-rich basin at Sulcis-Iglesiente (south-western Sardinia). It is formed of five large tombs dug into schist, almost all of them damaged by mineral extraction or farming work. The funerary architectural category is that of rock-cut tombs, which in Sardinia first appear during the first half of the 5th millennium cal BC (~7000-6500 BP) at Cuccuru s'Arriu (central-eastern Sardinia); during the second half of the millennium it evolved into the type of complex-plan hypogeum, locally known as *domu de janas*. These tombs numbered more than three thousand during the 4th millennium cal BC (~6000-5000 BP) and were present across the entire island, with greater concentrations in north-western Sardinia.

The five *domus* at San Benedetto had simple elongated plans with concave ceiling chambers. Tomb II was investigated during the 1960s[364,365,383]. The discovery was immediately held to be of extreme importance, as for the first time a *domu de janas* had been found sealed and unviolated[384].

This tomb stands out from the others in the necropolis for its greater size and for its cruciform plan, that consisted of a central sub-trapezoidal chamber, from which an opening in each of three walls led to peripheral rooms with a curved profile plan. The entrance, facing to the south-east, was discovered closed by a quadrangular stone slab. Similar stone slabs covered the entrances to the inner rooms.

The archaeological deposit was covered by a wind-carried sterile layer roughly 50 cm thick, that spread through all the internal spaces. Below this, touching the floor, the skeletal remains of 35 individuals were found, distributed across the four internal chambers. Based on the absence of anatomical connection and on the presence of only selected postcranial elements (mostly long bones and pelvic remains), it was immediately apparent that human remains were in secondary deposition. Preliminary studies of the remains were made by C. Maxia (see ivi bibliography in ref [364]) and subsequently by R. Floris[365]. The crania were present in each of the rooms: 15 in the central chamber, 10 in the north-west chamber, 5 to the north-east and 5 in the southern chamber. They were, however, more numerous in the two larger chambers, while equally distributed between the two, smaller, lateral rooms, that contained 5 each. The distribution of the crania (Supplementary Fig. 3.17) does not appear accidental; the spaces between them vary between a minimum of 4 cm and a maximum of 60 cm.

On the low raised doorstep of the entrance to the central chamber the excavation revealed the presence of ash, burnt wood and semi-combusted fauna remains, which are probably related to funerary practices performed inside the tomb, perhaps the last of such rites, considering their position at the entrance to the *domus de janas*. The archaeological deposit, 34/38 cm thick, was delimited to the south-east by vertical stones and slabs, while in proximity to the entrances to the north-west and north-east chambers it was partially covered by two horizontal slabs. The prevailing distribution of the crania was along the southern wall. On the basis of the distances between the crania (minimum 5 cm, maximum 75 cm) it is possible to identify a group of 6 to the south-east, another group of 6 to the west, 2 to the north and 1 to the east. In each of the two groups of 6 skulls, 4 were placed against the south wall and 2 were slightly further away.

In the north-west chamber, taking account of the distances between the crania (minimum 6 cm, maximum 76 cm) it is possible to see two groups of 4 crania close to the southern corner and the north-eastern wall respectively; two crania were placed close to the north-west wall.

In the north-eastern chamber the crania (separated by distances of between 4 cm and 21 cm) were distributed in a group of four close to the south-east wall with one isolated cranium next to the north-west wall.

Lastly, in the southern chamber, in which the distances between the crania were 6 cm and 34 cm, there was a group of 2 crania close to the south-east wall and a group of 3 more orthogonally aligned with the north-western wall.

The archaeological finds and radiocarbon dates[365–367] make it possible to attribute this context to the Sardinian Final Neolithic and in particular to the first phase of the Ozieri cultural facies (table 1). The new radiocarbon determinations confirm the use of the tomb within the same chronological range.

| **Lab. n.** | **Individual** | **Date BP** | **Date cal. BC (95.4%)** | **Bibliography** |
| --- | --- | --- | --- | --- |

| | | | | |
|---|---|---|---|---|
| AA78327 | 4 | 5078 ± 58 | 3982-3710(95.4%) | Lai 2009 |
| AA78328 | 5 | 5044 ± 58 | 3963-3706 (93.5%)<br>3672-3658 (1.9%) | Lai 2009 |
| AA78330 | 25 | 4984 ± 52 | 3946-3856 (22.4%)<br>3845-3834 (1.2%)<br>3819-3647(71.8%) | Lai 2009 |
| AA78329 | 18 | 4969 ± 52 | 3943-3862 (16.2%)<br>3814-3642 (79.3%) | Lai 2009 |
| Beta-72233 | Unknown | 4920 ± 70 | 3945-3857 (10.5%)<br>3818-3622 (80.0%)<br>3582-3530 (5.0%) | Floris 2001 |

**Supplementary Table S3.4 . Radiocarbon dates from Tomb II at San Benedetto (previous to this study).** The data were calibrated in OxCal v.4.4.4 using the IntCal20 atmospheric curve[33,385].

The pottery finds include forms and decorations typical of the Ozieri I repertoire; hemispherical and truncated cone bowls, carenated cups, ollas, undecorated vases or vessels with impressed or incised decorations. Amongst the stone tools the presence of artefacts made from raw materials of non-local origin is worthy of note: large blades and bladelets of Perfugas flint[386] and obsidian arrow heads from Monte Arci. Other finds included a polished greenstone axe, an elliptical bead in black anthracite and a spheroidal perforated stone in pink granite.

###### 3.4.1.2 Previous genetic studies on Sardinian material

Two recent genome-wide ancient DNA studies have contributed key data for understanding population continuity and change in Sardinia from the Neolithic through the historical period. Marcus et al.[387] present an extensive genome-wide ancient DNA dataset focused specifically on Sardinia. The study includes 70 individuals from more than 20 archaeological sites, spanning from the Middle/Late Neolithic to the Medieval period. Six Neolithic individuals are grouped under Sar-MN (Middle/Late Neolithic Sardinia) and derive from four funerary contexts, three *domus de janas* and a cave site. All individuals included in this group are directly radiocarbon dated on human remains. Fernandes et al. [79] present a genome-wide ancient DNA study investigating population dynamics across the western Mediterranean islands, including Sardinia, Sicily, and the Balearic Islands. Sardinia is not the primary focus of the study, but several Sardinian individuals are included and analysed within a comparative framework.

Genome-wide analyses in Marcus et al.[387] indicate that Neolithic individuals from Sardinia (grouped as Sar-MN) form a genetically homogeneous cluster and show limited internal differentiation. Neolithic inhabitants of Sardinia are characterised by ancestry profiles typical of western European Neolithic populations, with no evidence for distinct substructure corresponding to archaeological phases. Formal admixture modelling shows that Neolithic Sardinians can be described primarily as deriving from Early European Farmer–related ancestry, combined with a moderate contribution of Western Hunter-Gatherer (WHG) ancestry, estimated at approximately 15–20%. This ancestry composition remains stable throughout the Neolithic sequence and shows close similarity to that observed in contemporaneous western Mediterranean populations. Individuals from the later Early Copper Age (Sar-ECA) are genetically indistinguishable from Neolithic Sardinians[387]. Measures of genetic differentiation between Sar-MN and Sar-ECA are extremely low, and models of direct population continuity cannot be rejected.

Results from Fernandes et al.[79] are similar: the Neolithic individuals from Sardinia (grouped as Sardinia_Neolithic, n = 3) are genetically homogeneous and cluster with mainland European Early and Middle Neolithic farmers. They are described as a mixture of Anatolia Neolithic–related ancestry and Western Hunter-Gatherer (WHG) ancestry. Most Copper Age individuals from Sardinia form a clade with the Neolithic group, supporting strong genetic continuity. A notable exception is one individual showing substantial North African–related ancestry.

###### 3.4.1.3 List of sites included and brief description:

**Grutta I de Longu Fresu (Seulo, central-eastern Sardinia; Marcus et al.[387]):** A small cave used during the Late Neolithic for mortuary purposes, with later reuse. Human remains attributed to the Neolithic derive from disturbed burials at the rear of the cave and are directly radiocarbon dated.

**Noeddale (Ossi, north-western Sardinia; Marcus et al.[387]):** A *domu de janas* attributed to the Late Neolithic with later reuse. Two individuals from Tomb III were included in the aDNA dataset and directly dated.

**S'Isterridolzu (Ossi, north-western Sardinia; Marcus et al.[387]):** A *domu de janas* necropolis with evidence of later reuse. Three individuals from Tomb IV were sampled; only one yielded a radiocarbon date consistent with the Late Neolithic and was included in the Sar-MN group.

**Su Crucifissu Mannu, t. 12 (Porto Torres, north-western Sardinia; Marcus et al.[387]):** the tomb belongs to a *domus de janas* necropolis, used from the late Neolithic to the early Bronze Age. One individual yielded a radiocarbon date consistent with the Late Neolithic and was included in the Sar-MN group.

**Anghelu Ruju (Alghero, north-western Sardinia; Fernandes et al.[387]):** A *domu de janas* necropolis with evidence for multi-period use from the Late Neolithic through the Bronze Age. One individual yielded a radiocarbon date consistent with the Late Neolithic, and the authors attributed to the same layer other four individuals. Only three individuals were ultimately included in the analysis due to contamination of the remaining samples.

| ***Site*** | ***Sample ID*** | ***Dating Lab Code*** | ***Radiocarbon date (BP)*** | ***Calibrated (cal BC, 95.4%)*** |
|---|---|---|---|---|
| *Grutta I de Longu Fresu* | *LON003* | *MAMS-39524* | *5481 ± 32 BP* | *4442–4252* |

| | | | | |
|---|---|---|---|---|
| *Grutta I de Longu Fresu* | *LON004* | *MAMS-39525* | *5375 ± 33 BP* | *4333–4058* |
| *Noeddale* | *NOE002 (Noe5/)* | *MAMS-39527* | *5352 ± 32 BP* | *4325–4053* |
| *S'Isterridolzu* | *SID005 (SI7)* | *MAMS-39532* | *5316 ± 31 BP* | *4310–4047* |
| *S'Isterridolzu* | *S1 2* | *MAMS-26898* | *5309 ± 30 BP* | *4246-4046* |
| *Noeddale* | *NOE001 (Noe4)* | *MAMS-39526* | *5306 ± 32 BP* | *4246–4000* |
| *Noeddale* | *Noe 2* | *MAMS-26899* | *5296 ± 30 BP* | *4240-3998* |
| *S'Isterridolzu* | *S1 1* | *MAMS-26897* | *5279 ± 30 BP* | *4234-3991* |
| *Grutta I de Longu Fresu* | *MA79 / LF1002* | *OxA-22195* | *5258 ± 34 BP* | *4231–3982* |
| *Anghelu Ruju* | *I15946 (2485, Tomb F)* | *PSUAMS-6699* | *5205±30 BP* | *4158-3958* |

| | | | | |
|---|---|---|---|---|
| *Su Crucifissu Mannu, t.22* | *SUC008* | *MAMS-38301* | *5042 ± 21* | *3947-3779* |
| *Noeddale* | *Noe 3* | *MAMS-26900* | *4853 ± 27 BP* | *3704-3532* |
| *Anghelu Ruju* | *I15943 (2482, Tomb F)* | *No direct date. Layer date based on PSUAMS-6699* | | |
| *Anghelu Ruju* | *I15945 (2484, Tomb F)* | | | |
| *Anghelu Ruju (contaminated, not included in analysis)* | *I15944 (2483, Tomb F)* | | | |
| *Anghelu Ruju (contaminated, not included in analysis)* | *I15947 (2486, Tomb F)* | | | |

**Supplementary Table S3.5.** ***Neolithic individuals included in the genetic analyses by Olivieri et al.[388], Marcus et al.[387] and Fernandes et al.[79] .*** In the Supplementary Information of Marcus et al.[387], some Neolithic samples from Noeddale (NOE001, NOE002) and S'Isterridolzu (SID005) are associated with calibrated ranges inconsistent with their radiocarbon ages (~5300–5000 BP). Recalibration using OxCal v4.4 and IntCal20 places these samples in the late 5th–early 4th millennium cal BC (~6500-5500 BP), consistent with a

Neolithic attribution. These individuals are correctly assigned to the Sar-MN group in the original analyses.

Earlier genetic investigations of prehistoric Sardinian populations[388] were limited to mitochondrial DNA or targeted loci and did not provide genome-wide resolution. Olivieri et al.[388] analysed thousands of modern and 21 ancient mitogenomes from Sardinia and found that most Sardinian-specific mitochondrial lineages coalesce in the Neolithic-Copper Age, Nuragic, and post-Nuragic periods, with modern people from Sardinia forming a distinctive cluster closely associated with early European farmers and with a possible minor contribution from pre-Neolithic lineages. The five Neolithic samples included in the study derived from some of the sites mentioned above (Noeddale, S'Isterridolzu and Grutta I de Longu Fresu).

###### 3.4.1.4 Chronological position of San Benedetto relative to previous Neolithic aDNA data

The necropolis of San Benedetto is dated by multiple radiocarbon determinations to the Final Neolithic (~4000–3500 cal BC/~6000-5500 BP). While this interval formally falls within the broad Sar-MN temporal range of Marcus et al.[387], the San Benedetto assemblage post-dates most previously published Neolithic aDNA samples, which cluster primarily between ~ 4400 and 4000 cal BC (~6400-6000 BP, see also Fernandes et al.[79]). As a result, chronological overlap with the existing Sar-MN dataset is limited to the individual of Su Crucifissu Mannu, t. 22.

The San Benedetto site therefore extends Neolithic genetic sampling toward the first half of the 4th millennium cal BC (6000-5500 BP), immediately preceding the Early Copper Age, rather than duplicating the core chronological range already represented in aDNA studies. In addition, the sampling of 17 individuals from a single, sealed *domus de janas* tomb provides an unprecedented opportunity to investigate kinship structure, social organisation, and collective burial practices within a well-defined Neolithic community, adding contextual and social resolution to existing Neolithic genetic data from Sardinia.

**References:** [79,364,365,367,383,384,386–388]

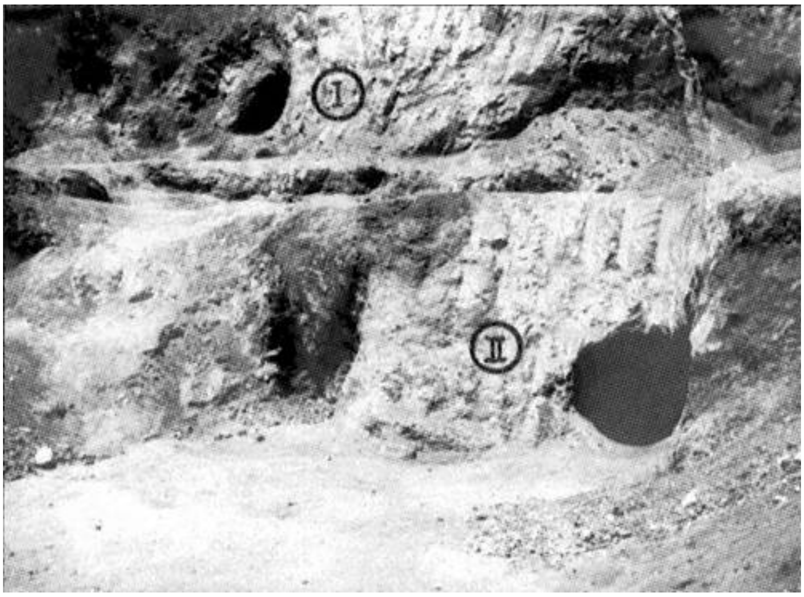

**Supplementary Fig. 3.17. The entrances to Tombs I and II of San Benedetto, brought to light and partially destroyed by a bulldozer (from Atzeni[383], fig. 1,2).**

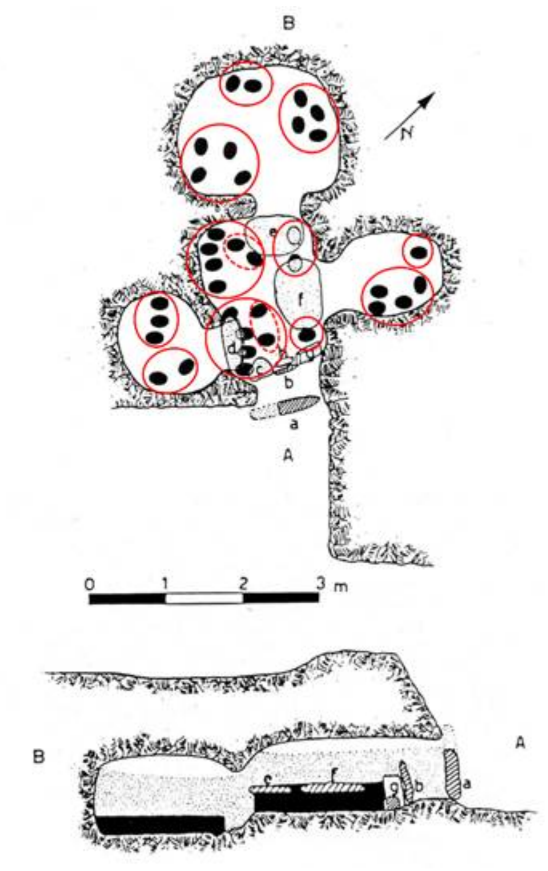

**Supplementary Fig. S3.18. Plan and section of Tomb II at San Benedetto (adapted by M.G. Melis from Atzeni[383], fig. 6).**

#### 3.5 The archaeological context of the Portuguese samples

*Rita Peyroteo Stjerna, Catarina Costeiro, Eduardo Porfirio, António Faustino Carvalho, Bettina Schulz Paulsson, Ana Maria Gama da Silva*

##### 3.5.1 The Tholos of Praia das Maçãs

***Description of architecture***

The prehistoric monument of Praia das Maçãs is located near to the top of the northern slope of Outeiro das Mós, on the north bank of Ribeira de Colares, close to the sea. This monument was excavated in rocks of the Lower, Middle and Upper Cenomanian period. This monument is composed of several funerary structures, with diverse architectural features, namely an artificial cave and a tholos.

Brief description of research history: Prehistoric burial structures were identified in the 1920s, during agricultural work. The first archaeological excavation took place in 1961, led by V. Leisner and O. V. Ferreira, and allowed the identification of a funerary complex, consisting of an artificial cave built in the late Neolithic and a tholos built and widely used during the Chalcolithic, with subsequent reuses. In the late 1970s, J.L Gonçalves excavated the entrance and an area of about 125 $m^2$ outside, especially on the northern slope. Between the 1980s and 2020, geological and petrographic analyses, radiocarbon dates, studies of archaeological materials and anthropological remains were carried out.

Since 2020, the Sintra municipality has been developing a project for the safeguarding of the monument that includes conservation and restoration work, supported by an archaeological research and excavation program. These works made it possible to document a complex stratigraphic sequence, which covers a temporal gap marked between the middle/end of the 4th millennium BC and the present day (5500/5000 cal BP to 0 BP), with special emphasis on the uses of the 3rd millennium BC (5000-4000 BP). They also made it possible to record and describe in greater detail the architectural and structural characteristics of this monument, as well as the ceramic deposits on the exterior, and to carry out a retrospective stratigraphic analysis, through the interrelationship between recent data and information obtained in 1961 and 1979.

Brief description of the context: The western chamber (artificial cave), with a sub-circular shape, was excavated in the argillaceous geological substrate and uses the limestone blocks that outcrop on the surface as a covering. Access to this chamber consisted of a paved ramp with a step. Several individuals were buried inside, accompanied by objects from the Late Neolithic/Early Chalcolithic (3500 – 2800 BC/~5500-4800 BP), including arrowheads, flint blades, bone and ivory pins, disc-shaped beads and decorated schist plaques.

The tholos consists of a large chamber with an oval shape and walls made of rows of stones, which is accessed via a long corridor and atrium. In these spaces, during the Chalcolithic (2800 – 2200 BC/~4800-4200 BP), a significant number of individuals of both sexes and different ages were deposited, together with ceramic vessels and numerous votive artefacts in bone and limestone. Fragments of containers with bell beaker decoration and copper weapons were collected in the corridor and at the entrance to the chamber.

In the external area of the tholos, multiple intentional depositions of whole or broken prehistoric ceramic vessels were identified, probably associated with ritual celebrations in memory of the ancestors.

**References:** [197,358,389–402]

Archive of the bones/materials: Museu Geológico / Museu Arqueológico de São Miguel de Odrinhas - Câmara Municipal de Sintra / Museu Nacional de Arqueologia

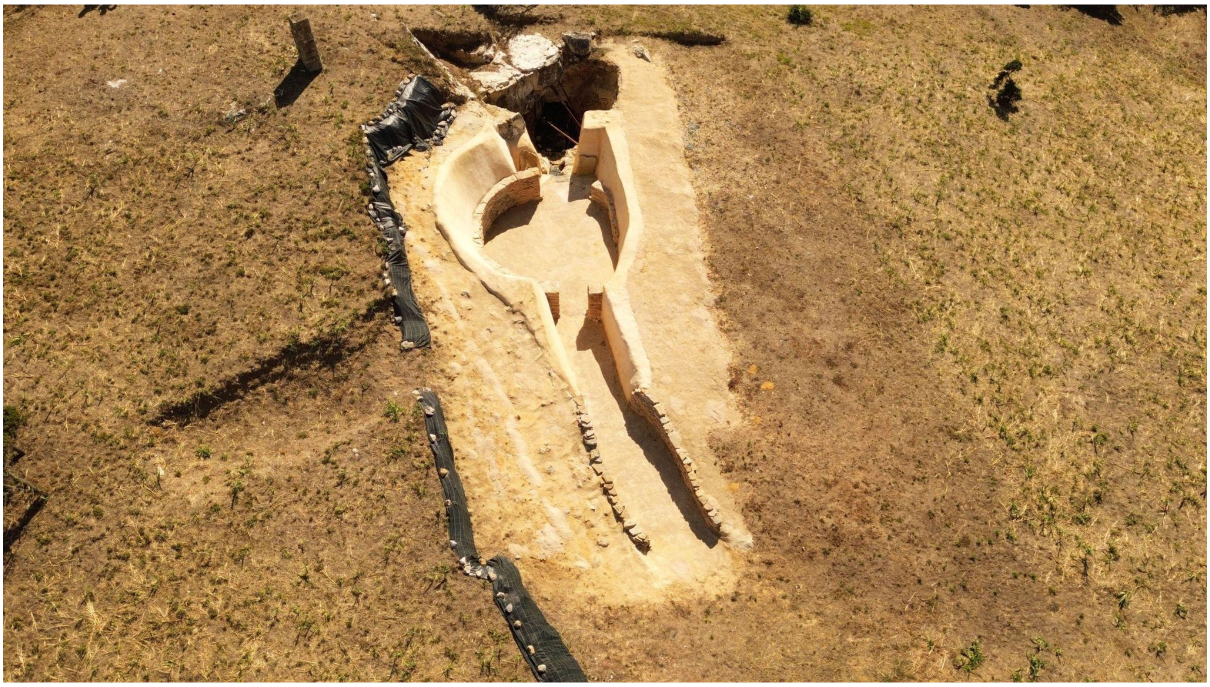

**Supplementary Fig. S3.19. Praia_Macas_001: Orthophotographic record of the prehistoric monument at Praia das Maçãs.** The monument is a so-called tholos with a round megalithic chamber and a corridor.

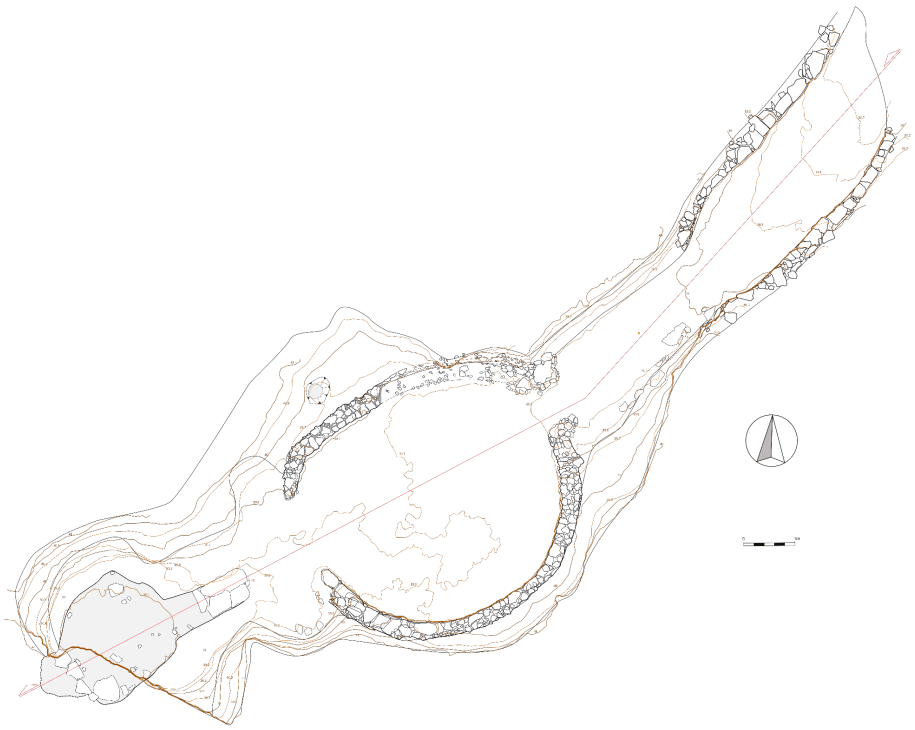

**Supplementary Fig. S3.20. General plan of the prehistoric monument at Praia das Maçãs.**

##### 3.5.2 Dolmen Pedras da Granja

Description of architecture: Dolmen Pedras da Granja is located on S. João das Lampas Plateau.

The burial chamber of this dolmen would have a polygonal plan, with seven upright megaliths. The work carried out during the 20th century, was not sufficient to identify the remains of the corridor and the mound.

Brief description of research history: This site was excavated in the 1950s[403] and 1970s[404], with some of its slabs being destroyed in the 1990s. In archaeological excavations, a significant and diverse set of artefacts and osteological remains were identified.

Brief description of the context: By the time the dolmen was identified, only two large sloping slabs, with almost 2,6m in height, remained of the original megalithic tomb, which was part of a dry-stone wall that separated the rural road from a property. During the archaeological works, some more orthostats were identified *in situ*, buried under the road, partially defining the funerary chamber.

This dolmen was used between the end of the 4th and throughout the 3rd millennium BC (~5250-4000 BP), identifying reuses in the Bronze Age and Iron Age.

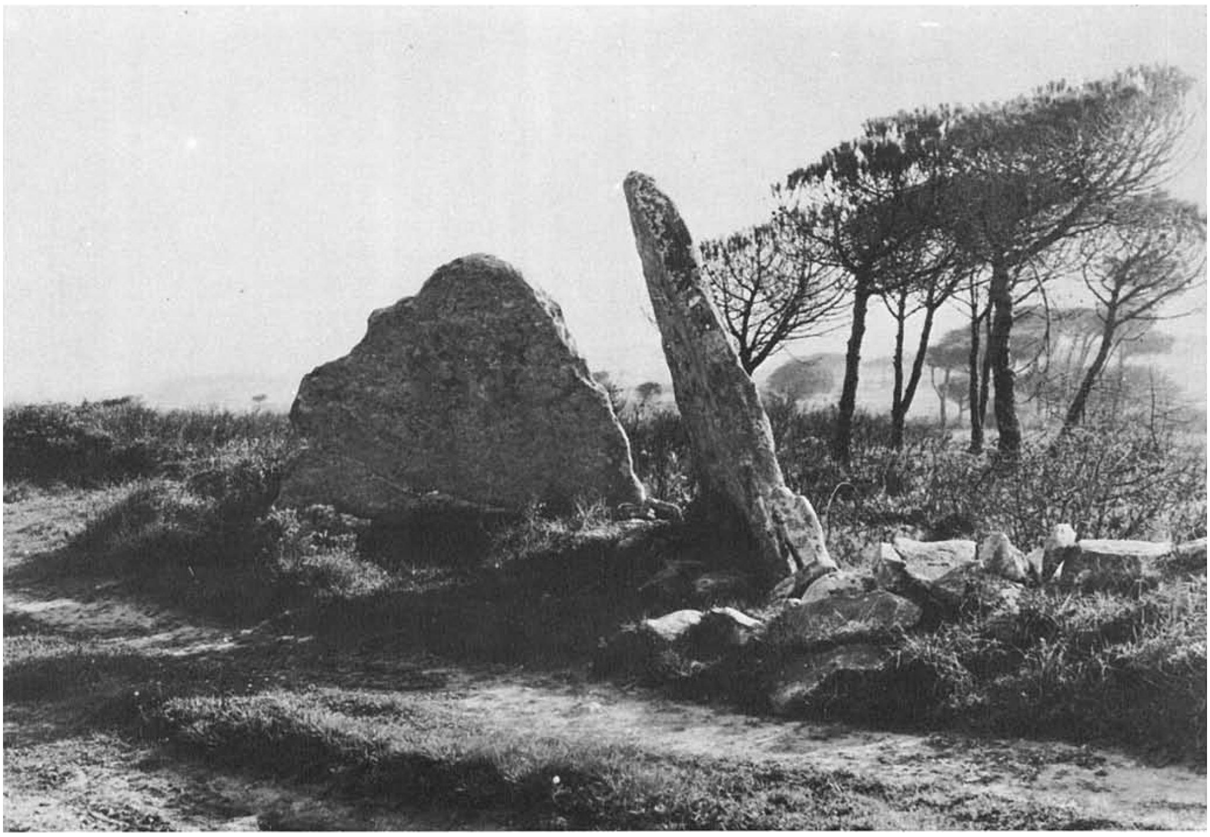

**Supplementary Fig. S3.21. Anta Pedras Granja: Archaeological perspective of the Anta das Pedras da Granja during the 1977 excavation campaign.**

**References:** [396,403–405]

Archive of the bones/materials: Museu Geológico / Museu Arqueológico de São Miguel de Odrinhas - Câmara Municipal de Sintra

##### 3.5.3 Samarra

Description of architecture: This site is near the mouth of the river of Samarra, on the northern cliff of the beach. The funerary context was identified on a natural plateau located near the top of the aforementioned slope. No preserved stratigraphy was identified at this site, but only secondary deposits with archaeological and human osteological remains. The authors interpreted this as a result of the material having slipped from the top of the slope. They suggested that it, originally, could have been a rock shelter adapted for funerary use, or a burial monument of a different kind, but it shouldn't have had megalithic characteristics as only numerous small limestone slabs were identified in this area, with the exception of the large slab lifted by the workers when they discovered the tomb.

Brief description of research history: In July 1948, some workers looking for stone to build a holiday home moved a large slab, under which they identified many bones, ceramics and some polished stone tools. Immediately, they began the uncontrolled excavation of this deposit. O. da Veiga Ferreira and J. Camarate França were both warned about the situation and visited the site in the company of Prof. Mendes Correia. Until November 1948, the first two researchers excavated the site, prospected the surrounding area and also excavated the Monte da Pedranta settlement, on the opposite shore of the beach, which would be contemporary with Samarra's tomb.

Brief description of the context: The archaeological material identified during the excavations at Samarra comprised flint tools (arrowheads and blades), splinters and small chips of quartz, one polished stone axe and various pieces of polished limestone, such as cylindrical idols and a hafted adze. Numerous beads made of various rocks, a perforated shell and various elements made of bone such as the idols with narrowed neck, nail-headed pins, a comb, a cylindrical box decorated with engravings and handles for various instruments. The ceramics were very fragmented, which made it difficult to reconstitute their typology, despite all this, some hemispherical and calotte-shaped bowls was identified. No decorated ceramics was found, but only one vessel covered with red engobe and decorated with four grooves near the rim. A significant number of individuals of both sexes and different ages were deposited in this tomb.

The archaeological materials collected indicate that this tomb was used during the 3rd millennium BC (~5000-4000 BP).

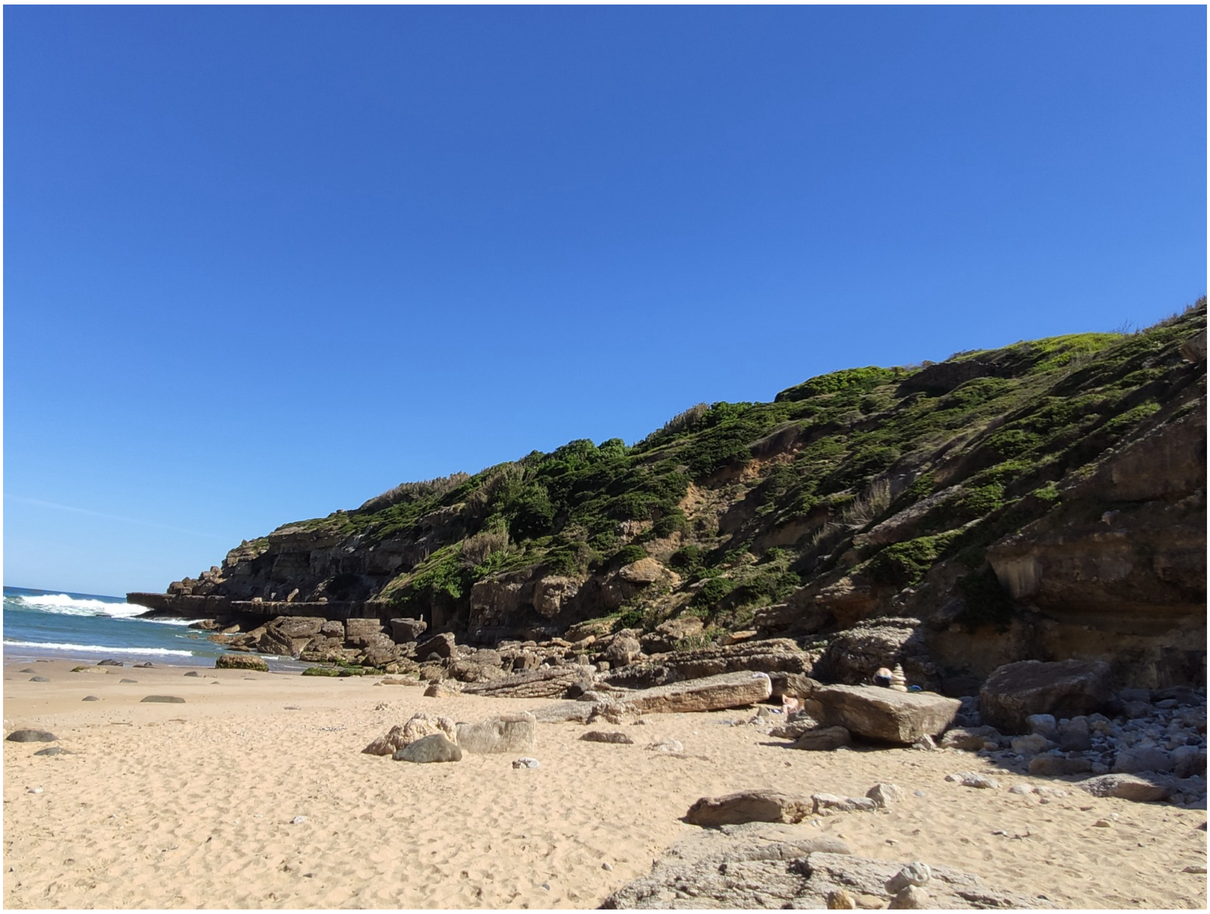

**Supplementary Fig. S3.23. Samarra: mouth of the Samarra stream and the northern coastal cliff, on whose summit the Samarra funerary monument is located.**

**References:** [197,396,401]

Archive of the bones/materials: Museu Geológico / Museu Arqueológico de São Miguel de Odrinhas - Câmara Municipal de Sintra

3.5.4 Bom Santo Cave, Lisbon, Portugal.

**Description of architecture**: This is a burial-cave, located in the Montejunto mountain range, which is topographically structured in three levels: the lower one shows no traces of human presence; the middle level contains five rooms with funerary deposits and a ritual area in Bracelets Room (a built 'altar' with an anthropomorphic stele); and the upper level has four rooms with funerary occupation. A total of 285 $m^2$ was used as a necropolis. Its only entrance opens onto the Tagus valley, at 350 meters a.s.l., in the upper level.

**Brief description of research history**: This cave was discovered in 1993 during speleological surveying of Montejunto carried out by the Associação de Estudos Subterrâneos e Defesa do Ambiente. The extent of the necropolis and its excellent state of preservation triggered systematic speleo-archaeological survey of the whole cave area and the archaeological excavation of Rooms A and B, near the entrance, in 1993–2001. The study of the human remains and grave goods was carried out in 2012–2014 in the framework of the research project 'The Bom Santo Cave and the Neolithic societies of Portuguese Estremadura, $6^{th-}4^{th}$ millennia BC', based at the University of Algarve (Faro, Portugal) and funded by the Portuguese Foundation for Science and Technology.

**Brief description of the context**: The excavation of Rooms A and B revealed a great number of human remains (8961 individual bones and teeth) and associated grave goods (scarce plain pottery, flint blades and geometrics, polished stone axes and adzes, bone tools, and personal adornments). This assemblage is typical of the beginning of megalithism in Portugal, i.e. the Middle Neolithic. Grave goods and human remains were commingled since funerary practices and rituals implied the recurrent rearrangement of the mortuary space. Only two individuals out of a MNI=72 were found in partial anatomical connection. After osteological analysis it was possible to determine 25 and 26 male and female individuals, respectively. Age-at-death determination revealed 20 non-adults (0-13 years-old).

**Existing C-14 dates and analyses**: Given the commingled nature of the deposit, all analyses were performed on a sample of 35 mandibles (49% of MNI) to avoid repetition of results. Only 15 individuals were radiocarbon dated; however, results consistently established the 3800–3380 BC (5750–5330 BP) time interval, in good accord with the above cultural attribution. $\delta^{13}C/\delta^{15}N$ values indicated a reliance on terrestrial food sources but with a relevant percentage of aquatic (=freshwater) component in diets: 19 out of 35 individuals (54%) show >30% of aquatic foods, likely obtained in the Flandrian paleo-estuary of the Tagus river valley. $^{87}Sr/^{86}Sr$ ratios revealed a large amplitude between its extreme values, with the baseline indicating that 11 out of 35 individuals (31%) are non-local, with values suggesting an origin from the neighboring inner Alentejo province, where one of the major megalithic territories in country are known. A previous batch of 14 mtDNA analyses (nine validated results) showed a marked heterogeneity in haplogroups (H, U5, J, K, T2, HV0) with ancestries going back to the region's last hunter-gatherers (H, U5) and the earliest farmers (K, T2) and admixture events taking place during the Early Neolithic.

Individuals #18 and #23, that provided genetic results within this project, are presently being directly dated by AMS; both must correspond to the time interval referred to above. The individuals description is as follows[406]:

Individual #18 - classed by osteological analysis as an adult female. $\delta^{13}C/\delta^{15}N$ values indicate 23% of aquatic resources in diet ($\delta^{13}C$=10.10‰; $\delta^{15}N$=-19.57‰) whereas $^{87}Sr/^{86}Sr$ ratios (0.710697 ± 0.000011) indicate a local individual.

Individual #23 - classed by osteological analysis as an adult male. $\delta^{13}C/\delta^{15}N$ values indicate 29% of aquatic resources in diet ($\delta^{13}C$=10.62‰; $\delta^{15}N$=-19.46‰) whereas $^{87}Sr/^{86}Sr$ ratios (0.712183 ± 0.000010) indicate a non-local individual.

**References**: [406–408]

**Archive of the bones/materials**: National Museum of Archaeology (Lisbon, Portugal).

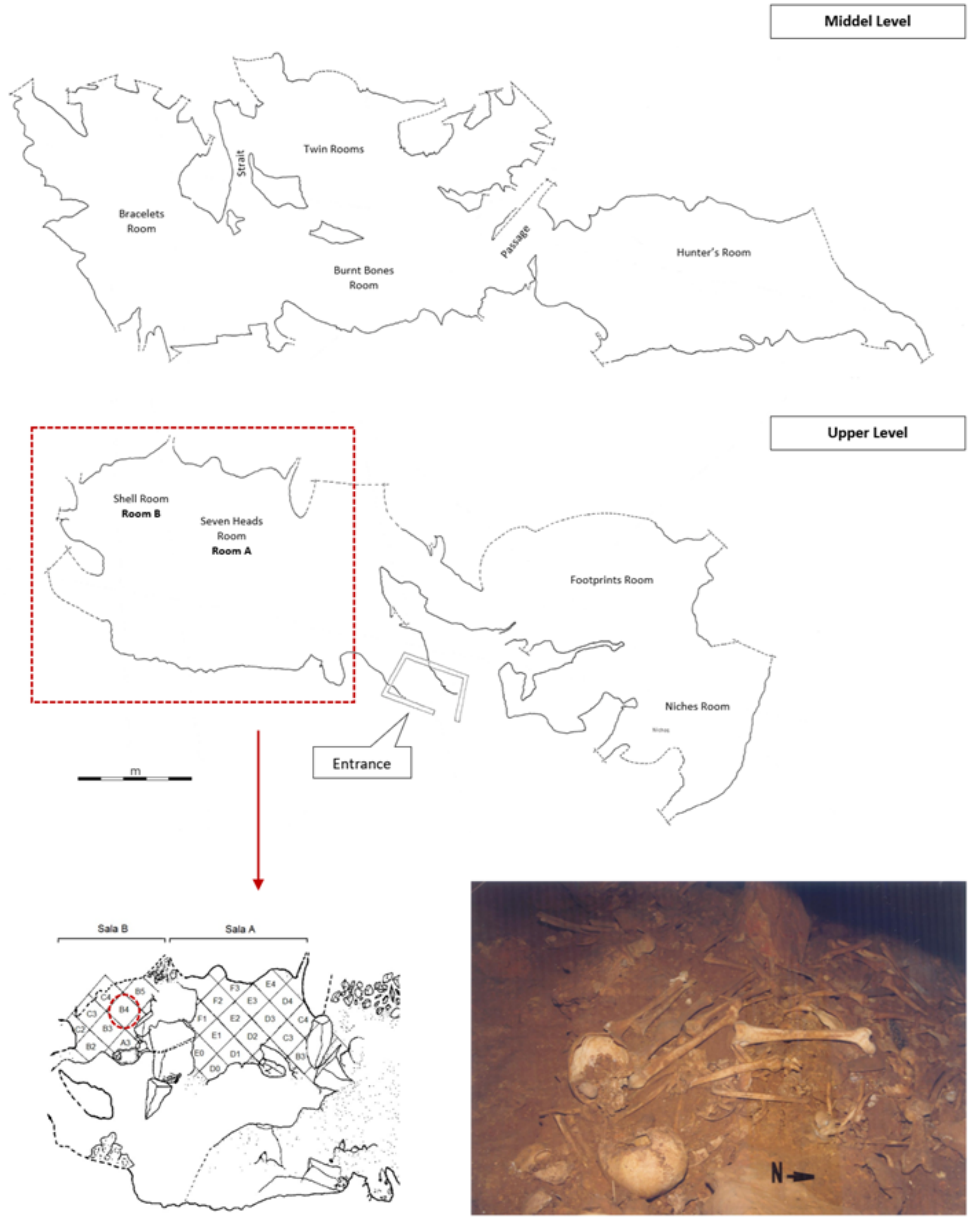

**Supplementary Fig. S3.24. Middle and Upper Levels of Bom Santo Cave with funerary rooms (above) and excavation areas in Rooms A and B with photo of the human remains during their exhumation in square unit B4 (below).** Photo courtesy of António Faustino Carvalho.

##### 3.5.5 Trigache 2, Lisbon, Portugal

**Description of architecture:**

The megalithic necropolis of Trigache is located in the Sintra region of western Portugal. Trigache 2 is situated approximately 400 m northwest of Trigache 1 on the slightly elevated Casinhola plateau. The monument comprises a partially preserved earthen mound (*mamoa*) containing a passage grave with a southeast-facing corridor leading to a circular chamber originally estimated to have measured c. 5.5 m in diameter. Only two orthostats (a and b) remain identifiable, of which orthostat a may have belonged to the corridor. The passage is approximately 2.1 m long, widening from 0.9 to 1.3 m, and is constructed of small orthostats that do not appear to have been broken.

**Brief description of research history**:

The site was first identified by F. C. Ribeiro and later excavated and documented by Vera Leisner and O. V. Ferreira in 1961. The assemblages from Trigache 2 and the neighbouring tomb 3 appear to have been partially mixed, complicating the interpretation of the burial contexts. Archival research based on the Leisner documentation subsequently clarified the attribution of artefacts and the architectural characteristics of the monument[197] (Boaventura, 2009).

**Brief description of the context**:

The archaeological deposits were concentrated within the chamber and corridor. The osteological assemblage consisted of highly fragmented human bones and teeth, including a concentration of cranial fragments adjacent to one of the orthostats.

The lithic assemblage comprises elongated flakes, blades, geometric microliths, predominantly trapezes, and numerous arrowheads, mainly with convex bases. Polished stone tools, including adzes and axes, were recovered from both the chamber and the vestibule. Bone artefacts include decorated hairpins and possible combs, while stone ornaments comprise an anthropomorphic idol plaque and fragments of a decorated, perforated lunula. The ceramic assemblage includes Bell Beaker vessels with incised and impressed decoration, indicating a secondary phase of use. Overall, the finds suggest primary funerary use during the late fourth and early third millennium BC (5500-4500 BP), followed by a reuse during the Bell Beaker period.

**Existing C-14 dates and analyses:**

An initial radiocarbon dating attempt on an occipital bone sample failed due to insufficient collagen. A second dating on a cranial fragment (Beta-239755) yielded an interval of 3090–2890 cal BC/5040–4840 cal BP (with an 89.8% probability for 3030–2890 cal BC/4980–4840 cal BP), corroborating the suggested use of the site during the late 4th to early 3rd millennium BC (5500-4500 BP).

**References:** [197,396,409,410]

**Archive of the bones/materials:** Museo Geologico Lisbon, Portugal).

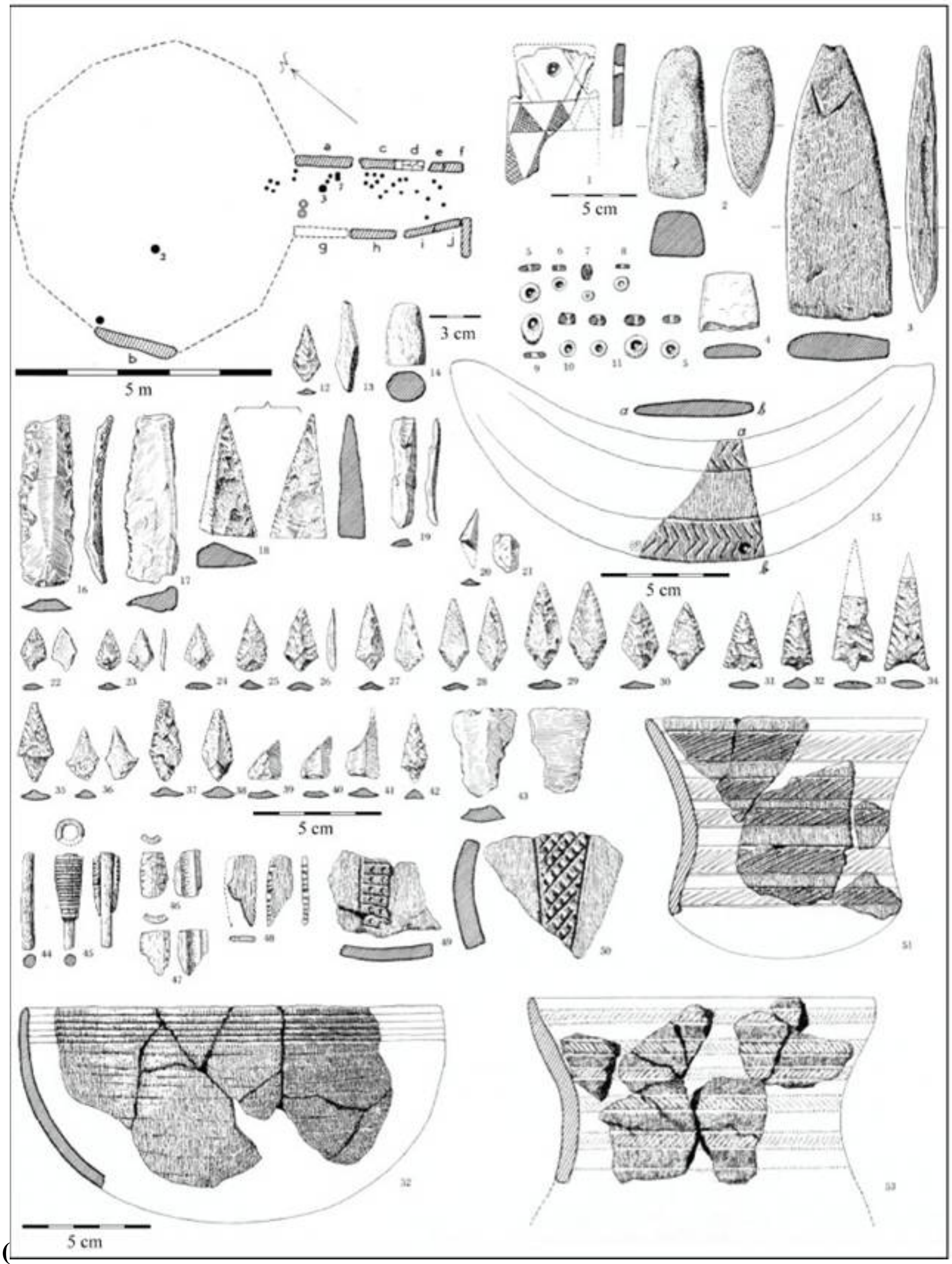

**Supplementary Fig. S3.25. Material from the megalithic grave Trigache 2 (Leisner & Leisner 1965).** Reproduced with kind permission of the Madrid Department of the German Archaeological Institute.

##### 3.5.6 Trigache 4, Lisbon, Portugal

***Description of Architecture:***

Trigache 4 is located approximately 400 m northwest of Trigache 1 on the Casinhola plateau in the Sintra region of western Portugal. The monument consists of a chamber and a corridor. The chamber, partially cut into the Cenomanian bedrock, originally measured approximately 5 m in diameter and preserves two inward-leaning orthostats. The corridor is approximately 2.5 m long and 1.2 m wide. No capstone is preserved, suggesting collapse of the monument. A natural rock formation divides the chamber into two unequal sections, and a possible posthole with stone packing has been identified, although its function remains uncertain.

***Brief Description of Research History:***

The monument was first documented by F. C. Ribeiro and subsequently investigated by Vera Leisner and O. V. Ferreira, who published detailed descriptions during the late 1950s and early 1960s. Although some aspects of the monument's original architecture, particularly the proposed dome-like structure, remain debated, subsequent archival research based on Ribeiro's field documentation and the Leisner archive has clarified several architectural features.

***Brief Description of the Context:***

The burial chamber contained disturbed human remains, with the best-preserved concentration recovered adjacent to orthostat b. Two crania (MG179.24 and MG179.25) were associated with two flint blades at a depth of 0.24 m. Additional finds included a dagger (MG179.03) and a modern whistle (MG179.90) from higher levels. The corridor yielded fragmented human remains together with a horse phalanx (MG179.02) bearing traces of red pigment, fragments of decorated bone hairpins, tubular bone artefacts, and chipped stone tools. No polished stone tools or ceramics were recovered. The assemblage is consistent with a Late Neolithic funerary context.

***Existing C-14 Dates and Analyses***:

A radiocarbon determination obtained from cranium MG179.24 yielded a calibrated age of 3340–2930 cal BC/5290–4880 cal BP (95.4%), confirming use of the monument during the late fourth and early third millennium BC (5500-4500 BP).

**References:** [197,396,409,410]

***Archive of the bones/materials:*** Museo Geologico Lisbon, Portugal).

***Photos/plans***

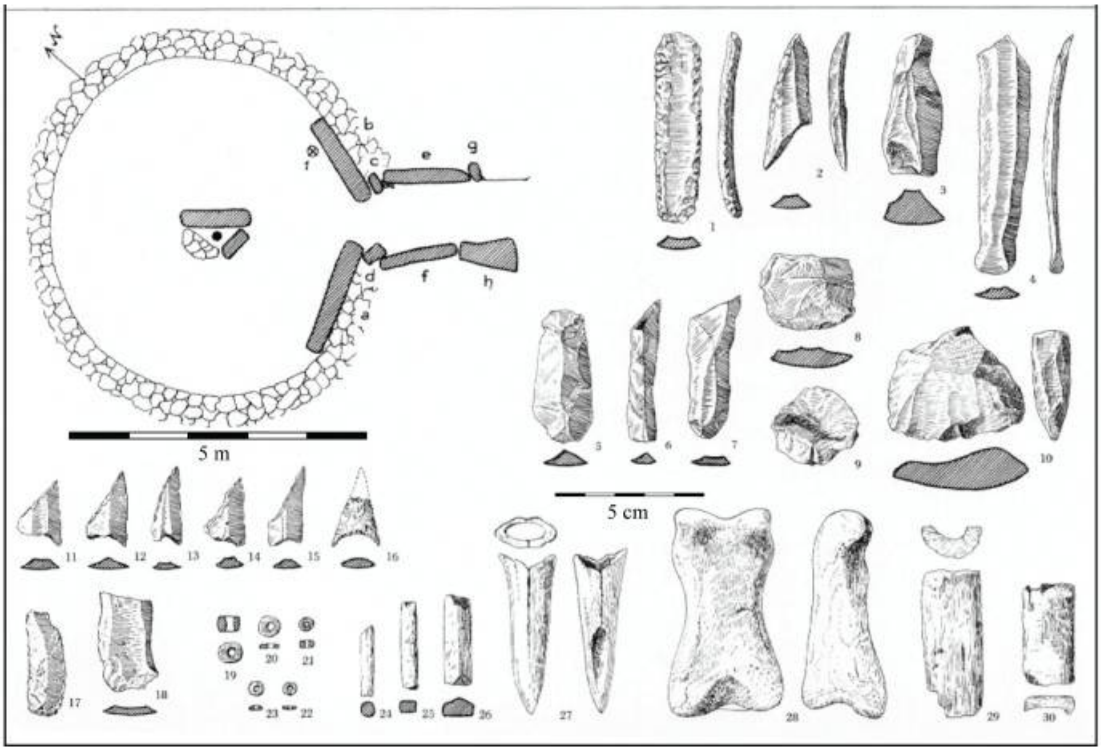

**Supplementary Fig. S3.26. Material from the megalithic grave Trigache 4 (Leisner & Leisner 1965).** Reproduced with kind permission of the Madrid Department of the German Archaeological Institute.

##### 3.5.7 Monte Abraão, Lisbon, Portugal

***Description of Architecture:***

Monte Abraão is located in the municipality of Sintra, near Lisbon, on a gently sloping plateau overlooking the Ribeira das Jardas valley. The monument consists of a single-chambered dolmen with a corridor. The chamber, constructed from large orthostats partially embedded in the bedrock, is approximately 5 m in diameter and was originally covered by a capstone, which is no longer preserved. The corridor measures approximately 2.5 m in length and 1 m in width.

***Brief Description of Research History:***

The monument was first described by Francisco Carlos Ribeiro and subsequently investigated by Vera Leisner and Octávio da Veiga Ferreira during the early 1960s. Their excavations documented the surviving architecture and recorded the extensive disturbance of the monument prior to excavation.

***Brief Description of the Context:***

The osteological assemblage consists of highly fragmented and disarticulated human remains, including cranial fragments and long bones. No articulated burials were preserved.

***Existing C-14 Dates and Analyses:***

A radiocarbon determination on a cranial fragment (MG179.24.01) yielded a calibrated age of 3090–2890 cal BC/5040–4840 cal BP (95.4%), confirming the use of the monument during the late fourth and early third millennium BC (5500-4500 BP).

**References:** [197,396,411]

***Archive of the bones/materials:*** Museo Geologico Lisbon, Portugal).

***Photos/plans***

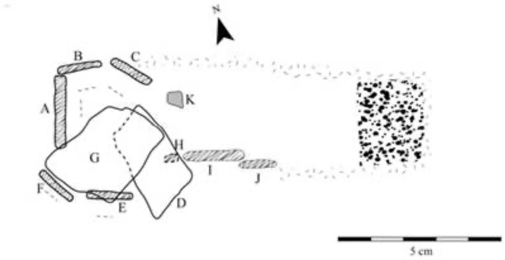

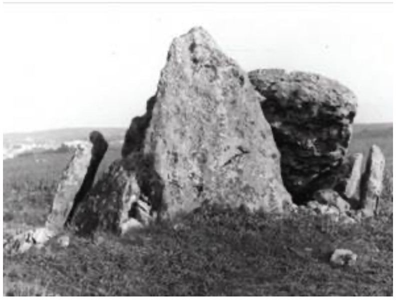

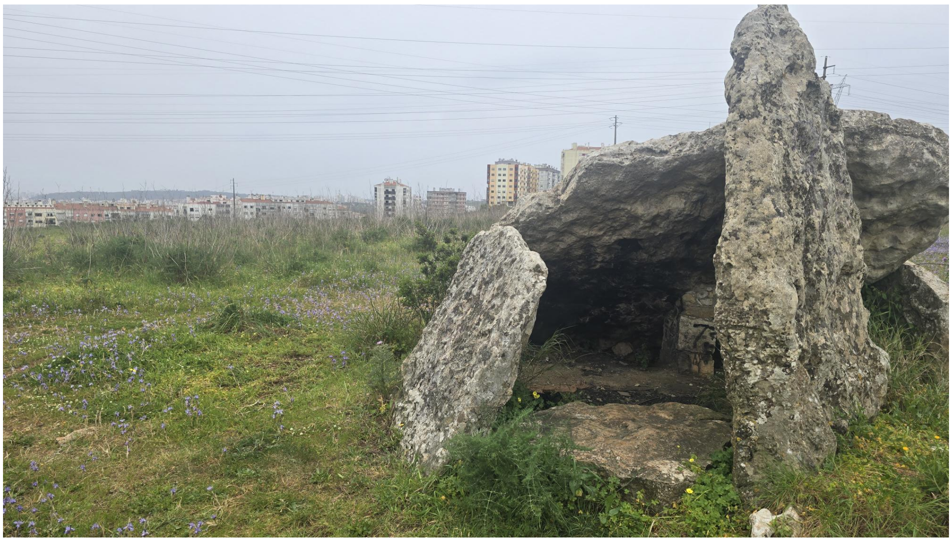

**Supplementary Fig. S3.27. The megalithic grave Monte Abraão**. Photo courtesy of Bettina Schulz Paulsson

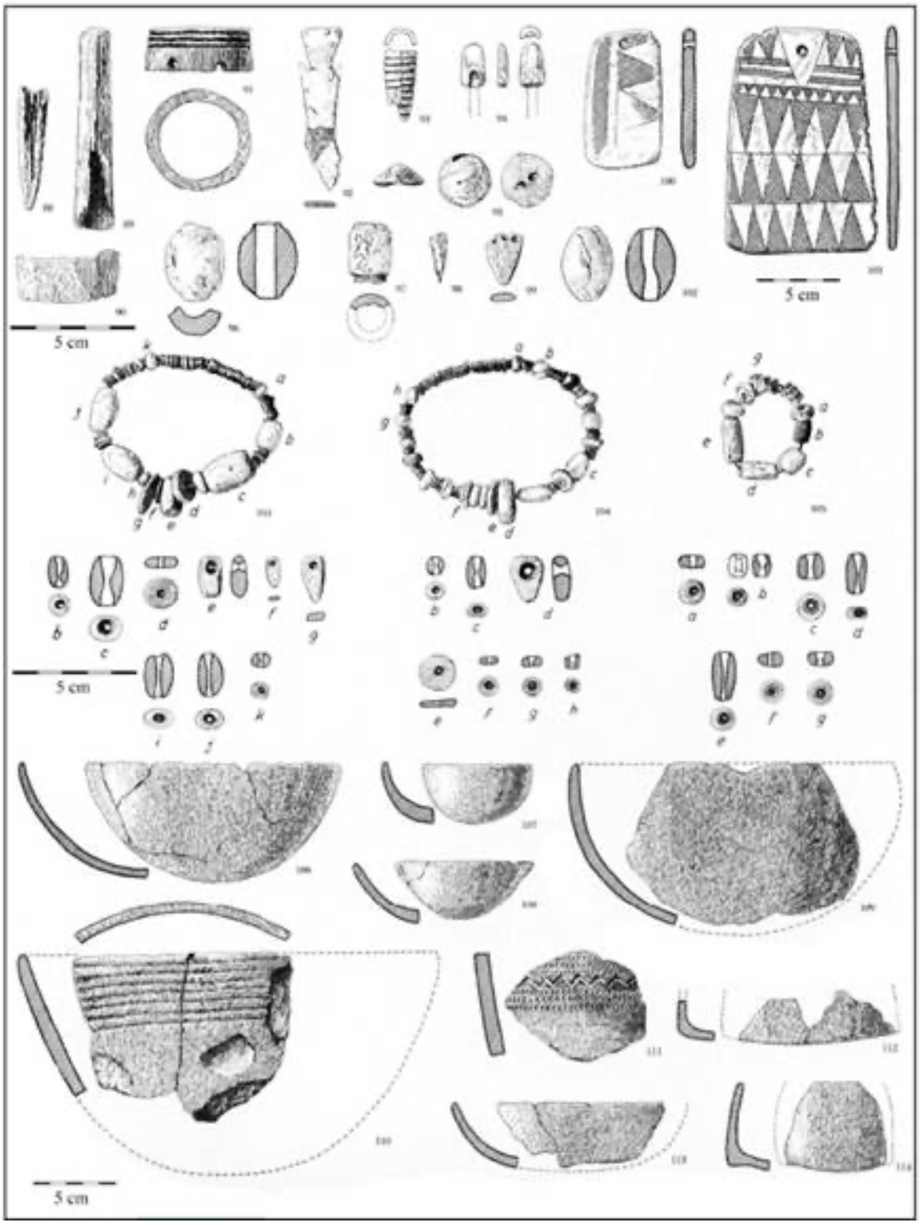

**Supplementary Fig. S3.28. Material from the megalithic grave Monte Abraão, Lisbon, Portugal (Leisner & Leisner 1965)**. Reproduced with kind permission of the Madrid Department of the German Archaeological Institute.

##### 3.5.8 Cova da Onça (Tagus Valley, Portugal)

**Site Context and History**

Cova da Onça, situated between 2 to 8 meters above sea level on the right bank of the Magos in the Tagus Valley, holds significant historical importance as the first shell midden identified in Portugal, discovered in April 1863[412] (Ribeiro 1884). The site was located near the Cabeço dos Ossos midden, but both locations have since been destroyed by recent agricultural activity. The human remains are curated at the Museu Geológico (MG) in Lisbon. The collection represents a Minimum Number of Individuals (MNI) of 32 (Cunha and Cardoso 2003). The skeletal material is curated in a disarticulated state, with individuals mixed in containers lacking specific stratigraphic information or unique excavation identifiers[413–416].

**Chronology and Research Significance.** Recent Bayesian modeling has refined the site's chronology, confirming a Late Mesolithic burial sequence[414]. The primary period of burial activity is estimated to have occurred between approximately 7600 to 7350 cal BP, with an earlier phase dated between 7920 to 7617 cal BP, coinciding with highly favorable estuarine conditions. Crucially, the latest burial activity at the site, estimated between 7488 to 7258 cal BP, is coeval with the earliest Neolithic phases in Portugal. This temporal overlap, combined with early previous studies showing a consistently high marine protein signature (with δ13C values from -16.1 to -14.9‰ and δ15N from 11.9 to 13.6‰)[414], makes Cova da Onça a vital case study for the NEOSEA project in investigating the genomic and dietary transitions between the last hunter-gatherers and the coastal megalithic farming communities in the Portugal.

**References:** [412–416]

#### 3.6. Radiocarbon Dates for the newly obtained genetic results

*Bente Philippsen & Bettina Schulz Paulsson*

For all 46 samples from Sardinia and Portugal that yielded new genetic data in this study, we generated AMS radiocarbon dates and stable isotope measurements.

##### 3.6.1 AMS dating of human bones

Radiocarbon dating and stable isotope measurements ($\delta^{13}C$, $\delta^{15}N$) were performed at the National Laboratory for Age Determination, University Museum, Norwegian University of Science and Technology. Surface contamination was removed from the bones using a scalpel. The bone pieces were comminuted to fragments of about 1-3 mm in diameter, to increase the surface area and allow for better access for the chemicals, while minimizing the damaging effect on the collagen fibrils that e.g. drilling or pulverizing can have. The collagen was extracted from the bone samples using a modified Longin procedure[417,418]. For less well-preserved samples, the method was modified further, employing weaker reagents and colder temperatures, to minimise damage and to maximise collagen yield. The extracted collagen was freeze-dried and split up for radiocarbon dating and stable isotope analysis. For radiocarbon dating, the samples were combusted, graphitised, and dated on the lab's 1MV AMS system (manufactured by High Voltage Engineering[417]). Dates were calculated and reported as described in Nadeau et al.[419]. Stable isotope measurements were done in triplicate using a Flash 2000 elemental analyser coupled to a Delta V mass spectrometer via a ConFlo IV interface (all delivered by Thermo Scientific). The samples were directly measured against $N_2$ and $CO_2$ reference gases, which are calibrated against the IAEA-600 caffeine standard. Other international and in-house standards were used for quality control. Stable isotope ratios were used to estimate the risk of reservoir effects. The dates were calibrated with the terrestrial calibration curve IntCal20[33], using the programme OxCal v.4.4.4[32].

##### 3.6.2 Results

For the 90 samples in the sample batches described here, in total 94 collagen extractions were attempted (several attempts for a few samples). Four extractions did not yield any collagen (TRa-21177 NEOSEA69, TRa-21185 NEOSEA414, TRa-21186 NEOSEA146, and TRa-21189 NEOSEA149) and were repeated with new samples from the same individuals (TRa-22233 to 22236). The other 90 extractions resulted in collagen yields between 0.04% and 14.85%, with an average collagen yield of 5.29% +/- 3.67%. Six of these were too small to attempt radiocarbon dating; they were the ones with very low collagen yields (0.17%, 0.43%, 0.48%, 0.91%, 1.15%, 6.16%). One of these was repeated (TRa-22237 to replace TRa-21191). The others all fulfilled the quality criteria for collagen, with $C:N_{atomic}$ ratios between 3.06 and 3.52 (average 3.15+/-0.07). Even the sample with the lowest non-zero collagen yield, TRa-22237, yielded enough collagen for a radiocarbon date, thanks to the large starting mass of 1000 mg. Its $C:N_{atomic}$ ratio is with 3.52 comparatively high, but still within the acceptable range for non-ultrafiltered collagen.

3.6.3 The calibrated radiocarbon ages

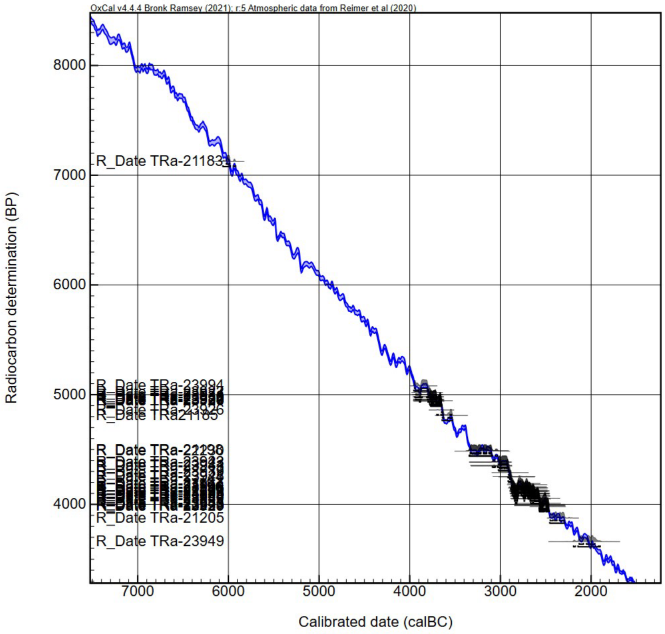

**Supplementary Fig. S3.29. Radiocarbon dates plotted against the calibration curve for samples yielding new genetic data from six megalithic tombs, including tholoi and hypogea, in Portugal and Sardinia.**

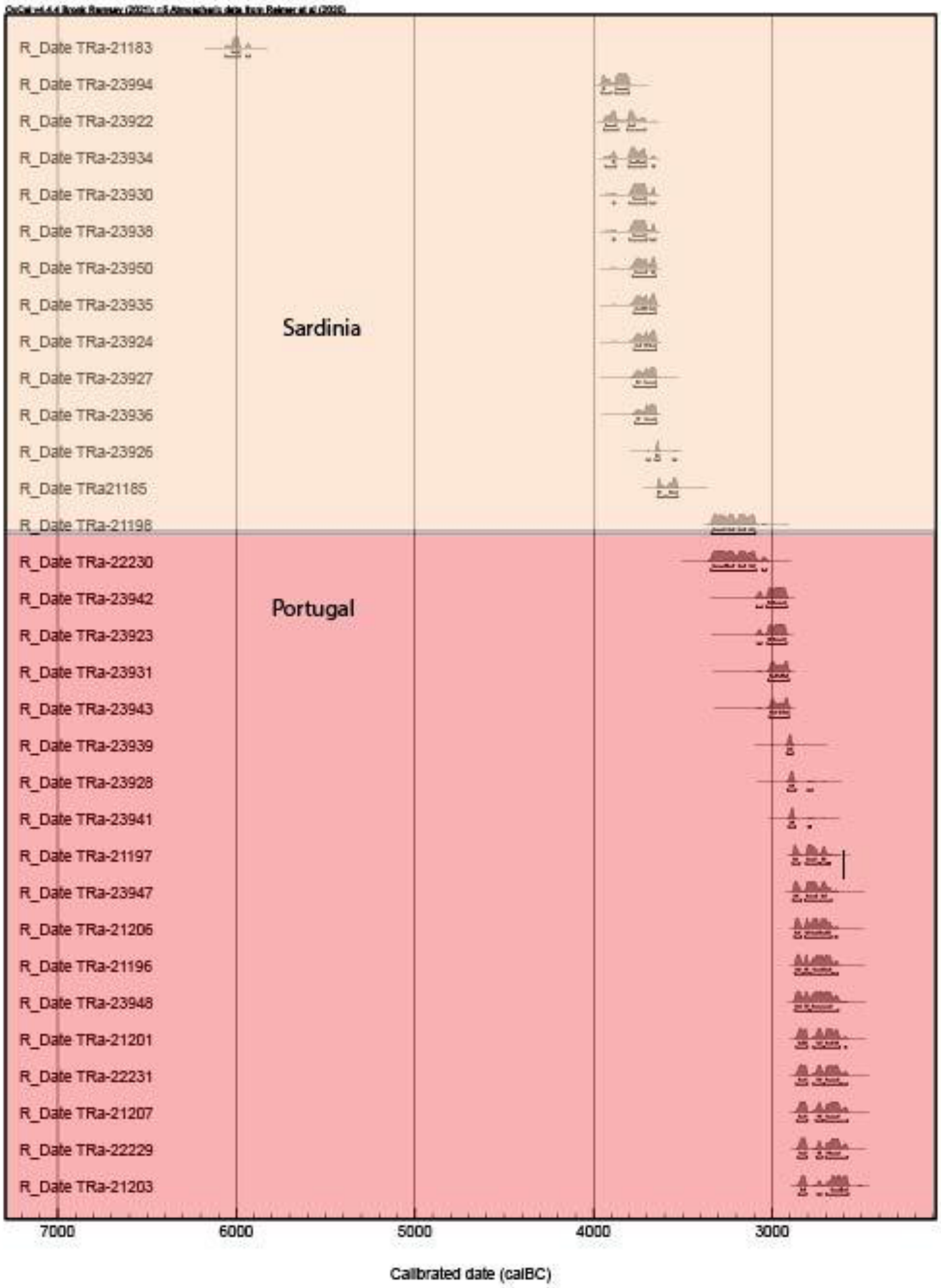

R_Date TRa-21183
R_Date TRa-23994
R_Date TRa-23922
R_Date TRa-23934
R_Date TRa-23930
R_Date TRa-23938
R_Date TRa-23950
R_Date TRa-23935
R_Date TRa-23924
R_Date TRa-23927
R_Date TRa-23936
R_Date TRa-23926
R_Date TRa21185
R_Date TRa-21198
R_Date TRa-22230
R_Date TRa-23942
R_Date TRa-23923
R_Date TRa-23931
R_Date TRa-23943
R_Date TRa-23939
R_Date TRa-23928
R_Date TRa-23941
R_Date TRa-21197
R_Date TRa-23947
R_Date TRa-21206
R_Date TRa-21196
R_Date TRa-23948
R_Date TRa-21201
R_Date TRa-22231
R_Date TRa-21207
R_Date TRa-22229
R_Date TRa-21203
Sardinia
Portugal
7000
6000
5000
4000
3000
Calibrated date (calBC)

**Supplementary Fig. S3.30. Probability Distributions of Radiocarbon Dates for Samples Yielding New Genetic Data from Six Megalithic Tombs, Tholoi, and Hypogea in Portugal and Sardinia.**

| Name<br>Show all<br>Show structure | | Unmodelled (BC/AD) | | | | | | Select<br>All<br>Visible | Page break |
|---|---|---|---|---|---|---|---|---|---|
| | | from | to | % | from | to | % | | |
| R_Date TRa-21183 | | -6024 | -5984 | 68.3 | -6061 | -5923 | 95.4 | 2 | |
| R_Date TRa-23994 | | -3948 | -3809 | 68.3 | -3956 | -3801 | 95.4 | 3 | |
| R_Date TRa-23922 | | -3933 | -3772 | 68.3 | -3944 | -3710 | 95.4 | 4 | |
| R_Date TRa-23934 | | -3891 | -3711 | 68.3 | -3934 | -3660 | 95.4 | 5 | |
| R_Date TRa-23930 | | -3777 | -3710 | 68.3 | -3892 | -3655 | 95.4 | 6 | |
| R_Date TRa-23938 | | -3777 | -3710 | 68.3 | -3892 | -3655 | 95.4 | 7 | |
| R_Date TRa-23950 | | -3768 | -3658 | 68.3 | -3782 | -3653 | 95.4 | 8 | |
| R_Date TRa-23935 | | -3765 | -3656 | 68.3 | -3776 | -3653 | 95.4 | 9 | |
| R_Date TRa-23924 | | -3763 | -3654 | 68.3 | -3774 | -3652 | 95.4 | 10 | |
| R_Date TRa-23927 | | -3760 | -3653 | 68.3 | -3774 | -3650 | 95.4 | 11 | |
| R_Date TRa-23936 | | -3756 | -3652 | 68.3 | -3769 | -3649 | 95.4 | 12 | |
| R_Date TRa-23926 | | -3650 | -3635 | 68.3 | -3702 | -3539 | 95.4 | 13 | |
| R_Date TRa21185 | | -3640 | -3533 | 68.3 | -3644 | -3528 | 95.4 | 14 | |
| R_Date TRa-21198 | | -3331 | -3103 | 68.3 | -3340 | -3096 | 95.4 | 15 | |
| R_Date TRa-22230 | | -3331 | -3100 | 68.3 | -3345 | -3034 | 95.4 | 16 | |
| R_Date TRa-23942 | | -3022 | -2927 | 68.3 | -3089 | -2917 | 95.4 | 17 | |
| R_Date TRa-23923 | | -3020 | -2929 | 68.3 | -3082 | -2919 | 95.4 | 18 | |
| R_Date TRa-23931 | | -3011 | -2913 | 68.3 | -3022 | -2907 | 95.4 | 19 | |
| R_Date TRa-23943 | | -3010 | -2910 | 68.3 | -3017 | -2906 | 95.4 | 20 | |
| R_Date TRa-23939 | | -2911 | -2891 | 68.3 | -2918 | -2885 | 95.4 | 21 | |
| R_Date TRa-23928 | | -2901 | -2881 | 68.3 | -2915 | -2776 | 95.4 | 22 | |
| R_Date TRa-23941 | | -2898 | -2881 | 68.3 | -2909 | -2784 | 95.4 | 23 | |
| R_Date TRa-21197 | | -2880 | -2705 | 68.3 | -2887 | -2676 | 95.4 | 24 | |
| R_Date TRa-23947 | | -2878 | -2701 | 68.3 | -2886 | -2668 | 95.4 | 25 | |
| R_Date TRa-21206 | | -2871 | -2679 | 68.3 | -2876 | -2638 | 95.4 | 26 | |
| R_Date TRa-21196 | | -2866 | -2672 | 68.3 | -2873 | -2632 | 95.4 | 27 | |
| R_Date TRa-23948 | | -2868 | -2670 | 68.3 | -2874 | -2630 | 95.4 | 28 | |
| R_Date TRa-21201 | | -2852 | -2633 | 68.3 | -2866 | -2586 | 95.4 | 29 | |
| R_Date TRa-22231 | | -2850 | -2630 | 68.3 | -2866 | -2581 | 95.4 | 30 | |
| R_Date TRa-21207 | | -2849 | -2627 | 68.3 | -2863 | -2580 | 95.4 | 31 | |
| R_Date TRa-22229 | | -2846 | -2625 | 68.3 | -2856 | -2580 | 95.4 | 32 | |
| R_Date TRa-21203 | | -2839 | -2581 | 68.3 | -2852 | -2574 | 95.4 | 33 | |
| R_Date TRa-23925 | | -2837 | -2581 | 68.3 | -2848 | -2576 | 95.4 | 34 | |
| R_Date TRa-23946 | | -2833 | -2580 | 68.3 | -2847 | -2573 | 95.4 | 35 | |
| R_Date TRa-21209 | | -2659 | -2574 | 68.3 | -2839 | -2500 | 95.4 | 36 | |
| R_Date TRa-23944 | | -2662 | -2503 | 68.3 | -2844 | -2496 | 95.4 | 37 | |
| R_Date TRa-23932 | | -2623 | -2497 | 68.3 | -2834 | -2475 | 95.4 | 38 | |
| R_Date TRa-23940 | | -2621 | -2496 | 68.3 | -2630 | -2476 | 95.4 | 39 | |
| R_Date TRa-23983 | | -2569 | -2476 | 68.3 | -2574 | -2472 | 95.4 | 40 | |
| R_Date TRa-23945 | | -2566 | -2474 | 68.3 | -2572 | -2468 | 95.4 | 41 | |
| R_Date TRa-22228 | | -2562 | -2471 | 68.3 | -2571 | -2466 | 95.4 | 42 | |
| R_Date TRa-23933 | | -2562 | -2471 | 68.3 | -2571 | -2466 | 95.4 | 43 | |
| R_Date TRa-23929 | | -2561 | -2469 | 68.3 | -2571 | -2462 | 95.4 | 44 | |
| R_Date TRa-21205 | | -2451 | -2297 | 68.3 | -2457 | -2293 | 95.4 | 45 | |
| R_Date TRa-23949 | | -2132 | -1959 | 68.3 | -2195 | -1901 | 95.4 | 46 | |

**Supplementary Fig. S3.31. Calibrated ranges (95.4 and 68.2% probability of the newly obtained samples).**

The Sardinian samples from the San Benedetto hypogea form a tight chronological cluster in the first third of the fourth millennium cal BC (3956–3649 cal BC/5906–5599 cal BP, 95.4% probability). In contrast, the Portuguese samples span a broader chronological interval of 3702–1901 cal BC/5652–3851 cal BP (95.4 % probability).

### Supplementary Note S4. Vegetation, sea-level and climatic change in Iberia and Northern France

*Jessie Woodbridge & Ralph Fyfe*

This section provides a synthesis of past vegetation, landscape, sea level and climate changes using palaeoenvironmental evidence from literature to place human migrations and cultural diffusion into the context of regionally variable environmental changes.

#### 4.1 Vegetation

In northern Iberia the periods between 4100–3800 cal. BC (6100–5800 cal. BP) and (4600–4300 cal. BC/6600–6300 cal. BP) were characterised by semi-closed cultural landscapes, diversification of anthropogenic indicators, and local-scale forest clearances, which led to shifts in dominant tree communities[420]. High rainfall and clearance associated with human land use is interpreted as having favoured the spread of beech (*Fagus*) forests ~6400 cal. BP[420]. Agriculture started in the lower altitude regions of northern Iberia 7700–6800 cal. yrs BP and reached the highlands from 6000–4500 cal. yrs BP[421]. Anthropogenic forest clearance for pastoral farming is also described as having involved deliberate use of fire in Northern Iberia[421]. Vegetation was dominated by pine (*Pinus),* birch (*Betula)* and deciduous oak (*Quercus*) (9250–5100 cal. yr BP) with increases in ash (*Fraxinus*) ~6000 cal yr BP. Mixed pine-birch and hazel (*Corylus*) mountain forests persisted for several millennia during the mid-Holocene. The role of climate in shaping the mountain landscapes is suggested to have decreased as human impact increased from ~5100 cal. yr BP[421].

Pollen datasets from northern Iberia show the mid-Holocene (8500–5500 cal. yrs BP) to be characterised by irregular high fire frequency linked with drier and warmer conditions with the period 6000–5000 cal yr BP reaching maximum fire occurrence. These patterns are reflected in various records from the same latitude and are interpreted as indicating regional climatic control[422]. Introduction of agriculture and husbandry started around 7600 cal yr BP in the Pyrenees and during the middle-to-late Neolithic (6500–4300 cal yr BP) grazing activities extended towards high-mountains zones[422]. Pollen records indicate abrupt changes in local vegetation structure between 7250–5500 cal. yrs BP with shifts between *Corylus*, Poaceae and *Betula*. Cereal (Cerealia) pollen, indicating human cultivation, appears ~7100 and 5600 cal yr BP[422]. Between 6000 and 5000 cal yr BP, at the Middle to Late Neolithic transition, fire episodes co-occur with cereal pollen (at 5700, 5550 and 5150 cal. yr BP) indicating a link between forest/woodland clearance and agriculture[422], which is similarly reflected in other records. Cultivation seems to have a large but temporary impact on vegetation during these periods, which is likely reflective of non-permanent agriculture and is also evidenced by increasing cover of tree species favoured by these temporary clearances. Similarly, Zapolska et al.[423] identified high human indicator pollen values between 6200-5700 cal. yrs BP, which correlate with population estimates and suggest high levels of anthropogenic land-cover transformations in western France. Vegetation cover of the mid-Holocene substantially differed from the state of potential natural vegetation due to the cumulative effects of early human alterations of land-cover[423].

In western France, Reille & Andrieu[424] identified the expansion of alder (*Alnus*) around 5810 and 5950 BP, which corresponds with the first appearance of cereal pollen. They describe human pressure as being responsible for a peak in grasses (Poaceae) between 5950 and 5295 cal. yrs BP. In records from the Loire region, Cyprien et al.[425] identified that the Atlantic marine transgression (between 7000 and 5000 cal. yrs BP) caused the level of brackish water to increase between 6740 and 5010 cal. yrs BP. Human activities had an early effect with low intensity deforestation taking place around 6600 cal. yrs BP in addition to the rearing of animals[425]. Cultivated plants were present towards the middle-end of the Neolithic period wheat (*Triticum aestivum*), rye (*Secale cereale*), buckwheat (*Fagopyrum esculentum*), and flax (*Linum usitatissimum*) at about 5600 cal. yrs BP. Cyprien et al.[425] identified tree species replacing one another over time (e.g. *Quercus* (oak) and *Tilia* (linden/lime) taking over from *Corylus* (hazel) and *Fagus* (beech) with marshy areas covered by *Alnus* and *Vitis* (vines), and freshwater and saltwater marshes supporting herbaceous vegetation (Chenopodiaceae (goosefoot) and Poaceae). Sub-halophytic vegetation suggests that brackish waters altered the coastal landscape from 6740 BP to 5000 BP, which occurred at the same time as the marine transgression (7000 to 5000 cal. yrs BP)[425].

Through analysis of multiple pollen sites from the central west coast of France, Joly & Visset[426] revealed that the area has undergone significant geomorphological changes via a succession of marine transgressions and regressions, although this pattern is not necessarily substantiated in other datasets. From the end of the Mesolithic period the woodland ecosystem saw clearance activities followed by cultivation (e.g. cereals and *Juglans* (walnut)) according to these specific records. Joly & Visset[426] describe the beginning of the Atlantic period (7800–5700 cal. yrs BP) as being characterised by rapid transformation of the forest ecosystem (*Quercus, Corylus, Fagus, Carpinu*s and *Ulmus*) marked by abrupt changes (e.g. rapid decline in *Tilia* at 6100 BP). Characteristic oscillations of arboreal taxa are thought to reflect forest clearance while sea level progressively increased across the coastal plain[426]. Shifting relationships were identified between human communities and vegetation revealing a transition from anthropisation of ecosystems to established anthropogenic systems[426]. They identified rapid sedimentation during the Neolithic period (6700–4000 BP/5500–2500 BC), which is interpreted as resulting from climate changes, variations in sea level, and increases in human activity leading to exposure of soils and erosion.

During the Early Neolithic period (6700–5800 BP/5500–4700 BC) they identified that agro-pastoral practices were sporadic, but were rapidly becoming more regular. This was followed by sharp vegetation shifts e.g. transitions in forest composition resulting from clearance followed by reversion to woodland. Vegetation shifts follow charcoal rich layers indicating fires on wetland fringes[426]. Cereals were also observed at the beginning of this period in all pollen sequences indicating widespread cultivation. The increase in anthropogenic pressure occurs shortly after the Flandrian transgression stabilises, therefore a decrease in marine influence and stabilisation of ecological conditions is interpreted as having favoured the spread of agricultural activities[426].

An extension of salt meadows noted in this period played a role in the development of pastoral activities and may have been important for human communities on the fringes of these transition zones[426]. Pastoral activities were also present on the waterway slopes where grasslands extended and were accompanied by presence of ruderals and heathlands[426]. On hillsides, walnut (*Juglans*) was identified around 6000 cal. BC/8000 BP and *Castanea* (chestnut) from 5000 BP, although these patterns are not necessarily captured in other datasets, so cannot be substantiated. During the middle Neolithic (5800–4700 BP/4700–3500 BC), crops become diversified with rye (*Secale*) and buckwheat (*Fagopyrum*) observed around 4300 BC, which are also noted in other studies in the broader region, and are accompanied by periods of land clearance. Rye is not an indigenous species in this region, so its appearance means that it was cultivated[426]. The Flandrian transgression appears to present a rise in sea level leading to agricultural practices possibly being moved further away from the coast due to increasing marine influence, although grazing activities persisted in this coastal area according to the study by Joly and Viesset[426].

#### 4.2 Sea level

Between 6800 and 4200 cal. yrs BP stabilisation of sea level has been identified in northwest Iberia during which freshwater wetlands were reestablished[427]. On the western coast of France, Joly & Visset[426] identify sea level trends as transgressive (rising) initially (6300-6200 BP) and regressive (falling) from 6300-6200 and 5700-5500 cal. yrs BP in this particular dataset. Joly & Visset[426] describe the transgressive phase between 6300–6200 BP as a rapid rise in sea level characterised by the expansion of saltwater mudflats, although patterns differ from other records. The regressive sea-level phases between 6300–6200 and 5700–5500 cal. yrs BP are described as having consequences for the surrounding landscape (expansion or retreat of freshwater marsh land and salt mudflats) and human activity in the area. Changing sea level may have influenced human population and migration patterns in coastal areas during this time.

In a study of Holocene relative sea-level (RSL) change along the north coast of Brittany, Regnauld et al.[428] identified almost continuous sea level rise from 110 m to present position. This trend was interrupted by short periods of negative sea level change around 10000 BP, 4000 BP and 3000 BP and extreme surge events around 2460 BP and 7310 BP[428]. Relatively constant and gradually rising sea level therefore appears to have characterised the period between 6600 BP (4600 BC) and 5800 BP (3800 BC). In a study of RSL change in the northwest coast of Brittany, Goslin et al.[429] estimated sea level to have risen continuously over the last 8000 years with a major inflection around 6000 BP. They identified large vertical discrepancies between the RSL records of Brittany and southwest England and inferred that the two regions underwent a very different pattern and amplitude of subsidence during the last 8000 years. They reveal rapid rise between 8000 and 6000 BP, followed by a continuous 6.5 m RSL rise since 7000 BP with a marked slowdown in RSL rise around 6000 BP[429]. In a RSL database for the Atlantic coast of Europe, García-Artola et al.[430] identified that rates of sea level change were highest during the early Holocene, and during the mid-to late-Holocene rates decreased over time. They highlight that deglaciation of the British-Irish and Fennoscandian Ice Sheets led to large-scale spatial variability in RSL change.

#### 4.3 Climate

Direct reconstruction of climatic changes in northwest Spain and western France largely draw on isotopic analysis (d18O and d13C) from speleothem records, isotopes from benthic organisms from shallow marine cores, lake levels, and sedimentologically-based reconstructions of storm conditions from coastal locations. Authors also draw on wider European changes. Climatic conditions in the region of northwest Iberia and southwest France are influenced both by the Mediterranean and Atlantic systems, with the strength of influence controlled by geographical location. Explanatory drivers of change typically include the strength of the NAO, the position of the Azores High, and periodic disturbance of Atlantic deep-water circulation on a 1500 year cycle (Bond events).

As outlined in Bernal Wormull et al.[431], the Holocene climate of the western Mediterranean region is broadly divided into three phases: humid conditions from 11700-7000 BP, increased hydroclimatic variability and higher temperatures between 7000-5500 BP, and a decrease in humidity since ~5500 BP associated with the onset of the current Mediterranean-type climate. Records generally agree that the period from ~5500 BP was marked by increased aridity in northwest Spain and southwest France. This is associated with major disturbances in the North Atlantic deep-water circulation. The analysis of a shallow marine core off northern Spain by Pena et al.[432] also suggested reduced terrestrial input to the near-coastal zone, caused by reduced river runoff between 5700-5000 BP, indicative of decreased rainfall and increased evaporation. This is also associated with colder sea surface temperatures identified in de Menocal et al.[433].

In a reconstruction of Holocene precipitation change in Iberia, Ilvonen et al.[434] found that precipitation values were highest from 8000 to 6000 cal. yrs BP and lake level reconstructions show highest levels from 8100 to 4000 cal. BP. The Holocene Thermal Maximum, which occurred at about 6000 years BP (4000 BC), is described as having been characterised by a summer temperature anomaly of +1°C and high precipitation[435] and is linked to northern hemisphere vegetation expansion[436]. According to Davis et al.[437], studies indicate a range of dates for the timing of the mid-Holocene Thermal Maximum between 7000 and 6000 BP. These include dates of between 7900 and 6700 BP from pollen data[438], 6200 BP from chironomids[439], maximum tree-line altitudes at 6300-4500 BP[440], and land ice cover was also at a minimum at 6200 BP[441]. van der Horst et al.[421] describe increasingly oceanic climatic conditions during the Holocene, i.e. higher moisture availability throughout the year caused by decreasing insolation seasonality, which favoured the spread of more shade-tolerant broad-leaved deciduous trees. According to Renssen et al.[442], the main climatic parameter influencing plant distribution and vegetation structure in northern Iberia may have been effective precipitation rather than temperature with vegetation changes interpreted as mainly reflecting changes in moisture availability[443] in addition to human activity and associated changes in fire frequency[444]. During the mid-Holocene, wet summers led to year-round moisture availability resulting in a homogeneous seasonal precipitation regime[445].

After 6000 cal. yrs BP, summer precipitation decreased towards present-day values while winter precipitation increased[445]. As shown by pollen-based temperature reconstructions[446] and glacier studies in the Alps, the mid-Holocene saw increasing summer temperature (Holocene Thermal Maximum) at the continental scale[422,446].

Human behaviours may also be influenced by changes in seasonal patterns in the climate system. Baldini et al.[447] modelled seasonal patterns of rainfall using their isotopic data from a speleothem in northern Spain, noting a key hinge-point at 6700 BP: they argue for a fundamental shift in seasonal patterns of rainfall. Unlike most of the rest of the Holocene, rainfall became strongly biased towards the summer (wetter summers and drier winters). The end of this reversed polarity rainfall seasonality interval coincided with the establishment of Mediterranean conditions in southern Iberia at 5300 BP, and a return to the Holocene "normal" of wetter winters and drier summers.

The western Atlantic seaboard is also susceptible to changes in storminess across the Holocene. Van Vliet-Lanoe et al.[448] have summarised episodes of storm conditions from sedimentological sequences around the Brittany region for the last 6500 years. There are major storm events identified around 5500-5100 BP linked to geomorphic change (episodes of dune building, breaching of coastal marsh systems, construction of storm terraces). These storm events are linked to human responses including abandonment of coastal areas[449,450].

Within a Mediterranean wide synthesis of palaeoclimate records, Finne et al.[451] identified wetter conditions in southern Iberia between 8500 and 6100 BP and oscillating climatic patterns after 6000 BP followed by a trend towards increasing aridity. In contrast, northern Iberia remained wetter during this period. Drier conditions were identified in the Mediterranean from 7900 to 3700 BP with rapid shifts in standardised climate z-scores 6700–

6300 BP, 4500–4300 BP, and 3500–3300 BP. The synthesis revealed a south/east versus north/west Mediterranean hydro-climate dipole throughout the Holocene[451]. They infer that climate in southern Iberia was generally wetter from 9700 to 5700 BP, and from 7100 BP there is a trend toward drier conditions. Northern Iberia is characterised by alternating periods of dry and wet climate from 10000 to 4700 BP. The datasets are interpreted as representing periods of relative stability in the hydro-climate in the Mediterranean followed by more rapid (centennial-scale) change, for example, between 6700–6300 BP. There is also a high degree of spatiotemporal hydro-climate variability during the Holocene, as well as variability within regions. They infer that the hydro-climatic history of northern Iberia seems to be linked to the North Atlantic. From 6000 BP, until approximately 3000 BP, southern Iberia shows a trend from wetter to drier conditions whereas other regions show little variability[451]. A palaeoclimate record from southeast Iberia presented in Fyfe et al.[452] shows that the early Holocene was drier with increasing aridity at 6200 BP followed by a shift to wetter climate ~6000 BP, followed again by a shift to drier conditions ~5800 BP, and generally wetter conditions into the mid-Holocene. The record from northeast Iberia presented by Fyfe et al.[452] shows a drier early Holocene followed by a shift to wetter conditions ~6400 BP.

#### 4.4 Summary

The period of population movement between 4100–3800 cal BC (6100-5800 cal. yrs BP) and 4600–4300 cal BC (6600-6300 cal. yrs BP) was accompanied by wetter and warmer climate perhaps providing opportunity for populations to colonise new areas. This was accompanied by stable sea level during some periods and episodes of sea-level transgression and regression during other periods, which provided new opportunities to colonise areas or led to movement of populations into areas more suited to cultivation (e.g. shifts from sea level rise to sea level fall around 6000 cal. yrs BP on the west coast of France). These time periods are also characterised by increasing evidence of anthropogenic activity in pollen records from northern Iberia and western France with crops becoming more diversified. It may have become easier for people to move within the landscape and grow crops in new areas during the mid-Holocene Thermal Maximum (or mid-Holocene climate optimum). While agricultural activity appears to have moved from coastal areas when sea level changes made land unsuitable for crops, grazing continued in coastal areas. These phases are characterised by evidence of intensifying human land use (i.e. forest clearance, deliberate use of fire, and cultivation) and warmer, wetter climate. There are also differences evident between records from higher and lower altitudes with agricultural indicators and human use of fire occurring later at higher altitudes. Evidence of non-permanent agriculture is also a characteristic of these periods indicated by sporadic appearance of land-use indicators and deliberate use of fire, which created cleared spaces allowing trees, such as birch (*Betula*), to colonise. Oscillations within arboreal taxa assemblages are evident in numerous records, which is thought to relate to the first signs of forest clearance. In western coastal France, a decrease in marine influence and more constant ecological conditions may have optimised the spread of agricultural activities.

| **Region** | **Key events in palaeoenvironmental records** |
|---|---|

| | Vegetation | Climate | Sea level |
|---|---|---|---|
| Western France | Rapid forest-ecosystem transformations and oscillations linked to clearance (7800–5700 cal. yrs BP) | Increasingly oceanic conditions through the Holocene. Mid-Holocene Thermal Maximum with higher summer temperatures and high precipitation around 7000–6000 BP. | Transgressive phase 6300–6200 BP (rapid rise; expansion of saltwater mudflats), then regressive phases 6300–6200 and 5700–5500 BP. Continuous rise with slowdown around 6000 BP inferred from Brittany records. |
| Iberia | North: Shifts in dominant tree communities with local clearances during mid-Holocene. Abrupt local vegetation changes 7250–5500 BP; agricultural signals appear. Northwest: early agriculture advancing to highlands by mid–late Neolithic. | North: Mid-Holocene drier/warmer with high fire frequency; maximum around 6000–5000 BP. Effective precipitation a key control; moisture-availability changes drive vegetation shifts. Northwest: Holocene Thermal Maximum near 6000 BP with warmer summers and high precipitation. | Northwest: freshwater wetlands reestablished during sea-level stabilisation (6800–4200 cal. yrs BP).<br><br>Large spatial variability in RSL due to deglaciation patterns. Atlantic coast: rates highest early Holocene, decreasing mid- to late Holocene. |
| Germany and Central Europe | Shift from dense forests toward more open environments between 6500 and 6000 BP, with increasing signals of human land use | Rapid warming around 6000 BP in southern Scandinavia created more habitable conditions for farming and human settlement | On the North Sea coast of Europe, sea level rose steadily through the Holocene. |

| | | | |
|---|---|---|---|
| Southern Britain | Vegetation shifted from mixed woodland to more human-modified open landscapes during mid-Holocene. Marsh expansion and increasing ecological diversity in coastal lowlands. | ~7200–6700 BP North Atlantic region underwent climate oscillations, and between ~6400–6200 BP, increasing marine influence continued and regional climates stabilised into the mid-Holocene 'optimum'. | Rising postglacial sea levels submerged Doggerland between 8000 and 7000 BP. Newly available coastal areas in Britain as sea-level change stabilised ~7000 BP. |
| Scandinavia | Expanded forest cover and improved agricultural viability around 6000 BP. | Increase in summer temperature at 6,000 cal. yr BP in northern Europe. Climatic warming facilitating adoption of farming and associated clearings. | |
| Mediterranean | Italy: drought-tolerant evergreens (e.g., Quercus ilex) expand from ~6400 BP.<br><br>Sardinia: 7500–5500 BP declining seasonality and frequent fires; after ~5500 BP evergreen oak expands with higher lake levels. | Italy: Humid peak to ~6200 BP (within ~9400–6200 BP wet phase), then aridification strengthens after ~6000 BP<br><br>Sardinia: High seasonality and fire 8100–7500 BP; did not mirror stronger mid-Holocene aridification seen elsewhere. | |

| | | | |
|---|---|---|---|
| Portugal | Shrublands (e.g. Cistaceae/Ericaceae) expand after ~5000 BP. | Distinct arid phases ~7000 and 6400–6200 BP; increasing anthropogenic influence alongside drying after ~5000 BP. | Coastal floodplains stabilise as rise decelerates ~6400–5200 BP. |

**Table 4.1. Summary of key regional trends in vegetation, climate and sea level change.**

### References

1. Schulz Paulsson, B. *Time and Stone: The Emergence and Development of Megaliths and Megalithic Societies in Europe*. (Archaeopress Publishing Ltd, 2017).

2. Müller, J., Kruckenberg, C., Großmann, R. & Luckner, J. A map of European megaliths. *J. Neolit. Archaeology* https://doi.org/10.12766/jna.2023.6 (2023) doi:10.12766/jna.2023.6.

3. Montelius, O. Förbindelse mellan Skandinavien och vestra Europa före Kristi födelse. *Svenska Fornminnesföreningens Tidskrift* **7**, 124–155 (1889).

4. Montelius, O. Orienten och Europa. *Antiqvarisk Tidskrift för Sverige XIII* 1–252 (1905).

5. Kossinna, G. Der Ursprung der Urfinnen und Urindogermanen und ihre Ausbreitung nach dem Osten. *Mannus* **I–II**, 17–52 (1910).

6. Sprockhoff, E. Handbücher der Urgeschichte Deutschlands 3. (1938).

7. Childe, V. G. *The Dawn of European Civilisation*. (Kegan Paul, London and Edinburgh, 1925).

8. Childe, V. G. *The Prehistory of European Society*. (Penguin Books, Harmondsworth, 1958).

9. Childe, V. G. *Prehistoric Migrations in Europe*. (Aschehaug, Oslo, 1950).

10. Childe, V. G. *Prehistoric Communities of the British Isles*. (W. & R. Chambers, London and Edinburgh, 1940).

11. Daniel, G. E. *The Prehistoric Chamber Tombs of England and Wales*. (Cambridge University Press, Cambridge, England, 1960).

12. Daniel, G. *The Megalith Builders of Western Europe*. (Hutchinson, London, 1963).

13. Renfrew, C. Megaliths, Territories and Populations. in *Acculturation and Continuity in Atlantic Europe* (ed. de Laet, S. J.) 45–58 (De Tempel, Brugge, 1976).

14. Renfrew, C. *Before Civilisation: The Radiocarbon Revolution and Prehistoric Europe*. (Penguin, London, 1973).

15. Chapman, R. The emergence of formal disposal areas and the “problem” of megalithic tombs in Europe. in *The Archaeology of Death* (eds. Chapman, R., Kinnes, I. & Randsborg, K.) 71–82 (Cambridge University Press, Cambridge, 1981).

16. Sherratt, A. The genesis of megaliths: Monumentality, ethnicity and social complexity in Neolithic north-west Europe. *World Archaeol.* **22**, 147–167 (1990).

17. Müller, J. *et al.* Zur absolutchronologischen Datierung der europäischen Megalithen. in *Tradition und Innovation* (ed. Festschrift, C.) 63–109 (1998).

18. Müller, J. *et al. Zur Entstehung Der Europäischen Megalithik*. 51–81 (Beier & Beran, Mannheim/Weißbach, 1999).

19. *Neolithic and Copper Age Monuments: Emergence, Function and the Social Construction of the Landscape*. (Archaeopress, Oxford, 2014).

20. Midgley, M. *The Megaliths of Northern Europe*. (Routledge, London & New York, 2008).

21. *Megaliths and Identities: Early Monuments and Neolithic Societies from the Atlantic to the Baltic. 3rd European Megalithic Studies Group Meeting, 13th-15th of May 2010 at Kiel University*. (Habelt, Bonn, 2011).

22. *As Time Goes By? Monumentality, Landscapes and the Temporal Perspective*. (Habelt, Bonn, 2012).

23. Schulz Paulsson, B. Radiocarbon dates and Bayesian modeling support maritime diffusion model for megaliths in Europe. *Proc. Natl. Acad. Sci. U. S. A.* **116**, 3460–3465 (2019).

24. Schulz Paulsson, B. Transmission symbolique et la mer dans l'Europe mégalithique (4700-2500 cal av). *Anthropologie* **130**, (2026).

25. Cassen, S. *et al. CARNAC. Récit Pour Un Imagier*. (Laboratoire de recherche archéologie et architectures / CreAAH, 2021).

26. García Sanjuán, L., Sánchez Díaz, F. & Morell Rovira, B. The "Megalithisation" of Iberia: A spatio-temporal model. *L'Anthropologie* **126**, 102993 (2022).

27. da Silva, N. A. *et al.* Long-distance genetic relatedness in megalithic central Europe. *Science* **392**, 839–844 (2026).

28. Bueno Ramírez, P., de Balbín Behrmann, R. & Barroso Bermejo, R. Megalithic art in the Iberian Peninsula: Thinking about graphic discourses in the European Megaliths. in *Fonctions, utilisations et représentations de l'espace dans les sépultures monumentales du Néolithique européen* 185–203 (Presses universitaires de Provence, Aix-en-Provence, 2016).

29. Cassen, S. *et al.* Real and ideal European maritime transfers along the Atlantic coast during the Neolithic. *Doc. Praehist.* **46**, 308–325 (2019).

30. Scarre, C. M., Verdonkschot, J., Scarre, C., González-García, A. C. & Criado-Boado, F. Time, mobility and society: New approaches to megalithic monumentality in western and northern Europe. in *Megalithic Societies: Old Questions, New Narratives* 1–17 (Archaeopress, Oxford, 2026).

31. Bronk Ramsey, C. Deposition models for chronological records. *Quat. Sci. Rev.* **27**, 42–60 (2008).

32. Bronk Ramsey, C. Bayesian analysis of radiocarbon dates. *Radiocarbon* **51**, 337–360 (2009).

33. Reimer, P. J. *et al.* The IntCal20 Northern hemisphere radiocarbon age calibration curve (0–55 cal kBP). *Radiocarbon* **62**, 725–757 (2020).

34. Hofmanová, Z. *et al.* Early farmers from across Europe directly descended from Neolithic Aegeans. *Proc. Natl. Acad. Sci. U. S. A.* **113**, 6886–6891 (2016).

35. Kılınç, G. M. *et al.* The demographic development of the first farmers in Anatolia. *Curr. Biol.* **26**, 2659–2666 (2016).

36. Lazaridis, I. *et al.* Genomic insights into the origin of farming in the ancient Near East. *Nature* **536**, 419–424 (2016).

37. Lipson, M. *et al.* Parallel palaeogenomic transects reveal complex genetic history of early European farmers. *Nature* **551**, 368–372 (2017).

38. Mathieson, I. *et al.* The genomic history of southeastern Europe. *Nature* **555**, 197–203 (2018).

39. Shennan, S. *Cambridge World Archaeology: The First Farmers of Europe: An Evolutionary Perspective*. (Cambridge University Press, Cambridge, England, 2018).

40. Horejs, B. *et al.* The Aegean in the early 7th millennium BC: Maritime networks and colonization. *J. World Prehist.* **28**, 289–330 (2015).

41. Koptekin, D. *et al.* Out-of-Anatolia: Cultural and genetic interactions during the Neolithic expansion in the Aegean. *Science* **388**, eadr3326 (2025).

42. Brami, M. N. A graphical simulation of the 2,000-year lag in Neolithic occupation between Central Anatolia and the Aegean basin. *Archaeol. Anthropol. Sci.* **7**, 319–327 (2015).

43. Maniatis, Y. & Adaktylou, F. Revenia-Korinos: One of the earliest Neolithic settlements in North Greece as evidenced by radiocarbon dating. *Radiocarbon* **63**, 1025–1051 (2021).

44. Krauß, R., Marinova, E., De Brue, H. & Weninger, B. The rapid spread of early farming from the Aegean into the Balkans via the Sub-Mediterranean-Aegean Vegetation Zone. *Quat. Int.* **496**, 24–41 (2018).

45. Porčić, M. The beginning of the Neolithic in the central Balkans: Knowns and unknowns. *Doc. Praehist.* **51**, 178–193 (2024).

46. Brami, M. *et al.* Was the fishing village of Lepenski Vir built by Europe's first farmers? *J. World Prehist.* **35**, 109–133 (2022).

47. Gelabert, P. *et al.* Social and genetic diversity in first farmers of central Europe. *Nat. Hum. Behav.* **9**, 53–64 (2025).

48. Manning, K. *et al.* Animal exploitation in the early neolithic of the Balkans and central Europe. 237–252 (2013).

49. Orton, D. Herding, settlement, and chronology in the Balkan Neolithic. *Eur. J. Archaeol.* **15**, 5–40 (2012).

50. Conolly, J., Manning, K., Colledge, S., Dobney, K. & Shennan, S. Species distribution modelling of ancient cattle from early Neolithic sites in SW Asia and Europe. *The Holocene* **22**, 997–1010 (2012).

51. Colledge, S., Conolly, J. & Shennan, S. The evolution of Neolithic farming from SW Asian origins to NW European limits. *Eur. J. Archaeol.* **8**, 137–156 (2005).

52. Porčić, M., Blagojević, T., Pendić, J. & Stefanović, S. The Neolithic Demographic Transition in the Central Balkans: population dynamics reconstruction based on new radiocarbon evidence. *Philos. Trans. R. Soc. Lond. B Biol. Sci.* **376**, 20190712 (2021).

53. Porčić, M., Blagojević, T., Pendić, J. & Stefanović, S. The timing and tempo of the Neolithic expansion across the Central Balkans in the light of the new radiocarbon evidence. *J. Archaeol. Sci. Rep.* **33**, 102528 (2020).

54. Blagojević, T., Porčić, M., Penezić, K. & Stefanović, S. Early Neolithic population dynamics in the Eastern Balkans and the Great Hungarian Plain. *Doc. Praehist.* **44**, 18–33 (2017).

55. Weninger, B. Barcode seriation and concepts of Gauge Theory. The 14C-Chronology of Starčevo, LBK, and early Vinča. *Quat. Int.* **560–561**, 20–37 (2020).

56. Oross, K. *et al.* 'It's still the same old story': The current southern Transdanubian approach to the Neolithisation process of central Europe. *Quat. Int.* 154–178 (2020).

57. Linden, M. V. & Silva, F. Dispersals as demographic processes: testing and describing the spread of the Neolithic in the Balkans. *Philos. Trans. R. Soc. Lond. B Biol. Sci.* **376**, 20200231 (2021).

58. Bocquet-Appel, J.-P., Naji, S., Linden, M. V. & Kozlowski, J. K. Detection of diffusion and contact zones of early farming in Europe from the space-time distribution of 14C dates. *J. Archaeol. Sci.* **36**, 807–820 (2009).

59. Ivanova, M., De Cupere, B., Ethier, J. & Marinova, E. Pioneer farming in southeast Europe during the early sixth millennium BC: Climate-related adaptations in the exploitation of plants and animals. *PLoS One* **13**, e0197225 (2018).

60. Bánffy, E. *et al.* The Alsónyék story: towards the history of a persistent place. *Bericht der Römisch-Germanischen Kommission* **94**, 283–318 (2017).

61. Stadler, P. Settlement of the Early Linear Ceramics Culture at Brunn am Gebirge, Wolfholz site. *Doc. Praehist.* **32**, 269–278 (2005).

62. Bánffy, E. *The 6th Millennium BC Boundary in Western Transdanubia and Its Role in the Central European Transition//The Szentgyörgyvölgyi-Pityerdomb Settlement*. (2004).

63. Jakucs, J. *et al.* The beginnings of the earliest LBK in Transdanubia. *Quat. Int.* 246–260 (2016).

64. Bánffy, E. & Höhler-Brockmann, H. Burnt daub talking: The formation of the LBK longhouse (a work hypothesis). *Quat. Int.* **560–561**, 179–196 (2020).

65. Berger, J., Metallinou, G. & Guilaine, J. Vers une révision de la transition méso-néolithique sur le site de Sidari (Corfou, Grèce) : nouvelles données géoarchéologiques et radiocarbone, évaluation des processus post-dépositionnels. 209–228 (2013).

66. Binder, D. Le complexe de la Céramique Imprimée en Méditerranée centrale et nord-occidentale : une synthèse chronoculturelle (VIIe et VIe millénaires AEC). in *Céramiques imprimées de Méditerranée occidentale (VIe millénaire AEC) : données, approches et enjeux nouveaux / Western Mediterranean Impressed Wares (6th millennium BCE): New data, approaches and challenges. Actes de la séance de la Société préhistorique française de Nice (mars 2019)* (eds. Binder, D. & Manen, C.) 27–124 (Paris, 2022).

67. Gomart, L. *et al.* Spiralled patchwork in pottery manufacture and the introduction of farming to Southern Europe. *Antiquity* **91**, 1501–1514 (2017).

68. Orton, D., Gaastra, J. & Linden, M. V. Between the Danube and the deep blue sea: Zooarchaeological meta-analysis reveals variability in the spread and development of neolithic farming across the western Balkans. *Open Quat.* **2**, (2016).

69. Lightfoot, E., Boneva, B., Miracle, P. T., Šlaus, M. & O'Connell, T. C. Exploring the Mesolithic and Neolithic transition in Croatia through isotopic investigations. *Antiquity* **85**, 73–86 (2011).

70. Malone, C. The Italian neolithic: A synthesis of research. *J. World Prehist.* **17**, 235–312 (2003).

71. Fiorentino, G., Orazio, M., Calcagnile, L., Quarta, G. & Agostino, A. Climate changes and human-environment interactions in the Apulia region of southeastern Italy during the Neolithic period. The Holocene. **23**, 1297–1316 (2013).

72. Reed, K. From the field to the hearth: plant remains from Neolithic Croatia (ca. 6000–4000 cal bc). *Veg. Hist. Archaeobot.* **24**, 601–619 (2015).

73. Rowley-Conwy, P., Gourichon, L., Helmer, D. & Vigne, J.-D. Early domestic animals in Italy, Istria, the Tyrrhenian Islands, and southern France. in *The Origins and Spread of Domestic Animals in Southwest Asia and Europe* (eds. Colledge, S., Conolly, J., Dobney, K., Manning, K. & Shennan, S.) 161–194 (Left Coast Press, 2013).

74. Spiteri, C., Muntoni, I. M. & Craig, O. E. Dietary practices at the onset of the neolithic in the western Mediterranean revealed using a combined biomarker and isotopic approach. in *Times of Neolithic Transition along the Western Mediterranean* 253–279 (Springer International Publishing, Cham, 2017).

75. Lelli, R. *et al.* Examining dietary variability of the earliest farmers of south-eastern Italy. *Am. J. Phys. Anthropol.* **149**, 380–390 (2012).

76. Salque, M. *et al.* New insights into the Early Neolithic economy and management of animals in Southern and Central Europe revealed using lipid residue analyses of pottery vessels. *Anthropozoologica* **47**, 45–62 (2012).

77. Robb, J. *Cambridge Studies in Archaeology: The Early Mediterranean Village: Agency, Material Culture, and Social Change in Neolithic Italy*. (Cambridge University Press, Cambridge, England, 2007). doi:10.1017/cbo9780511499647.

78. Gomart, L. *et al.* Technical frontiers and social boundaries in the Impressed Ware complex: ceramic manufacture as a proxy of Neolithisation processes in the Western Mediterranean. in *Céramiques imprimées de Méditerranée occidentale (VIe millénaire AEC): données, approches et enjeux nouveaux / Western Mediterranean Impressed Wares (6th millennium BCE): New data, approaches and challenges* (eds. Binder, D. & Manen, C.) 125–146 (Société préhistorique française, Paris, 2022).

79. Fernandes, D. M. *et al.* The spread of steppe and Iranian-related ancestry in the islands of the western Mediterranean. *Nat. Ecol. Evol.* **4**, 334–345 (2020).

80. Yu, H. *et al.* Genomic and dietary discontinuities during the Mesolithic and Neolithic in Sicily. *iScience* **25**, 104244 (2022).

81. Antonio, M. L. *et al.* Ancient Rome: A genetic crossroads of Europe and the Mediterranean. *Science* **366**, 708–714 (2019).

82. Rivollat, M. *et al.* Ancient genome-wide DNA from France highlights the complexity of interactions between Mesolithic hunter-gatherers and Neolithic farmers. *Sci. Adv.* **6**, eaaz5344 (2020).

83. Lugliè, C. Your path led trough the sea … The emergence of Neolithic in Sardinia and Corsica. *Quat. Int.* **470**, 285–300 (2018).

84. Costa, L. J. *Corsica Préhistorique. Peuplement d'une Île et Modes de Vie Des Sociétés Insulaires (IXe-IIe Millénaires Av)*. (Paris, Errance, 2004).

85. De Lanfranchi, F. Sepoltura a inhumazione a Monteleone (Bonifacio). 24–29 (1995).

86. Lugliè, C. Mesolitico e Neolitico. in *Atti della XLIV Riunione Scientifica dell'Istituto Italiano di Preistoria e Protostoria, Cagliari, Barumini, Sassari, 23-28 novembre 2009* 31–47 (Istituto Italiano di Preistoria e Protostoria, Firenze, 2009).

87. Atzeni, E. Notiziario: Su Carroppu di Sirri (Carbonia). *Rivista di Scienze Preistoriche* 478–479 (1972).

88. Usai, L. Il Neolitico medio. in *Atti della XLIV Riunione Scientifica dell'Istituto Italiano di Preistoria e Protostoria, Cagliari, Barumini, Sassari, 23–28 novembre 2009* 49–58 (Istituto italiano di preistoria e protostoria, Firenze, 2009).

89. Melis, M. G. Sardinian megalithic and rock-cut tombs in the context of the prehistoric western Mediterranean. in *Megalithic societies: Old questions, new narratives* (eds. Higginbottom, G. M., Verdonkschot, J., Scarre, C., González-García, A. C. & Criado-Boado, F.) (Archaeopress Archaeology, 2025).

90. Santoni, V. Alle origini dell'ipogeismo in Sardegna: Cabras-Cuccuru S'Arriu, la necropoli del Neolitico medio. in *L'ipogeismo nel Mediterraneo: origini, sviluppo, quadri culturali* (ed. Melis, M. G.) 369–397 (Sassari, 2000).

91. Lugliè, C. & Santoni, V. La necropoli ipogeica di Cuccuru is Arrius (Cabras-Oristano). in *Vasi a Bocca Quadrata. Evoluzione delle conoscenze. Nuovi approcci interpretativi* (ed. Mottes, E.) 487–496 (Trento, 2021).

92. Leandri, F. *et al.* Le site du Monte Revincu (Santo-Pietro-di-Tenda, Haute-Corse): contribution à la connaissance du Néolithique moyen de la Corse. in *Corse et Sardaigne préhistoriques: relations et échanges dans le contexte méditerranéen* (eds. D'Anna, A.,

Cesari, J., Ogel, L. & Vaquer, J.) 165–184 (Imprimerie Nationale Éditions, Bastia, 2007).

93. D'Anna, A. *Étude Des Sites Mégalithiques Corses Dans Leur Contexte Chrono-Culturel*. 38–42 (1998).

94. Tramoni, P., Anna, A., Pinet, L., Guendon, J. & Orsini, J. *Auvergne et Midi. Actes Des Ve Rencontres Méridionales de Préhistoire Récente*. 523–536 (Figari, Corse-du-Sud, 2004).

95. Grosjean, R. & Liégeois, J. Les coffres mégalithiques de la région de Porto-Vecchio. *L'Anthropologie* **68**, 527–548 (1964).

96. Leandri, F. *et al.* Bilan des recherches sur le gisement néolithique moyen du Monte Revincu. in *Archéologie des rivages méditerranéens: 50 ans de recherche* (eds. Delestre, X. & Marchesi, H.) 317–322 (Santo-Pietro-di-Tenda, Haute-Corse; Paris, Errance, 2010).

97. Melis, M. G. The chronology of Monte d'Accoddi (Sardinia, Italy) – new radiocarbon dates. *Interdiscip. Archaeol.* **XIV**, 31–40 (2023).

98. Zilhão, J. Radiocarbon evidence for maritime pioneer colonization at the origins of farming in west Mediterranean Europe. *Proc. Natl. Acad. Sci. U. S. A.* **98**, 14180–14185 (2001).

99. Isern, N., Zilhão, J., Fort, J. & Ammerman, A. J. Modeling the role of voyaging in the coastal spread of the Early Neolithic in the West Mediterranean. *Proc. Natl. Acad. Sci. U. S. A.* **114**, 897–902 (2017).

100. Hamon, C. & Manen, C. The mechanisms of neolithisation of Western Europe: Beyond a south/north approach. *Open Archaeol.* **7**, 718–735 (2021).

101. Manen, C. *et al.* Le sommet de l'iceberg ? Colonisation pionnière et néolithisation de la France méditerranéenne. *Bull. Soc. préhist. fr.* **116**, 317–361 (2019).

102. Ibáñez-Estévez, J. J., Bao, J. F. G., Gassin, B. & Mazzucco, N. Paths and rhythms in the spread of agriculture in the western Mediterranean: The contribution of the analysis of harvesting technology. in *Times of Neolithic Transition along the Western Mediterranean* 339–371 (Springer International Publishing, Cham, 2017).

103. Oms, F. X., Terradas, X., Morell, B. & Gibaja, J. F. Mesolithic-Neolithic transition in the northeast of Iberia: Chronology and socioeconomic dynamics. *Quat. Int.* **470**, 383–397 (2018).

104. Perrin, T., Manen, C., Valdeyron, N. & Guilaine, J. Beyond the sea… The Neolithic transition in the southwest of France. *Quat. Int.* **470**, 318–332 (2018).

105. van Willigen, S. Between Cardial and Linearbandkeramik: From no-man's-land to communication sphere. *Quat. Int.* **470**, 333–352 (2018).

106. Berger, J. Sédiments, dynamique du peuplement et climat au Néolithique ancien. 155–212 (2005).

107. Perrin, T. & Manen, C. Potential interactions between Mesolithic hunter-gatherers and Neolithic farmers in the Western Mediterranean: The geochronological data revisited. *PLoS One* **16**, e0246964 (2021).

108. Arzelier, A. *et al.* Neolithic genomic data from southern France showcase intensified interactions with hunter-gatherer communities. *iScience* **25**, 105387 (2022).

109. Perrin, T., Chambon, P., Gibaja, J. F. & Goude, G. *Le Chasséen, Des Chasséens… Retour Sur Une Culture Nationale et Ses Parallèles*. (Sepulcres de fossa, Cortaillod, Lagozza, 2016).

110. Colas, C. Regard technique sur les poteries du Chasséen entre 4500 et 3500 ans avant J.-C. en Auvergne et dans le Bassin parisien à travers le prisme de la coupe carénée. in *Le Chasséen, des Chasséens… Retour sur une culture nationale et ses parallèles, Sepulcres de fossa, Cortaillod, Lagozza* (eds. Perrin, T., Chambon, P., Gibaja, J. F. & Goude, G.) 123–140 (Archives d'Écologie Préhistorique, Toulouse, 2016).

111. Gernigon, K. Sphère d'interactions, complexe culturel : clefs de lecture de la variabilité géographique des expressions stylistiques du Chasséen. in *Le Chasséen, des Chasséens* (eds. Perrin, T., Chamblandes, J., Gibaja, J. F. & Gourde, G.) 29–42 (Cortaillod, Lagozza; Paris; Toulouse, 2014).

112. Delhon, C., Thiébault, S. & Berger, J.-F. Environment and landscape management during the Middle Neolithic in Southern France: Evidence for agro-sylvo-pastoral systems in the Middle Rhone Valley. *Quat. Int.* **200**, 50–65 (2009).

113. Helmer, D., Gourichon, L., Maamar, H. S. & Vigne, J. L'élevage des caprinés néolithiques dans le sud-est de la France : saisonnalité des abattages, relations entre grottes-bergeries et sites de plein air. *Anthropozoologica* **40**, 167–189 (2005).

114. Bréhard, S., Beeching, A. & Vigne, J.-D. Shepherds, cowherds and site function on middle Neolithic sites of the Rhône valley: An archaeozoological approach to the organization of territories and societies. *J. Anthropol. Archaeol.* **29**, 179–188 (2010).

115. Beeching, A. Habitats pérennes ou précaires au Néolithique. *Tech. Cult.* **56**, 48–61 (2011).

116. Branch, N. P., Black, S., Maggi, R. & Marini, N. A. F. The Neolithisation of Liguria (NW Italy): An environmental archaeological and palaeoenvironmental perspective. *Environ. Archaeol.* **19**, 196–213 (2014).

117. Gandelin, M. *Les Enceintes Chasséennes de Villeneuve-Tolosane et Cugnaux Dans Leur Contexte Du Néolithique Moyen Européen. Archives d'Ecologie Préhistoriques*. (2011).

118. Binder, D., Gratuze, B. & Vaquer, J. La circulation de l'obsidienne dans le sud de la France au Néolithique. *Rubricatum: Revista del Museu de Gavà* **5**, 189–199 (2012).

119. Lemercier, O. La transition du Néolithique moyen au Néolithique final dans le sud-est de la France : Recherches, données et scenarii. in *Quatrième millénaire. La transition du Néolithique moyen au Néolithique final dans le sud-est de la France et les régions voisines* (eds. Lemercier, O., Furestier, R. & Blaise, É.) vol. 27 305–321 (Publications de l'UMR 5140 / ADAL, Lattes, 2010).

120. Lemercier, O. Projet Collectif de Recherche "Nécropoles préhistoriques et protohistoriques de Passy." *Rapport d'activités 2010* (2010).

121. Pétrequin, P. & Weller, O. XVe siècle av. J.-C. : la reprise de la croissance démographique dans le Jura. in *Environnements et cultures à l'Âge du Bronze en Europe*

*occidentale* (eds. Richard, H., Magny, M. & Mordant, C.) 197–210 (Éditions du CTHS, Paris, 2007).

122. Colas, C., Praud, I., Bostyn, F., Cayol, N. & Le Digol, Y. Chasséen septentrional, qui es-tu ? Apports des découvertes récentes dans le nord-ouest de la France. in *Le Chasséen, des chasséens. Retour sur une culture nationale et ses parallèles : Sepulcres de fossa, Cortaillod, Lagozza* (eds. Perrin, T., Chambon, P., Gibaja, J. F. & Goude, G.) 123–140 (Archives d'Ecologie Préhistorique, 2016).

123. Binder, D. Le Chasséen et ses « cultures sœurs » : apports du colloque de 2014. in *Le Chasséen, des Chasséens. Retour sur une culture nationale et ses parallèles, Sepulcres de fossa, Cortaillod, Lagozza* (eds. Perrin, T., Chambon, P., Gibaja Bao, J. F. & Goude, G.) 537–544 (2016).

124. Crombé, P., Perdaen, Y. & Sergant, J. La néolithisation de la Belgique: quelques réflexions. in *Unité et diversité des processus de néolithisation sur la façade atlantique de l'Europe* (eds. Marchand, G. & Tresset, A.) 47–66 (2005).

125. Constantin, C. Fin du rubané, céramique du Limbourg et post-rubané : le néolithique le plus ancien en bassin parisien et en Hainaut. *Oxford: BAR International Series* (1985).

126. Bostyn, F. *Le Néolithique Ancien En Haute-Normandie*. (Maison des Sciences de l'Homme, Paris, 1994).

127. *The First Farmers of Central Europe*. (Oxford University Press, Oxford, 2013).

128. Praud, I. *et al.* The first Early Neolithic settlements in northwest France. *Gall. préhistoire* **58**, 139–144 (2018).

129. Dubouloz, J. *Datation Absolue Du Premier Néolithique Du Bassin Parisien: Complément et Relecture Des Données RRBP et VSG. Bulletin de La Société Préhistorique Française 100*. (2003).

130. Dietsch-Sellami, M.-F. *L'utilisation Des Plantes Sauvages Au Néolithique Dans Le Nord-Ouest de La France: Témoignages Carpologiques*. 127–136 (Lausanne, 2007).

131. Simões, L. G. *et al.* Genomic ancestry and social dynamics of the last hunter-gatherers of Atlantic France. *Proc. Natl. Acad. Sci. U. S. A.* **121**, e2310545121 (2024).

132. Chambon, P. La nécropole de Balloy les Réaudins, approche archéo-anthropologique. in *La Culture de Cerny, Nouvelle économie, nouvelle société au Néolithique* (eds. Constantin, C. & Mordant, D.) (1997).

133. Chambon, P. & Thomas, A. The first monumental cemeteries of Western Europe: the 'Passy type' necropolis in the Paris basin around 4500 BC. *Journal of Neolithic Archaeology* **12**, 1–14 (2011).

134. Chambon, P. *Les Morts Dans Les Sépultures Collectives Néolithiques En France. Du Cadavre Aux Restes Ultimes*. (CNRS Éditions, Paris, 2003).

135. Duhamel, P., Fonton, M. & Carré, H. *La Culture de Cerny, Nouvelle Économie, Nouvelle Société Au Néolithique*. 397–448 (APRAIF, La Culture de Cerny, nouvelle économie, nouvelle société au Néolithique; Nemours, 1997).

136. Lemercier, O. *Projet Collectif de Recherche "Nécropoles Préhistoriques et Protohistoriques de Passy*. (2011).

137. Rivollat, M. *et al.* Ancient DNA gives new insights into a Norman Neolithic monumental cemetery dedicated to male elites. *Proc. Natl. Acad. Sci. U. S. A.* **119**, e2120786119 (2022).

138. Desloges, J. Des architectures monumentales pour les morts: mémoire et pouvoir. in *La Hache et la Meule* (eds. Marcigny, C., Ghesquière, E. & Desloges, J.) 129–131 (Muséum du Havre, Le Havre, 2007).

139. Schulz Paulsson, B. *et al.* A highly precise chronology for the Le Plasker site. in *Le Plasker* (eds. Montassier, N. & Marcoux, B.) (Archeodunum, Sainte Luce sur Loire, 2024).

140. Klassen, L. *Frühes Kupfer Im Norden: Untersuchungen Zu Chronologie, Herkunft Und Bedeutung Der Kupferfunde Der Nordgruppe Der Trichterbecherkultur*. (Aarhus University Press, Aarhus, 2000).

141. Klassen, L. *Jade Und Kupfer. Untersuchungen Zum Neolithisierungsprozess Im Westlichen Ostseeraum Unter Besonderer Berücksichtigung Der Kulturentwicklung Europas 5500-3500 BC*. (Aarhus University Press, Aarhus, 2000).

142. Pétrequin, P. & Pétrequin, A.-M. Du jade pour les dieux. Un système religieux de l'Europe néolithique. Ve et IVe millénaires av. in *Jade for the gods. A religious system in Neolithic Europe. 5th and 4th millennia BC)* (Besançon, 2025).

143. Cassen, S. *et al.* Measuring distinction in the megalithic architecture of the Carnac region: from sign to material. *J. Neolit. Archaeology* https://doi.org/10.12766/jna.2010.45 (2011) doi:10.12766/jna.2010.45.

144. Herbaut, F. Les haches carnacéennes. in *Éléments d'Architecture* (eds. Cassen, S., Boujot, C. & Vaquero, J.) (Éditions Chauvinoises, Chauvigny, 2000).

145. Klassen, L., Pétrequin, P. & Cassen, S. The Power of Attraction. in *Sozialarchäologische Perspektiven* (eds. Hansen, S. & Müller, J.) 13–43 (Habelt, Bonn, 2011).

146. Pétrequin, P., Cassen, S., Croutsch, C. & Weller, O. Haches alpines et haches carnacéennes dans l'Europe du Ve millénaire. *Notae Praehistoricae* **17**, 135–150 (1997).

147. Pétrequin, P., Pétrequin, A.-M., Errera, M., Cassen, S. & Croutsch, C. Complexité technique et valorisation sociale. in *Normes techniques et pratiques sociales* (ed. Astruc, L.) 419–433 (2006).

148. Pétrequin, P. *et al.* Neolithic alpine axeheads, from the continent to great Britain, the isle of man and Ireland. *Analecta Praehistorica Leidensia* **40**, 261–279 (2008).

149. Pétrequin, P., Cassen, S. & Klassen, L. Zwischen Atlantik und Schwarzem Meer. in *Jungsteinzeit im Umbruch* 191–198 (Primus, Darmstadt, 2010).

150. Herbaut, F. & Querré, G. La parure néolithique en variscite dans le sud de l'Armorique. *Bull. Soc. préhist. fr.* **101**, 497–520 (2004).

151. Schulz Paulsson, B. *et al. The Time of Callaïs: Radiocarbon Dates and Bayesian Chronological Modelling*. 479–520 (Archaeopress, Oxford, 2019).

152. Boujot, C. & Cassen, S. Le développement des premières architectures funéraires monumentales en France occidentale. *Revue Archéologique de l'Ouest Supplément* **5**, 195–211 (1992).

153. Cassen, S., Boujot, C. & Vaquero, J. *Éléments d'Architecture: Exploration d'un Tertre Funéraire à Lannec Er Gadouer (Erdeven, Morbihan)*. (Association des Publications Chauvinoises, Chauvigny, 2000).

154. Soler, L. Les gestes funéraires des sépultures en coffre du Néolithique de la Goumoizière. in *Les cistes de Chamblandes et la place des coffres dans les pratiques funéraires du Néolithique moyen occidental* (eds. Moinat, P. & Chambon, P.) 115–133 (Paris/Lausanne, 2007).

155. Blanchard, A., Guyodo, J.-N., Schulz Paulsson, B. & Montassier, F. Le Plasker in Plouharnel (fifth millennium cal BC): a newly discovered section of the megalithic complex of Carnac. *Antiquity* **99**, 915–934 (2025).

156. Cassen, S. *Autour de La Table. Explorations Archéologiques et Discours Savants Sur Des Architectures Néolithiques à Locmariaquer, Morbihan (Table Des Marchands et Grand Menhir)*. (CNRS / Université de Nantes, Nantes, 2009).

157. Rivollat, M. *et al.* Extensive pedigrees reveal the social organization of a Neolithic community. *Nature* **620**, 600–606 (2023).

158. Patterson, N. *et al.* Large-scale migration into Britain during the Middle to Late Bronze Age. *Nature* **601**, 588–594 (2022).

159. Gaillard, F. Fouilles de dolmen de Port-Blanc (Saint-Pierre-Quiberon). *Bulletin et Mémoires de la Société d'Anthropologie de Paris* **VI**, 292–316 (1883).

160. Gouézin, P. *Au Sud Des Landes de Lanvaux (de Guidel à Quiberon)*. (Rennes, 2007).

161. Guilaine, J., Vaquer, J. & Zammit, J. *Grottes Sépulcrales Des Hautes Corbières. Les Grottes de Las Claousos à Auriac et de René Carrié à Termes (Aude). Toulouse: Monographie Des Archives d'Ecologie Préhistorique*. (EHESS, 2015).

162. Arias, P. Neighbours but diverse: social change in north-west Iberia during the transition from the Mesolithic to the Neolithic (5500—4000 cal bc). in *Going Over: The Mesolithic-Neolithic Transition in North-West Europe* 52–71 (British Academy, 2007).

163. Martins, H. *et al.* Radiocarbon dating the beginning of the Neolithic in Iberia: New results, new problems. *J. Mediterr. Archaeol.* **28**, 105–131 (2015).

164. Aubán, J. B., Manen, C. & Pardo-Gordó, S. Spatial and temporal diversity during the neolithic spread in the western Mediterranean: The first pottery productions. in *Times of Neolithic Transition along the Western Mediterranean* (eds. García-Puchol, O. & Salazar-García, D. C.) 373–397 (Springer International Publishing, Cham, 2017).

165. Bernabeu, J., García-Puchol, O. & Orozco, T. New insights relating to the beginning of the Neolithic in eastern Spain: Evaluating empirical data and modelled predictions. *Quat. Int.* **470**, 439–450 (2018).

166. Cámara Manzaneda, J., Clop García, X., García Rosselló, J. & Martín Cólliga, A. Pottery forming of the Cardial and Epicardial Neolithic wares: Analysis and systematisation of technological traces from the ceramic productions of Cova del Frare (NE Iberian Peninsula, 5200-4800 BCE). *J. Archaeol. Sci. Rep.* **43**, 103457 (2022).

167. Mazzucco, N. *et al.* Migration, adaptation, innovation: The spread of Neolithic harvesting technologies in the Mediterranean. *PLoS One* **15**, e0232455 (2020).

168. Oms, F. X. *et al.* Fases de ocupación y estratigrafía del asentamiento neolítico de Les Guixeres de Vilobí (Sant Martí Sarroca, Barcelona). *Trab. Prehist.* **78**, 257–276 (2021).

169. Saña, M. Domestication of animals in the Iberian Peninsula. in *The Origins and Spread of Domestic Animals in Southwest Asia and Europe* (eds. Colledge, S., Conolly, J., Dobney, K., Manning, K. & Shennan, S.) 195–220 (Left Coast Press, Walnut Creek, 2013).

170. Carvalho, A. F. When the Mediterranean met the Atlantic. A socio-economic view on Early Neolithic communities in central-southern Portugal. *Quat. Int.* **470**, 472–484 (2018).

171. Cámara Manzaneda, J. A., Camacho, A. & Barceló Álvarez, J. A. Pottery forming and technical traditions in the Early Neolithic of northeastern Iberia: The case of Cardial and Epicardial wares from Cova del Frare (Barcelona). *Trab. Prehist.* **79**, 234–253 (2022).

172. Oms, F. X. *et al.* The Neolithic in northeast Iberia: Chronocultural phases and $^{14}C$. *Radiocarbon* **58**, 291–309 (2016).

173. Rojo-Guerra, M. A., García-Martínez de Lagrán, Í. & Royo-Guillén, J. I. The beginning of the Neolithic in the mid-Ebro valley and in Iberia's Inland (Northern and Southern submeseta), Spain. *Quat. Int.* **470**, 398–438 (2018).

174. García Sanjuán, L., Wheatley, D. W. & Costa Caramé, M. E. The numerical chronology of the megalithic phenomenon in southern Spain: Progress and problems. in *Exploring Time and Matter in Prehistoric Monuments: Absolute Chronology and Rare Rocks in European Megaliths* (eds. Scarre, C., García Sanjuán, L. & Wheatley, D.) 121–159 (Junta de Andalucía, Antequera, 2011).

175. Góngora, Y. & Martínez, M. *Antigüedades Prehistóricas de Andalucía*. (Universidad de Granada, Granada, 1868).

176. Peyroteo-Stjerna, R. On death in the Mesolithic: Or the mortuary practices of the last Hunter-gatherers of the south-western Iberian peninsula, 7th-6th millennium BCE. *Studies in Archaeology* **20**, (2020).

177. Zilhão, J. New evidence from Galeria da Cisterna (Almonda) and Gruta do Caldeirão on the phasing of central Portugal's early Neolithic. *Open Archaeol.* **7**, 747–764 (2021).

178. Cardoso, J. L., Carvalho, A. F., Rebelo, P., Neto, N. & Simões, C. D. Individual vessels, individual burials? New evidence on early Neolithic funerary practices on the Iberian peninsula's western façade. *Eur. J. Archaeol.* **25**, 331–349 (2022).

179. Carvalho, A. F. *et al.* Hunter-gatherer genetic persistence at the onset of megalithism in western Iberia: New mitochondrial evidence from Mesolithic and Neolithic necropolises in central-southern Portugal. *Quat. Int.* **677–678**, 111–120 (2023).

180. García-Puchol, O. *et al.* A Bayesian approach for timing the neolithization in Mediterranean Iberia. *Radiocarbon* **60**, 181–205 (2018).

181. Peña-Chocarro, L., Zapata, L. & Iriarte, M. J. Neolithic plant economy in the Iberian Peninsula. *Veg. Hist. Archaeobot.* **14**, 330–341 (2005).

182. Soares, P. M. M., Cardoso, R. M., Semedo, Á., Chinita, M. J. & Ranjha, R. Climatology of the Iberia coastal low-level wind jet: weather research forecasting model high-resolution results. *Tellus A* **66**, 22377 (2014).

183. Fuertes-Prieto, M. N. *et al.* An overview of the Mesolithic in the northwest Atlantic and inland area of the Iberian Peninsula. *Quat. Int.* **677–678**, 95–110 (2023).

184. Fábregas Valcarce, R. *et al.* Vaso con decoración cardial de Cova Eirós (Triacastela, Lugo). *Trab. Prehist.* **76**, 147–160 (2019).

185. Prieto-Martínez, M. P. From Galicia to the Iberian Peninsula: Neolithic ceramics and traditions. in *Early Farmers, Late Foragers and Ceramic Traditions: On the Beginning of Pottery in Europe* (ed. Gheorghiu, D.) 94–112 (Cambridge Scholars Publishing, Newcastle upon Tyne, 2009).

186. Suárez Otero, J. Del yacimiento de A Cunchosa al Neolítico en Galicia. Primera aproximación al contexto cultural de la aparición del megalitismo en Galicia. in *O Neolítico Atlántico e as Orixes do Megalitismo* (ed. Rodríguez Casal, A.) 485–506 (Universidade de Santiago de Compostela, Santiago de Compostela, 1997).

187. Lantes-Suárez, O. *et al.* A prehistoric jade axe from Galicia (Northwestern Iberia): Researching its origin. *J. Lithic Stud.* **8**, (2021).

188. Schulz Paulsson, B. Ritual Practice, Symbols, and Cosmology. in *The Oxford Handbook of Scandinavian Archaeology to 1050 CE* (eds. Hedeager, L., Prescott, C. & Kristiansen, K.) (Oxford University Press, Oxford, 2026).

189. Fábregas Valcarce, R. & Vilaseco Vázquez, X. I. El Neolítico y el Megalitismo en Galicia: problemas teórico-metodológicos y estado de la cuestión. in *Muita gente, poucas antas? Origens, espaços e contextos do Megalitismo. Actas do II Colóquio Internacional sobre Megalitismo* (ed. Gonçalves, V. S.) vol. 25 281–305 (Instituto Português de Arqueologia, Lisboa, 2003).

190. Cámara Serrano, J. A. *El Ritual Funerario En La Prehistoria Reciente En El Sur de La Península Ibérica. BAR International Series 913*. (Archaeopress, Oxford, 2001).

191. Cruz, D. J. Cronologia dos monumentos com tumulus do noroeste peninsular e da Beira Alta. *Estudos Pré-Históricos III* 81–119 (1995).

192. Cruz, D. J. Expressões funerárias e cultuais no norte da Beira Alta (V-II milénios a.C.). Estudos Pré-Históricos VI. 149–166 (1998).

193. Jorge, V. O. Arqueologia social dos sepulcros megalíticos atlânticos : conhecimentos e perspectivas actuais. *Revista da Faculdade de Letras, Série II* **6**, 365–443 (1989).

194. Jorge, S. Later prehistoric monuments of Northern Portugal : some remarks. *Journal of iberian archaeology* **0**, 105–114 (1998).

195. Alonso, F. & Bello Diéguez, J. M. Neolítico Atlántico e as Orixes do Megalitismo. Actas do Coloquio Internacional (Santiago de Compostela, 1-6 de abril de 1996). 507–520 (1997).

196. Gallego, A. Balance de las actuaciones arqueológicas llevadas a cabo en la necrópolis megalítica de Cotogrande (Cabral) (Campañas de 1989-1992). in *Castrelos 5-6* 7–28 (1992).

197. Boaventura, R. As antas e o megalitismo da região de Lisboa. (Universidade de Lisboa, Faculdade de Letras, Departamento de História, 2009).

198. Whittle, A. A provisional model for the Mesolithic-Neolithic transition in Britain. in *The early Neolithic in northern Europe: new approaches to migration, movement and*

*social connection* (eds. Hofmann, D., Cummings, M. & Bjornevad-Ahlqvist, R.) 113–124 (Sidestone Press, Leiden, 2025).

199. Whittle, A. The politics of early Neolithic connectivity: Relations between Britain and Ireland. *Proc. Prehist. Soc.* **91**, 115–129 (2025).

200. Brace, S. & Booth, T. The genetics of the inhabitants of Neolithic Britain: a review. in *Ancient DNA and the European Neolithic: Relations and descent* (eds. Whittle, A., Pollard, J. & Greaney, S.) 123–146 (Oxbow Books, 2023).

201. Cassidy, L. Islands apart? Genomics perspectives on the Mesolithic-Neolithic transition in Ireland. in *Ancient DNA and the European Neolithic: Relations and descent* (eds. Whittle, A., Pollard, J. & Greaney, S.) 147–168 (Oxbow Books, 2023).

202. Thomas, J. Neolithization and population replacement in Britain: An alternative view. *Camb. Archaeol. J.* **32**, 507–525 (2022).

203. Pailler, Y. & Sheridan, A. Everything you always wanted to know about… la néolithisation de la Grande-Bretagne et de l'Irlande. Bulletin de la Société préhistorique française 106-1. 25–56 (2009).

204. Bocquet-Appel, J.-P., Naji, S., Linden, M. V. & Kozlowski, J. Understanding the rates of expansion of the farming system in Europe. *J. Archaeol. Sci.* **39**, 531–546 (2012).

205. Brace, S. *et al.* Ancient genomes indicate population replacement in Early Neolithic Britain. *Nat. Ecol. Evol.* **3**, 765–771 (2019).

206. Ariano, B. *et al.* Ancient Maltese genomes and the genetic geography of Neolithic Europe. *Curr. Biol.* **32**, 2668-2680.e6 (2022).

207. Collard, M., Edinborough, K., Shennan, S. & Thomas, M. G. Radiocarbon evidence indicates that migrants introduced farming to Britain. *J. Archaeol. Sci.* **37**, 866–870 (2010).

208. Shennan, S. *et al.* Regional population collapse followed initial agriculture booms in mid-Holocene Europe. *Nat. Commun.* **4**, 2486 (2013).

209. Bevan, A. *et al.* Holocene fluctuations in human population demonstrate repeated links to food production and climate. *Proc. Natl. Acad. Sci. U. S. A.* **114**, E10524–E10531 (2017).

210. Woodbridge, J. *et al.* The impact of the Neolithic agricultural transition in Britain: a comparison of pollen-based land-cover and archaeological 14C date-inferred population change. *J. Archaeol. Sci.* **51**, 216–224 (2014).

211. Whitehouse, N. J. *et al.* Neolithic agriculture on the European western frontier: the boom and bust of early farming in Ireland. *J. Archaeol. Sci.* **51**, 181–205 (2014).

212. Rowley-Conwy, P. *et al.* The earliest farming in Britain: towards a new synthesis. in *Farmers at the Frontier* 401–424 (Oxbow Books, 2020).

213. Woodman, P. & McCarthy, M. Contemplating some awful (ly interesting) vistas: importing cattle and red deer into prehistoric Ireland. *Neolithic settlement in Ireland and western Britain* 31–39 (2003).

214. Schulting, R. On the northwestern fringes: earlier Neolithic subsistence in Britain and Ireland as seen through faunal remains and stable isotopes. *The origins and spread of stock-keeping in the near East and Europe* 313–338 (2013).

215. Tresset, A. French connections II: of cows and men. *Neolithic settlement in Ireland and western Britain* 18–30 (2003).

216. Cramp, L. J. E. *et al.* Immediate replacement of fishing with dairying by the earliest farmers of the Northeast Atlantic archipelagos. *Proc. Biol. Sci.* **281**, 20132372 (2014).

217. Jones, G. & Rowley-Conwy, P. On the importance of cereal cultivation in the British Neolithic. in *The Origins and Spread of Domestic Plants in Southwest Asia and Europe* (eds. Colledge, S. & Conolly, J.) 391–419 (Left Coast Press, Walnut Creek, CA, 2007).

218. McClatchie, M. *et al.* Farming and foraging in Neolithic Ireland: an archaeobotanical perspective. *Antiquity* **90**, 302–318 (2016).

219. Roushannafas, T., Bogaard, A. & Charles, M. Geometric morphometric analysis of Neolithic wheat grains: insights into the early development of free-threshing forms. *Veg. Hist. Archaeobot.* **32**, 597–613 (2023).

220. Stevens, C. J. & Fuller, D. Q. Did Neolithic farming fail? The case for a Bronze Age agricultural revolution in the British Isles. *Antiquity* **86**, 707–722 (2012).

221. Edinborough, K. *et al.* New radiocarbon dates show early neolithic date of flint-mining and stone quarrying in Britain. *Radiocarbon* **62**, 75–105 (2020).

222. Wysocki, M. *et al.* Dates, diet, and dismemberment: Evidence from the Coldrum megalithic monument, Kent. *Proc. Prehist. Soc.* **79**, 61–90 (2013).

223. Bergh, S. & Hensey, R. Unpicking the chronology of Carrowmore: Unpicking the chronology of carrowmore. *Oxf. J. Archaeol.* **32**, 343–366 (2013).

224. Schulting, R. J., Murphy, E., Jones, C. & Warren, G. New dates from the north and a proposed chronology for Irish court tombs. *Proc. R. Ir. Acad. C Archaeol. Celt. Stud. Hist. Linguist. Lit.* **112**, 1–60 (2012).

225. Bayliss, A. *et al.* One thing after another: The date of the Ascott-under-Wychwood long barrow. *Camb. Archaeol. J.* **17**, 29–44 (2007).

226. Bayliss, A., Whittle, A. & Wysocki, M. Talking about my generation: The date of the West Kennet long barrow. *Camb. Archaeol. J.* **17**, 85–101 (2007).

227. Brophy, K. From big houses to cult houses: Early Neolithic timber halls in Scotland. *Proc. Prehist. Soc.* **73**, 75–96 (2007).

228. Whittle, A., Healy, F. & Bayliss, A. *Gathering Time: Dating the Early Neolithic Enclosures of Southern Britain and Ireland*. (Oxbow Books, Oxford, 2011).

229. Smyth, J. Tides of change? The house through the Irish neolithic. in *Tracking the Neolithic House in Europe* 301–327 (Springer New York, New York, NY, 2013).

230. Smyth, J. Breaking ground: an overview of pits and pit-digging in Neolithic Ireland. in *Regional Perspectives on Neolithic Pit Deposition* 13–29 (Oxbow Books, 2011).

231. McLaughlin, T. R. *et al.* The changing face of neolithic and bronze age Ireland: A big data approach to the settlement and burial records. *J. World Prehist.* **29**, 117–153 (2016).

232. Jeunesse, C. Pour une origine occidentale de la culture de Michelsberg ? in *Die Michelsberger Kultur und ihre Randgebiete - Probleme der Entstehung* (eds. Biel, J.,

Schlichtherle, H., Strobel, M. & Zeeb, A.) 29–45 (Hemmenhofen, Konrad Theiss, Stuttgart, 1998).

233. Jeunesse, C. The fifth millennium BC in central Europe. Minor changes, structural continuity: a period of cultural stability. in *Contacts, Boundaries and Innovation in the Fifth Millennium: Exploring Developed Neolithic Societies in Central Europe and Beyond* (eds. Gleser, R. & Hofmann, D.) 105–127 (Sidestone Press, Leiden, 2019).

234. Crombé, P. & Vanmontfort, B. The neolithisation of the Scheldt basin in western Belgium. in *Going Over: The Mesolithic-Neolithic Transition in North-West Europe* 262–285 (British Academy, 2007).

235. Linden, M. V. To tame a land: Archaeological cultures and the spread of the neolithic in western Europe. in *Investigating Archaeological Cultures* 289–319 (Springer New York, New York, NY, 2011).

236. Zimmermann, A., Wendt, K. P., Frank, T. & Hilpert, J. Landscape archaeology in central Europe. *Proc. Prehist. Soc.* **75**, 1–53 (2009).

237. Zimmermann, A., Richter, J., Frank, T. & Wendt, K. P. Landschaftsarchäologie II: Überlegungen zu Prinzipien einer Landschaftsarchäologie. *Bericht der Römisch-Germanischen Kommission* **85**, 37–95 (2004).

238. Colas, C. *et al.* Le monument funéraire Michelsberg ancien de Beaurieux la Plaine. in *Cahier d'archéologie romande* (ed. Besse, M.) 329–334 (Aisne, France; Lausanne, 2007).

239. Beau, A. *et al.* Multi-scale ancient DNA analyses confirm the western origin of Michelsberg farmers and document probable practices of human sacrifice. *PLoS One* **12**, e0179742 (2017).

240. Seidel, U. *Badisches Landesmuseum Karlsruhe (Hrsg.): Jungsteinzeit Im Umbruch. Die "Michelsberger Kultur" Und Mitteleuropa Vor 6000 Jahren. Katalog Zur Ausstellung Im Badischen Landesmuseum Schloss Karlsruhe*. 82–87 (Primus-Verlag, Darmstadt, 2010).

241. Jeunesse, C. & Seidel, U. Die Erdwerke. in *Jungsteinzeit im Umbruch. Die »Michelsberger Kultur« und Mitteleuropa vor 6000 Jahren. Katalog zur Ausstellung im Badischen Landesmuseum Karlsruhe 20.11.2010-15.5.2011* 58–61 (Darmstadt, 2010).

242. Seidel, U. *et al.* Die Zeit der großen Gräben: Modelle zur Chronologie des Michelsberger Fundplatzes von Heilbronn-Klingenberg „Schlossberg", Stadtkreis Heilbronn, Baden-Württemberg. *Praehist. Z.* **91**, (2016).

243. Zimmermann, A. Cultural cycles in Central Europe during the Holocene. *Quat. Int.* **274**, 251–258 (2012).

244. Chenal, F., Perrin, B., Barrand-Emam, H. & Boulestin, B. A farewell to arms: a deposit of human limbs and bodies at Bergheim, France, c. 4000 BC. *Antiquity* **89**, 1313–1330 (2015).

245. Geschwinde, M. & Raetzel-Fabian, D. *EWBSL: Eine Fallstudie Zu Den Jungneolithischen Erdwerken Am Nordrand Der Mittelgebirge (Beiträge Zur Archäologie in Niedersachsen 14)*. (Verlag Marie Leidorf, Rahden/Westf, 2009).

246. Rivollat, M. *et al.* When the waves of European Neolithization met: first paleogenetic evidence from early farmers in the southern Paris Basin. *PLoS One* **10**, e0125521 (2015).

247. Bogaard, A. *Neolithic Farming in Central Europe: An Archaeobotanical Study of Crop Husbandry Practices*. (Routledge, London, England, 2004). doi:10.4324/9780203358009.

248. Schier, W. Extensiver Brandfeldbau und die Ausbreitung der neolithischen Wirtschaftsweise im prähistorischen Mitteleuropa. *Prähistorische Zeitschrift* **84**, 155–175 (2009).

249. Kalis, A. J. Umwelt, Klima und Landnutzung im Jungneolithikum. in *Jungsteinzeit im Umbruch. Die „Michelsberger Kultur" und Mitteleuropa vor 6000 Jahren* (ed. Badisches Landesmuseum Karlsruhe) 37–43 (Primus-Verlag, Darmstadt, 2010).

250. Kreuz, A. *et al.* The Late Neolithic Michelsberg culture – just ramparts and ditches? A supraregional comparison of agricultural and environmental data. *Praehist. Z.* **89**, (2014).

251. Zerl, T. *et al.* Dynamics of early agriculture – multivariate analysis of changes in crop cultivation and farming practices in the Rhineland (Germany) between the 6th and early 4th millennium BCE. *J. Archaeol. Sci.* **183**, 106369 (2025).

252. Antolín, F. *et al.* An archaeobotanical and stable isotope approach to changing agricultural practices in the NW Mediterranean region around 4000 BC. The Holocene. **34**, 239–254 (2024).

253. Jeanty, A. *et al.* Archaeobiological evolution of barley (Hordeum vulgare) over the last eight millennia in the northwestern Mediterranean Basin. *Philos. Trans. R. Soc. Lond. B Biol. Sci.* **380**, 20240194 (2025).

254. Out, W. A. *et al.* Plant use at Funnel Beaker sites: combined macro- and microremains analysis at the Early Neolithic site of Frydenlund, Denmark (ca. 3600 bce). *Veg. Hist. Archaeobot.* https://doi.org/10.1007/s00334-024-01020-9 (2024) doi:10.1007/s00334-024-01020-9.

255. Bollongino, R. *et al.* 2000 years of parallel societies in Stone Age Central Europe. *Science* **342**, 479–481 (2013).

256. Skoglund, P. *et al.* Genomic diversity and admixture differs for Stone-Age Scandinavian foragers and farmers. *Science* **344**, 747–750 (2014).

257. Haak, W. *et al.* Massive migration from the steppe was a source for Indo-European languages in Europe. *Nature* **522**, 207–211 (2015).

258. Mittnik, A. *et al.* The genetic prehistory of the Baltic Sea region. *Nat. Commun.* **9**, 442 (2018).

259. Allentoft, M. E. *et al.* Population genomics of post-glacial western Eurasia. *Nature* **625**, 301–311 (2024).

260. Hofmann, D., Frieman, C. J., Furholt, M., Burmeister, S. & Johannsen, N. N. *Negotiating Migrations: The Archaeology and Politics of Mobility*. (Bloomsbury Academic, London, England, 2024).

261. Höhn, B. *Michelsberger Kultur in Der Wetterau*. (Rudolf Habelt, Bonn, 2002).

262. Mischka, D. *Eine Feinchronologische Studie Zur Besiedlungsgeschichte Anhand von Gräbern. 3 Vols. Frühe Monumentalität Und Soziale Differenzierung 20*. (Schleswig-Holstein; Bonn, 2022).

263. Mischka, D., Roth, G. & Struckmeyer, K. The Dąbki Site in Pomerania and the Neolithisation of the North European Lowlands (c. 5000-3000 cal BC). *Archäologie und Geschichte im Ostseeraum* **8**, 465–478 (2015).

264. Sjögren, K.-G. Neolithisation in Sweden, based on directly dated cereals. in *The Early Neolithic in Northern Europe: New Approaches to Migration, Movement and Social Connection* (eds. Hofmann, D., Cummings, V., Bjørnevad-Ahlqvist, M. & Iversen, R.) 205–232 (Sidestone Press, Leiden, 2025).

265. Torfing, T. D. & Feeser, I. *Northern Networks: An Analysis of Three Neolithic Enclosures from the Jutland Peninsula*. (Bonn, 2021).

266. Torfing, T. A new time: Bayesian models of an Early Neolithic enclosure in North-Western Denmark. *Dan. J. Archaeol.* **4**, 109–124 (2015).

267. Andersen, N. H. *The Sarup Enclosures*. (Jutland Archaeological Society, Aarhus, 1997).

268. Andersen, N. H. Sarup revisited: Archaeological Reality and Realities of Archaeology. *Acta Archaeol.* **89**, 31–62 (2018).

269. Artursson, M., Earle, T. & Brown, J. The construction of monumental landscapes in low-density societies: New evidence from the Early Neolithic of Southern Scandinavia (4000–3300 BC) in comparative perspective (November 5, 2015). *J. Anthropol. Archaeol.* **41**, 1–18 (2016).

270. Andersen, N. H. Bondestenalders rituelle mega-pladser. *Skalk* 6–13 (2022).

271. Schulz Paulsson, B. Scandinavian models: Radiocarbon dates and the origin and spreading of passage graves in Sweden and Denmark. *Radiocarbon* **52**, 1002–1017 (2010).

272. Klassen, L. *Along the Road: Aspects of Causewayed Enclosures in South Scandinavia and Beyond*. (Aarhus University Press, Aarhus, 2014).

273. Whittle, A. *et al.* Building for the Dead: Events, Processes and Changing Worldviews from the Thirty-eighth to the Thirty-fourth Centuries cal. bc in Southern Britain. *Camb. Archaeol. J.* **17**, 123–147 (2007).

274. Rudebeck, E. En tidigneolitisk långhög i Kristineberg. in *Monumentala gravformer i det äldsta bondesamhället* (ed. Larsson, L.) 77–109 (Department of Archaeology and Ancient History, Lund University, Lund, 2002).

275. Sjögren, K.-G. Mortuary Practices, Bodies and Persons in Northern Europe. in *The Oxford Handbook of Neolithic Europe* (eds. Fowler, C., Harding, J. & Hofmann, D.) 1005–1022 (Oxford University Press, Oxford, 2015).

276. *Megaliter, Menhirer Och Bebyggelse Från Tidigneolitikum, Senneolitikum Och Yngre Bronsålder–Yngre Järnålder*. (Sydsvensk Arkeologi, Malmö, 2021).

277. Rønne, P. Høj over høj. *Skalk* 3–8 (1979).

278. Rønne, P. Tidligneolitiske jordgrave og et megalitanlæg ved Horsens. *Antikvariske studier* **2**, 213–214 (1978).

279. Price, T. D. New information on the Stone Age graves at Dragsholm, Denmark. *Acta Archaeol.* **78**, 193–219 (2007).

280. Sjögren, K.-G. & Fischer, A. The chronology of Danish dolmens. Results from 14C dates on human bones. *J. Neolit. Archaeology* https://doi.org/10.12766/jna.2023.1 (2023) doi:10.12766/jna.2023.1.

281. Fritsch, B. *et al.* Dichtezentren und lokale Gruppierungen - Eine Karte zu den Großsteingräbern Mittel- und Nordeuropas. *J. Neolit. Archaeology* https://doi.org/10.12766/jna.2010.56 (2010) doi:10.12766/jna.2010.56.

282. Sjögren, K.-G. Megaliter och samhälle i Västsverige. in *Göteborgs universitet* vol. 27 (Göteborg, 2003).

283. Østmo, E. The northern periphery of the TRB: Graves and ritual deposits in Norway. *Acta Archaeol.* **78**, 111–142 (2007).

284. Andersen, N. H. Kult og ritualer i den ældre bondestenalder. *KUML* 13–58 (2000).

285. Jensen, J. Danmarks Oldtid. Bind 1: Stenalder 13.000-2.000 f. *Kr. København: Gyldendal* (2001).

286. Blank, M., Sjögren, K.-G. & Storå, J. Old bones or early graves? Megalithic burial sequences in southern Sweden based on 14C datings. *Archaeol. Anthropol. Sci.* **12**, (2020).

287. Dehn, T. & Hansen, S. I. Birch bark in Danish passage graves. *J. Dan. Archaeol.* **14**, 23–44 (2006).

288. Mannering, U., Possnert, G., Heinemeier, J. & Gleba, M. Dating Danish textiles and skins from bog finds by means of 14C AMS. *Journal of Archaeological Science* **37**, 261–268 (2010).

289. Nielsen, B. H., Christensen, T. & Frei, K. M. New insights from forgotten bog bodies: The potential of bog skeletons for investigating the phenomenon of deposition of human remains in bogs during prehistory. *Journal of Archaeological Science* **120**, 105166 (2020).

290. Chamberlain, A. T. & Pearson, M. P. *Earthly Remains: History and Science*. (British Museum Press, London, England, 2001).

291. Sjögren, K.-G. *et al.* Early Neolithic human bog finds from Falbygden, western Sweden: New isotopic, osteological and histological investigations. *J. Neolit. Archaeology* https://doi.org/10.12766/jna.2017.4 (2017) doi:10.12766/jna.2017.4.

292. Müller, J. *Separation, Hybridisation, and Networks: Globular Amphora Sedentary Pastoralists ca. 3200-2700 BCE (Scales of Transformation in Prehistoric and Archaic Societies, 17)*. (Sidestone Press, Leiden, 2023).

293. Tassi, F. *et al.* Genome diversity in the Neolithic Globular Amphorae culture and the spread of Indo-European languages. *Proceedings of the Royal Society B: Biological Sciences* **284**, 20171540 (2017).

294. Schroeder, H. *et al.* Unraveling ancestry, kinship, and violence in a Late Neolithic mass grave. *Proc. Natl. Acad. Sci. U. S. A.* **116**, 10705–10710 (2019).

295. Immel, A. *et al.* Genome-wide study of a Neolithic Wartberg grave community reveals distinct HLA variation and hunter-gatherer ancestry. *Commun. Biol.* **4**, 113 (2021).

296. Papac, L. *et al.* Dynamic changes in genomic and social structures in third millennium BCE central Europe. *Sci. Adv.* **7**, eabi6941 (2021).

297. Broodbank, C. *et al.* Oued Beht, Morocco: a complex early farming society in north-west Africa and its implications for western Mediterranean interaction during later prehistory. *Antiquity* **98**, 1199–1218 (2024).

298. Broodbank, C. & Lucarini, G. The dynamics of Mediterranean Africa, ca. 9600-1000 BC: an interpretative synthesis of knowns and unknowns. *J. Mediterr. Archaeol.* **32**, 195–267 (2019).

299. Ballouche, A. & Marinval, P. Premiers indices d'agriculture et de pastoralisme sur la façade nord-ouest de l'Afrique (site de Kaf Taht el-Ghar, Maroc). *Revue d'Archéométrie* **27**, 121–128 (2003).

300. Roberts, C. N. *et al.* Mediterranean landscape change during the Holocene: Synthesis, comparison and regional trends in population, land cover and climate. *The Holocene* **29**, 923–937 (2019).

301. Schuhmacher, T. X. Elefanten und Elfenbein auf der Iberischen Halbinsel und in Nordwestafrika: Interdisziplinäre Studien zu Austauschsystemen im 3. und der ersten Hälfte des 2. *Jts. v. Chr* **16**, (2016).

302. Schuhmacher, T. X., Cardoso, J. L. & Banerjee, A. Sourcing African ivory in Chalcolithic Portugal. *Antiquity* **83**, 983–997 (2009).

303. Bokbot, Y. La civilización del Vaso Campaniforme en Marruecos y la cuestión del sustrato Calcolítico precampaniforme. in *El Campaniforme en la Península Ibérica y su Contexto Europeo* (eds. Rojo Guerra, M. A., Garrido-Pena, R. & García-Martínez de Lagrán, I.) 137–161 (Universidad de Valladolid, Valladolid, 2005).

304. Hilmi, M., Mahamoud, A., Aziz El Agbani, M. & Qninba, A. Biodiversity and ecological values of a part of the lower valley of Oued Beht located between the dams of El Kansera and Ouljet Essoltane. **44**, 7–17 (2022).

305. Camps, G. & Camps-Fabrer, H. *La Nécropole Mégalithique Du Djebel Mazela à Bou Nouara*. (Arts et Métiers Graphiques, Paris, 1964).

306. Camps, G. *Aux Origines de La Berbérie. Monuments et Rites Funéraires Protohistoriques*. (Arts et Métiers Graphiques, Paris, 1961).

307. Camps, G. Allées couvertes (Kabylie). *Encyclopédie berbère* **4**, 531–533 (1986).

308. Sanmartí, J. The megalithic necropolises of the eastern Maghreb. in *Megaliths of the World* 1111–1124 (Archaeopress Publishing Ltd, 2022).

309. Koehler, H. Les civilisations mégalithiques du Maroc, le Menhir de Mesora. *Bull. Soc. préhist. fr.* **29**, 413–420 (1932).

310. Muscat, I. *Megalithism and Monumentality in Prehistoric North Africa (Unpublished Dissertation)*. (University of Malta, 2012).

311. Brooks, N., Clarke, J., Garfi, S. & Pirie, A. The archaeology of Western Sahara: results of environmental and archaeological reconnaissance. *Antiquity* **83**, 918–934 (2009).

312. Fregel, R. *et al.* Ancient genomes from North Africa evidence prehistoric migrations to the Maghreb from both the Levant and Europe. *Proc. Natl. Acad. Sci. U. S. A.* **115**, 6774–6779 (2018).

313. Simões, L. G. *et al.* Northwest African Neolithic initiated by migrants from Iberia and Levant. *Nature* **618**, 550–556 (2023).

314. Olalde, I. *et al.* The genomic history of the Iberian Peninsula over the past 8000 years. *Science* **363**, 1230–1234 (2019).

315. *JADE 2. Objets-Signes et Interprétations Sociales Des Jades Alpins Dans l'Europe Néolithique*. (Presses Universitaires de Franche-Comté, Besançon, 2017).

316. *Jade. Grandes Haches Alpines Du Néolithique Européen. Ve et IVe Millénaires Av. J.-C.* (Presses Universitaires de Franche-Comté / Centre de Recherche Archéologique de la Vallée de l'Ain, Besançon / Gray, 2012).

317. Pétrequin, P. *et al.* Les haches en « jades alpins » en Bulgarie. Axeheads of « Alpine jades » in Bulgaria. in *Les Cahiers de la MSHE Ledoux 17; Série Dynamiques Territoriales 1)* (ed. Pétrequin, A.-M.) 1231–1279 (Besançon, 2012).

318. Querré, G., Herbault, F. & Calligaro, T. Transport of Neolithic variscites demonstrated by PIXE analysis. *Xray Spectrom.* **37**, 116–120 (2008).

319. Real, F. Estudo mineralógico de adornos de cor verde do Neolítico antigo da Gruta do Caldeirão. *Lisboa: IPPAR (Trabalhos de Arqueologia)* **6**, 315–319 (1992).

320. Villalba, M. J., Edo, M. & Blasco, A. Explotación, manufactura, distribución y uso como bien de prestigio de la 'calaita' en el Neolítico: el ejemplo del complejo de Can Tintorer. in *Minerales y Metales en la Prehistoria Reciente* 41–70 (Valladolid, 1998).

321. Querré, G., Domínguez-Bella, S. & Cassen, S. La variscite ibérique. Exploitation, diffusion au cours du Néolithique. in *Roches et Sociétés de la Préhistoire. Entre Massifs cristallins et Bassins sédimentaires* (eds. Marchand, G. & Querré, G.) 307–315 (Presses Universitaires de Rennes, Rennes, 2012).

322. *Intervencions Arqueològiques a Les Mines de Gavà (Sector Serra de Les Ferreres). Anys 1998–2009. De La Variscita al Ferro: Neolític i Antiguitat*. (2009).

323. Morell-Rovira, B. *et al.* Cronología de las Minas neolíticas de Can Tintorer (Gavà, Barcelona): explotación minera y uso funerario. *Munibe Antropologia-Arkeologia* **74**, 39–54 (2023).

324. Gibaja Bao, J. F. Prácticas funerarias durante el Neolítico en Cataluña. *Mainake* **26**, 9–27 (2004).

325. Cassen, S. & Grimaud, V. *Signes Gravés, Signes Polis. Locmariaquer, Morbihan. Mane Er Hroech et Kerpenhir Dans Le Contexte Néolithique Des Rives Du Morbihan*. (Les Vaisseaux de Pierres, Carnac, 2026).

326. Tykot, R. H. Obsidian studies in the prehistoric central Mediterranean: After 50 years, what have we learned and what still needs to be done? *Open Archaeol.* **3**, (2017).

327. Manen, C. Spatial, chronological and cultural dynamics of the Neolithization in the Western Mediterranean. in *La Transition Néolithique en Méditerranée* (eds. Manen, C., Perrin, T. & Guilaine, J.) 405–418 (Errance, Paris, 2014).

328. Dolfini, A. From the Neolithic to the Bronze Age in central Italy: Settlement, burial, and social change at the dawn of metal production. *J. Archaeol. Res.* **28**, 503–556 (2020).

329. Freund, K. P. & Batist, Z. Sardinian obsidian circulation and early maritime navigation in the neolithic as shown through social network analysis. *J. Isl. Coast. Archaeol.* **9**, 364–380 (2014).

330. Terradas, X. *et al.* Neolithic diffusion of obsidian in the western Mediterranean: new data from Iberia. *J. Archaeol. Sci.* **41**, 69–78 (2014).

331. Léa, V. Diffusion of obsidian in the northwestern Mediterranean: Toward a new model of the Chassey culture? *J. Mediterr. Archaeol.* **25**, 147–173 (2012).

332. Bronk Ramsey, C. Dealing with outliers and offsets in radiocarbon dating. *Radiocarbon* **51**, 1023–1045 (2009).

333. Schulting, R., Lanting, J. & Reimer, P. New dates from Tumulus Saint-Michel, Carnac. in *Autour de la Table. Explorations archéologiques et discours savants sur des architectures néolithiques à Locmariaquer* (ed. Cassen, S.) 769–773 (Morbihan; Nantes, 2009).

334. Boujot, C. & Cassen, S. Tertres Armoricaines et Tumulus Carnacéens dans les contextes de la néolithisation de la France occidentale. in *Sépultures d'Occident et genèses des mégalithismes (9000-3500 avant notre ère)* (ed. Guilaine, J.) (Éditions Errance, Paris, 1998).

335. Fouquet, M. Rapport sur la découverte d’une grotte sépulcrale dans la butte de Tumiac, le 21 juillet 1853. *Bulletin de la Société Polymathique du Morbihan* **6**, 1–7 (1862).

336. Galles, R. Fouille du tumulus de Tumiac en Arzon. in *Imprimerie Galles* (Vannes, 1878).

337. Le Rouzic, Z. Tumulus de Tumiac, monument historique, commune d’Arzon (Morbihan). in *Restaurations faites dans la région* 5–17 (1934).

338. *La Bretagne et l’Europe Préhistoriques: Mémoire En Hommage à Pierre-Roland Giot*. (Association pour la diffusion des recherches archéologiques dans l’Ouest de la France, Rennes, 1990).

339. Cassen, S. Préhistoire récente entre Loire et Vilaine: tendances actuelles de l’enregistrement et nouvelles visées de l’interprétation. *Cahiers du Pays de Guérande* **44**, 4–14 (2004).

340. Gely, B. Nouvelles datations des restes humains néolithiques de la nécropole du Replat à Aime (Savoie). in *Bilan Scientifique de la région Rhône-Alpes 2003* 186 (DRAC Rhône-Alpes, Lyon, 2005).

341. Giot, P.-R. *La Bretagne Des Mégalithes*. (Éditions Ouest-France, Rennes, 2007).

342. Lecornec, J. Le complexe mégalithique du Petit-Mont à Arzon (Morbihan). *Rev. archéol. l Ouest* **4**, 37–56 (1987).

343. Lecornec, J. Le Petit Mont. in *Documents Archéologiques de l’Ouest* (Arzon, Morbihan. Rennes, 1994).

344. Giot, P.-R. *Barnenez, Carn, Guennoc*. (Université de Rennes, Rennes, 1987).

345. Sparfel, Y. & Pailler, Y. *Les Mégalithes de l'arrondissement de Brest*. (ICB, Rennes, 2009).

346. Delibrias, G., Guillier, M. & Labeyrie, J. Gif natural radiocarbon measurements V. *Radiocarbon* **12**, 421–444 (1970).

347. Mohen, J.-P. Les styles céramiques des Tumulus de Bougon, Deux-Sèvres. in *Le Néolithique de la France. Hommage à Gérard Bailloud* (eds. Guilaine, J. & Demoule, J.-P.) 207–216 (CNRS, Paris, 1986).

348. Mohen, J.-P. & Scarre, C. *Les Tumulus de Bougon. Complexe Mégalithique Du Ve Au IIIe Millénaire*. (Errance, Paris, 2002).

349. Billard, C., Guillon, M., Sunder, F. & Arbogast, R.-M. La sépulture collective néolithique de Val-de-Reuil, Les Varennes et ses structures associées de l'âge du Bronze. in *Actes du XXe colloque interrégional sur le Néolithique, Évreux 1993* (Presses Universitaires de Rennes, Rennes, 1995).

350. Chambon, P. & Salanova, L. Chronologie des sépultures du IIIe millénaire dans le bassin de la Seine. *Bull. Soc. préhist. fr.* **93**, 103–118 (1996).

351. Billard, C. Les allées sépulcrales: derniers caveaux funéraires collectifs. in *La Hache et la Meule* (eds. Marcigny, C., Ghesquière, E. & Desloges, J.) (Muséum du Havre, Le Havre, 2007).

352. *Défunts Néolithiques En Toulousain*. (Archives d'Ecologie Préhistorique / EHESS, Toulouse, 2008).

353. Zilhão, J. Gruta do Caldeirão. O Neolítico antigo. *Lisboa: Instituto Português do Património Arquitectónico e Arqueológico* **6**, (1992).

354. Silva, C. T. Octávio da Veiga Ferreira e o estudo do Megalitismo da Serra de Monchique e do Baixo Alentejo. *Estudos Arqueológicos de Oeiras* **16**, 301–315 (2008).

355. Gomes, L. F. C., Carvalho, P. M. S., Francisco, J. P. A., Perpétuo, J. A. & Marrafa, L. C. O dólmen de Areita (S. João da Pesqueira, Viseu). A Pré-História na Beira Interior. *Estudos Pré-Históricos* **6**, 33–94 (1998).

356. Leisner, V. & Ribeiro, L. Die Dolmen von Carapito. *Madrider Mitteilungen* **9**, 11–62 (1968).

357. Soares, J. & Silva, C. T. *Anta Grande Do Zambujeiro: Arquitectura e Poder. Intervenção Arqueológica Do MAEDS*. vol. 3 83–129 (Musa, 2010).

358. Boaventura, R. Chronology of megalithism in south-central Portugal. *Menga. Revista de Prehistoria de Andalucía* **1**, 159–192 (2011).

359. Gonçalves, J. L. M. Olelas e Pragança: duas fortificações calcolíticas da Estremadura. *O Arqueólogo Português* **8–10**, 31–40 (1990-1992).

360. Odriozola, C. P. *et al.* Las producciones de adorno personal en rocas verdes del SW peninsular: los casos de Leceia. *Estudos Arqueológicos de Oeiras* **20**, 605–622 (2013).

361. Sanches, M. J. *O Abrigo Do Buraco Da Pala (Mirandela) No Contexto Da Pré-História Recente de Trás-Os*. (1997).

362. Querré, G. & Cassen, S. Callaïs. La parure en callaïs (variscite, turquoise) du Néolithique européen. *Actes du colloque de Carnac* (2015).

363. Odriozola, C. P., Linares-Catela, J. A. & Hurtado-Pérez, V. Variscite source and source analysis: testing assumptions at Pico Centeno (Encinasola, Spain). *J. Archaeol. Sci.* **37**, 3146–3157 (2010).

364. Maxia, C. & Atzeni, E. La necropoli eneolitica di S. Benedetto di Iglesias. in *Atti della VIII e IX Riunione Scientifica. Trieste, 19-20 ottobre 1963; Calabria, 6-8 aprile 1964* (Istituto Italiano di Preistoria e Protostoria, Firenze, 1964).

365. Floris, R. I resti scheletrici della Tomba II della necropoli preistorica di San Benedetto-Iglesias. in *La collezione Pistis-Corsi ed il patrimonio archeologico del comune di Iglesias. Iglesias: Comune di Iglesias* (eds. Atzeni, E., Alba, L. & Canino, G.) 30–31 (2001).

366. Atzeni, E. La necropoli di cultura “Ozieri” a San Benedetto di Iglesias (CA). in *La collezione Pistis-Corsi ed il patrimonio archeologico del Comune di Iglesias* (eds. Atzeni, E., Alba, L. & Canino, G.) 25–29 (Comune di Iglesias, Iglesias, 2001).

367. Lai, L. Il clima nella Sardegna preistorica e protostorica: problemi e nuove prospettive. in *Istituto Italiano di Preistoria e Protostoria* (eds. Preistoria E La Protostoria Della Sardegna, L. & Della Xliv Riunione Scientifica Dellistituto Italiano Di Preistoria E Protostoria, A.) vol. I 313–324 (Firenze, 2009).

368. Germanà, F. *L’uomo in Sardegna Dal Paleolitico All’età Nuragica*. (Carlo Delfino, Sassari, 1995).

369. Silvester, C. M. & Hillson, S. A critical assessment of the potential for Structure-from-Motion photogrammetry to produce high fidelity 3D dental models. *Am. J. Phys. Anthropol.* **173**, 381–392 (2020).

370. Cantín, M., Muñoz, M. & Olate, S. Generation of 3D tooth models based on three-dimensional scanning to study the morphology of permanent teeth. *Int. J. Morphol.* **33**, 782–787 (2015).

371. Nobles, G., Çakirlar, C. & Svetachov, P. Bonify 1.0: evaluating virtual reference collections in teaching and research. *Archaeol. Anthropol. Sci.* **11**, 5705–5716 (2019).

372. Batallés, M., Varela Richard, S. T. L., Fariña, A. & Costoya, G. *Megafauna 3D: Fossil Digitalization for Education, Outreach and Research*. (2018).

373. Bertsatos, A. & Chovalopoulou, M.-E. Advances in osteometric sorting: Utilizing diaphyseal CSG properties for lower limb skeletal pair-matching: Skeletal pair-matching with CSG properties. *J. Forensic Sci.* **65**, 1400–1405 (2020).

374. Alonso, A. G., Abadía, M. F. & González, R. G. Digital epigraphy:: New technologies and 3D modelling. in *Epigraphy in the Digital Age* 47–54 (Archaeopress Publishing Ltd, 2021).

375. Dell'Unto, N. & Landeschi, G. *Archaeological 3D GIS*. (Routledge, London, 2022).

376. Derudas, P. & Berggren, Å. Expanding field-archaeology education: The integration of 3D technology into archaeological training. *Open Archaeol.* **7**, 556–573 (2021).

377. Green, S. Structure from Motion. in *The Encyclopedia of Archaeological Sciences* (ed. Sandra, L. L.) 1–3 (John Wiley & Sons, Inc, Hoboken, NJ, USA, 2018).

378. Green, S., Bevan, A. & Shapland, M. A comparative assessment of structure from motion methods for archaeological research. *J. Archaeol. Sci.* **46**, 173–181 (2014).

379. Horn, C., Peternell, M., Ling, J., Green, A. & Potter, R. Rock art in three dimensions: Comments on the use and possibilities of 3D rock art documentation. in *The 3 Dimensions of Digitalised Archaeology* 87–108 (Springer International Publishing, Cham, 2024).

380. Potter, R., Rönnlund, R. & Wallensten, J. An evaluation of Substance Painter and Mari as visualisation methods using the Piraeus Lion and its runic inscriptions as a case study. *Herit. Sci.* **11**, (2023).

381. Morris, C., Peatfield, A. & O'Neill, B. 'Figures in 3D': Digital Perspectives on Cretan Bronze Age Figurines. *Open Archaeol.* **4**, 50–61 (2018).

382. Pinotti, T., McColl, H., Sikora, M. & Czech, L. ybyra: Y-chromosome haplogroup calling using a tree-based scoring method. *bioRxiv* (2025) doi:10.1101/2025.11.20.689455.

383. Atzeni, E. La preistoria del Sulcis-Iglesiente. in *AA.VV., Iglesias - Storia e società, Iglesias: Rotary club* 7–57 (1987).

384. Atzeni, E. Documenti per la preistoria di Iglesias. in *AA.VV., Iglesias. Storia e società, Iglesias: Rotary Club Iglesias* 9–20 (1978).

385. Bronk Ramsey, C. *OxCal v4.4.4*. (2021).

386. Guilbeau, D. *Les Grandes Lame et Les Lames Par Pression Au Levier Du Néolithique et de l'Éneolithique En Italie*. (2010).

387. Marcus, J. H. *et al.* Genetic history from the Middle Neolithic to present on the Mediterranean island of Sardinia. *Nat. Commun.* **11**, 939 (2020).

388. Olivieri, A. *et al.* Mitogenome diversity in Sardinians: A genetic window onto an island's past. *Mol. Biol. Evol.* **34**, 1230–1239 (2017).

389. Dehn, W., Kalb, P. & Vortisch, W. Geologisch-Petrographische Untersuchungen an Megalithgräbern Portugals. *Madr. Mitt.* 1–28 (1991).

390. Costeira, C., Porfírio, E., Cardoso, J. L., Costa, A. & Simões, T. Monumento Pré-histórico da Praia das Maçãs - Leitura comparada dos dados da escavação arqueológica realizada em 1961 e das intervenções arqueológicas de 2020–2022. **33**, 253–320 (2023).

391. Costeira, C. *et al.* De Gibraltar aos Pirinéus - megalitismo, vida e morte na fachada atlântica peninsular: arqueologia. *Fabião* **25**, 125–155 (2024).

392. Costeira, C., Porfírio, E. & Melo, L. A Câmara Ocidental do Monumento Pré-histórico da Praia das Maçãs (Sintra): Contexto arqueológico, estudo antropológico e datação absoluta dos enterramentos humanos identificados em 2022. *Munibe Monogr. Anthropol. Archaeol. Ser.* **3**, (2025).

393. Gonçalves, J. L. M. O monumento pré-histórico da Praia das Maçãs: Arquitectura e cerâmica pré-campaniforme. *Boletim Cultural da Assembleia Distrital de Lisboa. Lisboa, 3ª Série* **85**, 125–135 (1979).

394. Gonçalves, J. L. M. *Monumento Pré-Histórico Da Praia Das Maçãs (Sintra). Notícia Preliminar. Sintria, I-II*. vol. I–II 29–58 (Sintria, Sintra, 1982–1983).

395. Jordão, P. & Mendes, P. As grutas artificiais da Estremadura Portuguesa: uma leitura crítica das arquiteturas. *Arqueologia e História* **58**, 43–78 (2007).

396. Leisner, V. *Die Megalithgräber Der Iberischen Halbinsel. Der Westen. Madrider Forschungen, 1/3. Berlim*. (Walter de Gruyter & C.°, 1965).

397. Leisner, V. & Ferreira, O. V. Primeiras datas de radiocarbono 14 para a cultura megalítica portuguesa. *Revista de Guimarães* **73**, 358–366 (1963).

398. Leisner, V., Zbyszewski, G. & Ferreira, O. V. *Les Monuments Préhistoriques de Praia Das Maçãs et de Casainhos*. vol. 16 (Memória, nova Série, 1969).

399. Machado, L. S. *Sepultura Pré-Histórica Da Praia Das Maçãs. O Arqueólogo Português*. vol. 27 (Lisboa, 1929).

400. Monteagudo, L. Versuch einer chronologischen gliederung der portugiesischen kupferzeit. *Madr. Mitt.* 61–78 (1966).

401. Silva, A. M. Os ossos humanos "esquecidos" da Praia das Maçãs: análise antropológica da amostra óssea do museu arqueológico de São Miguel de Odrinhas. *Conimbriga* **46**, 5–26 (2007).

402. Cardoso, J. & Monge Soares, A. M. Sobre a cronologia absoluta das grutas artificiais da Estremadura portuguesa. *Al-Madan. Série II* 10–13 (1995).

403. Serrão, E. C. As jazidas arqueológicas de Catrivana e o dolmen de «Pedra Erguida». Sintria. Sintra: Museu Arqueológico de São Miguel de Odrinhas. 11–28 (1982).

404. Zbyszewski, G. *et al.* Le monument de "Pedras da Granja" ou de "Pedras Altas" dans la "Várzea de Sintra." **3**, 197–239 (1977).

405. Ferreira, O. V. Inventário dos monumentos megalíticos dos arredores de Lisboa. in *Actas e Memórias do 1º Congresso Nacional de Arqueologia* vol. 1 215–233 (Lisboa, Lisboa, 1958).

406. Carvalho, A. F., Gonçalves, D., Díaz-Zorita Bonilla, M. & Valente, M. J. Multi-isotope approaches to the Neolithic cemetery-cave of Bom Santo (Lisbon): new data and comparisons with fourth millennium BC populations from central–southern Portugal. *Archaeol. Anthropol. Sci.* **11**, 6141–6159 (2019).

407. Carvalho, A. F. *Bom Santo Cave (Lisbon) and the Middle Neolithic Societies of Southern Portugal*. vol. 17 (Promontoria Monográfica, Faro, 2014).

408. Carvalho, A. F. *et al.* The Bom Santo Cave (Lisbon, Portugal): Catchment, diet, and patterns of mobility of a middle Neolithic population. *Eur. J. Archaeol.* **19**, 187–214 (2016).

409. Leisner, V. & Ferreira, O. V. Os monumentos megalíticos de Trigache e de A-de-Beja. in *Actas e Memórias do I Congresso Nacional de Arqueologia, Lisboa, 15-20 Dezembro de 1958* vol. 1 187–233 (Lisboa, 1959).

410. Leisner, V. & Ferreira, O. V. Monumentos megalíticos de Trigache e de A-da-Beja II: Monumentos megalíticos. *Comunicações dos Serviços Geológicos de Portugal* **40**, 300–337 (1961).

411. Ribeiro, F. C. *Estudos Prehistoricos Em Portugal. II: Monumentos Megalithicos Das Visinhanças de Bellas*. (Typographia da Academia, Lisboa, 1880).

412. Ribeiro, C. Les kioekkenmoeddings de la Vallée du Tage. in *Compte Rendu de la IX session du Congrès International d'Anthropologie et d'Archéologie Préhistoriques, Lisbonne 1880* 279–290 (Lisboa, 1884).

413. Peyroteo Stjerna, R. On Death in the Mesolithic: Or the Mortuary Practices of the Last Hunter-Gatherers of the South-Western Iberian Peninsula, 7th--6th Millennium BCE. (Uppsala University, Uppsala, 2016).

414. Peyroteo Stjerna, R. Chronology of the Burial Activity of the Last Hunter-Gatherers in the Southwestern Iberian Peninsula, Portugal. *Radiocarbon* **63**, 265–299 (2021).

415. Cunha, E. & Cardoso, F. New data on Muge shell middens: A contribution to more accurate numbers and dates. *Estudos Arqueológicos de Muge* **I**, 171–184 (2003).

416. Meiklejohn, C., Roksandic, M., Jackes, M. & Lubell, D. Radiocarbon dating of Mesolithic human remains in Portugal. *Mesolithic Miscellany* **20**, 4–16 (2009).

417. Seiler, M. *et al.* Status report of the Trondheim Radiocarbon Laboratory. *Radiocarbon* **61**, 1963–1972 (2019).

418. Longin, R. New method of collagen extraction for radiocarbon dating. *Nature* **230**, 241–242 (1971).

419. Nadeau, M.-J. *et al.* Status of the "new" AMS facility in Trondheim. *Nuclear Instruments and Methods in Physics Research Section B: Beam Interactions with Materials and Atoms* **361**, 149–155 (2015).

420. López-Merino, L., López-Sáez, J. A., Zapata, M. B. R. & García, M. J. G. Reconstructing the history of beech (Fagus sylvatica L.) in the north-western Iberian

Range (Spain): From Late-Glacial refugia to the Holocene anthropic-induced forests. *Rev. Palaeobot. Palynol.* **152**, 58–65 (2008).

421. van der Horst, A. *et al.* Late-glacial and Holocene shifts in the mountain landscapes of the Cantabrian range (northern Spain) in response to changing climate, fire occurrence and land use. *Quat. Sci. Rev.* **342**, 108899 (2024).

422. Rius, D., Vannière, B. & Galop, D. Holocene history of fire, vegetation and land use from the central Pyrenees (France). *Quat. Res.* **77**, 54–64 (2012).

423. Zapolska, A. *et al.* More than agriculture: Analysing time-cumulative human impact on European land-cover of second half of the Holocene. *Quat. Sci. Rev.* **314**, 108227 (2023).

424. Reille, M. & Andrieu, V. The late Pleistocene and Holocene in the Lourdes Basin, Western Pyrénées, France: new pollen analytical and chronological data. *Veg. Hist. Archaeobot.* **4**, (1995).

425. Cyprien, A.-L., Visset, L. & Carcaud, N. Evolution of vegetation landscapes during the Holocene in the central and downstream Loire basin (Western France). *Veg. Hist. Archaeobot.* **13**, (2004).

426. Joly, C. & Visset, L. Evolution of vegetation landscapes since the Late Mesolithic on the French West Atlantic coast. *Rev. Palaeobot. Palynol.* **154**, 124–179 (2009).

427. Gómez-Orellana, L., Ramil-Rego, P., Ferreiro da Costa, J. & Muñoz Sobrino, C. Holocene environmental change on the Atlantic coast of NW Iberia as inferred from the Ponzos wetland sequence. *Boreas* **50**, 1131–1145 (2021).

428. Regnauld, H., Jennings, S., Delaney, C. & Lemasson, L. Holocene sea-level variations and geomorphological response: An example from northern Brittany (France). *Quat. Sci. Rev.* **15**, 781–787 (1996).

429. Goslin, J. *et al.* A new Holocene relative sea-level curve for western Brittany (France): Insights on isostatic dynamics along the Atlantic coasts of north-western Europe. *Quat. Sci. Rev.* **129**, 341–365 (2015).

430. García-Artola, A. *et al.* Holocene sea-level database from the Atlantic coast of Europe. *Quat. Sci. Rev.* **196**, 177–192 (2018).

431. Bernal-Wormull, J. L. *et al.* New insights into the climate of northern Iberia during the Younger Dryas and Holocene: The Mendukilo multi-speleothem record. *Quat. Sci. Rev.* **305**, 108006 (2023).

432. Pena, L. D. *et al.* Climate fluctuations during the Holocene in NW Iberia: High and low latitude linkages. *Cont. Shelf Res.* **30**, 1487–1496 (2010).

433. deMenocal, P., Ortiz, J., Guilderson, T. & Sarnthein, M. Coherent high- and low-latitude climate variability during the holocene warm period. *Science* **288**, 2198–2202 (2000).

434. Ilvonen, L. *et al.* Spatial and temporal patterns of Holocene precipitation change in the Iberian Peninsula. *Boreas* **51**, 776–792 (2022).

435. García-Amorena, I. *et al.* CO2 radiative forcing during the Holocene Thermal Maximum revealed by stomatal frequency of Iberian oak leaves. *Biogeosciences Discussions* **5**, 3945–3964 (2008).

436. Thompson, A. J., Zhu, J., Poulsen, C. J., Tierney, J. E. & Skinner, C. B. Northern Hemisphere vegetation change drives a Holocene thermal maximum. *Sci. Adv.* **8**, eabj6535 (2022).

437. Davis, B. A. S., Brewer, S., Stevenson, A. C. & Guiot, J. The temperature of Europe during the Holocene reconstructed from pollen data. *Quat. Sci. Rev.* **22**, 1701–1716 (2003).

438. Seppä, H. & Birks, H. J. B. July mean temperature and annual precipitation trends during the Holocene in the Fennoscandian tree-line area: pollen-based climate reconstructions, The Holocene. **11**, 527–539 (2001).

439. Korhola, A., Weckström, J., Holmstrom, L. & Erasto, P. A quantitative Holocene climatic record from diatoms in northern Fennoscandia. *Quat. Res.* **54**, 284–294 (2000).

440. Seppä, H., Nyman, M., Korhola, A. & Weckström, J. Changes of treelines and alpine vegetation in relation to post-glacial climate dynamics in northern Fennoscandia based on pollen and chironomid records: HOLOCENE CLIMATE-VEGETATION DYNAMICS IN NORTHERN FENNOSCANDIA. *J. Quat. Sci.* **17**, 287–301 (2002).

441. Nesje, A., Matthews, J. A., Dahl, S. O., Berrisford, M. S. & Andersson, C. Holocene glacier fluctuations of Flatebreen and winter precipitation changes in the Jostedalsbreen region, western Norway, based on glaciolacustrine sediment records. *The Holocene* **11**, 267–280 (2001).

442. Renssen, H., Seppä, H., Crosta, X., Goosse, H. & Roche, D. M. Global characterization of the Holocene thermal maximum. *Quat. Sci. Rev.* **48**, 7–19 (2012).

443. Jalut, G., Esteban Amat, A., Bonnet, L., Gauquelin, T. & Fontugne, M. Holocene climatic changes in the Western Mediterranean, from south-east France to south-east Spain. *Palaeogeogr. Palaeoclimatol. Palaeoecol.* **160**, 255–290 (2000).

444. Colombaroli, D. *et al.* Response of broadleaved evergreen Mediterranean forest vegetation to fire disturbance during the Holocene: insights from the peri-Adriatic region. *Journal of Biogeography* **36**, 314–326 (2009).

445. Combourieu-Nebout, N. *et al.* Holocene vegetation and climate changes in the central Mediterranean inferred from a high-resolution marine pollen record (Adriatic Sea). *Clim. Past* **9**, 2023–2042 (2013).

446. Renssen, H. *et al.* The spatial and temporal complexity of the Holocene thermal maximum. *Nat. Geosci.* **2**, 411–414 (2009).

447. Baldini, L. M. *et al.* North Iberian temperature and rainfall seasonality over the Younger Dryas and Holocene. *Quat. Sci. Rev.* **226**, 105998 (2019).

448. Van Vliet-Lanoë, B. *et al.* Middle- to late-Holocene storminess in Brittany (NW France): Part II – The chronology of events and climate forcing. *The Holocene* **24**, 434–453 (2014).

449. Fernane, A. *et al.* Climate variability and storm impacts as major drivers for human coastal marsh withdrawal over the Neolithic period (Southern Brittany, NW France). *Palaeogeogr. Palaeoclimatol. Palaeoecol.* **435**, 136–144 (2015).

450. Eynaud, F. *et al.* Holocene climate dynamics on the European scale: Insights from a coastal archaeological record from the temperate Bay of Biscay (SW France). *Quat. Int.* **613**, 46–60 (2022).

451. Finné, M., Woodbridge, J., Labuhn, I. & Roberts, C. N. Holocene hydro-climatic variability in the Mediterranean: A synthetic multi-proxy reconstruction. *Holocene* **29**, 847–863 (2019).

452. Fyfe, R. M. *et al.* Prehistoric palaeodemographics and regional land cover change in eastern Iberia. *The Holocene* **29**, 799–815 (2019).
